# Rebalancing the motor circuit restores movement in a *Caenorhabditis elegans* model for TDP-43-toxicity

**DOI:** 10.1101/2023.10.24.563563

**Authors:** Mandy Koopman, Lale Güngördü, Leen Janssen, Renée I. Seinstra, Janet E. Richmond, Nathan Okerlund, René Wardenaar, Priota Islam, Andre E.X. Brown, Erik M. Jorgensen, Ellen A.A. Nollen

## Abstract

Amyotrophic lateral sclerosis (ALS) and frontotemporal dementia are caused by the abnormal accumulation of TAR DNA-binding protein 43 (TDP-43) in the cytoplasm of neurons. How TDP-43 accumulation leads to disease symptoms is not well-characterized. Here, we use a *C. elegans* model for TDP-43-induced toxicity to identify the biological mechanisms that lead to disease-related phenotypes. By applying deep behavioral phenotyping, we established a phenotypic fingerprint of TDP-43 worms. This fingerprint was compared to that of 294 *C. elegans* mutants, in which genes were mutated that are important for nervous system and muscle functioning. By using a computational clustering approach, we found that the release of acetylcholine and GABA was the primary defect in TDP-43 worms. We then functionally dissected the neuromuscular circuit to show that GABA transmission was more severely diminished compared to acetylcholine. Whereas the loss of GABA transmission was caused by a profound loss of GABA synapses, acetylcholine neurons appeared to be functionally silenced. Enhancing functional output of repressed acetylcholine neurons at the level of G-protein coupled receptors or through optogenetic stimulation restored neurotransmission, but inefficiently rescued locomotion. Surprisingly, rebalancing the excitatory and inhibitory input by simultaneous stimulation of GABA and acetylcholine input into muscles not only synergized the effects of boosting individual neurotransmitter systems, but instantaneously improved movement. Our results suggest that interventions accounting for the altered connectome may be more efficient in restoring motor function than those solely focusing on diseased neuron populations.

## Introduction

Multiple age-related neurodegenerative diseases, including amyotrophic lateral sclerosis (ALS), and frontotemporal dementia (FTD), are characterized by brain inclusions containing aggregated TAR DNA binding-protein 43 (TDP-43)^1–7^. TDP-43 inclusions are found in neurons in 97% of patients with ALS^8, 9^. Since missense mutations in TDP-43 explain only a small fraction of disease cases, a variety of insults seem to converge on TDP-43 aggregation as the cause of neurodegeneration^10^. The concomitant loss of TDP-43 from the nucleus and enrichment in the cytosol seems to be central to toxicity^1,2,11,12^. The altered localization of TDP-43 has been shown to induce several pathogenic processes within neurons, including glutamate-induced excitotoxicity, the dysregulation of RNA metabolism, impaired protein degradation and mitochondrial dysfunction^13–19^. How TDP-43 toxicity ultimately leads to functional and behavioral decline in patients remains to be elucidated.

Most research on TDP-43 diseases has centered around cell-intrinsic factors that underlie neuronal vulnerability and cell-autonomous mechanisms of degeneration. However, neuronal function, vulnerability and their consequences for behavior are not a sole reflection of cell-intrinsic effects, but also depend on connected synaptic networks^20, 21^. Emerging evidence reveals that pathogenesis of neurodegenerative diseases is related to the widespread and progressive changes in brain networking, including altered patterns of connectivity and transmission in brain and motor systems^22–24^. As part of a network, neurons are constantly receiving, processing and transmitting signals and homeostatically adjusting to this information stream^25^. Consequently, therapeutic interventions that aim to restore functional properties of diseased neurons may have unexpected effects in these adapted, or perhaps maladapted, networks. Thus, elucidating how damage to individual neurons affects connected networks may help to identify novel therapeutic targets.

Using rodent models to unravel the role of diseased neurons on networks is exceedingly difficult due to the complexity of their nervous system. However, the nematode C*aenorhabditis elegans*, which has a nervous system of only 302 cells, a fully established connectome, and comparable neurotransmitter systems to humans, could provide us with a simplified model for a systemic approach^26–33^. Behavioral and phenotypic readouts, such as posture and locomotion, are effective predictors of molecular and neuronal processes in *C. elegans*^34, 35^. Moreover, pan-neuronal overexpression of human TDP-43 in *C. elegans* has been shown to govern cellular toxicity, induce paralysis, and capture molecular hallmarks similar to ALS patients^36–43^.

Therefore, we used computational behavioral profiling to identify biological processes underlying movement dysfunction in a *C. elegans* model for TDP-43-induced toxicity. Comparing the phenotypic profiles to those of known neuronal mutants, we identified acetylcholine and GABA transmission as dominant biological processes responsible for hTDP-43 phenotypes. We then characterized the anatomy and electrophysiology of these neurotransmitter systems. Our findings suggest that an excitation-inhibition imbalance in this motor circuit underlies TDP-43-induced paralysis. Our results indicate that interventions that rebalance circuits, rather than improve individual neurotransmitter systems, may restore behavioral functions in neurodegenerative diseases.

## Results

### Behavioral profiling implicates GABA and acetylcholine transmission in the hTDP-43 phenotype

We and others have previously shown that pan-neuronal expression of human *TDP-43* in *C. elegans* (hereafter ‘hTDP-43 worms’) reproduces characteristic cellular features of ALS^36–43^. Expression of hTDP-43 causes severe movement defects, including swimming and crawling deficits and early paralysis^36–43^. Both the molecular features (TDP-43 levels) and motor impairments are enhanced when the growing temperature of hTDP-43 worms is shifted from 20 °C to 25 °C ^40–43^. Detailed feature measurements can be used to characterize the underlying biological processes affected by a gene of interest and identify the neuronal basis for a behavioral phenotype^35^. To elucidate the biological basis of the phenotypes seen in hTDP-43 worms, we performed phenotypic profiling using the ‘Tierpsy 256’ dataset. ‘Tierpsy 256’ is a standardized set of 256 features describing morphology, path, posture and velocities of whole worms^34, 44^. First, we assessed whether the Tierpsy profile is sufficient to distinguish hTDP-43 worms raised at 20 °C vs 25 °C, and whether phenotypes were distinguishable from control worms. Although we could visually distinguish hTDP-43 worms raised at different temperatures, clustering was only moderately robust. However, the phenotypic profiles between the control and hTDP-43 worms were clearly distinguishable (**Figure 1A,B**; **Supplementary Table 1.1**)

**Figure 1:**
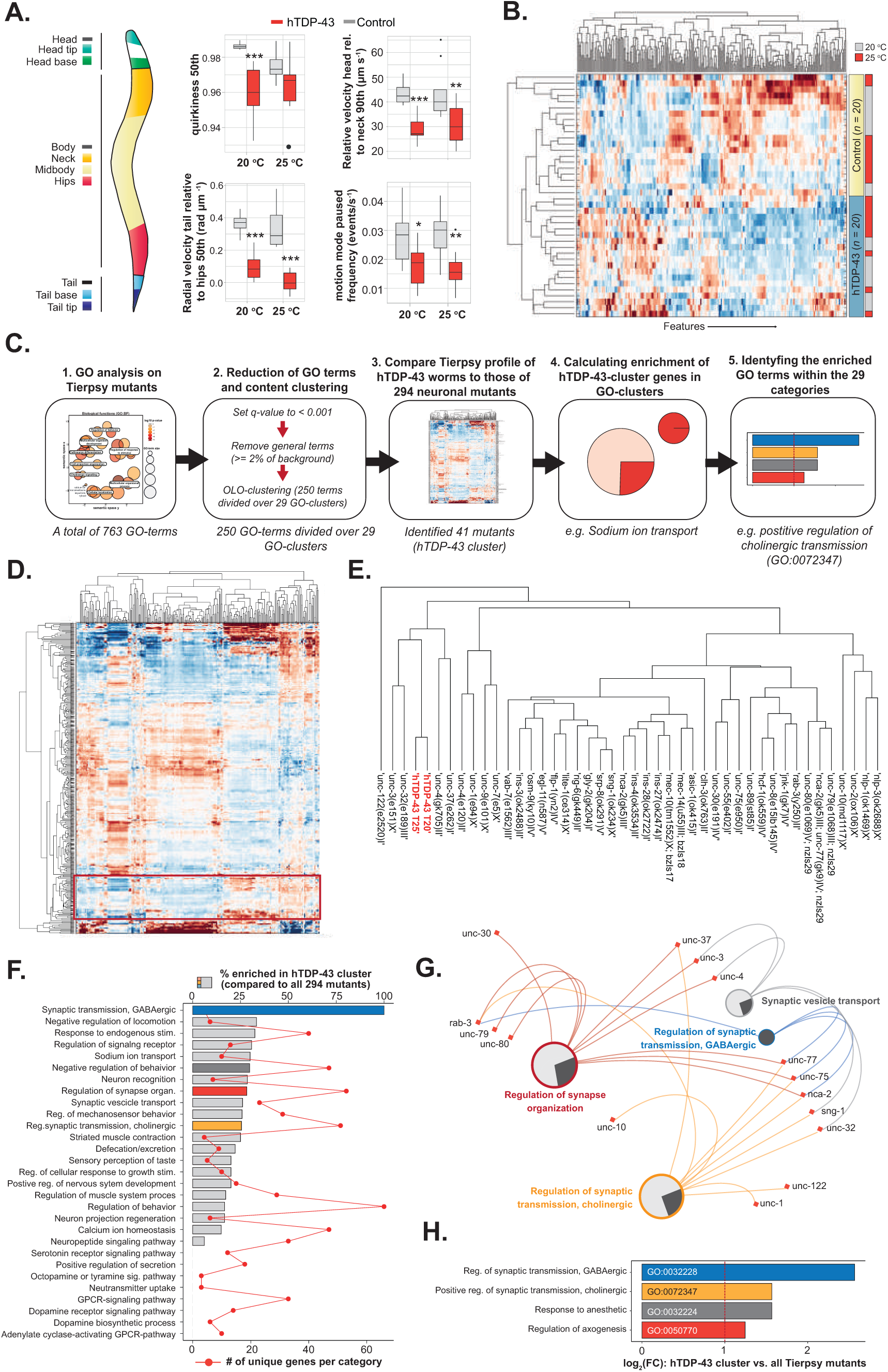
Phenomics screen identifies GABAergic and cholinergic transmission and neuronal excitability as processes underlying the phenotypic profile of hTDP-43 worms. **A)** schematic representation of worm segmentation used by ‘Tierpsy tracker’ to parameterize and quantify 256 features related to morphology, path, posture and velocities. Four examples of quantified features: quirkiness, a metric analogous of eccentricity (two-tailed unpaired Student’s t-test with Welch correction, 20 °C and 25 °C: p<0.001), radial velocity of the tail relative to the hips (50^th^ percentile) (two-tailed unpaired Student’s t-test with Welch correction, 20 °C: p<0.001, 25 °C: n.s.), motion mode paused frequency (two-tailed unpaired Student’s t-test, 20 °C: p=0.023, 25 °C: p =0.0018), and relative velocity head to neck (two-tailed unpaired Student’s t-test, 20 °C: p<0.001, 25 °C: p = 0.0037). **B)** Heatmap representing the ‘Tierpsy 256’ for hTDP-43 and control worms raised at 20 °C or 25 °C. **C)** A schematic representation of the analytical pipeline employed for the ‘phenomics’ approach OLO: Optimal leaf ordering. **D)** Heatmap comparing the average ‘Tierpsy 256’ profile of the hTDP-43 expressing worms at 20 °C and 25 °C to 294 established mutants and a control. The cluster with the hTDP43 strain and the most similar mutants is highlighted with the red square. **E)** Detail of the cluster highlighted in D. **F)** The GO-clusters, generated via ‘optimal leaf ordering (OLO)’ of the 250 unique GO terms associated with the 294 mutants. Gray bars show the percentual enrichment of genes in the hTDP-43 hits compared to the background of the 294 mutants. The colored bars indicate GO-clusters containing enriched GO-terms (see G and H). Red lines show the number of unique genes contained in each GO-cluster. **G)** A partial network showing the overlap of between four different GO-clusters. The circle-diagrams show the percentual enrichment (dark-gray) of the GO-cluster based on the hTDP-43 -hits. **H)** The most enriched GO-term per GO-cluster of the hTDP-43 hits that has at least one enriched term. The colors of the bars refer to the GO-cluster containing the GO terms (see F). * = p<0.05, ** = p<0.01, ** = p<0.001.

To determine the underlying biological processes affected by pan-neuronal hTDP-43 expression, we developed a computational pipeline to analyze behavioral features of known mutants to eventually compare to the Tierpsy profile of hTDP-43 worms (**Figure 1C-H**). The database comprises 9.203 short videos segmented to extract behavior and morphology features of *C. elegans* mutants, including the most canonical Unc (uncoordinated) and Egl (egg-laying mutants) as well as representative knockouts from receptor, channel and neuropeptide gene families^35^. We reanalyzed and summarized the movies of 294 neuronal gain- and loss-of-function mutants (our dataset hereafter referred to as ‘Tierpsy mutants’) according to the new ‘Tierpsy 256’ features (**Figure 1D,E**; **Supplementary Table 1.2**).

To segment these features into specific biological processes, we used a gene ontology (GO) enrichment analysis to categorize them (**Supplementary Table 1.3**; **Figure 1C**, 1; **Supplementary Table 1.4**). We identified 763 GO terms (biological function only), which were then reduced to 250 unique terms by eliminating general terms and selecting the most significant clusters (**Figure 1C**, 2; **Supplementary Table 1.5**). By means of optimal leaf ordering we ranked and grouped similar GO-terms with strongly overlapping gene content into 29 GO-clusters that represent the main biological processes in our screening dataset (**Figure 1F**; **Supplementary Table 1.5**). GO-clusters range from GABA signaling, to sodium ion transport, and muscle contraction (**Supplementary Table 1.5**).

**Figure 2:**
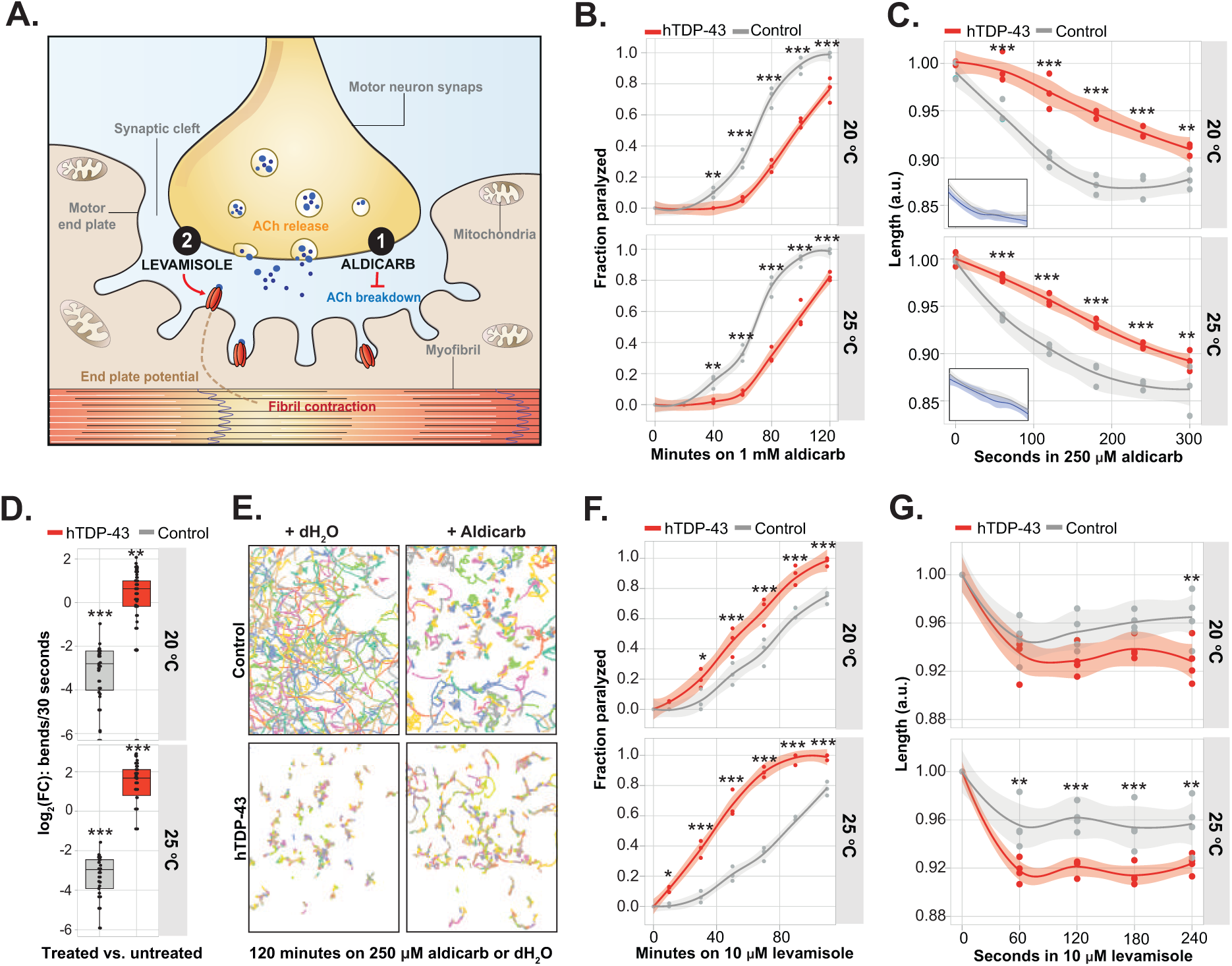
Cholinergic signaling is presynaptically impaired in worms expressing hTDP-43. **A)** Schematic of a cholinergic neuron and muscle fibrils. Levamisole stimulates post-synaptic acetylcholine receptors directly, while aldicarb inhibits acetylcholine breakdown by acetylcholine-esterases. **B)** Cholinergic neuronal transmission was measured by determining the onset of paralysis induced by the acetylcholine esterase inhibitor aldicarb. Two-way ANOVA for both 20 °C and 25 °C (treatment, genotype and interaction: p<0.001) with post-hoc Bonferroni multiple comparison tests: see asterisks. n =3 with 30-70 worms per experiment. **C)** Cholinergic neuronal transmission was measured by determining the decrease in body length induced by the acetylcholine esterase inhibitor aldicarb. Two-way ANOVA for both 20 °C and 25 °C (treatment, genotype and interaction: p<0.001) with post-hoc Bonferroni multiple comparison tests: see asterisks. n =3 with 20-60 worms per experiment. Small graphs show how the control worms (grey) compare to N2s (blue). **D)** Aldicarb normally induces paralysis of worms and a decline in movement speed or thrashing ability. The boxplots show the log_2_ fold-change of bends/30 seconds at the 120-minute interval compared to the 0-minutes timepoint for each strain. *n =* 50-80 worms per condition, Wilcoxon rank-test (p<0.011 for hTDP-43 at 20 °C, all other conditions: p<0.001). **E)** Representative crawling maps (10 minutes) of hTDP-43 and control worms grown at 25 °C. **F)** Postsynaptic sensitivity to acetylcholine was assessed with direct exposure to 10 μM levamisole. The onset of paralysis was assessed over time. Two-way ANOVA for 20 °C (treatment, genotype: p<0.001, interaction: p = 0.0019) and 25 °C (treatment, genotype and interaction: p<0.001) with post-hoc Bonferroni multiple comparison tests: see asterisks. n =3 with 30-70 worms per experiment. **G)** The decrease in body length upon 10 μM levamisole was assessed over time. Two-way ANOVA for 20 °C (treatment, genotype: p<0.001, interaction: n.s.) and 25 °C (treatment, genotype: p<0.001, interaction: p = 0.0155) with post-hoc Bonferroni multiple comparison tests: see asterisks. n=4 with 60-80 worms per experiment. Transparent areas in line-graphs represent 95% confidence interval. * = p<0.05, ** = p<0.01, ** = p<0.001.

We then compared the Tierpsy 256-signature of hTDP-43 worms to those of known neuronal mutants (**Figure 1C**, **3**; **Supplementary Table 1.2**) and identified 41 different mutants that closely resemble the hTDP-43-signature (**Figure 1D,E**; **Supplementary Table 1.2**). The hTDP-43 cluster includes *unc-30* and *unc-55*, which are genes involved in GABA neuron development. Moreover, the genes *unc-3*, *unc-4*, and *unc-37* are also included in the cluster; these transcription factors largely impact acetylcholine neurons. The gene *unc-10* encodes RIM which is a constituent of the active zone. Finally, it is notable that the cluster includes all of the genes encoding the homolog of the NALCN Na^+^ leak channel, including the pore-forming subunits *nca-1* (i.e. *unc-77*) and *nca-2* and the associated subunits *unc-79* and *unc-80.* Thus, from a geneticist point of view our phenomics approach suggests hTDP-43 disrupts GABA transmission, acetylcholine transmission as well as neuronal excitability.

**Figure 3:**
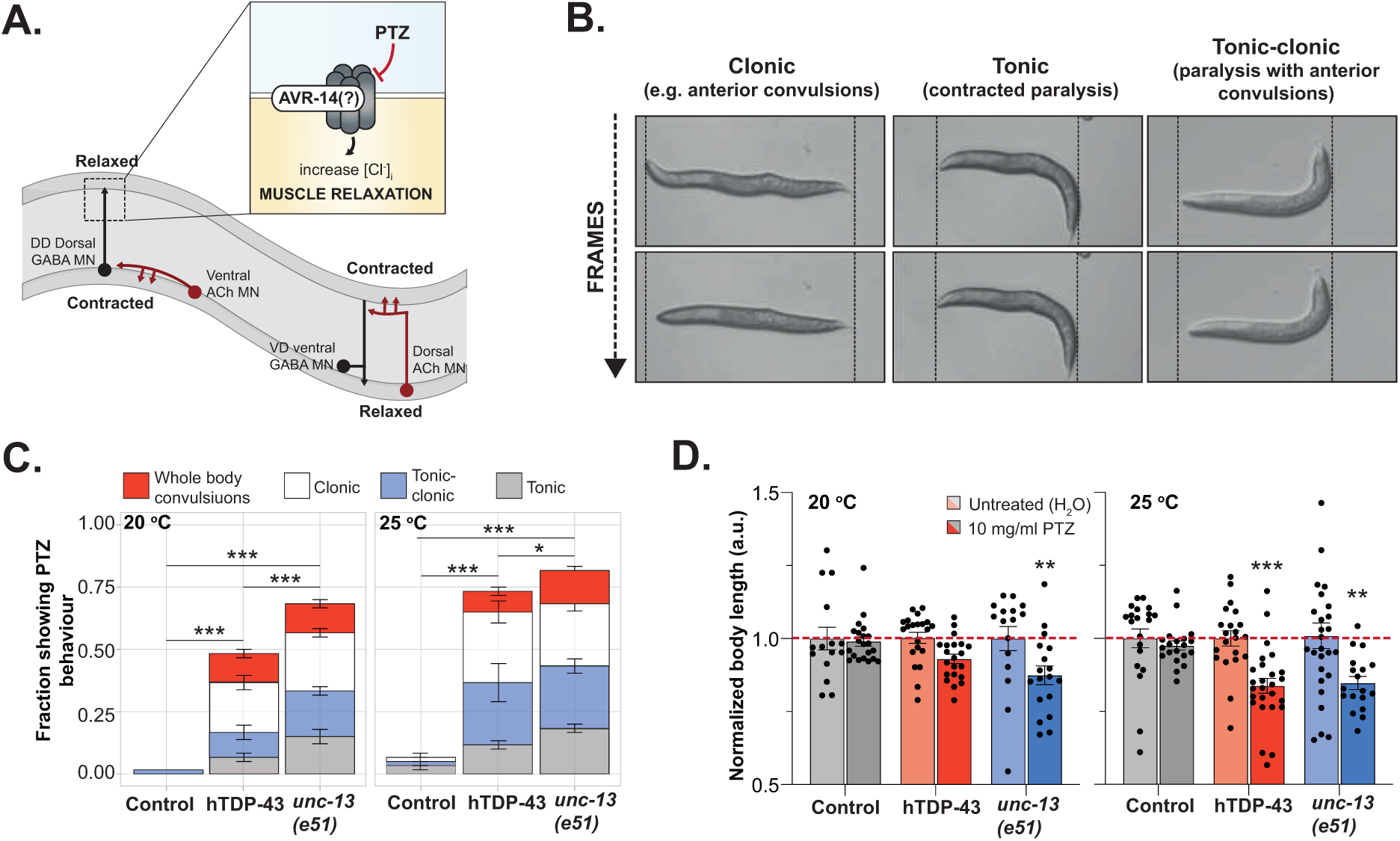
GABAergic signaling is impaired in hTDP-43 expressing animals. **A)** Schematic showing the connections between ventral cholinergic motoneurons and dorsal GABAergic neurons, and between dorsal cholinergic motoneurons and ventral GABAergic neurons. Dorsal cholinergic neurons excite the dorsal muscles and ventral cholinergic neurons excite the ventral muscles, while the connecting GABAergic neurons relax the muscles opposite to those undergoing contractions. PTZ binds likely the glutamate-gated chloride channel AVR-14 or the neurosteroid site of of UNC-49. **B)** Representative images of unc-13*(e51)* worms that show anterior convulsions (head bobs, tonic) or full body convulsions, contracted paralysis (shrinker, clonic), and tonic-clonic behaviour (head bobbing with a paralyzed body). **C)** The fraction of animals showing one of the aforementioned (see B) phenotypes after 30 minutes exposure to 10 mg/ml PTZ. *n* = 3 with 20 worms per experiment, one-way ANOVA for both 20 °C and 25 °C (p<0.001) with post-hoc Tukey. **D)** Normalized length of worms treated with 10 mg/ml PTZ for 30 minutes to their untreated controls. *n =3* with 20-26 worms per experiment, two-way ANOVA at 20 °C (treatment: p = 0.0020, strain and interaction: n.s.) and 25 °C (treatment: p<0.001, strain and interaction: n.s.) with post-hoc Sidak’s multiple comparison test, see asterisks. Error bars represent the S.E.M. * = p<0.05, ** = p<0.01, ** = p<0.001.

To determine which biological process was significantly enriched in the hTDP-43 cluster, we first investigated in which of the 29 GO-clusters the 41 identified genes were annotated. We found that the cluster surrounding hTDP-43 included all genes associated with the GO-cluster ‘synaptic transmission, GABAergic’ (**Figure 1C**, **4**-**5**; **Figure 1F-H**; **Supplementary Table 1.6; 1.7**). In contrast, genes involved in serotonergic or dopaminergic neurotransmission are absent within the hTDP-43 hits. Since the 29 GO-terms only give a global representation of the relationship between the more detailed GO-terms in our dataset and substantial overlap between the clusters exists (**Figure 1G**; **Supplementary Table 1.5**), we also looked specifically at the enrichment within the separate GO terms. We identified 13 GO terms in 4 of the GO-clusters that had at least a two-fold-change in enrichment (**Supplementary Table 1.7**). As expected, the enrichment of the GO-cluster ‘synaptic transmission, GABAergic’ is also evident in the clear enrichment of its underlying GO-terms (**Figure 1H**, GO enrichment (GOrilla): p = 6.78E-4). In addition, cholinergic signaling and processes related to synapse organization are also enriched and the hits in these categories overlapped (**Figure 1F-H**). In contrast to ‘synaptic transmission, GABAergic’ these GO-terms are strikingly influenced by the starting set of the genes (GO enrichment (GOrilla): n.s.) and merely based on chance. Thus, our computational phenomics approach identifies disrupted GABAergic signaling as predominant process responsible for the phenotypic profile of hTDP-43 worms. From a geneticist point-of-view altered cholinergic transmission and neuronal excitability invite a critical evaluation as well.

**Figure 4:**
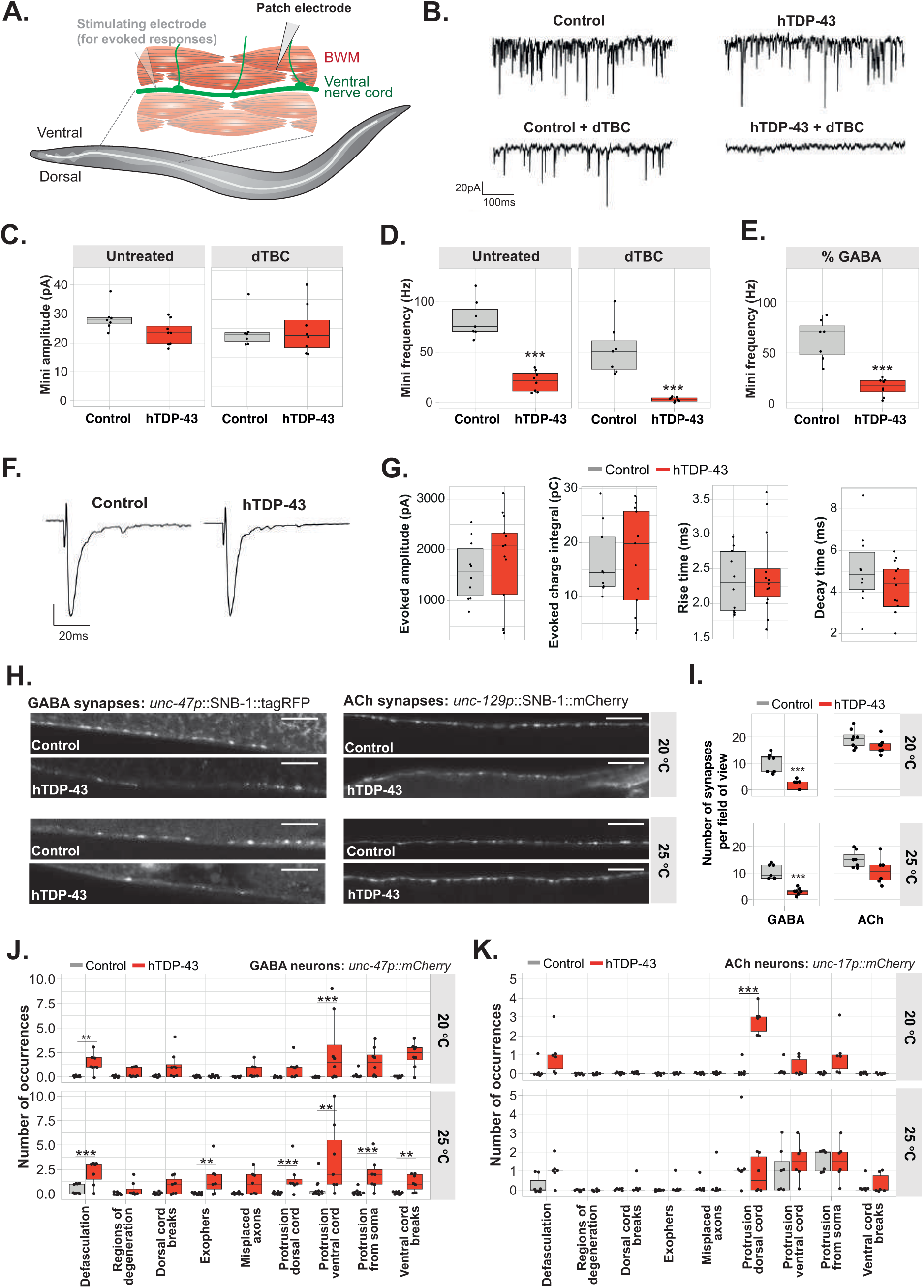
Degenerative and electrophysical phenotypes are especially existent in the GABAergic branch. **A)** Schematic representation of muscle patch-clamping in C. elegans. The transparent electrode left shows the location of a stimulation electrode for evoked responses. **B)** Representative traces of mPSCs before and after treatment with dTBC. **C)** Quantification of mEPSC amplitude before and after treatment with dTBC in control and hTDP-43 worms. n ≥ 6, Mann-Whitney U test, all: n.s. **D)** Quantification of mPSC frequency before and after dTBC treatment. n ≥ 6, test, Student’s t-test. **E)** Percentage of mPSCs remaining after dTBC treatment (indication of GABAergic minis), n ≥ 6, test, Student’s t-test. **F)** Representative traces of evoked EPSCs in hTDP-43 and control worms. **G)** Quantification of evoked amplitude, charge integral, rise time and decay time in hTDP-43 and control worms. n ≥ 10, two-way ANOVA (genotype, interaction: n.s., feature: p<0.001) with post-hoc Holm-Sidak. **H)** Representative images of control and hTDP-43 worms expressing SNB-1::mCherry or SNB-1::tagRFP in cholinergic or GABAergic synapses respectively. The dorsal cord was imaged. **I)** Quantification of the number of cholinergic and GABAergic synapses in hTDP-43 worms. n ≥ 7, Inter-strain Mann-Whitney U test were performed. Scale bar = 10 microns. **J)** Quantification of pre-established neurodegenerative phenotypes in worms expressing mCherry in GABAergic neurons. Inter-strain Mann-Whitney U-tests were performed per phenotype. **K)** Quantification of pre-established neurodegenerative phenotypes in worms expressing mCherry in cholinergic neurons. n ≥ 6, Inter-strain Mann-Whitney U-tests were performed per phenotype. FDR-corrections were applied (q<0.01) to correct for multiple comparisons. FDR-corrections were applied (q<0.01) to correct for multiple comparisons. * = p<0.05, ** = p<0.01, ** = p<0.001.

**Figure 5:**
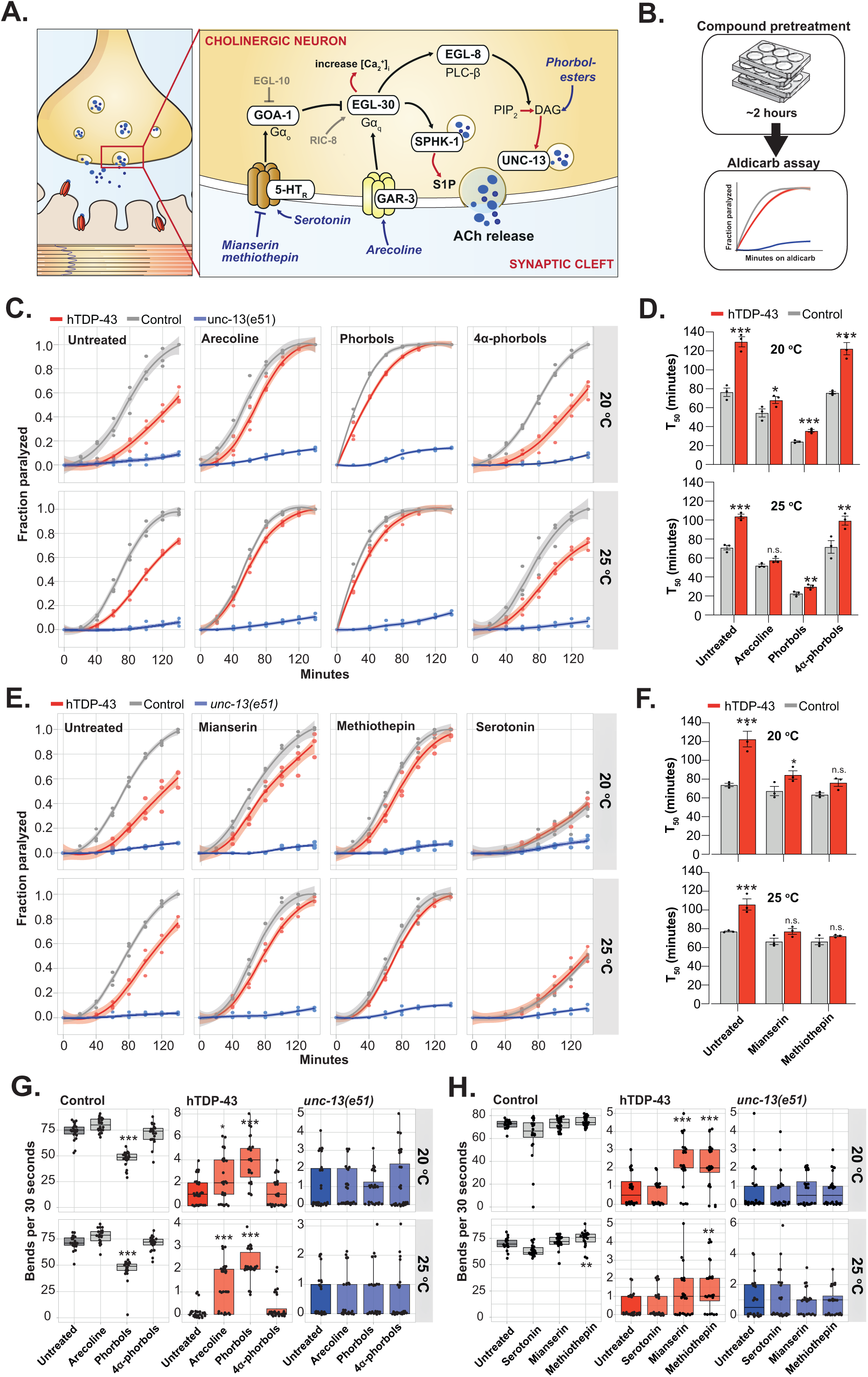
Enhanced acetylcholine-release through facilitation with phorbol-esters or by stimulating of G-protein-coupled-receptors indicate a functional synaptic machinery. **A)** Schematic showing the molecular pathways involved in cholinergic facilitation. Stimulation of GAR-3 and 5-HT receptors have opposite results in the facilitation of acetylcholine release. **B)** Schematic showing the experimental outline for the aldicarb-assays in D and G. **C)** Cholinergic neuronal transmission, after pretreatment with the depicted compounds, was measured by determining the onset of paralysis on 0.5 mM aldicarb plates. *n = 3*. Transparent areas show the S.E.M. **D)** The average time in which half of the worms are paralyzed on aldicarb in C is represented as the mean T_50_, *n =3*, two-way ANOVA on ln-transformed data at 20 °C (treatment, genotype: p<0.001, interaction: p = 0.043) and 25 °C (treatment, genotype: p<0.001, interaction: n.s.) with post-hoc Holm-Sidak’s. **E)** Cholinergic neuronal transmission, after pretreatment with the depicted compounds, was measured by determining the onset of paralysis on 0.5 mM aldicarb plates. *n = 3*. Transparent areas show the S.E.M. **F)** The average time in which half of the worms are paralyzed on aldicarb in E is represented as the mean T_50_, *n =3*, two-way ANOVA on ln-transformed data at 20 °C (treatment, genotype: p<0.001, interaction: p = 0.0191.) 25 °C (treatment, genotype: p<0.001, interaction: 0.0331) with post-hoc Holm-Sidak’s. **G)** Thrashing ability, in bends per 30 seconds, of worms pretreated with the depicted compounds. Intra-strain comparisons were performed only, *n* = 15-20, Kruskal-Wallis for control at 20 °C and 25 °C (p<0.001), for hTDP-43 at 20 °C and 25°C (p<0.001) and for unc-13(e51) 20 °C and 25 °C (n.s.) with post-hoc Dunn’s. One representative experiment of a triplicate is shown. **H)** Thrashing ability, in bends per 30 seconds, of worms pretreated with the depicted serotonin-related compounds. Only intra-strain comparisons were performed, *n* = 15-20, Kruskal-Wallis for control at 20 °C (p=0.0118), and 25 °C (p<0.001), for hTDP-43 at 20 °C (p<0.001) and 25°C (p = 0.0034) and for *unc-13*(e51) 20 °C and 25 °C (n.s.) with post-hoc Dunn’s. One representative experiment of a triplicate is shown. Error bars represent the S.E.M. and transparent areas in line-graphs represent 95% confidence interval. * = p<0.05, ** = p<0.01, ** = p<0.001.

### Presynaptic acetylcholine release is impaired in hTDP-43 worms and coexist with post-synaptic hyperreactivity

To validate conclusions from our phenomics study we performed pharmacological assays to investigate cholinergic and GABAergic transmission in the hTDP-43 worms. We first assessed the function of acetylcholine (ACh) neurons by measuring the sensitivity of animals to the acetylcholinesterase inhibitor aldicarb. Aldicarb enhances acetylcholine transmission by inhibiting the breakdown of the neurotransmitter^45^ (**Figure 2A**, **1**). Consequently, the accumulation of ACh at cholinergic synapses will lead to muscle hypercontraction, paralysis, and a decline in movement speed or thrashing ability. We found that hTDP-43 worms required more time to be paralyzed by aldicarb than wild-type control animals, which suggested that they are partially resistant to accumulating ACh (**Figure 2B**). However, since hTDP-43 worms are largely immobile to start with, using time-to-paralysis as a readout could skew the results. Therefore, we used body contraction in response to aldicarb as an additional readout^46^. Again, hTDP-43 worms took longer to respond to aldicarb in this assay, confirming their partial resistance (**Figure 2C**). Strikingly, although the movement of wild-type animals decreased when exposed to aldicarb, movement of the largely immobile hTDP-43 worms was increased up to 3-fold at 25 °C (**Figure 2D, E**). These results suggest that reduced ACh signaling contributes to the hTDP-43 induced motor impairment.

Aldicarb resistance can be caused by reduced ACh release from motor neurons (MNs) or reduced numbers of ACh receptors in the muscles. To determine if ACh receptors were reduced or absent, we used the nicotinic agonist levamisole to directly activate receptors on the body muscles^47^ (**Figure 2A**, 2). Surprisingly, levamisole paralyzed worms faster and the contraction was significantly stronger in hTDP-43 worms compared to the controls, and this effect was more pronounced at 25 °C (**Figure 2F, G**). The levamisole data suggests that the observed aldicarb-resistance of hTDP-43 worms is most likely underestimated due to compensatory increase in cholinergic sensitivity of the muscle. Thus, our data shows that a presynaptic cholinergic defect is coexistent with post-synaptic hyperreactivity in hTDP-43 worms.

### GABA signaling is impaired in hTDP-43 worms as assayed by pharmacology

Coordinated locomotor behavior in *C. elegans* requires both excitatory inputs from acetylcholine neurons and inhibitory signaling from GABA neurons at neuromuscular junctions (**Figure 2H**). Both aldicarb sensitivity and levamisole sensitivity are highly influenced by GABAergic signaling^48–50^. To investigate the effect of hTDP-43 expression on GABA signaling, worms were exposed to the non-specific chloride channel blocker, pentylenetetrazole (PTZ). PTZ exposure in the absence of GABA transmission causes a unique ‘head-bobbing’ phenotype^51^. The target of PTZ in strains lacking GABA transmission is not known. Nematodes lacking GABA transmission, including through loss of the muscle GABA-gated chloride channel UNC-49, still respond to PTZ. One of the targets may include the glutamate-gated chloride channel AVR-14, because it is sensitive to picrotoxin^52^. AVR-14 is expressed in the acetylcholine MNs, RMD and SMD, which innervate the head muscles. Disinhibition of these MNs may drive the spontaneous head-bobbing observed in GABA-deficient strains. We assayed the hTDP-43 worms for PTZ-sensitivity by assessing *a priori* defined convulsive phenotypes in unc-13(e51) mutant (**Figure 2I**)^53^. hTDP-43 worms showed strong convulsion phenotypes after a 30-minute exposure to PTZ at both 20 °C and 25 °C (**Figure 2J**). Additionally, hTDP-43 worms showed a shrinking phenotype similar to *unc-13(e51)* worms after 30-minute PTZ exposure (**Figure 2K**). Thus, GABAergic signaling is severely impaired in hTDP-43 worms. Taken together, these results show that both excitatory ACh and inhibitory GABA transmission are diminished in hTDP-43 worms.

### Frequency but not the amplitude of postsynaptic currents is reduced in hTDP-43 worms

To directly test if acetylcholine and GABA MNs exhibit defective neurotransmitter release in hTDP-43 worms, we used voltage-clamp electrophysiology on dissected worms to assess synaptic transmission at the neuromuscular junction (NMJ) (**Figure 3A**). First, we recorded the endogenous miniature postsynaptic currents (PSCs or ‘minis’), which reflect the basal activity of motor neurons at the presynapse, and subsequent postsynaptic responses to neurotransmitters^54^. To distinguish between cholinergic and GABAergic minis we recorded before and after perfusion of the cholinergic receptor antagonist d-tubocurarine (dTBC) (**Figure 3B**). Mini amplitudes were not significantly different between controls and hTDP-43 worms, in either untreated worms or after dTBC application suggesting that postsynaptic receptor fields were intact (**Figure 3C**). However, the mini frequency was significantly reduced in hTDP-43 worms, demonstrating that vesicle fusion at synapses was severely decreased (**Figure 3D**). Treatment by dTBC demonstrated that defect was primarily due to elimination of GABAergic events in hTDP-43 worms (**Figure 3E**). The sole cholinergic events (Δ =mini – mini_TBC_) were trending towards a significant decrease, even though the experiment was underpowered (Δ: 11.82 ± 6.545, p = 0.0941, *n* = 7. Post-hoc power: 0.38). The combination of a reduction in mini frequency with normal mini amplitudes suggests that the defect is in presynaptic release rather than postsynaptic reception or vesicle loading.

The continued presence of cholinergic endogenous events suggests that the total paralysis of hTDP-43 worms and aldicarb resistance could be indicative of altered volume transmission by neuropeptides or modulatory transmitters, which will not be apparent in dissected preparations, or presynaptic defects in evoked cholinergic release at the level of excitation/secretion. To further investigate the transduction of electrical stimuli (excitation) into synaptic vesicle fusion with the plasma membrane (secretion), we recorded the evoked excitatory currents (EPSCs) in hTDP-43 worms and their controls (**Figure 3F**). We found no differences in the evoked amplitude, charge transferred, rise or decay phase (**Figure 3G**). These results show that when the normal process of membrane depolarization is bypassed in dissected preparations, acetylcholine transmission in hTDP-43 worms is indistinguishable from control worms. Thus, a combination of pharmacology and electrophysiology reveals severely impaired GABA transmission and a reversible functional repression of acetylcholine neurons in hTDP-43 worms.

### GABA synapse numbers are severely reduced in hTDP-43 worms

The profound defect in GABA transmission could be due to a decrease in the rate of vesicle fusion at GABA synapses or a decrease in the number of GABA synapses. To assay synapses directly we expressed a tagged synaptic vesicle protein in GABA neurons (*Punc-47::*SNB-1::tagRFP) or in acetylcholine neurons (*Punc-129*::SNB-1::mCherry) (**Figure 3H**). Although the number of SNB-1 puncta in acetylcholine neurons appeared slightly lower in the hTDP-43 background, neither synapse size nor number were significantly reduced relative to control worms (**Figure 3H,I**; **Supplementary Figure 1A**). In contrast, in GABA neurons, SNB-1 puncta were almost completely absent in hTDP-43 worms, which was accompanied by a decrease in synaptic area for the remaining synapses (**Figure 3H,I**; **Supplementary Figure 1A**).

The lack of synaptic markers could be due to a specific loss of synapses along existing axons, or the retractions of axons themselves. To assay the morphology of the motor neurons, we crossed hTDP-43 worms with strains expressing mCherry in GABA neurons (*unc-47* promoter) or in acetylcholine neurons (*unc-17* promoter) (**Supplementary Figure 1B-E**). Animals were scored for structural abnormalities; mild phenotypes included defasciculation of the ventral cord, and protrusions from either the ventral or dorsal cord. Moderate defects included misdirected or branched axons, aberrant soma and exophers, or an occasional break in the dorsal or ventral nerve cords. Severe defects exhibit degeneration of the neurons, a phenotype which combines multiple defects, and often include breaks in the ventral nerve cord (**Figure 3J,K**, **Supplementary Figure 1B-E**).

For the most part, axons from GABA neurons were still present in the dorsal nerve cord, demonstrating that there is a specific loss of synapses from GABA neurons in hTDP-43 worms. However, GABA neurons exhibited neurodegenerative phenotypes present at both 20°C and 25°C, including defasciculation and protrusions from the ventral cord (**Figure 3J**). In addition, aberrant structures adjacent to cell bodies were present at 25°C; these are possibly herniated protrusions called exophers^55^. Although degeneration was significant, we did not observe motoneuron loss within the pool of 19 GABA motoneurons.

Despite the presence of some mild degenerative phenotypes, the morphology of the acetylcholine motoneurons appeared largely unaffected by the presence of hTDP-43 (**Figure 3K**, **Supplementary Figure 1B,D**). Since mCherry was expressed from an extrachromosomal array, some variability in the number of mCherry cell bodies was observed in both control and hTDP-43 animals, consistent with loss of the array. However, we did observe more frequent loss of mCherry expression in subsets of neurons in hTDP-43 animals grown at 25 °C. (**Supplementary Figure 1F**). Taken together, these data show that GABAergic motor neurons are more severely impacted in hTDP-43 worms and that the abolished GABA transmission is likely due to a decrease in the number of GABA neuromuscular junctions.

### Cholinergic facilitation enhances movement of hTDP-43 worms and indicates altered G-protein signaling

Next, we investigated the apparent contradiction of aldicarb resistance, suggesting a defect in ACh release, on the one hand and normal electrophysiological recordings at the other hand. One possibility is that modulatory pathways regulated by G-protein signaling are defective in hTDP-43 worms. Volume transmission by neuropeptides or modulatory transmitters would not be apparent in the dissected preparation used for electrophysiology. The Gα_q_-pathway can be activated upstream of Gα_q_ by the metabotropic acetylcholine receptor agonist arecoline^56, 57^; the Gα_q_-pathway can be activated downstream of G α _q_ by the DAG analog phorbol-12-myristate-13-acetate (phorbol ester)^58^ (**Figure 4A, B**). Arecoline and phorbol esters nearly restored aldicarb sensitivity in the hTDP-43 worms (**Figure 4C,D**) but had no effect on *unc-13* mutants which lack the DAG binding protein at synapses^33, 58–61^. Controls demonstrated the effect was not caused by off-target effects of phorbol ester or acted postsynaptically (**Supplementary Figure 2A**). These data suggest that hTDP-43 worms may have intact, but reduced Gα_q_ signaling.

The Gα_o_-pathway is known to antagonize the Gα_q_ -pathway in *C. elegans*^62, 63^. We found that inhibiting the Gα_o_ pathway could also rescue acetylcholine neurotransmission. Aldicarb sensitivity was rescued in hTDP-43 worms to wild-type levels when the G_o_-pathway was inhibited by the serotonin antagonists mianserin and metiothepin (**Figure 4E, F**)^64^. Conversely, the addition of serotonin caused a rightward shift in aldicarb sensitivity, collapsing the curves onto each other, which was stronger in control than hTDP-43 worms. Since the G_o_ pathway is strongly dependent on the G_q_ pathway to exert inhibitory effects on neurotransmitter release ^62, 63^, our data suggest that functional G_q_-signaling might be counteracted by high levels of G_o_-signaling in hTDP-43 worms. Thus, these data suggest that hTDP-43 primarily acts to decrease neurotransmission at neuromuscular junctions by interfering in G-protein signaling.

Finally, we asked whether the paralysis of hTDP-43 worms is entirely due to altered G-protein signaling. To test this possibility, we incubated hTDP-43 worms and their controls on plates containing Gα_q_-activating or Gα_o_-inhibiting compounds. While phorbol esters, arecoline, mianserin and methiothepin all greatly increase aldicarb sensitivity, their effects on locomotion were less striking (**Figure 4G,H**). Nevertheless, the almost completely immobile hTDP-43 worms became mobile on compounds that activate Gα_q_ signaling, with phorbol esters inducing the largest increase in mobility. In addition, activation of Gα_q_- and inhibition of Gα_o_ -signaling also increased coiling of the animals, which reduced normal forward locomotion (**Supplementary Figure 2B-D**). Taking together, these data suggest that both synaptic priming and fusion machinery are functional in cholinergic neurons of hTDP-43 worms, and suggest that alterations in the extent of upstream G-protein signaling might be involved in the reduced acetylcholine release, but do not fully account for the full paralysis phenotype observed in hTDP-43 worms.

### Pharmacological and genetical modulation of L-type calcium channels or calcium-activated large conductance potassium channels ameliorate movement defects in hTDP-43 worms

Another explanation for the discrepancy between aldicarb resistance and electrophysiological recordings in hTDP-43 worms is impaired coupling between neuronal excitation and secretion. Given a functional synaptic priming and fusion machinery (i.e. secretion; Figure 4), we focused on effects of Ca^2+^ at the presynaptic site and changes in membrane potential (i.e excitation)^65^. As these activities require functional synaptic ion channels, we assessed their function, using an explorative compound-screen with *a priori* selected ion channel targeting compounds. We selected compounds that cover potassium, calcium and sodium channels and include both antagonistic and agonistic players (**Supplementary Figure 3A,B**; Supplementary File 2)^66–82^. Priority was given to compounds previously tested and validated in *C. elegans* and with established targets (Supplementary File 2). Finally, to verify our methodology we included two compounds that are known to enhance thrashing rate in hTDP-43 worms: arecoline, as evidenced by our own results above, and the anti-epileptic drug ethosuximide^83–86^.

We developed a pipeline that involved a broad primary screen, hit selection and subsequent rescreening (**Supplementary Figure 3A**). First, dose-response and time-response curves of all selected compounds were generated by assessing the thrashing frequency of hTDP-43 worms. This approach was validated by arecoline and ethosuximide showing the expected increase in thrashing frequency of hTDP-43 worms, although the positive effect of ethosuximide was limited to the 20 °C condition only (**Supplementary Figure 3C,D**). Compounds that increased the thrashing frequency to a level that exceeded the 95% bootstrapped confidence interval of the untreated worms and had a dose-response relationship of R^2^>0.3 were considered potential hits (Supplementary File 2). Next, a triplicated rescreen was performed in hTDP-43, control and *unc-13(e51)* worms with the compounds and accompanying concentrations determined in the primary screen (**Supplementary Figure 3,4 and 5**). We identified an L-type calcium channel agonist (nefiracetam) and two potassium channel blockers (iberiotoxin and 4-aminopyrdine) as positive hits capable of increasing thrashing frequency in hTDP-43 worms after a treatment period of only 2 hours (**Supplementary Figure 3E-M**). Those compounds had no significant effects in *unc-13(e51)* (**Supplementary Figure 4**) and control worms (**Supplementary Figure 5**).

The observed increase in thrashing frequency upon modulation of specific ion channels is small. Nevertheless, one should take the short treatment window, to avoid potential side-effects, and the relatively inefficient uptake of compounds by *C. elegans* into account ^87–90^. To assess the full potency of the targets connected to the identified compounds in enhancing movement in hTDP-43 worms we took a genetic approach. Two of the identified compounds have established targets: nefiracetam potentiates L-type Ca^2+^ channels^74^ of which *C. elegans* has only one (EGL-19) and iberiotoxin specifically antagonizes Ca^2+^-dependent SLO-1 K^+^ channel ^78^, (**Figure 5A**). Hence, we constructed *egl-19* gain-of function, hTDP-43; *egl-19(ad695),* and *slo-1* loss-of-function, hTDP-43; *slo-1(js118),* double mutants to determine whether these genetic mutations recapitulate the action of the drugs. As a negative control we created an *egl-19* loss-of-function, hTDP-43; *egl-19(n582),* strain (**Figure 5A**), and crossed the control worms with all mutant strains as well.

First, we explored whether the double mutants improved aldicarb sensitivity and movement. As expected, both *egl-19(gf)* and *slo-1(lf)* mutations alone caused a left-shift in the aldicarb curve, but importantly both mutations restored aldicarb sensitivity of hTDP-43 worms (**Figure 5B**; **Supplementary Figure 6A**). This, in combination with unaltered levamisole-sensitivity (**Supplementary Figure 6B**) indicates that mutations in ion-channel activity expected to increase Ca^2+^ influx appear to enhance cholinergic release in hTDP-43 worms^68^. Subsequently, we looked at several movement phenotypes. Both the thrashing frequency and maximal crawling speed were significantly increased in hTDP-43 worms with either the *egl-19(gf)* or *slo-1(lf)* mutation, especially at 20 °C (**Figure 5C,D**). In contrast, the movement capacity of control worms was actually decreased by the same mutations, which has been previously described^68, 91, 92^. Next, we investigated whether the rescue was temporary or long-lasting. Hence, we looked at movement of the worms at adulthood day 4 and day 8 (**Figure 5E**) and observed a similar decrease in paralysis rate of hTDP-43 worms when EGL-19 or SLO-1 function was enhanced or inhibited, respectively. Conversely, disruption of EGL-19-function in a hTDP-43 background was lethal at an early age.

Importantly, the observed suppression of hTDP-43 toxicity by *slo-1(lf)* or *egl-19(gf)* mutations was not due to reduced hTDP-43 transgene expression (**Supplementary Figure 6C**). In fact, the hTDP-43 levels were increased in both the *egl-19(lf)* and *egl-19(gf)* background. This, together with the observation that *egl-19(gf)* mutants are seemingly hypercontracted (**Supplementary Figure 6D,E**) may partially account for the different extent of motor improvement caused by *egl-19(gf)* and *slo-1(lf)* mutations. Given the rescuing effects of *egl-19(gf)* and *slo-1(lf)* mutations on the movement capacity of hTDP-43 worms, we investigated whether these mutations also impacted endogenous mPSC frequency. The gain-of-function mutation in *egl-19* increased the mPSC frequency in hTDP-43 worms to WT levels, while decreasing the frequency in control worms (note, *n*=3, efforts to patch clamp muscles of dissected *egl-19* mutant worms were less successful due to muscle fragility) (**Figure 5F, G**). In contrast, the *slo-1(lf)* mutation had no significant effect on mPSCs in either the control or hTDP-43 worms (**Figure 5F, G**).

Taking together, these results reveal that improved function in L-type calcium channels or inhibition of calcium-activated large-conductance BK potassium channels partially rescue the movement defects in a hTDP-43 background. Moreover, our pharmacological and genetic data suggest that alterations in excitation/secretion coupling^68, 92, 93^ may underly the observed decrease in neurotransmission and impaired movement of hTDP-43 worms

### Optogenetic stimulation reveals incongruency between activation of individual circuit cell types and movement in hTDP-43 worms

Both EGL-19 and SLO-1 are expressed in neurons and muscle^68, 73, 92–95^. In fact, EGL-19 is the predominant voltage-gated ion channel in body-wall muscle ^73, 94, 95^, while SLO-1 has been suggested to control Ca^2+^ transients in muscles cells^94^. As a consequence, mutations in these genes will affect both neuronal and muscle membrane potential, making it impossible to easily attribute the observed effects to a specific cell type of the neuromuscular unit. For example, the observed hypercontraction of *egl-19(gf)* mutants has been attributed to the muscle compartment^92^. In order to further assess to what extent activation of individual cell types of the motor circuit can restore movement of TDP-43 worms, we opted for an optogenetic approach (**Figure 6A**).

**Figure 6:**
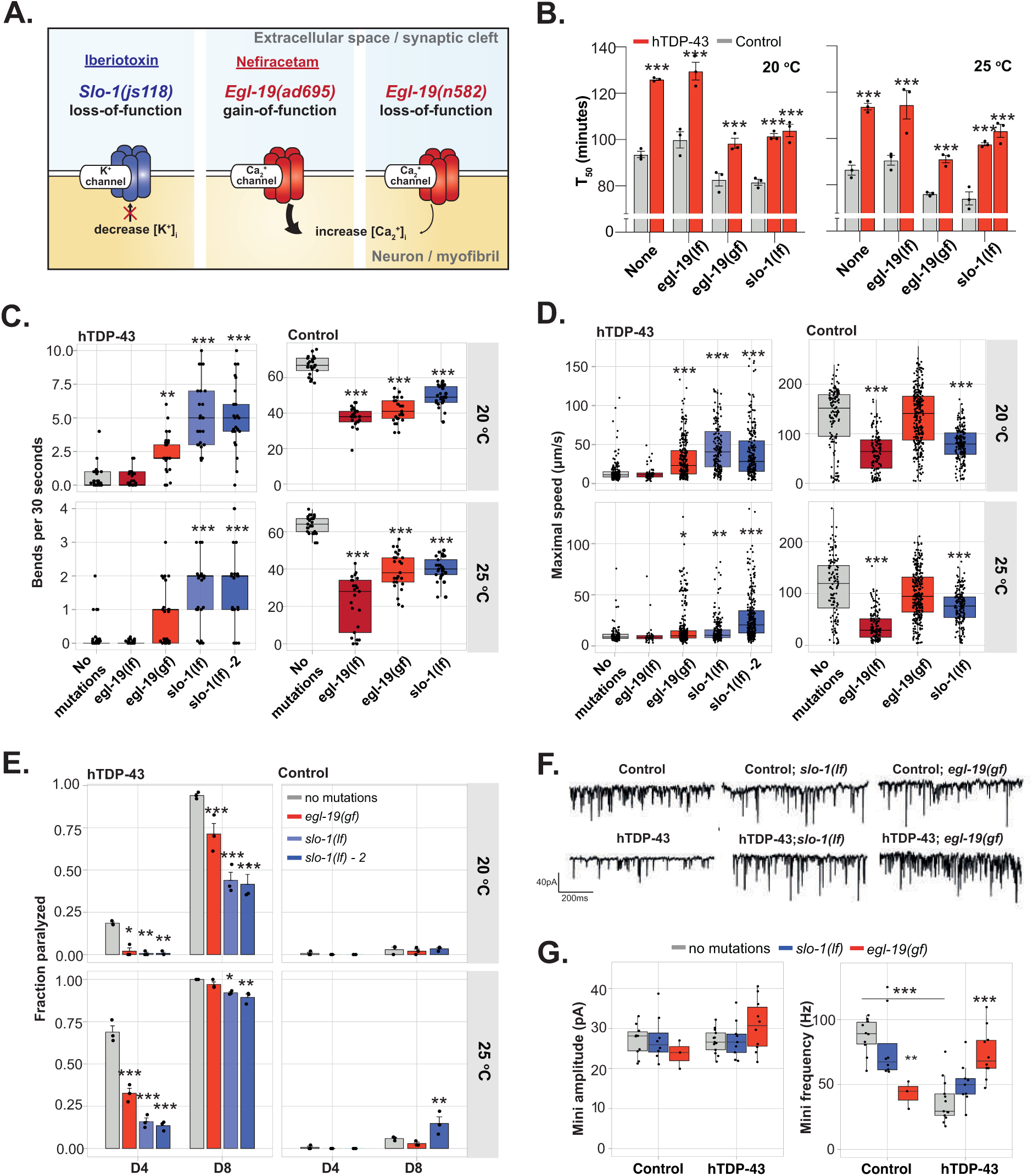
Gain-of-function of *egl-19* and loss-of-function of *slo-1* increase acetylcholine release and rescuesmovement defects in hTDP-43 worms. **A)** Schematics of the molecular targets identified by the compounds screen. Iberiotoxin inhibits SLO-1 channels and can be genetically mimicked by a *slo-1(lf)* mutation. Nefiracetam activates EGL-19 and can be genetically phenocopied by an *egl-19(gf)* mutation. **B)** The average time in which half of the worms are paralyzed on aldicarb represented as the mean T_50_, *n = 3*, two-way ANOVA at 20 °C (mutation, genotype: p<0.001, interaction: p =0.0089) and *25 °C (mutation, genotype: p<0.001, interaction: n.s.) with post-hoc Holm-Sidak’s.* ***C)*** *Thrashing ability, in bends per 30 seconds. On*ly intra-strain comparisons were performed, *n = 25*, Kruskal-Wallis for control at 20 °C and 25 °C (p<0.001), for hTDP-43 at 20 °C and 25°C (p<0.001.) with post-hoc Dunn’s. One representative experiment of a triplicate is shown. **D)** Maximum crawling speed on solid media. Only intra-temperature comparisons were performed, n = 60-270, two-way ANOVA at 20 °C (genotype, mutation, interaction: p<0.001) and 25 °C (genotype, mutation, interaction: p<0.001), post-hoc Dunnett’s. One representative experiment of a triplicate is shown. **E)** Paralysis rate at D4 and D8 of adulthood. Only intra-strain comparisons were performed, *n = 3,* two-way ANOVA for control at 20 °C (time: p = 0.0039, mutation, interaction n.s.) and 25 °C (time: p<0.001, mutation: p = 0.0117, interaction: p = 0.0094) and for hTDP-43 at 20 °C and 25°C (time, mutation, interaction: p<0.001) with post-hoc Dunnett’s. **F)** Representative traces of mPSCs of from hTDP-43, *slo-1(lf)* and *egl-19(gf)* single and double mutants. **G)** Quantification of mini amplitude and frequency in the single and double mutants. Mini amplitude: n ≥ 7 (except for *egl-19*), One-way ANOVA (n.s.). Mini frequency: n ≥ 7 (except for *egl-19*), One-way ANOVA (p<0.001) with post-hoc Holm-Sidak. Only intra-strain comparisons and direct comparisons between hTDP-43 and control worms (without additional mutations) were performed. Error bars represent the S.E.M. * = p<0.05, ** = p<0.01, ** = p<0.001.

We first expressed channelrhodopsin-2 (ChR2) specifically in cholinergic neurons to remotely control Ca^2+^-influx and concurrent activity of these neurons (**Supplementary Figure 6A,B**). Worms were exposed to a 3-3 light stimulus, in which light was OFF for 3 seconds, followed by light ON (∼1.6 mW/mm^2^) for 3 seconds. The change in body length during this light stimulus was used as a measure for cholinergic signaling^96–98^. We found drastically increased photo-induced body-contractions in worms expressing hTDP-43 compared to its controls (**Figure 6B-D**). This exaggerated response to cholinergic stimulation could not be ascribed to compensatory mechanisms within the muscle. In fact, direct photostimulation of muscle cells expressing ChR2 did not reveal differences in the extent of body contractions between hTDP-43 and control worms (**Figure 6E,F**).

Normally, concomitant GABA release is triggered by photostimulation of cholinergic neurons directly innervating contralateral GABAergic neurons. Since, GABA-signaling normally dampens photo-evoked contractions^97^, we controlled Ca^2+-^influx specifically in GABAergic neurons with ChR2 (**Supplementary Figure 7A,B**) and assessed the worms’ ability to elongate. In contrast to the exaggerated cholinergic response, we observed a clear reduction in the GABAergic output as evidenced by a diminished body elongation (**Figure 6G,H**). The decrease in photo-evoked body elongation was especially pronounced at 25 °C. In line with our functional, morphological and electrophysical evidence for deterioration of GABAergic neurons, these findings suggest a loss of GABAergic function, especially at 25 °C, which remains even persistent after photo-induced depolarization. Therefore, it is likely that the increased photo-evoked contractions after cholinergic stimulation are at least partially due to a loss of GABAergic signaling.

Subsequently, we explored the possibility that the paralysis of hTDP-43 worms can be partially or entirely rescued by increased Ca^2+^ influx in either cholinergic or GABAergic neurons. We exposed worms expressing ChR2 specifically in cholinergic or GABAergic neurons to 30 seconds of blue light (∼1.0 mW/mm^2^) and quantified their thrashing frequency before and during this lightning regime (**Figure 6I, J**). Counterintuitively, we found that GABAergic stimulation increased the thrashing frequency significantly in hTDP-43 worms during photo-stimulation, opposite to the effect seen in control worms (**Figure 6J**). Immediate photo-induced relaxation was followed by enhanced thrashing and this increased mobility was maintained for at least 30 seconds after stimulation. Apparently, a small Ca^2+^-induced increase in inhibitory output of the remaining GABAergic synapses results in an increased thrashing frequency. Indeed, GABAergic *unc-47* mutants that are unable to load GABA in synaptic vesicles do not change their thrashing frequency when stimulated (**Supplementary Figure 7C**). In contrast, cholinergic photo-activation, with concomitant indirect GABAergic neuronal stimulation, was less effective in enhancing motility, as a large portion of worms became hypercontracted and switched between ‘coiling’, ‘knotting’ and sporadic thrashing, comparable to G_q_-stimulation (**Figure 6I**). Neither GABAergic nor cholinergic stimulation of worms that were grown without ATR caused similar changes in swimming behavior (**Supplementary Figure 7D,E**).

Taking everything together, we show that introducing a depolarizing current specifically into GABA or acetylcholine neurons results in enhanced neurotransmission. In particular, the activation of GABA neurons via directed Ca^2+^-influx ameliorates the movement defects seen in hTDP-43 worms, suggesting a role of ion influx or resting potential in staggered neurotransmission. Additionally, our results also suggest that the muscle excitation-to-inhibition ratio is shifted towards excitation. Based on the collective optogenetic, structural and electrophysiological data, we propose a model in which cholinergic and GABAergic output are imbalanced (**Figure 6K**, step I). In this model, maximal cholinergic stimulation will preserve this imbalance (**Figure 6K**, step II), while sole GABAergic stimulation of remaining synapses will partially restore the balance and concurrently enhance movement (**Figure 6K**, step III).

### Rebalancing muscle excitation-to-inhibition ratio towards inhibition enhances movement in hTDP-43 worms

Having established a potential role of ion channels in hTDP-43 pathology, we decided to further examine whether post-synaptic alterations in muscle excitation-to-inhibition (E/I) ratio contribute to movement dysfunction. We made use of the well-established pathways that regulate muscle contraction (**Figure 7A**). First, we investigated the transcript levels of numerous structural and receptor genes of the post-synaptic unit with qPCR. We found that both GABAergic (*unc-49*) and cholinergic (e.g. *acr-16, lev-1, unc-63*) receptors are strikingly upregulated at 25 °C, suggesting post-synaptic transcriptional adaptation (**Figure 7B**). Moreover, neither the functional capacity of the muscle (**Figure 6E,F**) nor its structure is changed (**Supplementary Figure 8A**). Phalloidin staining of muscle cells did not reveal morphological aberrations (**Supplementary Figure 8A**). Nevertheless, we did observe delayed relaxation kinetics in the body wall muscle of hTDP-43 worms that might be indicative of inefficient calcium handling and impaired relaxation (**Supplementary Figure 8B**). In addition, hTDP-43 worms tend to be hypercontracted when their contraction indices are compared to controls worms (**Supplementary Figure 4D, 8C**).

**Figure 7:**
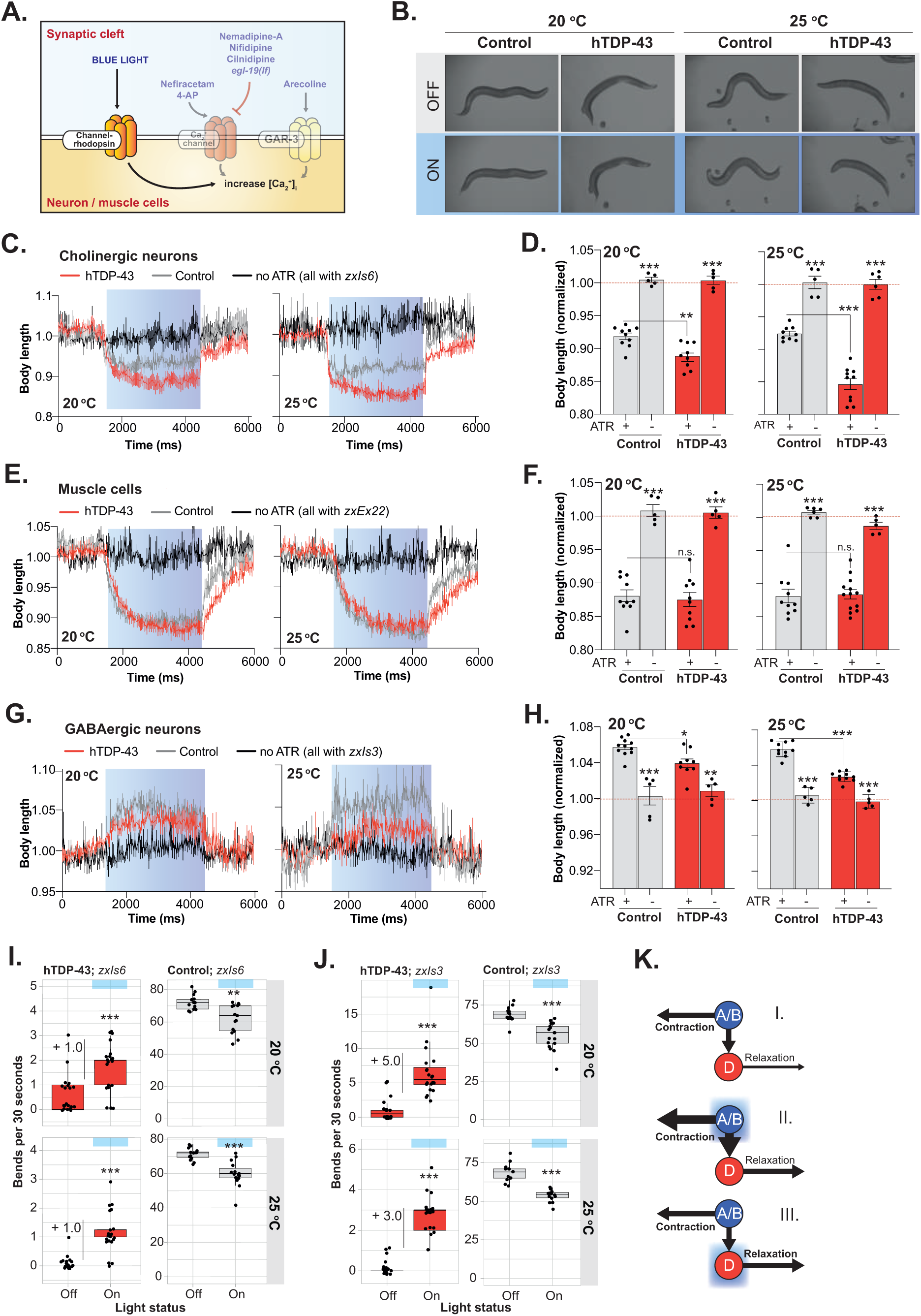
Optogenetic stimulation of neuronal calcium influx in either GABAergic or cholinergic neurons increases mobility in hTDP-43 worms. **A)** Schematic showing the different pathways involved in increasing intracellular calcium levels. With the use of the channelrhodopsin (ChRd), calcium-gating can be induced by blue light. **B)** Representative pictures showing the body length of worms when blue light is turned off or on. The worms shown have ChRd expressed under the *unc-17* promoter. **C)** Representative graphs showing body length changes during blue-light illumination for 3 seconds (blue bar) in worms carrying ChRd in cholinergic neurons (P*unc-17*), *n =* 7-8 worms per condition. **D)** Mean relative body length during blue-light illumination of worms carrying ChRd in cholinergic neurons (P*unc-17*), *n =* 8-9 (5 for -ATR), one-way ANOVA for both 20 °C and 25 °C (p<0.001) with post-hoc Bonferroni’s multiple comparisons tests. Asterisks belonging to the non-ATR conditions, show intra-genotypic comparisons only. One representative experiment of a triplicate is shown **E)** Representative graphs showing body length changes during blue-light illumination for 3 seconds (blue bar) in worms carrying ChRd in their muscle cells (P*myo-3*), *n =* 10-12 worms per condition. **F)** Mean relative body length during blue-light illumination of worms carrying ChRd in muscle cells (P*myo-3*), *n =* 8-10 (5 for -ATR), one-way ANOVA for both 20 °C and 25 °C (p<0.001) with post-hoc Bonferroni’s multiple comparisons tests. Asterisks belonging to the non-ATR conditions, show intra-genotypic comparisons only. One representative experiment of a triplicate is shown **G)** Representative graphs showing body length changes during blue-light illumination for 3 seconds (blue bar) in worms carrying ChRd in GABAergic neurons (P*unc-47*), *n =* 6-8 worms per condition. **H)** Mean relative body length during blue-light illumination of worms carrying ChRd in GABAergic neurons (P*unc-47*), *n =* 8-9 (5 for -ATR), one-way ANOVA for both 20 °C and 25 °C (p<0.001) with post-hoc Bonferroni’s multiple comparisons tests. Asterisks belonging to the non-ATR conditions, show intra-genotypic comparisons only. One representative experiment of a triplicate is shown. **I)** Thrashing ability, in bends per 30 seconds, of worms not exposed to blue light (‘off’) or exposed to blue light (‘on’). Worms express ChRd in their cholinergic neurons. Only intra-strain comparisons were performed, *n* = 15-20, Kruskal-Wallis for control at 20 °C (p=0.0024), and 25 °C (p<0.001), for hTDP-43 at 20 °C and 25°C (p<0.001) with post-hoc Dunn’s. One representative experiment of a triplicate is shown. **J)** Thrashing ability, in bends per 30 seconds, of worms not exposed to blue light (‘off’) or exposed to blue light (‘on’). Worms express ChRd in their GABAergic neurons. Only intra-strain comparisons were performed, *n* = 15-20, Kruskal-Wallis for control at 20 °C and 25 °C (p<0.001), for hTDP-43 at 20 °C and 25°C (p<0.001) with post-hoc Dunn’s. **K)** Hypothetical model based on the optogenetics experiments. I. Suggested situation in hTDP-43 worms: GABAergic signaling is lower than cholinergic output. II. enhancing cholinergic output also indirectly affects GABAergic activity via the specific wiring of the nervous system. It however does not change the disbalance between the two branches. III. Specific stimulation of GABAergic neurons does not directly affect cholinergic output and can therefore be used to restore the suggested E/I disbalance. One representative experiment of a triplicate is shown. Error bars represent S.E.M. * = p<0.05, ** = p<0.01, ** = p<0.001.

**Figure 8:**
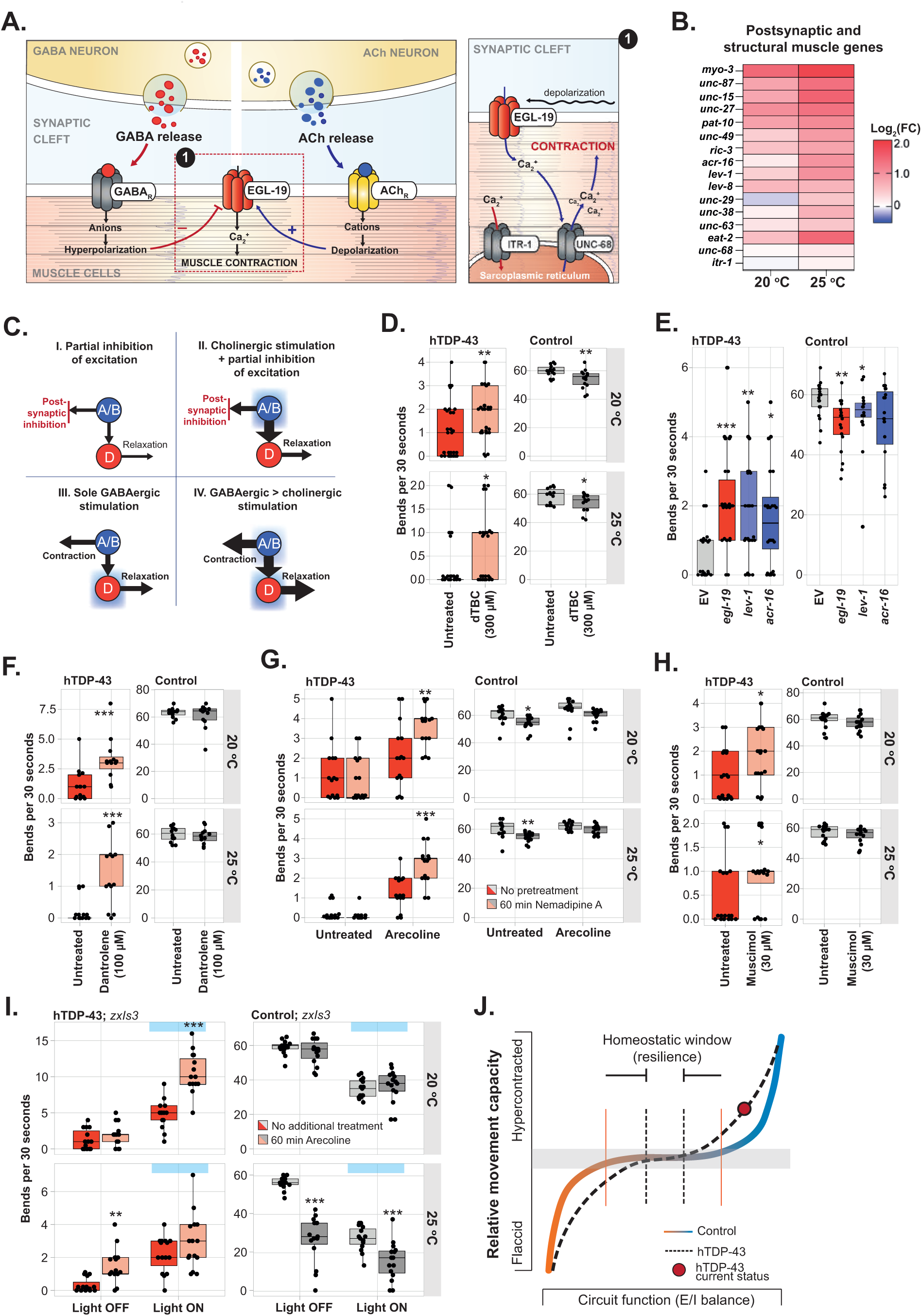
Balancing the muscle excitation-to-inhibition ratio enhances movement in hTDP-43 worms. **A)** Simple representation of the genes and pathways involved in muscle contraction and relaxation. GABAergic signaling actively relaxes the muscle, while cholinergic signaling enhances the post-synaptic potential and induces calcium influx via EGL-19. The calcium signal is enhanced via the ryanodine receptor UNC-68 at the sarcoplasmic reticulum. **B)** A heatmap showing the log_2_ fold-change of genes that are part of in the post-synaptic muscle compartment, normalized by *pmp-3*, in hTDP-43 worms compared to their controls at 20 °C and 25 °C. **C)** Extended hypothetical models that are based on the idea that the E/I balance is disturbed and excitation > inhibition (**Figure 7K**). We speculate that 4 different treatments could restore this balance (I-IV). **D)** Thrashing ability, in bends per 30 seconds, with or without pretreatment with 300 μM dBTC. Only intra-strain comparisons were performed, *n* = 15, Mann-Whitney U-test, except for control at 20 °C: Student’s t-test. One representative experiment of a triplicate is shown. **E)** Thrashing frequency after treatment with different RNAis from L1 in a muscle-specific RNAi background. Only intra-strain comparisons were performed, *n* = 15-25, Kruskal-Wallis test for hTDP-43 (p < 0.001) and control (p = 0.0106) with post-hoc Dunn’s. One representative experiment is shown. **F)** Thrashing frequency of worms treated with the UNC-68 antagonist Dantrolene. Two-tailed unpaired Student’s t-test, *n = 15.* One representative experiment of a triplicate is shown. **G)** Thrashing frequency of worms that received no pretreatment or pretreatment with Nemadipine A and were subsequently exposed to arecoline or water (untreated). Only intra-strain comparisons were performed, *n* = 15-25, two-way ANOVA for hTDP43 at 20 °C (arecoline: p<0.001, interaction: p = 0.0124, pretreatment: n.s.) and at 25 °C (arecoline, interaction: p<0.001, pretreatment: p = 0.001), for controls at 20 °C (arecoline: p = 0.0018, pretreatment: p = 0.0012, interaction: n.s.) and at 25 °C (arecoline: p = 0.0015, pretreatment: p = 0.0014, interaction: n.s) with post-hoc Holm-Sidak’s. **H)** Thrashing ability, in bends per 30 seconds, with or without pretreatment with 30 μM muscimol. Only intra-strain comparisons were performed, *n* = 15, Mann-Whitney U-test. One representative experiment of a triplicate is shown. **I)** Thrashing frequency of worms that received no pretreatment or pretreatment with arecoline and in which GABAergic neurons were photo-activated. Only intra-strain comparisons were performed, *n* = 15-25, two-way ANOVA for hTDP43 at 20 °C and at 25 °C (GABA, pretreatment, interaction: p < 0.001), for controls at 20 °C (GABA: p <0.001, pretreatment: p = 0.0014, interaction: n.s.) and at 25 °C (GABA: p < 0.0015, pretreatment, interaction: n.s) with post-hoc Holm-Sidak’s. **J)** Hypothetical model of narrowed homeostatic window in hTDP-43 worms. Due to decreased neuronal output, small perturbations of components of the neuromuscular system will lead to larger responses in hTDP-43 worms. The red dot annotates the current state of hTDP-43 worms, the neuromuscular circuit is flipped to an increased E/I ratio that, due to a narrowed homeostatic window, results in a hypercontracted state and decreased ability to move. Error bars represent S.E.M. * = p<0.05, ** = p<0.01, ** = p<0.001.

With all the aforementioned observations, we expanded our excitation-to-inhibition model of hTDP-43 worms and generated four scenarios that, if accurate, would provide additional evidence for a system-wide shift between excitation and inhibition (**Figure 7C**). First, we argued that partially inhibiting cholinergic signaling post-synaptically would shift the E/I balance towards inhibition (**Figure 7C**, scenario I). Therefore, we treated worms with the AChR-antagonist dTBC and subsequently assessed their thrashing frequency in liquid. While small, we found a significant increase in thrashing frequency (**Figure 7D**). Complementarily, we crossed our hTDP-43 worms with a muscle-specific RNAi strain and knocked down several post-synaptic genes from L1 required for muscle contraction: *egl-19, acr-16, lev-1*, *unc-63* and *unc-68*^99–103^. Only *egl-19, acr-16* and *lev-1* RNAi initiated a decrease in transcript levels in at least 2 out of 4 replicates at 20 °C (**Supplementary Figure 8D**). At 25 °C, however, RNAi-knockdown was inefficient as transcript levels were unchanged. *Unc-68* RNAi was lethal in hTDP-43 worms, therefore we tried inhibiting this ryanodine receptor with its antagonist dantrolene^104^. The knockdown or inhibition of post-synaptic receptors and ion channel all increased the thrashing frequency of hTDP-43 worms (**Figure 7E, F**).

Secondly (**Figure 7C**, scenario II), because cholinergic signaling is reduced in hTDP-43 worms, we hypothesized that enhancing cholinergic signaling (and thus indirectly GABAergic signaling as well) together with post-synaptic inhibition might further increase the movement capacity of hTDP-43 worms. Hence, we either inhibited post-synaptic genes pharmacologically (*egl-19* and *unc-68*) or transcriptionally (*egl-19*, *lev-1*, *acr-16*) and treated worms with the G_q_-enhancer arecoline. Again, we did observe significantly increased movement capacity when post-synaptic receptors or ion channels were inhibited (**Figure 7G**, **Supplementary Figure 8E**). Moreover, substituting arecoline for optogenetic stimulation of cholinergic neurons while post-synaptic genes were blocked resulted in a similar increase in motility (**Supplementary Figure 8F,G**).

Thirdly (**Figure 7C**, scenario III), while inhibition of the cholinergic branch likely shifts the E/I balance and enhances movement, direct stimulation of GABARs should do the same. As expected, direct stimulation of the GABARs with muscimol also increased the thrashing frequency of hTDP-43 worms (**Figure 7H**). Noteworthy, pharmacological inhibition of the post-synaptic receptors or channels involved in muscle contraction or activation of receptors inducing muscle relaxation converge and both result in a reversal of the hypercontracted state of hTDP-43 worms (**Supplementary Figure 7C**).

Fourthly, both GABAergic and cholinergic signaling are decreased in hTDP-43 worms but seemingly to a different extent. Therefore, we hypothesized that enhancing both branches simultaneously but at different magnitudes (**Figure 7C**, scenario IV) might increase motility even further. We pre-treated worms expressing ChR2 in GABAergic neurons with the cholinergic facilitating agent arecoline and subsequently photo-activated GABAergic neurons. This combination treatment results in a fold-change of 10, the highest rescue of hTDP-43 worms acquired so far (**Figure 7I**; **Supplementary Figure 8H**). Nevertheless, the effect appeared to be limited to hTDP-43 worms that were grown at 20 °C.

Altogether, our results suggest that diminished circuit activity together with a prevailing imbalance between excitation and inhibition exists at the structural and functional level in hTDP-43 worms and contributes to movement dysfunction. Unexpectedly, in a system with highly degenerated inhibitory neurons, hTDP-43 worms still benefit from a re-established E/I ratio in terms of improved movement.

## Discussion

Using a phenogenomic approach on a *C. elegans* model for pan-neuronal expression of hTDP-43, we identified altered cholinergic and GABAergic neurotransmission as main biological processes associated with the phenotypic profiles of hTDP-43 worms. We reveal that hTDP-43 worms display an overall reduction in neurotransmission in combination with an imbalance between excitatory and inhibitory signals of the nervous system. Activation of excitatory cholinergic neurons via G-protein coupled receptor pathways or the modulation of ion channels could enhance neurotransmission but only slightly improved motor function. Such activation, in addition, resulted in adverse movement effects including hypercontraction, spastic paralysis and coiling. Excitingly, restoring and balancing the functions of both excitatory and inhibitory cells simultaneously, synergized the effect of cell-autonomous interventions and, moreover, rescued effective locomotion without the aforementioned unfavorable side effects. Our results suggest that rebalancing neuronal circuits, rather than enhancing the function of individual cell types within the network, is required to restore function in neurodegenerative diseases.

### Different origins of impaired neurotransmission between GABAergic and cholinergic neurons

Our findings show that GABAergic neurons were more damaged than cholinergic neurons in hTDP-43 worms. The morphological abnormalities and degenerative character of the GABAergic branch are in line with the less reversible nature of its functional output, as shown by optogenetic stimulation. Whether our observations represent increased vulnerability of GABAergic neurons to hTDP-43 or merely reflect the differences in expression levels, remain to be determined. The synaptobrevin promotor used to drive pan-neuronal expression of hTDP-43 is not equally expressed in GABAergic and cholinergic neurons^105^, potentially resulting in increased loads of hTDP-43 in GABAergic neurons.

Deficits in cholinergic neurotransmission, on the other hand, can be completely reversed via cholinergic facilitation pathways and modulation of specific ion channels, as evidenced by the reversal of aldicarb-resistance. Together with the absence of synaptic degeneration and evidence for a functional synaptic machinery, these observations suggest that cholinergic neurons are functionally repressed in the presence of hTDP-43. Nonetheless, since a loss of GABAergic output also modulates aldicarb sensitivity^106^ it is not unlikely that the seemingly reverted cholinergic transmission is confounded by GABAergic degeneration. Thus, while cholinergic neurons appear to be only functionally silenced in a hTDP-43 background, we cannot exclude that some function has already been irreversibly lost.

### Cholinergic hypoactivity as consequence of altered input of the connectome

The reduction in endogenous post-synaptic currents together with the absence of synaptic degeneration points towards intrinsic hypoexcitability as a potential mechanism underlying the hypoactive state of cholinergic neurons in hTDP-43 worms. This hypothesis is strengthened by the positive effects of unidirectional modulation of ion channels, driving depolarization, on neurotransmission and locomotion. Both the pharmacological and genetic activation of L-type calcium channels and inhibition of calcium-activated large conductance BK potassium channels results in enhanced post-synaptic currents, neurotransmission and movement in hTDP-43 worms. Intriguingly, those interventions not only resulted in a direct improvement of motor function, but also positively impacted paralysis in the long term.

However, while spontaneous transmitter release is reduced by the presence of hTDP-43, evoked responses are intact, which does not per se point to intrinsic hypoexcitability. Given our results, the connectome of cholinergic neurons likely also contributes to the observed hypoactivity. Serotonergic neurons, known to inhibit synaptic transmission at the *C. elegans* neuromuscular junction^62, 64^, could play a role in silencing cholinergic neurons in hTDP-43 worms. We found that antagonizing serotonin receptors, thereby shutting down the GOA-1-dependent G_o_-pathway, resulted in a strikingly reverted aldicarb resistance and thus enhanced cholinergic neurotransmission. Simultaneously, we report that the spare functional capacity of cholinergic neurons can be recruited via stimulation of the GAR-3-EGL-30 pathway, which is normally activated via volume transmission of extrasynaptically released ACh^57^.

Therefore, we suggest that the altered input of the worm’s connectome to cholinergic neurons contributes to the reduced functional output in a hTDP-43 background. In such a framework it remains to be determined whether modulation of intrinsic excitability via ion channels may simply overshoot existing inhibitory and modulatory input to cholinergic neurons, or actually revert an intrinsic hypoexcitable state. The latter is not unlikely, since activation of EGL-30 also indirectly increases neuronal calcium levels^57, 107^. Thus, while we cannot conclude whether the repressed cholinergic function is solely secondary to alterations in the connectome, we do show that spare functional capacity resides within this neuron population.

### hTDP-43 expression results in a disbalanced, hyperreactive motor circuit

Considering the finding that animals with laser-ablated GABAergic MNs perform normal locomotion, albeit with decreased amplitude^108^, the activation of cholinergic neurons had an unexpected low effect on movement capacity. The rescue of motor impairment was always small, even when the previously determined effectors underlying cholinergic hypoactivity were specifically accounted for. In hTDP-43 worms cholinergic stimulation results in exaggerated down-stream responses, as evidenced by pharmacological and optogenetic-induced coiling, hypercontraction and spastic paralysis. These results suggest the existence of hyperreactivity to excitatory signals of the motor system at the level of the muscular compartment.

Inhibitory motoneurons normally dampen evoked contractions and are therefore important in counteracting excitatory signals^97, 108^. GABAergic input is, in turn, also necessary for rapid, high-frequency undulation^109^. Therefore, the loss of inhibitory synapses in hTDP-43 worms may account for both the hyperreactive response to excitatory signals and the reduced thrashing frequency. Considering the highly degenerative character of the GABAergic branch, the finding that stimulation of the remaining inhibitory system resulted in an increase in movement capacity was equally exciting and unexpected. Intriguingly, locomotion could be even further enhanced when different multi-drug approaches, considering the prevailing disbalance between inhibition and excitation and intrinsic properties of single neuron populations, were applied. We reveal that a combinatory intervention targeting cholinergic GPCRs and enhancing calcium-induced activation of GABAergic neurons, effectively restores motor function.

Considering all the aforementioned observations, we propose that hTDP-43 toxicity causes changes in the muscle excitation-to-inhibition ratio. In **Figure 7J** we show a chair-shaped homeostatic curve (coloured line) (based on models in: ^110, 111^) that demonstrates the window (between solid orange vertical lines) in which normal motor function can be maintained despite changes in the E/I ratio. Based on our observations, we propose that increases of the E/I ratio beyond the homeostatic window will lead to hypercontracted paralysis, whereas decreases could lead to flaccid paralysis. As functional output of the motor system declines, as in the hTDP-43 worms, the homeostatic plateau narrows (dashed black vertical lines). Consequently, small perturbations in the E/I ratio will result in a drastic change in contraction status, exemplified by the large effects of compound treatment or genetic interventions on the contraction index of hTDP-43 worms. From this perspective, the reduced cholinergic output might actually be an adaptive response to maintain a balanced circuit. Also, the magnitude of the observed rescue in the ion channel mutants warrants a critical side note when assuming this model. Both a loss-of-function of SLO-1 and a gain-of-function EGL-19 increase post-synaptic calcium transients^92, 94^ and therefore likely preserve or even enhance the E/I imbalance at the muscular compartment.

It remains to be determined whether the observed post-synaptic hyperreactivity (e.g. muscle hypercontraction) is only the consequence of GABAergic dysfunction or also reflects an adaptive response of the post-synaptic unit itself. Transcript levels of post-synaptic receptors hint towards increased expression levels of both cholinergic and GABAergic receptors, suggesting that network adaptations are not limited to neuronal tissue but also involve muscle cells. Clearly, hTDP-43 affects the neuromuscular system at different levels, ranging from alterations in intrinsic properties of individual neuron populations, altered synaptic input and even imbalanced communication at the neuro-muscular interface.

### From phenotypic profiles to new therapeutic strategies

ALS causes death within 2-5 years after onset and has to date no cure or effective treatment^112, 113^. With only one approved disease-modifying drug, Riluzole, treatment of ALS revolves around symptomatic care^114–117^. Undoubtedly, the lack of success in clinical trials underlines the apparent complexity and unknown etiology of ALS but also emphasizes the need for reassessment of the methods used in ALS drug development ^118, 119^.

While the degeneration of glutamatergic upper motor neurons and cholinergic lower motor neurons are characteristic for ALS, classifying ALS as a strict neuromuscular disease has become untenable as neuronal susceptibility is not confined to motor neurons (MNs)^120–122^. There is evidence from studies on ALS to suggest that connecting neurons, including gamma, sensory and serotonergic neurons, may contribute to motoneuron pathology ^123–129^. The idea that MN vulnerability cannot be solely ascribed to intrinsic properties of MNs but likely also encompasses input via synaptic connections^122–127, 130, 131^, suggests that drug discovery studies should examine the full connectome of the motor system instead of focusing solely on MNs.

Here, we show that a phenomics approach can be used to identify the neuronal basis for the observed behavioral abnormalities in a *C. elegans* model for TDP-43-induced neurodegeneration. By leveraging our predictions, the ensuing systemic approach yielded several (poly-)therapeutic interventions that restored motor dysfunction at the behavioral level. Thus, we provide the field with a novel pipeline making use of emergent properties of the full motor connectome to identify and test druggable targets in TDP-43 pathologies. In such methodology, therapeutic interventions do not solely rely on altered neuronal properties but do consider the maladapted network as well. The work presented here provides a basis for focusing on how hTDP-43 affects neuronal and network function and to identify the molecular and cellular players involved. In addition, given the multifactorial and complex nature of ALS, our work also underlines that poly-therapeutic interventions should be considered as promising approaches. This strategy is clearly not limited to ALS, but might prove its strength in other TDP-43 proteinopathies and neurodegenerative diseases as well. Therefore, future research should aim to elucidate whether promising therapeutics that failed clinical trials may still harbor beneficial synergistic effects when combined with other interventions.

Our phenomics approach complements other work with *C. elegans* ALS models^132^. We used an unbiased set of behavioral features and phenotypic profiles as a starting point for further investigation, instead of performing genetic screens, drug screens or hypothesis-driven interventions to identify modifiers, targets and therapeutics^83, 133–139^. The behavioral effects of our interventions also differ, as we do see instant effects of treatment which do not depend on long-term interventions. As evidenced by our and other studies, the combination of representative ALS-mimicking models and simplistic motor circuits can be valuable in identifying the biological mechanisms contributing to motor dysfunction in ALS ^138^. Whether our findings in *C. elegans* can be translated to human patients remains to be determined. Worms lack for example the typical bifurcation of upper and lower motor neurons that are characteristic for human, have direct GABAergic signaling at the neuromuscular junction, and generate not sodium but calcium-dependent all-or-none action potentials at the muscle^108, 140, 141^.

Despite these critical differences, changes in cortical inhibition are thought to precede the clinical onset of motor dysfunction in ALS by several months^23, 142^, implying that GABAergic alterations may be an early sign of ALS^143^. Consistently, reduced GABA levels in the motor cortex of ALS patients have been reported^23, 125, 143, 144^ and degeneration of parvalbumin-positive cortical interneurons have been observed^145, 146^. Our results support the idea that a loss of inhibition is an early sign in TDP-43-induced ALS. Moreover, we show that recruiting the spare capacity of already degenerative GABAergic neurons provides a way of restoring motor function. In line with that, studies in rodent models and neuron cultures have highlighted GABAergic modulation as promising intervention in ALS^147, 148^. Noteworthy, also altered excitability has been proposed to represent one of the earliest modifications in a cascade of pathological events in ALS (as reviewed in: ^20, 149^).

Overall, our findings reinforce the importance of considering the surrounding network of neurons in the initiation and progression of clinical symptoms in neurodegenerative diseases. The ability to study the output of a full connectome with diseased elements is not only crucial for elucidating the components contributing to pathology, but also for assessing the often-unexpected side effects of therapeutic interventions. Neurons are embedded within complex circuits in which their activity, in addition to cell-intrinsic processes, is extrinsically regulated. Likely, even small alterations in the output of individual neurons will affect its connected network and may result in constant adaptations of the whole circuitry, including non-neuronal tissue. Therefore, interventions solely aiming to restore function of diseased neurons might actually aggravate disease symptoms by disrupting the newly established homeostatic status of the system. Whether our findings in a *C. elegans* model for TDP-43 induced toxicity might translate to higher model-organisms and eventually ALS patients remains to be determined.

## Methods

### Strains and maintenance

Standard conditions were used for *C. elegans* propagation at 20 °C^150^. Animals were synchronized by hypochlorite bleaching, unless stated differently, hatched overnight in M9 buffer at 20 °C and subsequently cultured on NGM agar plates seeded with OP50. Worms were grown from L1 until adulthood D1 at either 20 °C or 25 °C. For ageing experiments, worms were transferred ∼72 hours after plating to NGM plates containing 5-Fluoro-2’deoxy-uridine (FUdR) to inhibit growth of offspring. The following genotypes were used: Table 1.

**Table 1:** genotypes used.

| Strain | Description | Genotype | Remark |
| --- | --- | --- | --- |
| N2 | Bristol wild isolate | Wildtype |  |
| OW1601 | CL6049 6x backcrossed with N2. Named: hTDP-43 worms. | dvIs62[*snb-1p::hTDP-43/3'long UTR + mtl-2p::GFP*]X | Gift of Chris Link |
| OW1603 | CL2122 6x backcrossed with N2. Named: control worms. | dvIs15[(*pPD30.38*)*unc-54 (vector) + (pCL26)mtl-2p::GFP*] | Gift of Chris Link |
| OW1640 | *egl-19* loss-of-function in hTDP-43 background | *egl-19(n582)IV*; dvIs62 [*snb-1p::hTDP-43/ 3'Long UTR + mtl-2p::GFP*]X |  |
| OW1641 | *egl-19* gain-of-function in hTDP-43 background | *egl-19 (ad695)IV*; dvIs62 [*snb-1p::hTDP-43/ 3'Long UTR + mtl-2p::GFP*]X |  |
| OW1642 | *egl-19* loss-of-function in control background | *egl-19 (n582)IV*; dvIs15 [(*pPD30.38*) *unc-54 (vector) + (pCL26) mtl-2p::GFP*] |  |
| OW1643 | *egl-19* gain-of-function in control background | *egl-19(ad695)IV*; dvIs15[(*pPD30.38*) *unc-54 (vector) + (pCL26) mtl-2p::GFP*] |  |
| OW1644 | Muscular *ChR2::YFP* expression in hTDP-43 background | zxEx22 [*myo-3p::ChR2 (H134R)::YFP + lin-15(+)*]; dvIs62 [*snb-1p::hTDP-43/ 3'Long UTR + mtl-2p::GFP*]X |  |
| OW1645 | Muscular *ChR2::YFP* expression in control background | zxEx22 [*myo-3p::ChR2(H134R)::YFP + Lin-15 (+)*]; dvIs15 [(*pPD30.38*) *unc-54 (vector) + (pCL26) mtl-2p::GFP*] |  |
| OW1646 | GABAergic *ChR2::YFP* expression in hTDP-43 background | zxIs3 [*unc-47p::ChR2 (H134R)::YFP + lin-15 (+)*]; dvIs62 [*snb-1p::hTDP-43/ 3'Long UTR + mtl-2p::GFP*]X |  |
| OW1647 | GABAergic *ChR2::YFP* expression in control background | zxIs3 [*unc-47p::ChR2 (H134R)::YFP + lin-15 (+)*]; dvIs15 [(*pPD30.38*) *unc-54 (vector) + (pCL26)mtl-2p::GFP*] |  |
| OW1648 | Cholinergic *ChR2::YFP* expression in hTDP-43 background | zxIs6 [*unc-17p::ChR2 (H134R)::YFP + lin-15 (+)*]V; dvIs62 [*snb-1p::hTDP-43/ 3'Long UTR + mtl-2p::GFP*]X |  |
| OW1649 | Cholinergic *ChR2::YFP* expression in control background | zxIs6 [*unc-17p::ChR2 (H134R)::YFP + lin-15 (+)*] V; dvIs15 [(*pPD30.38*) *unc-54 (vector) + (pCL26) mtl-2p::GFP*] |  |
| OW1654 | Muscle-specific RNAi in control background | *rde-1(ne219)V*; kzIs [*hlh-1p::rde-1 + sur-5p:: NLS::GFP*]; dvIs15 [(*pPD30.38*) *unc-54 (vector) + (pCL26) mtl-2p::GFP*]; |  |
| OW1655 | Muscle-specific RNAi in hTDP-43 background | *rde-1(ne219)V*; kzIs [*hlh-1p::rde-1 + sur-5p:: NLS::GFP*]; dvIs62 [*snb-1p::hTDP-43/3'long UTR + mtl-2p::GFP*]X; |  |
| OW1656 | *slo-1* loss-of-function in hTDP-43 background | *slo-1(js118) III*; dvIs62 [*snb-1p::hTDP-43/3'long UTR + mtl-2p::GFP*]X |  |
| OW1657 | *slo-1* loss-of-function in control background | *slo-1(js118)III*; dvIs15 [(*pPD30.38*)*unc-54 (vector) + (pCL26)mtl-2p::GFP*] |  |
| OW1658 | Cholinergic mCherry expression in hTDP-43 background | dvIs62[*snb-1p::hTDP-43/3'long UTR + mtl-2p::GFP*]; oxEx838 [*unc-17p::mcherry + lin-15 (+)*] |  |
| OW1659 | Cholinergic mCherry expression in control background | dvIs15 [(*pPD30.38*) *unc-54 (vector) + (pCL26) mtl-2p::GFP*]; oxEx838[*unc-17p::mcherry + lin-15(+)*] |  |
| OW1660 | GABAergic mCherry expression in hTDP-43 background | oxIs608[*unc-47p::mcherry*]V; dvIs15[(*pPD30.38*) *unc-54 (vector) + (pCL26) mtl-2p::GFP*] |  |
| OW1661 | GABAergic mCherry expression in control background | oxIs608 [*unc-47p::mcherry*]V; dvIs15 [(*pPD30.38*) *unc-54 (vector) + (pCL26) mtl-2p::GFP*] |  |
| OW1663 | GABAergic synapses tagged with tagRFP in control background | dvIs15 [(*pPD30.38*)*unc-54 (vector) + (pCL26)mtl-2p::GFP*]; oxEx1411 [*Punc-47::snb-1::tagRFP, Pcc::GFP*] |  |
| OW1664 | GABAergic synapses tagged with tagRFP in hTDP-43 background | dvIs62[*snb-1p::hTDP-43/3'Long UTR + mtl-2p::GFP*]X; oxEx1411 [*unc-47p::snb-1::tagRFP, Pcc::GFP*] |  |
| KG1645 | Cholinergic synapses tagged with tagRFP | ceIs61[Punc-129::flp-3::venus, Punc-129::mCherry-snb-1, Ptx-3::mCherry]II |  |
| EG10039 | Cholinergic synapses tagged with tagRFP in hTDP-43 background | ceIs61[Punc-129::flp-3::venus, Punc-129::mCherry-snb-1, Ptx-3::mCherry] II; dvIs62[pmtl-2::GFP; psnb-1::hTDP-43] X |  |
| MT7929 | *unc-13* loss-of-function | *unc-13 (e51)I* |  |
| ZX460 | Cholinergic *ChR2::YFP* expression | *Lin-15B&lin-15A(n765) X; zxls6[unc-17p::ChR2(H134R)::YFP + lin-15(+)] V* | Gift of A. Gottschalk |
| ZX463 | GABAergic *ChR2::YFP* expression in a *unc-47* loss-of-function background | *unc-47 (e307)III; zxls3 [unc-47p::ChR2(H134R)::YFP; lin-15(+)]* | Gift of A. Gottschalk |

### Western blotting/SDS-PAGE

Worms were collected in PBS supplemented with Complete Protease Inhibitor (Roche; #11697498001) and then sonicated for 15 cycles: 30 sec on, 15 sec off. Worm debris was spun down and the protein concentration was determined using the PierceTM BCA Protein Assay Kit (Thermo Fischer; #23225) according manufacturer’s protocol. 20-30µg protein was dissolved in laemmli buffer (5x) and boiled at 95°C for 10 minutes. Subsequently, samples were loaded on a 12% Tris-Glycine acrylamide gel. In some cases, worms were directly picked into laemmli buffer (5x) and boiled at 95°C for 10 minutes before loading. Next the gel was transferred to a 0.2µm nitrocellulose membrane (Bio-Rad; #1620112) or an activated 0.2µm PVDF membrane (Millipore;#ICEQ00010). Membranes were blocked with 5% milk in PBS-T (0.1%) for 1 hour. For protein detection the following antibodies were used: anti-tdp-43 (abnova; # H00023435-M01); 1:1000 in 5% milk, anti-tubulin (Sigma Aldrich; # T6074); 1:10.000 in 5% milk. Primary antibodies were incubated overnight at 4°C. Incubation with the secondary antibody was at a 1:10 000 dilution for 1 hour at room temperature. Goat anti-mouse IgG (H+L)-HRP conjugate (Bio-Rad; #170-6516) and goat anti-Rabbit IgG (H+L)-HRP conjugate (Bio-Rad; #170-6515). The antibody binding was visualized using Amersham ECL Prime Western Blotting Detection Reagent (GE healthcare; #RPN2236) and imaged with the ImageQuant LAS400 Imaging unit (GE Healthcare).

### Quantitative PCR

To assess the gene expression levels (see Table 2), total RNA from day 1 adult C. elegans was isolated using TRizol (Invitrogen; #15596018) according manufacturer’s protocol. The RNA quality and concentration were assessed with a NanoDrop 2000 spectrophotometer (Thermo Scientific). From 1µg total RNA, cDNA was made using the RevertAid H Minus First Strand cDNA Synthesis kit (Thermo Scientific; #K1632), using random hexamer primers. The Quantitative real-time PCR was performed with a Roche LightCycler 480 Instrument II (Roche diagnostics), using 2µl of 10 times diluted cDNA. To detect the cDNA amplification, SYBR green dye (Bio-Rad; #172-5125) was used. The following PCR program was used: 95°C for 10 minutes, followed by 40 cycles at 95 °C for 15 seconds, 60 °C for 30 seconds and 72 °C for 15 seconds, the program ended with 95°C for 5 seconds, 65 °C for 1 minute and 97 °C. Relative transcript levels were quantitated using a standard curve of pooled cDNA samples. Expression levels were normalized against pmp-3; an endogenous reference gene.

**Table 2:**
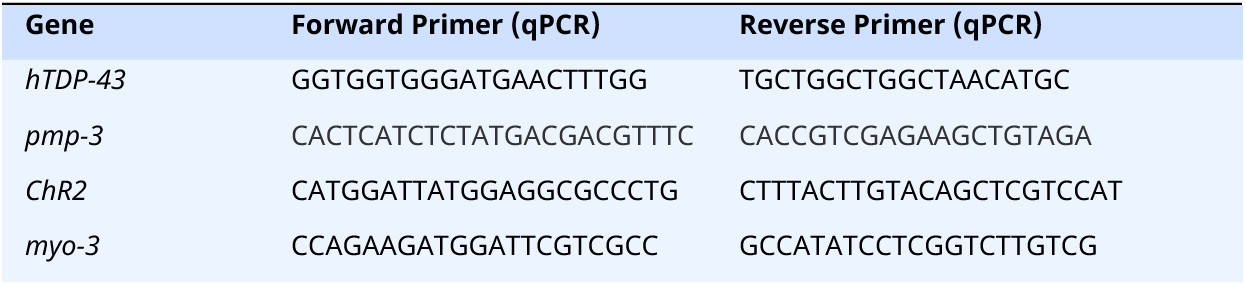
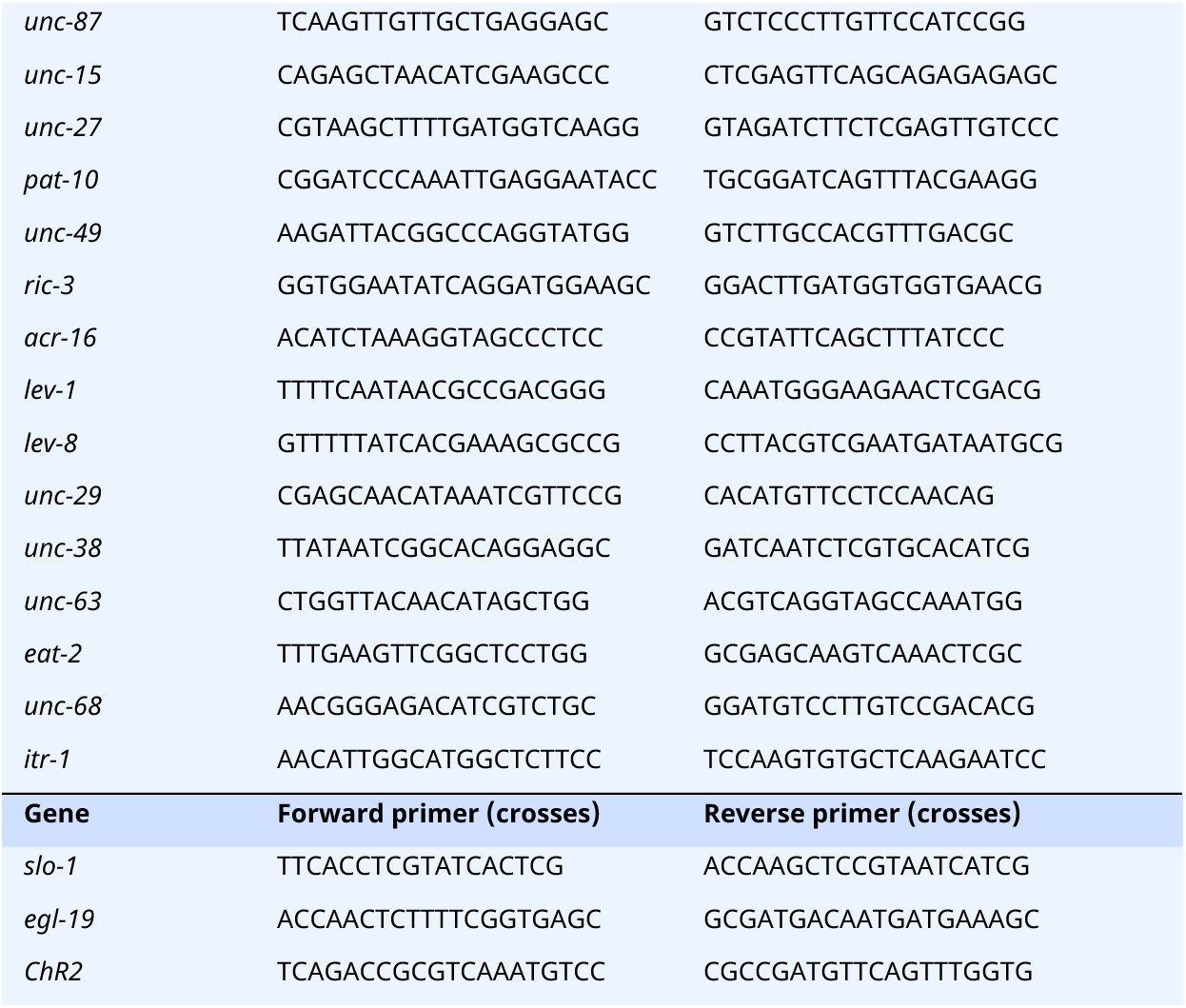
Primers.

### Tierpsy tracker and phenomics

#### Generating phenotypic profile

On the day of tracking, five day 1 adults were picked from the incubated plates to each of the imaging plates (see below) and allowed to habituate for 30 min before recording for 15 min. Imaging plates were 35 mm plates with 3.5 mL of low-peptone (0.013% Difco Bacto) NGM agar (2% Bio/Agar, BioGene) to limit bacteria growth. Imaging plates were seeded with 50 μl of a 1:10 dilution of OP50 in M9 the day before tracking and left to dry overnight with the lid on at room temperature. A total of 10 worms per conditions were recorded. For the analysis, worms were segmented, tracked, and skeletonized using Tierpsy Tracker^34, 44^. Motion, path, and posture features were extracted and a subset of 256 features were selected strains based on their usefulness in classifying a previously analyzed set of mutant worms^34, 44^.

#### Computational approach

The phenotypic profile of hTDP-43 worms was compared with a database of previously analyzed mutants ^35^. These mutants were reanalyzed according to the latest Tierpsy 256 and represent 274 unique (mutated) genes. The function enrichGO of the Bioconductor R package clusterProfiler^151^ (version 3.14.3) was used to test whether certain gene ontology categories were enriched or depleted amongst the unique genes in the Tierpsy mutants (settings: OrgDb=org.Ce.eg.db, version 3.10.1;^152^, keyType=WORMBASE, universe=all analyzed genes, qvalueCutoff=0.05, minGSSize=1, maxGSSize=100000). GO-terms from the “Biological Process”-aspect with a q-value<0.001 and covering less than 2% of the background list were determined to be significant and specific for our screening dataset. GO-terms were further grouped based on overlapping gene content. Using the hierarchical clustering with optimal leaf ordering from the R package seriation^153, 154^. GO-terms were ordered based on similar gene content and subsequently assigned to the same cluster if they shared 50% or more of the same genes with the nearest GO-term, resulting in 29 GO-clusters. GO-terms were considered enriched for mutant hits if the hits ratio within the term exceeded the hits ratio within the overall screening dataset background by a fold change of 2 or more.

### Crawling and thrashing

For crawling ability, we recorded the free crawling of worms on empty NGM agar plates for 30 seconds (unless stated differently) with the WF-NTP platform and analyzed the crawling speed and generated crawling maps with the WF-NTP software^155^. The thrashing frequency was acquired by transferring worms to an empty NGM agar plate flooded with M9 and subsequent recording for 30 seconds. The WF-NTP software was used to derive the thrashing frequency, but also changes in eccentricity. In addition, since uncoordinated worms do skew centroid-based trackers sometimes in our hands, we recounted always (at least) one experiment manually by randomly counting at least 15 worms per movie.

### Paralysis

At day 4 after synchronization (D1-adulthood), 50 worms per condition (unless stated differently) were transferred to 6-cm NGM plates containing FUDR (10 worms per plate). Plates were kept at 20 or 25 °C, as described in the figures. Animals were tested for paralysis every day by tapping their nose/tail with a platinum wire as described previously^156^. Worms that failed to show a touch-response (i.e. moving their nose, but not their body) were scored as paralyzed. Worms that did not move, did not show a touch-response and had no pharyngeal pumping were considered death and were excluded from the assay.

### Contraction index and length measurements

A high-resolution microscopy system was used to image the worms, consisting of an Olympus SZ51 microscope coupled with an IphoneX via a Carson HookUpz 2.0. smartphone adapter. Images of different conditions were always taken with the same optical and digital magnification. The animal width and length were subsequently determined using ImageJ software (NIH Image, Bethesda, MD, USA). The length was measured using a segmented-line line fit from nose to tail and the width was measured just posterior to the vulva. The contraction index was calculated by dividing the width of the worm by its length.

### Aldicarb and levamisole assay

#### Using paralysis

Aldicarb and levamisole (Sigma-Aldrich) plates were made by adding aldicarb or levamisole directly to unseeded NGM plates, giving a final concentration of 10 or 100μM for levamisole and 0.5 or 1mM for aldicarb. Drug plates were left at room temperature for 4-6 hours to allow for the drugs to equilibrate into the agar. In each experiment, 50-75 worms were placed on drug plates and prodded every 20 minutes over a 140-minute period for aldicarb and a 110-minute period for levamisole to determine if they retained the ability to move. Worms that failed to respond at all after two harsh touches were classified as paralyzed. Each experiment was triplicated. Where indicated, animals were pretreated for 2 hours with 2 μg/ml phorbol 12-myristate 13-acetate (PMA) – Sigma-Aldrich: P8139, 2 μg/ml 4*a*-phorbol 12-myristate 13-acetate (4*a*-PMA) – Sigma-Aldrich: P148, 5 mM arecoline hydrobromide – abcam: ab141044, 100 μM methiothepin mesylate salt – Sigma-Aldrich: M149, 0.4 mg/ml serotonin – Sigma-Aldrich: H7752 or 50 μM mianserin hydrochloride – Sigma-Aldrich: M2525.

#### Using length

Assays were performed as described in^157^, but with some significant changes. For aldicarb: synchronized worm populations were pipetted (∼50-70 worms) at D1 of adulthood onto 3-cm NGM agar plates. Each condition as plated double. Then either 250 μM aldicarb or dH_2_O was directly applied onto the plates and a reference image was taken of the worms (T=0). Subsequently, pictures were taken at 60-minute intervals over a set time course of 5 h. For levamisole: worms were pipetted at unseeded NGM plates and 100 μM levamisole or dH_2_O was directly applied onto the 3-cm plates. Recording started immediately after soaking worms in the liquid and pictures were taken at 20-seconds intervals over a set time course of 5 minutes. Worm lengths were determined using ImageJ software. Pictures were binarized (maxEntropy or mean) and skeletonized in such a way that the background was reduced and only worms remained visible. The length of the skeleton was used to define the length of the worm (area-selection tool). Individual worm lengths whilst on aldicarb or levamisole were calculated, and represented as a fraction of worm length compared to time-matched controls.

### PTZ assay

6-well plates NGM agar plates (3 mL each well) were prepared before the experiments and kept at 4°C. For the induction of head-bobbing convulsions, 100 mg/mL PTZ (Sigma-Aldrich, P6500-25G) was prepared freshly on the day and added to each well, giving a final concentration of 10mg/mL PTZ. Plates were left at room temperature for ∼2 hours to allow for the drugs to equilibrate into the agar. Around 30 adult D1 worms were added to each well and filmed with a microscope-smartphone set-up (see *contraction index*) after 30 and 60 minutes.

### Compound screen and rescreen

Adult D1 worms were washed with M9 and approximately 30 worms were transferred per well of 6-well plates. For the explorative screen, each 6-well plate was filled with 5 different concentrations of a specific compound in a log-dose scale and a control-solvent that was similar to the diluting agent of the matched compound (see Table 3), to establish dose-response and time-response relationships. A total of 16 compounds were screened and all compounds were either dissolved in DMSO or water. Plates were covered with parafilm to prevent evaporation and plates were incubated at room temperature at a shaker of 50 rpm. Movies were recorded after 30, 60, and 120 minutes. Except for iberiotoxin treated worms, which were recorded only after 30 minutes. The movies were analyzed by manual counting (at least 20 worms per well were scored) and dose-response curves were generated. The explorative screen was performed twice. Compounds that increased the thrashing frequency to a level that exceeded the 95% bootstrapped confidence (R = 1000, BCa-method) interval of the untreated worms and had a dose-response relationship of R^2^>0.3 were considered as potential hits. Hits were three times retested at a single time-interval of 120 minutes and with one pre-established concentration in the same way as for the explorative screen. Every compound had its own untreated control in a matching solvent, so intra-conditional comparisons could be made. All compounds were prepared freshly before each experiment and all movies were recorded at a framerate of 20 fps (30s total).

**Table 3:** Compounds.

| Compounds | Concentrations explorative screen (μM) | Solvent |
| --- | --- | --- |
| Nemadipine-A (Abcam, ab145991) | 300-100-30-10-3-0 | 2% DMSO |
| Bay K 8644 (Tocris, 1544) | 100-30-10-3-1-0 | 1% DMSO |
| Nefiracetam (Tocris, 2851) | 100-30-10-3-1-0 | 1% DMSO |
| Nifedipine (Sigma-Aldrich, N7634) | 1000-300-100-30-10-0 | 1% DMSO |
| Retigabine (Supelco, 90221) | 1000-300-100-30-10-0 | 1% DMSO |
| Linopirdine dihydrochloride (Tocris, 1999) | 300-100-30-10-3-0 | Water |
| Cilnidipine (Sigma-Aldrich, C1493) | 100-30-10-3-1-0 | 3% DMSO |
| Quinidine (Tocris, 4108) | 1000-300-100-30-10-0 | 1% DMSO |
| Lidocaine (Tocris, 3057) | 1000-300-100-30-10-0 | 1% DMSO |
| Carbamazepine (Sigma-Aldrich, C4024) | 300-100-30-10-3-0 | 1% DMSO |
| 4-Aminopyridine (Sigma-Aldrich, 275875) | 1000-300-100-30-10-0 | Water |
| Clofilium tosylate (Sigma-Aldrich, C2365) | 1000-300-100-30-10-0 | Water |
| Arecoline hydrobromide (abcam, ab141044) | 100-30-10-3-1-0 (mM*) | Water |
| Ethoxsumide (Sigma-Aldrich, E7138) | 10-3-0 (mM*) | Water |
| Iberitoxin (Sigma-Aldrich, I5904) | 1 -0.3-0.1 -0.03 -0.01 | Water |
| Veratidine (Tocris, 2918) | 100-30-10-3-1-0 | 3% DMSO |

### Pretreatment pharmacological assays

Worms were incubated with the compound for either 1 hour in liquid (30 μM muscimol – abcam: ab120094, 300 μM tubacurarine hydrochloride pentahydrate – Sigma-Aldrich: T2379, 100 μM dantrolene sodium salt – Sigma-Aldrich: D9175) or 2 hours on solid substrates (2 μg/ml phorbol 12-myristate 13-acetate (PMA) – Sigma-Aldrich: P8139, 2 μg/ml 4*a*-phorbol 12-myristate 13-acetate (4*a*-PMA) – Sigma-Aldrich: P148, 5 mM arecoline hydrobromide – abcam: ab141044, 100 μM methiothepin mesylate salt – Sigma-Aldrich: M149, 0.4 mg/ml serotonin – Sigma-Aldrich: H7752 or 50 μM mianserin hydrochloride – Sigma-Aldrich: M2525). Subsequently, worms were recorded in liquid (i.e. either the compound or added M9) with the WF-NTP platform and analyzed with the associated software. For muscimol and tubacurarine dose-response curves were created running from 1000-300-100-30-10-0 μM and the concentration with the maximized effect was picked for further experimentation. For combination treatments with arecoline (e.g. nemadinpine-A and dantrolene), worms were first treated for an hour with the specified compound and subsequently incubated for 30 minutes with 5 mM arecoline.

### RNAi experiments

RNAi experiments were performed on NGM agar plates containing 1 mM isopropylthio-β-D-galactoside (IPTG) and 50 mg/mL ampicillin that were seeded with RNAi bacteria. The following RNAi’s were tested: *unc-68, unc-63, acr-16, lev-1, itr-1,* and *egl-19.* Except for *egl-19*, all RNAi were derived from the Ahringer library^158, 159^. *egl-19* was amplified with forward primer: GAAGAACCGCGGGATATCCTCGTCGTTGCAGTATC (SacII site) and reverse primer: GCCGCCTCTAGAAAAAGTGTTCGAATTCCTTCTCC (Xbal site) and cloned into a L4440 vector before being transformed into Ht115 bacteria. Worms were kept from L1 to D1 at the RNAi plates and were subsequently tested in 12-wells plate for the ability to thrash after pretreatment for 1 h with 5 mM arecoline or water (control). Thrashing frequency was only scored of those conditions that had a clear knockdown of the gene of interest.

### Optogenetics

For optogenetic experiments, transgenic worms were cultivated in the dark at 20 °C on NGM plates with or without 0.2 mM all-trans retinal (ATR) added to the OP50. A final concentration of 0.2 mM ATR was obtained by mixing 0.4 μl of a 100 mM ATR stock solution in ethanol (Sigma-Aldrich) with the 200 μL E. coli that was spread on each 6 cm plate. For single-worm illumination, worms were freely moving on unseeded 3-cm NGM plates and kept in frame by manual location while videos were recorded with a microscope-smartphone set-up (see *contraction index*). For ‘short-term’ experiments related to changes in body length, worms were exposed to a 3-3-3 lightening regime, in which light was OFF for 3 seconds, followed by light ON for 3 seconds and light OFF for another 3 seconds. Light intensity was always 1.6 mW/mm^2^ unless stated differently. Changes in length were assessed with a perimeter or midline approach in ImageJ^160^. For multiple-worm illumination in relation to motility, worms were collected in M9 buffer and plated on an empty 3-cm plate that was flooded with 1.5 mL M9. Worms were recorded with the WF-NTP set-up at a framerate of 20 fps and a light intensity of 1.0 mW/mm^2^. We generated separate movies for light OFF (30 seconds) and light ON (30 seconds).

### Phalloidin staining and confocal imaging

Synchronized worms were washed of NGM agar plates, pellets were resuspended in PBS and put on ice. Next, tubes were submerged and snap frozen in liquid nitrogen to freeze crack the worms in suspension. Subsequently, worms were fixed in cold acetone for 5 minutes at −20°C. The acetone was removed and worms were washed once with PBS before adding a Texas Red-X phalloidin (ThermoFisher Scientific #T7471) staining solution (1:10 dilution of 400X stock solution in DMSO with PBS). Worms were stained in the dark for 30 minutes at room temperature. Finally, the staining solution was removed and worms were kept in the dark at 4°C in PBS until imaging. For the confocal imaging worms were placed on a glass slide with agar pad and covered with a cover slip. Confocal Z-stack images were made of the body wall muscles in the region between the pharynx and vulva. All imaging was performed using a Leica SP8X DLS confocal microscope (ex. 595nm, em. 600-789nm) with a 40x oil immersion.

### Nervous system imaging

#### Preparation

10-20 adult worms of each genotype were placed on a lawn of OP50 bacteria and allowed to lay embryos for two hours. After two hours the adult worms were removed and the embryos transferred to a 20 °C or to a 25 °C incubator and raised for 72 hours. At the end of this time the adult worms were transferred into a small volume of M9 culture medium and fixed using a “hot fix” protocol. Roughly fifty worms were picked into a small volume (∼25 microliters) of M9 in a 1.5 mL Eppendorff tube and then 300 uL of hot 4% PFA (55C) was added to the worms. The Eppendorf tube containing worms and fixative was placed at 55 °C for 5 minutes. After the incubation, the worms were three times washed with M9 and spun down in a tabletop microcentrifuge (∼15 seconds at 2,000 x g). After the final wash most of the supernatant was removed and the fixed worms were stored at 4 C in the dark for up to one week.

#### Imaging

To prepare fixed worms for imaging, as much of the supernatant as possible was removed (down to ∼5-10 microliters) and 10 microliters of Vectashield added. The worms were taken up in Vectashield, dropped onto a glass slide, and covered with a glass coverslip. The slide was sealed with nail polish. Fixed worms were imaged on a Zeiss LSM 880 confocal microscope using 20x and 63x objectives. Fixed worms were taken and imaged blind to their identity; after image acquisition, all images of all worms for a given set of four conditions (TDP-43 expressing and control, 20 and 25 °C, for a given set of neurons of interest) were mixed using ImageJ for blind analysis and then scored for the phenotypes of interest. For the synapses, the dorsal side of the worm was imaged.

### Electrophysiology

Electrophysiological methods were as previously described^54^ with the following modifications: Following worm stabilization and dissection, ventral body wall muscle cells were recorded in the whole-cell voltage-clamp mode (holding potential −60 mV) using an EPC-10 patch-clamp amplifier and digitized at 1 kHz. The extracellular solution consisted of (in mM): NaCl 150; KCl 5; CaCl_2_ 5; MgCl_2_ 4, glucose 10; sucrose 5; HEPES 15 (pH 7.4, ∼340mOsm). The patch pipette was filled with (in mM): KCl 120; KOH 20; MgCl_2_ 4; (*N*-tris[Hydroxymethyl] methyl-2-aminoethane-sulfonic acid) 5; CaCl_2_ 0.25; Na^2^ATP 4; sucrose 36; EGTA 5 (pH 7.2, ∼315mOsm). To isolate spontaneous inhibitory postsynaptic currents 10^-4^ M d-tubocurarine (dTBC) was added to the extracellular solution to block acetylcholine receptors. Evoked responses were stimulated with a 2ms depolarizing 20 V pulse delivered via a loose patch pipette placed on the anterior ventral nerve cord using. Data were acquired using Pulse software (HEKA, Southboro, Massachusetts, US) and subsequently analyzed and graphed using Pulsefit (HEKA), Mini Analysis (Synaptosoft Inc., Decatur, Georgia, US) and Igor Pro (Wavemetrics, Lake Oswego, Oregon, US).

### Statistics and visualization

Statistical analyses were done in R and Graphpad Prism 9. The used statistical tests can be found in the different figures and are based on several criteria. In short, (log-)normality tests were always performed on the collected data to test for gaussian distributions (e.g. Shapiro-Wilk). Based on the distribution of the data the appropriate test was selected (parametric or non-parametric). When more than two groups were compared, multiple-testing correction was always applied. When inequality of variance was expected between two groups (based on the experimental design) student’s t-test were always performed with a Welsch’s correction (The Welch test must be specified as part of the experimental design, and not decided upon *a posterori;* ^161, 162^). Post-hoc testing after a one- or two-way ANOVA was only performed when the initial test gave significant results. All experiments were replicated three times, unless stated differently. *: p ≤ 0.05, **: p ≤ 0.01, ***: p ≤ 0.001. All data was visualized with R or Graphpad Prism 9 and color-adjusted in Adobe Illustrator.

## Acknowledgments

We thank Christopher Link for kindly sharing the hTDP-43 and control strains *elegans*. We thank Alexander Gottschalk for sharing several strains and fruitful tips and trick on optogenetics with *C. elegans*. Some strains were provided by the CGC, which is funded by NIH Office of Research Infrastructure Programs (P40 OD010440). This project was funded by a BCN-Brain grant (to M.K.) and a grant of Stichting ALS Nederland (686990) (to E.A.A.N. and M.K.), HORIZON EUROPE Marie Sklodowska-Curie Actions, HealthAge (812830) (to E.A.A.N), Medical Research Council (UK), (MC-A658-5TY30) (to A.E.X.B), Ubbo Emmius Fonds Alumni Chapter Gooische Groningers (to E.A.A.N), The Dutch Research Council (NWO) (Aspasia (015.014.005), (to E.A.A.N).

## Author contributions

LJ, MK and RW performed the computational analyses. AEXB and AP performed the Tierpsy experiments and the clustering with the Tierpsy mutants. MK designed and performed most of the phenotypic, pharmaceutical, optogenetic and genetic assays with extensive help of LG and input of EN. RIS and LG crossed all the mutant strains and performed the qPCRs and western blots. WH cloned RNAi constructs. NO and JR performed the confocal imaging and electrophysiology respectively with the help and input of EJ and MK. MK analyzed all data, created all the figures and performed statistical analyses. MK, EN, JR, EJ, were extensively involved in discussing experiments and follow-up steps. MK wrote the manuscript with help of EN, LJ and LG.

---

**Supplementary Figure 1:**
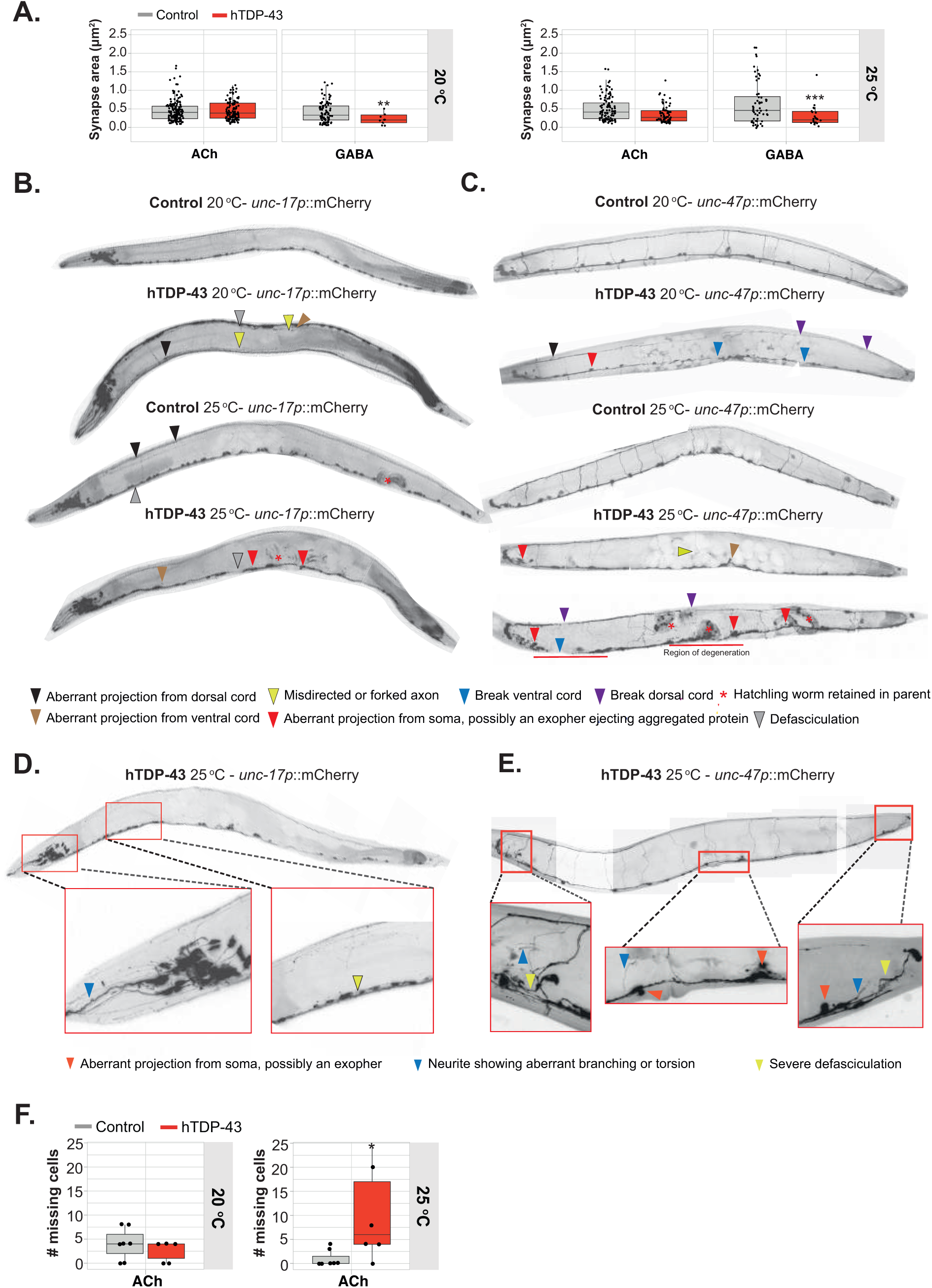
Neurodegenerative signs in hTDP-43 worms. **A)** Quantification of the area of cholinergic and GABAergic synapses in hTDP-43 worms. *n =* 10-150, Inter-strain Student’s t-tests with Welsch correction were performed. **B)** Representative images of control and hTDP-43 worms expressing mCherry in cholinergic neurons. Arrows refer to specific phenotypes. **C)** Representative images of control and hTDP-43 worms expressing mCherry in GABAergic neurons. Arrows refer to specific phenotypes. **D)** Representative images of hTDP-43 worms expressing mCherry in cholinergic neurons. Arrows refer to specific phenotypes. **E)** Representative images of hTDP-43 worms expressing mCherry in GABAergic neurons. Arrows refer to specific phenotypes. **F)** Normally, 45 ACh cell bodies were observed in the ventral nerve cord. In some animals stereotypical groups of soma did not express mCherry, consistent with the loss of the extrachromosomal array in that lineage. *n* ≥ 5, Intra-temperature Mann-Whitney U-tests were performed per phenotype. * = p<0.05, ** = p<0.01, ** = p<0.001.

**Supplementary figure 2:**
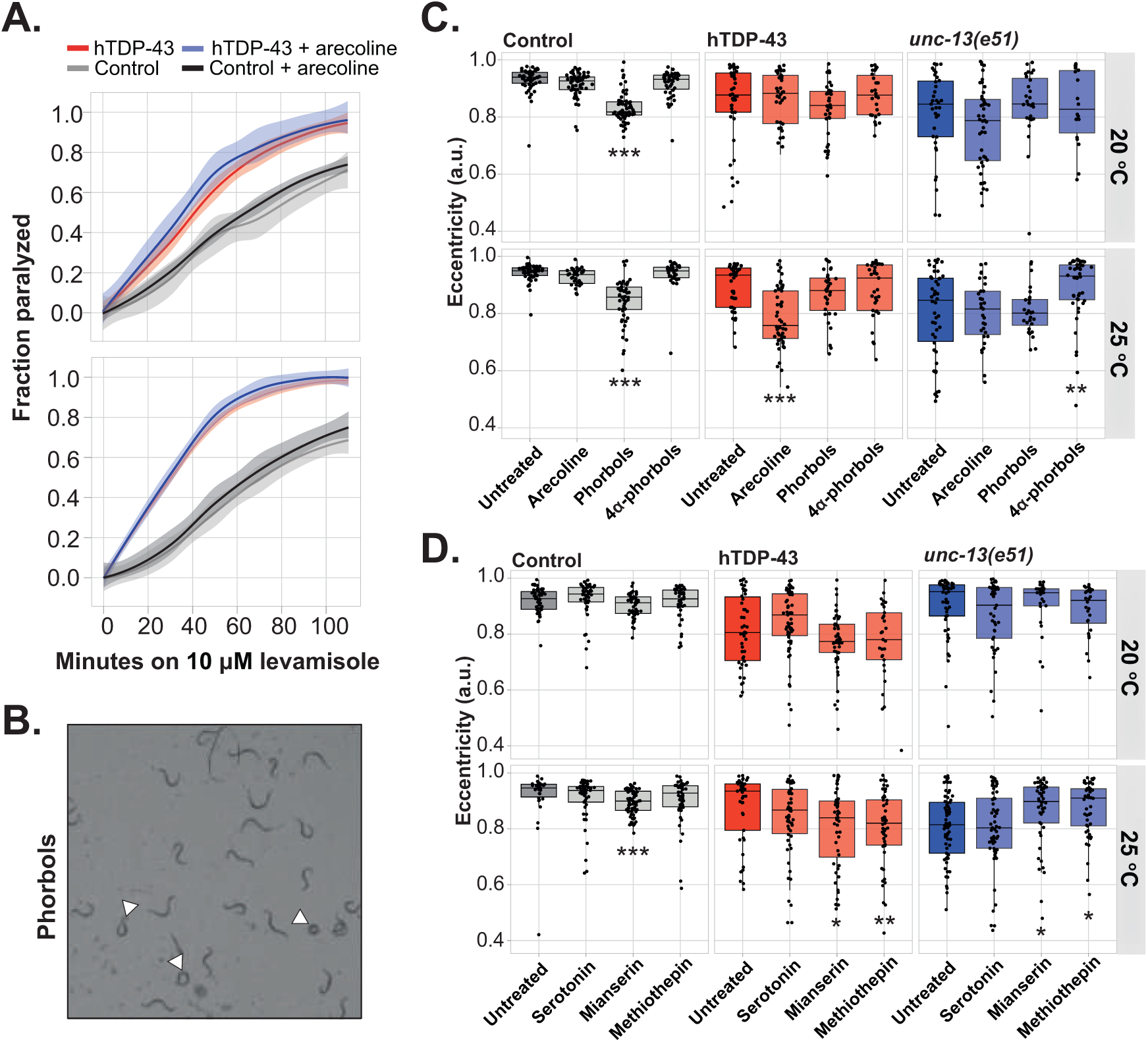
Enhancing cholinergic signaling with aldicarb ameliorates movement phenotypes in hTDP-43 worms that have severe cholinergic and GABAergic signaling defects. **A)** Pretreatment with arecoline has only consequences at the pre-synaptic level (aldicarb) but not at the post-synaptic level (levamisole). The average of three experiments with 10 uM levamisole is shown. **B)** Example image of treated control worms. Clearly, the eccentricity of control worms decreases (roundness increases) when treated with phorbol-esters. **C)** Quantification of eccentricity after pretreatment with the depicted compounds. Only intra-strain comparisons were performed, *n* = 30-60, Kruskal-Wallis for control at 20 °C and 25 °C (p<0.001), for hTDP-43 at 20 °C (n.s.) and 25°C (p < 0.001) and for *unc-13*(e51) 20 °C (n.s.) and 25 °C (p<0.001) with post-hoc Dunn’s. One representative experiment of a triplicate is shown. **D)** Quantification of eccentricity after pretreatment with the depicted compounds. Only intra-strain comparisons were performed, *n* = 30-80, Kruskal-Wallis for control at 20 °C (p = 0.0026) and 25 °C (p<0.001), for hTDP-43 at 20 °C (p = 0.0051) and 25°C (p = 0.0104) and for *unc-13*(e51) 20 °C (n.s.) and 25 °C (p = 0.0018) with post-hoc Dunn’s. One representative experiment of a triplicate is shown. Transparent areas in line-graphs represent 95% confidence interval * = p<0.05, ** = p<0.01, ** = p<0.001.

**Supplementary Figure 3:**
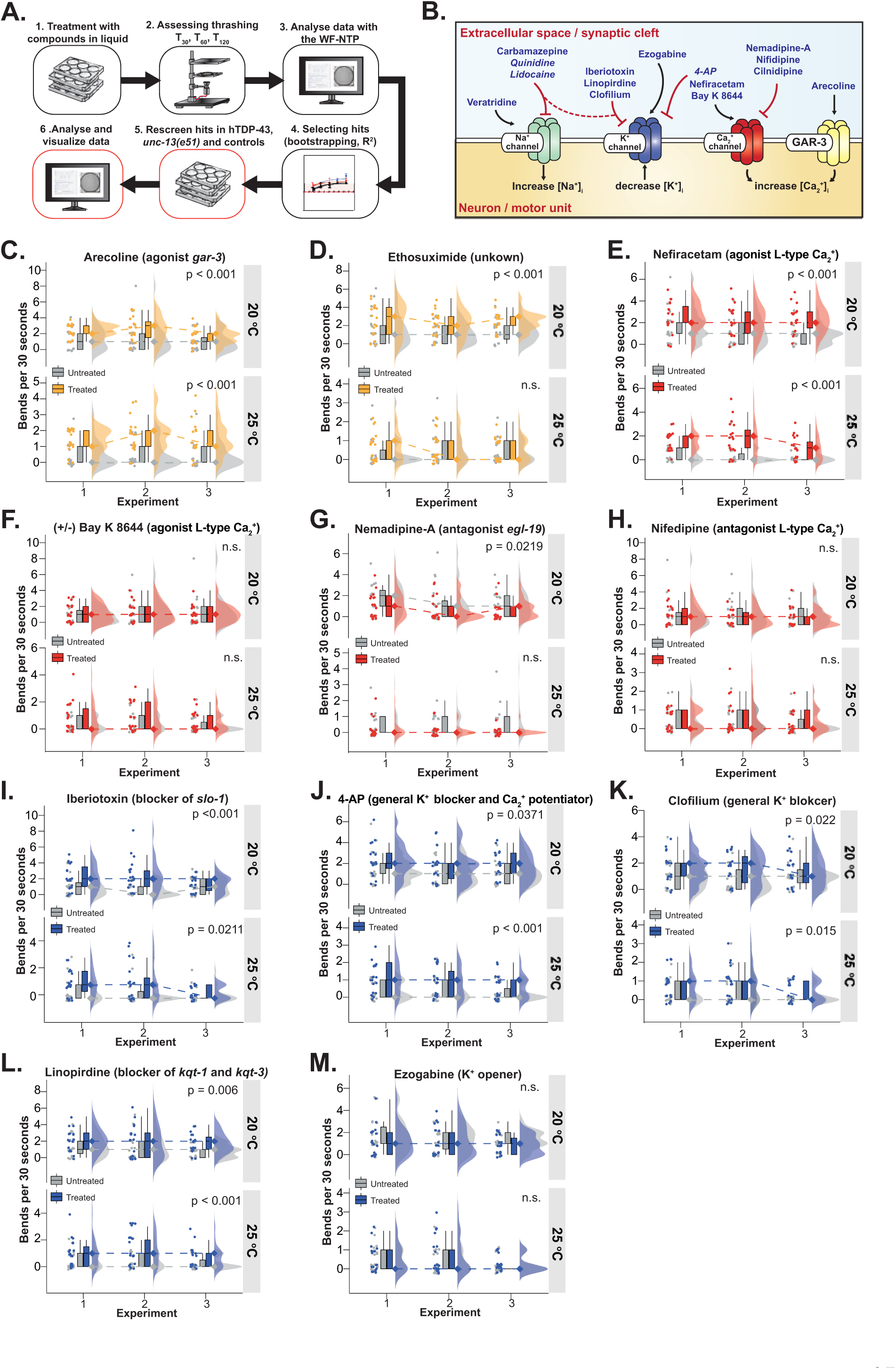
An informed compound screens reveals the potency of calcium channel-activators and potassium-channel blockers to increase the mobility of hTDP-43 worms. **A)** Pipeline used to perform the explorative compound screen and to select and rescreen hits. **B)** Overview of the compounds used in the explorative screen. The colors of the ion channels correspond with the colors in D-I. Names that are written in italics refer to compounds with dual actions. **C)** Thrashing ability of hTDP-43 worms treated with Arecoline, *n =* 20 per experiment, two-way ANOVA at 20 °C and 25 °C (treatment: p<0.001, interaction, experiment: n.s.). **D)** Thrashing ability of hTDP-43 worms treated with Ethosuximide, *n =* 20 per experiment, two-way ANOVA at 20 °C (treatment: p<0.001; interaction, experiment: n.s.) and 25 °C (treatment, interaction, experiment: n.s.). **E)** Thrashing ability of hTDP-43 worms treated with Nefiracetam, *n =* 20 per experiment, two-way ANOVA at 20 °C and 25 °C (treatment: p<0.001; interaction, experiment: n.s.) **F)** Thrashing ability of hTDP-43 worms treated with Bay K 8644, *n =* 20 per experiment, two-way ANOVA at 20 °C and 25 °C (treatment, interaction, experiment: n.s.). **G)** Thrashing ability of hTDP-43 worms treated with Nemadipine A, *n =* 20 per experiment, two-way ANOVA at 20 °C (treatment: p = 0.0219; interaction, experiment: n.s.) and 25 °C (treatment, interaction, experiment: n.s.). **H)** Thrashing ability of hTDP-43 worms treated with Nifedipine, *n =* 20 per experiment, two-way ANOVA at 20 °C and 25 °C (treatment, interaction, experiment: n.s.). **I)** Thrashing ability of hTDP-43 worms treated with Iberiotoxin, *n =* 20 per experiment, two-way ANOVA at 20 °C (treatment: p<0.001; interaction, experiment: n.s.) and 25 °C (treatment: p = 0.0021; interaction: n.s., experiment: 0.0211). **J)** Thrashing ability of hTDP-43 worms treated with 4-AP, *n =* 20 per experiment, two-way ANOVA at 20 °C (treatment: p = 0.0371; interaction, experiment: n.s.) and 25 °C (treatment: p<0.001; interaction, experiment: n.s.). **K)** Thrashing ability of hTDP-43 worms treated with Clofilium, *n =* 20 per experiment, two-way ANOVA at 20 °C (treatment: p=0.0215; interaction, experiment: n.s.) and 25 °C (treatment: p = 0.0145; interaction: n.s., experiment: 0.0179). **L)** Thrashing ability of hTDP-43 worms treated with Linopirdine, *n =* 20 per experiment, two-way ANOVA at 20 °C (treatment: p = 0.0064; interaction, experiment: n.s.) and 25 °C (treatment: p < 0.001; interaction, experiment: n.s.). **M)** Thrashing ability of hTDP-43 worms treated with Ezogabine, *n =* 20 per experiment, two-way ANOVA at 20 °C and 25 °C (treatment, interaction, experiment: n.s.). Dotted lines show the interaction effects per compound. The p-values in figure D-I represent the main treatment effect. * = p<0.05, ** = p<0.01, ** = p<0.001. Colors of the plots corresponds to the schematic in B.

**Supplementary Figure 4:**
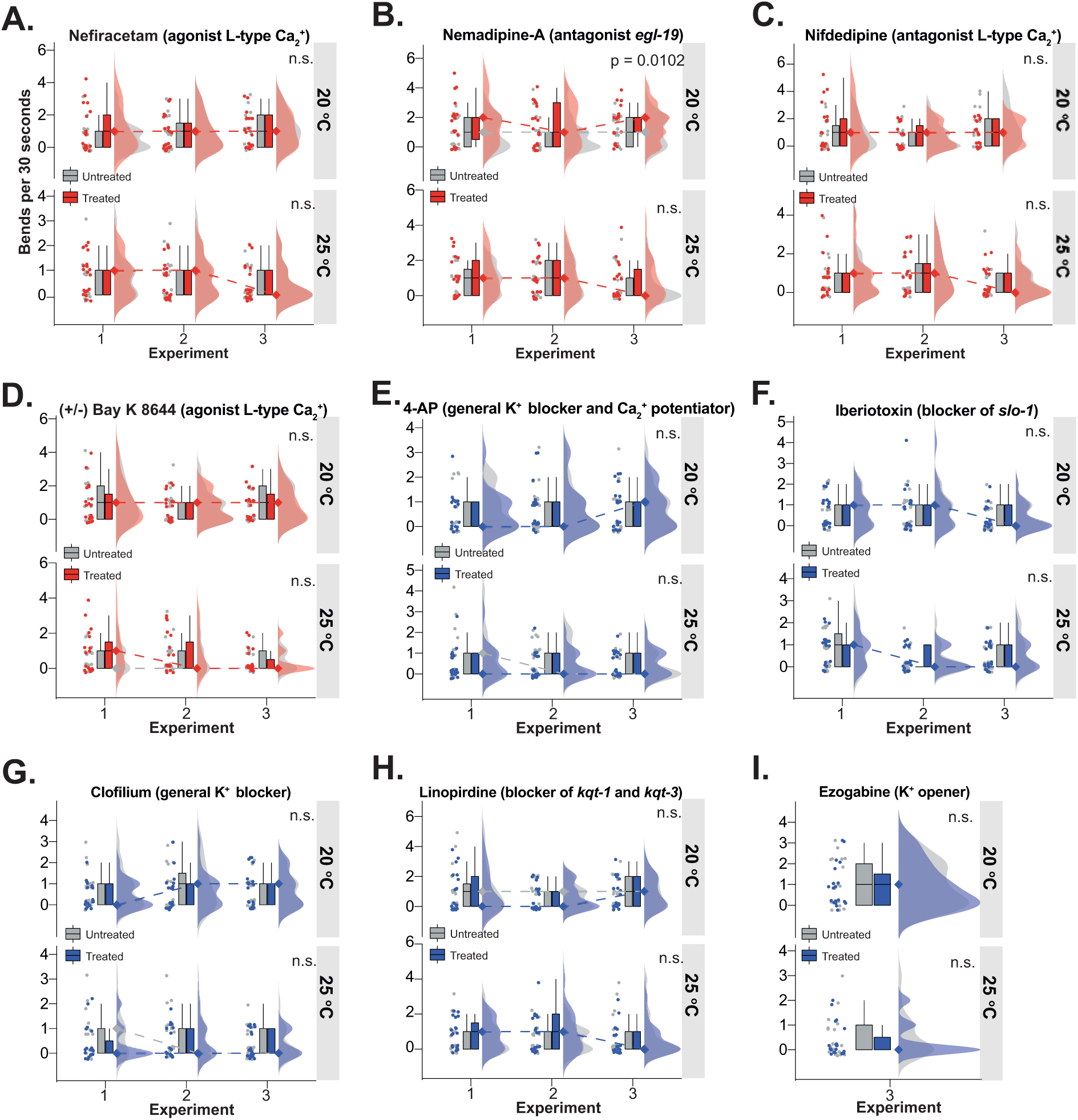
Ion-channel modulators have no effect in *unc-13*(e51) worms. **A)** Thrashing ability of control worms treated with Nefiracetam, *n =* 20 per experiment, two-way ANOVA at 20 °C and 25 °C (treatment, interaction, experiment: n.s.). **B)** Thrashing ability of hTDP-43 worms treated with Nemadipine A, *n =* 20 per experiment, two-way ANOVA at 20 °C (treatment: p = 0.0102; interaction, experiment: n.s.) and 25 °C (treatment, interaction, experiment: n.s.). **C)** Thrashing ability of hTDP-43 worms treated with Nifedipine, *n =* 20 per experiment, two-way ANOVA at 20 °C and 25 °C (treatment, experiment, interaction: n.s.). **D)** Thrashing ability of hTDP-43 worms treated with Bay K 8644, *n =* 20 per experiment, two-way ANOVA at 20 °C and 25 °C (treatment, interaction, experiment: n.s.). **E)** Thrashing ability of hTDP-43 worms treated with 4-AP, *n =* 20 per experiment, two-way ANOVA at 20 °C and 25 °C (treatment, interaction, experiment: n.s.). **F)** Thrashing ability of hTDP-43 worms treated with Iberiotoxin, *n =* 20 per experiment, two-way ANOVA at 20 °C and 25 °C (treatment, interaction, experiment: n.s.). **G)** Thrashing ability of hTDP-43 worms treated with Clofilium, *n =* 20 per experiment, two-way ANOVA at 20 °C and 25 °C (treatment, interaction, experiment: n.s.). **H)** Thrashing ability of hTDP-43 worms treated with Linopirdine, *n =* 20 per experiment, two-way ANOVA at 20 °C and 25 °C (treatment, interaction, experiment: n.s.). **I)** Thrashing ability of hTDP-43 worms treated with Ezogabine, *n =* 20, Mann-Whitney U test at 20 °C and 25 °C. Colors of the plots corresponds to the schematic in Supplementary Figure 3B, annotated p-values correspond to the main treatment effect in each comparison. * = p<0.05, ** = p<0.01, ** = p<0.001

**Supplementary Figure 5:**
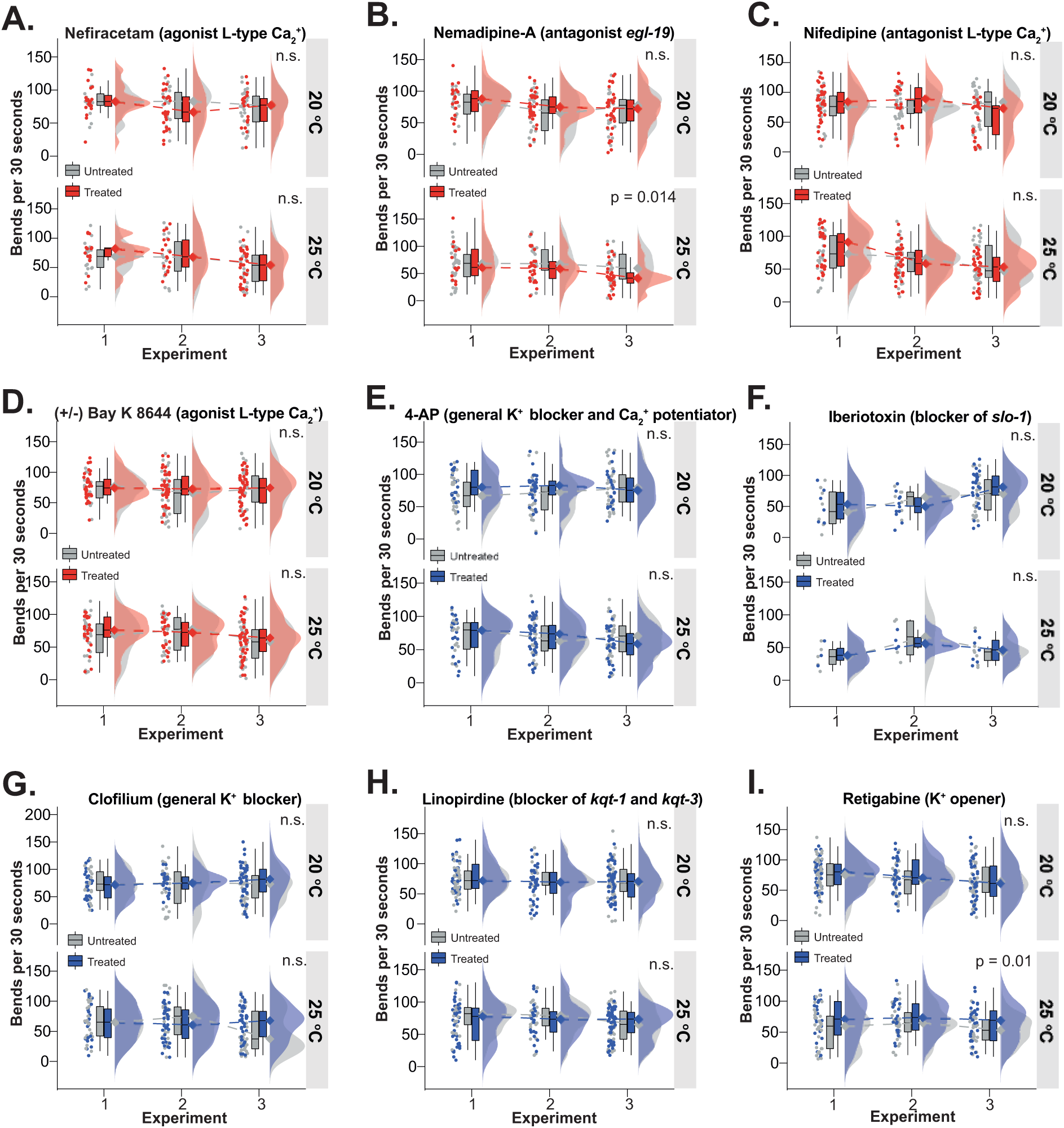
Ion-channel modulators have no effects on motility in control worms. Thrashing ability of control worms treated with Nefiracetam, *n =* 20 per experiment, two-way ANOVA at 20 °C and 25 °C (treatment, interaction, experiment: n.s.). **B)** Thrashing ability of hTDP-43 worms treated with Nemadipine A, *n =* 20 per experiment, two-way ANOVA at 20 °C (treatment, interaction, experiment: n.s.) and 25 °C (treatment, interaction: n.s., experiment: p = 0.0135). **C)** Thrashing ability of hTDP-43 worms treated with Nifedipine, *n =* 20 per experiment, two-way ANOVA at 20 °C (treatment: n.s., interaction: p = 0.0269; experiment: p = 0.0179) and 25 °C (treatment, interaction: n.s., experiment: p < 0.001). **D)** Thrashing ability of hTDP-43 worms treated with Bay K 8644, *n =* 20 per experiment, two-way ANOVA at 20 °C and 25 °C (treatment, interaction, experiment: n.s.). **E)** Thrashing ability of hTDP-43 worms treated with 4-AP, *n =* 20 per experiment, two-way ANOVA at 20 °C and 25 °C (treatment, interaction, experiment: n.s.). **F)** Thrashing ability of hTDP-43 worms treated with Iberiotoxin, *n =* 20 per experiment, two-way ANOVA at 20 °C (treatment, interaction: n.s., experiment: p = 0.0042) and 25 °C (treatment, interaction, experiment: n.s.). **G)** Thrashing ability of hTDP-43 worms treated with Clofilium, *n =* 20 per experiment, two-way ANOVA at 20 °C and 25 °C (treatment, interaction, experiment: n.s.). **H)** Thrashing ability of hTDP-43 worms treated with Linopirdine, *n =* 20 per experiment, two-way ANOVA at 20 °C and 25 °C (treatment, interaction, experiment: n.s.). **I)** Thrashing ability of hTDP-43 worms treated with Ezogabine, *n =* 20 per experiment, two-way ANOVA at 20 °C (treatment, interaction: n.s., experiment: p = 0.0349) and 25 °C (treatment: p = 0.0096; interaction, experiment: n.s.). Colors of the plots corresponds to the schematic Supplementary Figure 3B, annotated p-values correspond to the main treatment effect in each comparison. * = p<0.05, ** = p<0.01, ** = p<0.001

**Supplementary Figure 6:**
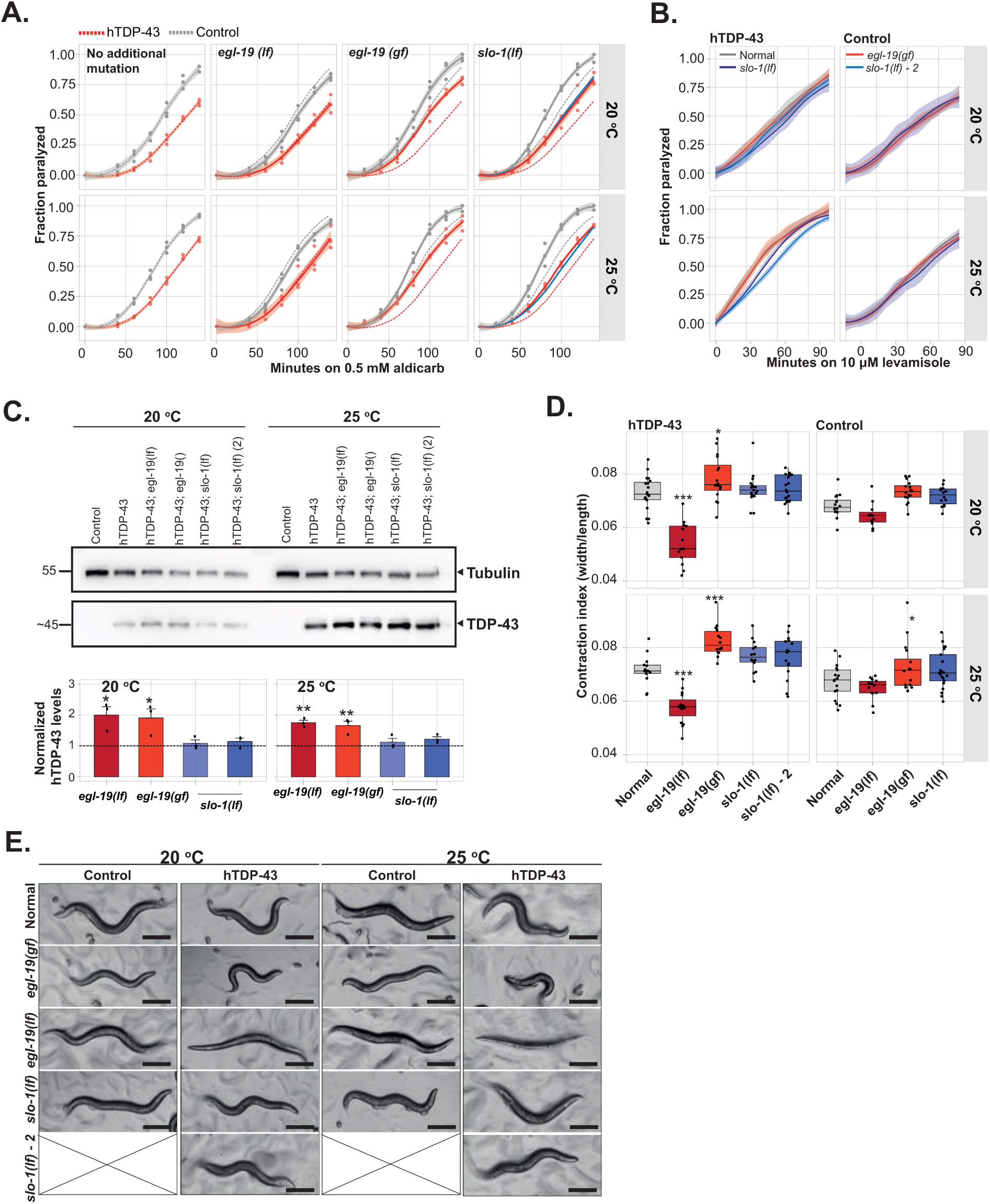
Gain-of-function of *egl-19* and loss-of-function of *slo-1* increase acetylcholine release presynaptically. **A)** Cholinergic neuronal transmission in the different depicted double mutants was measured by determining the onset of paralysis on 0.5M aldicarb plates. *n = 3*. Transparent areas show the S.E.M. **B)** Postsynaptic sensitivity to 10 μM levamisole expressed in the onset of paralysis. *n = 3*. Transparent areas show the S.E.M. **C)** Western blot of hTDP-43 and its quantification, *n = 3*, one-way ANOVA for 20 °C (p = 0.0072) and 25 °C (p<0.001) with post-hoc Dunnettt’s. **D)** Contraction index (width/length) of control and hTDP-43 worms with additional mutations in the background, *n = 14-19*, two-way ANOVA for 20 °C (mutation, interaction: p<0.001, genotype: n.s.) and 25 °C (genotype, interaction: p<0.001, mutation: p = 0.0013) with post-hoc Dunnett’s. **E)** Representative images of the different double mutants. Scale bar = 200 microns. Error bars represent the S.E.M. * = p<0.05, ** = p<0.01, ** = p<0.001.

**Supplementary Figure 7:**
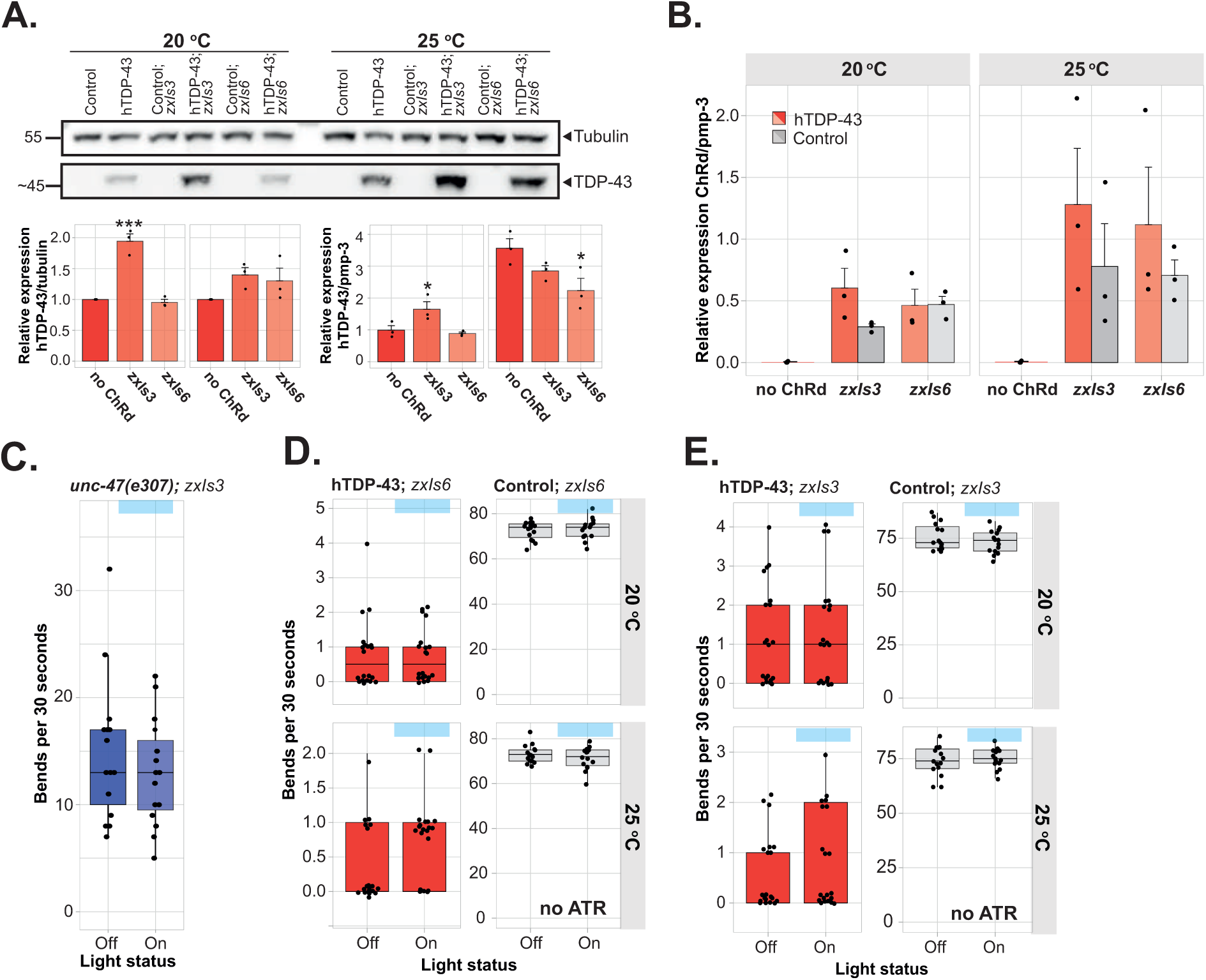
Enhanced movement after photo-stimulation requires GABAergic signaling and ATR treatment. **A)**. Western blot and accompanying quantification of hTDP-43 levels in worms expressing ChRd either in their cholinergic of GABAergic neurons, *n =* 3, one-way ANOVA at 20 °C (p<0.001) and 25 °C (n.s.) with post-hoc Dunnett’s compared. Relative expression of hTDP-43, normalized by *pmp-3*, at different temperatures as measured with a quantitative PCR is also shown, *n* = 3, one-way ANOVA at 20 °C (p = 0.038) and 25 °C (n.s.) with post-hoc Dunnett’s. **B)** Relative expression of ChRd, normalized by *pmp-3*, at different temperatures as measured with a quantitative PCR, *n* = 3, two-way ANOVA at 20 °C (genotype, interaction: n.s., construct: p<0.001) and 25 °C (genotype, interaction: n.s., construct: p = 0.0068) with post-hoc Sidak’s. **C)** Thrashing ability, in bends per 30 seconds, of *unc-47(e307)* worms grown with ATR and either not exposed to blue light (‘off’) or exposed to blue light (‘on’). Worms express ChRd in their GABAergic neurons. *n* = 15-20, two-tailed unpaired Student’s t-test (n.s.). **D)** Thrashing ability, in bends per 30 seconds, of worms not grown on ATR and either not exposed to blue light (‘off’) or exposed to blue light (‘on’). Worms express ChRd in their cholinergic neurons. Only intra-strain comparisons were performed, *n* = 15-20, Mann-Whitney U test for hTDP-43 at 20 °C and 25 °C (n.s.) and two-tailed unpaired Student’s t-test for control worms at 20 °C and 25 °C (n.s.). One representative experiment of a triplicate is shown. **E)** Thrashing ability, in bends per 30 seconds, of worms not grown on ATR and either not exposed to blue light (‘off’), or exposed to blue light (‘on’). Worms express ChRd in their GABAergic neurons. Only intra-strain comparisons were performed, *n* = 15-20, Mann-Whitney U test for hTDP-43 at 20 °C, 25 °C and controls at 20 °C (n.s.) and two-tailed unpaired Student’s t-test for control worms at 25 °C (n.s.). One representative experiment of a triplicate is shown. * = p<0.05, ** = p<0.01, ** = p<0.001.

**Supplementary Figure 8:**
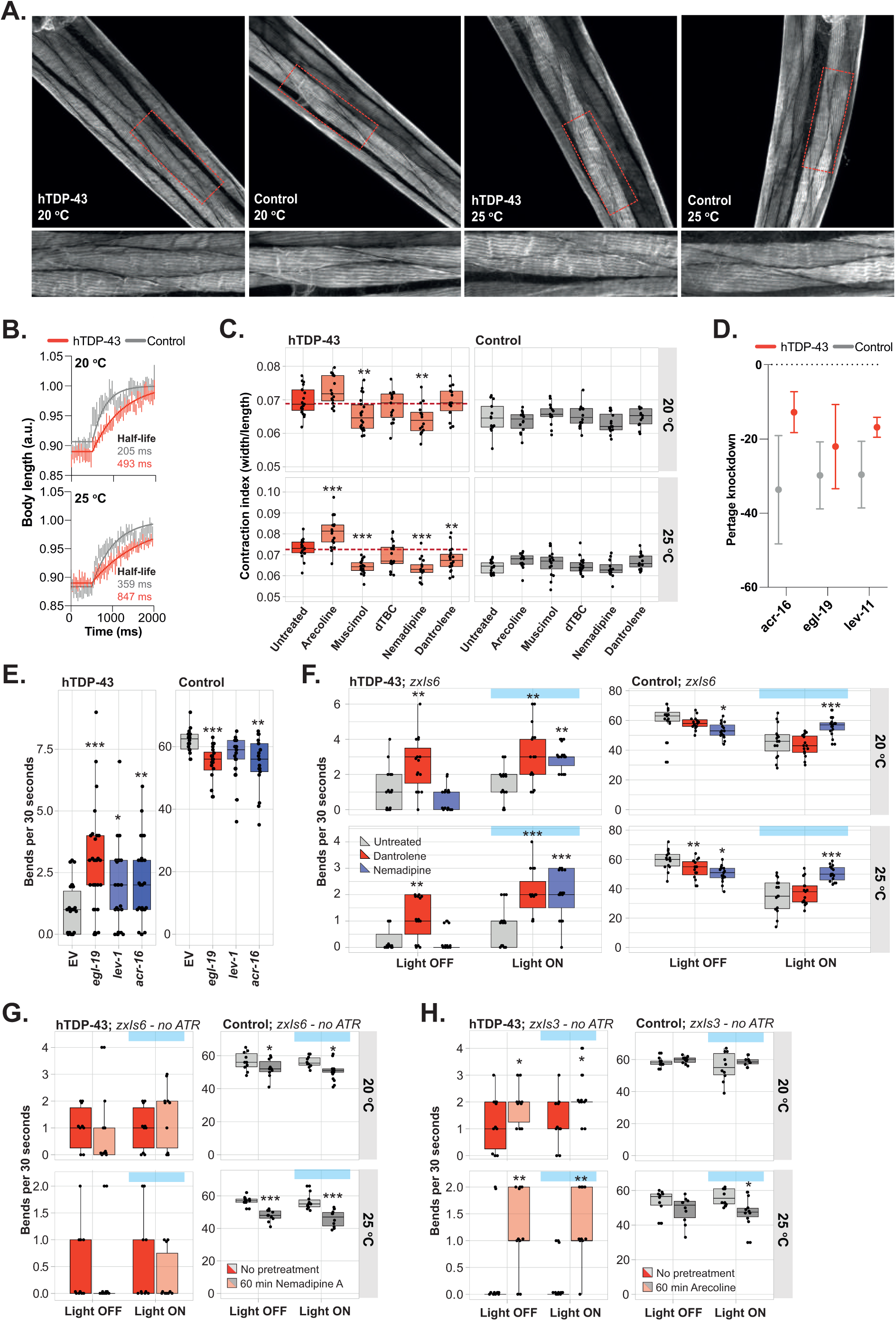
Balancing the muscle excitation-to-inhibition ratio with optogenetics and RNAi. **A)** Representative images of muscle fibers stained with phalloidin in hTDP-43 and control worms. **B)** The normalized body-length after maximal contraction (optogenetically induced, see Figure 7). Extra-sum-of-squares F test, examining K (length ms^-1^) and the half-time for both curves at 20 °C and 25 °C (p<0.001). **C)** Contraction index (width/length) of control and hTDP-43 worms after treatment with different compounds, *n = 13-19*, two-way ANOVA for 20 °C (genotype, treatment, interaction: p<0.001) and 25 °C (genotype, interaction: p<0.001, mutation: p = 0.0013) with post-hoc Dunnett’s. **D)** Knockdown of *egl-19*, *acr-16,* and *lev-1* in hTDP-43 and control worms at 20 °C. **E)** Thrashing frequency after treatment with different RNAis from L1 in a muscle-specific RNAi background combination with short-term arecoline treatment. Only intra-strain comparisons were performed, *n* = 15-25, Kruskal-Wallis test for hTDP-43 and control worms (p<0.001) with post-hoc Dunn’s. One representative experiment of a duplicate is shown. **F)** Thrashing frequency of worms that received no pretreatment or pretreatment with Dantrolene or Nemadipine A and in which cholinergic neurons were photo-activated. Only intra-strain comparisons were performed, *n* = 15-25, two-way ANOVA for hTDP43 at 20 °C (pretreatment, photo-stimulation: p<0.001, interaction: p = 0.0034, pretreatment: n.s.) and at 25 °C (pretreatment, photo-stimulation: p<0.001, interaction: p = 0.0069, pretreatment: n.s.), for controls at 20 °C and 25 °C (interaction, photo-stimulation: p<0.001, pretreatment: n.s.) with post-hoc Holm-Sidak’s. One representative experiment of a triplicate is shown. **G)** Thrashing frequency of worms grown without ATR that received no pretreatment or pretreatment with Nemadipine A and in which cholinergic neurons were photo-activated. Only intra-strain comparisons were performed, *n* = 10-15, two-way ANOVA for hTDP43 at 20 °C and at 25 °C (pretreatment, photo-stimulation, interaction: n.s.), and for controls at 20 °C (pretreatment: p = 0.0032, photo-stimulation, interaction: n.s.), and 25 °C (pretreatment: p < 0.001, photo-stimulation, interaction: n.s.) with post-hoc Holm-Sidak’s. One representative experiment of a triplicate is shown. **H)** Thrashing frequency of worms grown without ATR that received no pretreatment or pretreatment with arecoline and in which GABAergic neurons were photo-activated. Only intra-strain comparisons were performed, *n* = 10-15, two-way ANOVA for hTDP43 at 20 °C (pretreatment: p = 0.0028, photo-stimulation, interaction: n.s.) and at 25 °C (pretreatment: p < 0.001, photo-stimulation, interaction: n.s.), and for controls at 20 °C (pretreatment, photo-stimulation, interaction: n.s.), and 25 °C (pretreatment: p =0.0043, photo-stimulation, interaction: n.s.) with post-hoc Holm-Sidak’s. One representative experiment of a triplicate is shown. * = p<0.05, ** = p<0.01, ** = p<0.001.

## References

1. Neumann, M., et al. Ubiquitinated TDP-43 in Frontotemporal Lobar Degeneration and Amyotrophic Lateral Sclerosis. Science 314, 130–133 (2006).

2. Arai, T., et al. TDP-43 is a component of ubiquitin-positive tau-negative inclusions in frontotemporal lobar degeneration and amyotrophic lateral sclerosis. Biochemical and biophysical research communications 351, 602–611 (2006).

3. Riku, Y., et al. Differential motor neuron involvement in progressive muscular atrophy: a comparative study with amyotrophic lateral sclerosis. BMJ Open 4, e005213 (2014).

4. Kosaka, T., et al. Primary lateral sclerosis: Upper-motor-predominant amyotrophic lateral sclerosis with frontotemporal lobar degeneration - immunohistochemical and biochemical analyses of TDP-43. Neuropathology 32, 373–384 (2012).

5. Matej, R., Tesar, A. & Rusina, R. Alzheimer’s disease and other neurodegenerative dementias in comorbidity: A clinical and neuropathological overview. Clinical biochemistry 73, 26–31 (2019).

6. Cortese, A., et al. Widespread RNA metabolism impairment in sporadic inclusion body myositis TDP43-proteinopathy. Neurobiol Aging 35, (2014).

7. Nalbandian, A., et al. The Multiple Faces of Valosin-Containing Protein-Associated Diseases: Inclusion Body Myopathy with Paget’s Disease of Bone, Frontotemporal Dementia, and Amyotrophic Lateral Sclerosis. J Mol Neurosci 45, 522–531 (2011).

8. Ling, S., Polymenidou, M. & Cleveland, D. Converging Mechanisms in ALS and FTD: Disrupted RNA and Protein Homeostasis. Neuron 79, 416–438 (2013).

9. Tan, R. H., Ke, Y. D., Ittner, L. M. & Halliday, G. M. ALS/FTLD: experimental models and reality. Acta Neuropathol 133, 177–196 (2017).

10. Renton, A., et al. A Hexanucleotide Repeat Expansion in C9ORF72 Is the Cause of Chromosome 9p21-Linked ALS-FTD. Neuron 72, 257–268 (2011).

11. Da Cruz, S. & Cleveland, D. W. Understanding the role of TDP-43 and FUS/TLS in ALS and beyond. Current opinion in neurobiology 21, 904–919 (2011).

12. Vanden Broeck, L., et al. TDP-43 Loss-of-Function Causes Neuronal Loss Due to Defective Steroid Receptor-Mediated Gene Program Switching in Drosophila. Cell Reports 3, 160–172 (2013).

13. Gordon, D. et al. Single-copy expression of an amyotrophic lateral sclerosis-linked TDP-43 mutation (M337V) in BAC transgenic mice leads to altered stress granule dynamics and progressive motor dysfunction. Neurobiology of disease 121, 148–162 (2019).

14. Fratta, P., et al. Mice with endogenous TDP-43 mutations exhibit gain of splicing function and characteristics of amyotrophic lateral sclerosis. The EMBO Journal 37 (2018).

15. Watanabe, S., et al. ALS-linked TDP-43M337V knock-in mice exhibit splicing deregulation without neurodegeneration. Molecular Brain 13, 1–8 (2020).

16. White, M. A., et al. TDP-43 gains function due to perturbed autoregulation in a Tardbp knock-in mouse model of ALS-FTD. Nature neuroscience 21, 1138 (2018).

17. Osaka, M., Ito, D. & Suzuki, N. Disturbance of proteasomal and autophagic protein degradation pathways by amyotrophic lateral sclerosis-linked mutations in ubiquilin 2. Biochemical and biophysical research communications 472, 324–331 (2016).

18. Brockington, A., et al. Unravelling the enigma of selective vulnerability in neurodegeneration: motor neurons resistant to degeneration in ALS show distinct gene expression characteristics and decreased susceptibility to excitotoxicity. Acta Neuropathol 125, 95–109 (2013).

19. Selvaraj, B. T., et al. C9ORF72 repeat expansion causes vulnerability of motor neurons to Ca2+-permeable AMPA receptor-mediated excitotoxicity. Nature communications 9, 1–14 (2018).

20. Gunes, Z. I., Kan, V. W. Y., Ye, X. & Liebscher, S. Exciting Complexity: The Role of Motor Circuit Elements in ALS Pathophysiology. Frontiers in neuroscience 14, 573 (2020).

21. Seeley, W. W., Crawford, R. K., Zhou, J., Miller, B. L. & Greicius, M. D. Neurodegenerative Diseases Target Large-Scale Human Brain Networks. Neuron 62, 42–52 (2009).

22. Nasseroleslami, B., et al. Characteristic Increases in EEG Connectivity Correlate With Changes of Structural MRI in Amyotrophic Lateral Sclerosis. Cerebral cortex 29, 27–41 (2019).

23. Vucic, S., Nicholson, G. A. & Kiernan, M. C. Cortical hyperexcitability may precede the onset of familial amyotrophic lateral sclerosis. Brain 131, 1540–1550 (2008).

24. McMackin, R., et al. Measuring network disruption in neurodegenerative diseases: New approaches using signal analysis. *Journal of Neurology*, Neurosurgery, and Psychiatry 90, 1011–1020 (2019).

25. Turrigiano, G. G. & Nelson, S. B. Homeostatic plasticity in the developing nervous system. Nature reviews. Neuroscience 5, 97–107 (2004).

26. White, J. G., Southgate, E., Thomson, J. N. & Brenner, S. The Structure of the Nervous System of the Nematode Caenorhabditis elegans. Phil. Trans. R. Soc. Lond. B 314, 1–340 (1986).

27. Fang-Yen, C., Alkema, M. J. & Samuel, A. D. T. Illuminating neural circuits and behaviour in Caenorhabditis elegans with optogenetics. Phil. Trans. R. Soc. B 370, 20140212 (2015).

28. Richmond, J. Synaptic function. WormBook, 1–14 (2005).

29. White, B. H. What genetic model organisms offer the study of behavior and neural circuits. Journal of neurogenetics 30, 54–61 (2016).

30. Yan, G., et al. Network control principles predict neuron function in the Caenorhabditis elegans connectome. Nature 550, 519–523 (2017).

31. Augustin, I., Rosenmund, C., Südhof, T. C. & Brose, N. Munc13-1 is essential for fusion competence of glutamatergic synaptic vesicles. Nature 400, 457–461 (1999).

32. Brose, N., Hofmann, K., Hata, Y. & Südhof, T. C. Mammalian Homologues of Caenorhabditis elegans unc-13 Gene Define Novel Family of C2-domain Proteins. The Journal of biological chemistry 270, 25273–25280 (1995).

33. Richmond, J. E., Davis, W. S. & Jorgensen, E. M. UNC-13 is required for synaptic vesicle fusion in C. elegans. Nature neuroscience 2, 959–964 (1999).

34. Javer, A., et al. An open-source platform for analyzing and sharing worm-behavior data. Nature methods 15, 645–646 (2018).

35. Yemini, E., Jucikas, T., Grundy, L. J., Brown, A. E. X. & Schafer, W. R. A database of Caenorhabditis elegans behavioral phenotypes. Nature methods 10, 877–879 (2013).

36. Ash, P. E. A., et al. Neurotoxic effects of TDP-43 overexpression in C. elegans. Human molecular genetics 19, 3206–3218 (2010).

37. Vaccaro, A., et al. Mutant TDP-43 and FUS Cause Age-Dependent Paralysis and Neurodegeneration in C. elegans. PLoS ONE 7, e31321 (2012).

38. Zhang, T., Hwang, H., Hao, H., Talbot, C. & Wang, J. Caenorhabditis elegans RNA-processing Protein TDP-1 Regulates Protein Homeostasis and Life Span. The Journal of biological chemistry 287, 8371–8382 (2012).

39. Liachko, N. F., Guthrie, C. R. & Kraemer, B. C. Phosphorylation Promotes Neurotoxicity in a Caenorhabditis elegans Model of TDP-43 Proteinopathy. The Journal of neuroscience 30, 16208–16219 (2010).

40. Koopman, M., Güngördü, L., Seinstra, R. I. & Nollen, E. A. A. Neuronal overexpression of human TDP-43 in Caenorhabditis elegans causes a range of sensorimotor phenotypes. microPublication biology (2023).

41. Koopman, M., Güngördü, L., Seinstra, R. I. & Nollen, E. A. A. Neuronal overexpression of hTDP-43 in Caenorhabditis elegans impairs different neuronally controlled behaviors and decreases fecundity. microPublication biology (2023).

42. Koopman, M., Güngördü, L., Seinstra, R. I., Hogewerf, W. & Nollen, E. A. A. Neuronal overexpression of hTDP-43 in Caenorhabditis elegans mimics the cellular pathology commonly observed in TDP-43 proteinopathies. microPublication biology (2023).

43. Koopman, M., Güngördü, L., Seinstra, R. I. & Nollen, E. A. A. Neuronal overexpression of hTDP-43 in Caenorhabditis elegans impairs motor function. microPublication biology (2023).

44. Javer, A., Ripoll-Sánchez, L. & Brown, A. E. X. Powerful and interpretable behavioural features for quantitative phenotyping of Caenorhabditis elegans. Phil. Trans. R. Soc. B 373, 20170375 (2018).

45. Miller, K. G., et al. A Genetic Selection for Caenorhabditis elegans Synaptic Transmission Mutants. Proceedings of the National Academy of Sciences - PNAS 93, 12593–12598 (1996).

46. Mulcahy, B., Holden-Dye, L. & O’Connor, V. Pharmacological assays reveal age-related changes in synaptic transmission at the Caenorhabditis elegans neuromuscular junction that are modified by reduced insulin signalling. Journal of experimental biology 216, 492–501 (2013).

47. Lewis, J. A., Fleming, J. T., Maclafferty, S., Murphy, H. & Wu, C. The levamisole receptor, a cholinergic receptor of the nematode Caenorhabditis elegans. Molecular pharmacology 31, 185–193 (1987).

48. Dabbish, N. S. & Raizen, D. M. GABAergic synaptic plasticity during a developmentally regulated sleep-like state in C. elegans. The Journal of neuroscience 31, 15932–15943 (2011).

49. Wu, C., et al. Enhancing GABAergic Transmission Improves Locomotion in a Caenorhabditis elegans Model of Spinal Muscular Atrophy. eNeuro 5, ENEURO.0289-18.2018 (2018).

50. Chaya, T., et al. A C. elegans genome-wide RNAi screen for altered levamisole sensitivity identifies genes required for muscle function. G3 (Bethesda, Md.) 11 (2021).

51. Williams, S. N., Locke, C. J., Braden, A. L., Caldwell, K. A. & Caldwell, G. A. Epileptic-like convulsions associated with LIS-1 in the cytoskeletal control of neurotransmitter signaling in Caenorhabditis elegans. Human molecular genetics 13, 2043–2059 (2004).

52. Etter, A., et al. Picrotoxin Blockade of Invertebrate Glutamate-Gated Chloride Channels: Subunit Dependence and Evidence for Binding Within the Pore. Journal of neurochemistry 72, 318–326 (1999).

53. Locke, C., et al. Paradigms for Pharmacological Characterization of C. elegans Synaptic Transmission Mutants. Journal of Visualized Experiments 18, 847 (2008).

54. Richmond, J. E. & Jorgensen, E. M. One GABA and two acetylcholine receptors function at the C. elegans neuromuscular junction. Nature neuroscience 2, 791–7 (1999).

55. Melentijevic, I., et al. C. elegans neurons jettison protein aggregates and mitochondria under neurotoxic stress. Nature 542, 367–371 (2017).

56. Chan, J. P., Hu, Z. & Sieburth, D. Recruitment of sphingosine kinase to presynaptic terminals by a conserved muscarinic signaling pathway promotes neurotransmitter release. Genes & development 26, 1070–1085 (2012).

57. Chan, J. P., et al. Extrasynaptic Muscarinic Acetylcholine Receptors on Neuronal Cell Bodies Regulate Presynaptic Function in Caenorhabditis elegans. The Journal of neuroscience 33, 14146–14159 (2013).

58. Lackner, M. R., Nurrish, S. J. & Kaplan, J. M. Facilitation of Synaptic Transmission by EGL-30 G qα and EGL-8 PLCβ : DAG Binding to UNC-13 Is Required to Stimulate Acetylcholine Release. Neuron 24, 335–346 (1999).

59. Maruyama, I. N. & Brenner, S. A phorbol ester/diacylglycerol-binding protein encoded by the unc-13 gene of Caenorhabditis elegans. Proceedings of the National Academy of Sciences - PNAS 88, 5729–5733 (1991).

60. Sassa, T., et al. Regulation of the UNC-18-Caenorhabditis elegans Syntaxin Complex by UNC-13. The Journal of neuroscience 19, 4772–4777 (1999).

61. Richmond, J. E., Weimer, R. M. & Jorgensen, E. M., An open form of syntaxin bypasses the requirement for UNC-13 in vesicle priming. Nature 412, 338–341 (2001).

62. Miller, K. G., Emerson, M. D. & Rand, J. B. Goα and Diacylglycerol Kinase Negatively Regulate the Gqα Pathway in C. elegans. Neuron 24, 323–333 (1999).

63. Hajdu-Cronin, Y. M., Chen, W. J., Patikoglou, G., Koelle, M. R. & Sternberg, P. W. Antagonism between G(o)alpha and G(q)alpha in Caenorhabditis elegans: the RGS protein EAT-16 is necessary for G(o)alpha signaling and regulates G(q)alpha activity. Genes & development 13, 1780–1793 (1999).

64. Nurrish, S., Ségalat, L. & Kaplan, J. M. Serotonin inhibition of synaptic transmission: Galpha(0) decreases the abundance of UNC-13 at release sites. Neuron 24, 231–242 (1999).

65. Wang, Z. Origin of quantal size variation and high-frequency miniature postsynaptic currents at the Caenorhabditis elegans neuromuscular junction. Journal of neuroscience research 88, 3425–3432 (2010).

66. Gierbolini, J., Giarratano, M. & Benbadis, S. R. Carbamazepine-related antiepileptic drugs for the treatment of epilepsy - a comparative review. Expert opinion on pharmacotherapy 17, 885–888 (2016).

67. Franks, C. J., et al. Ionic Basis of the Resting Membrane Potential and Action Potential in the Pharyngeal Muscle of Caenorhabditis elegans. Journal of Neurophysiology 87, 954–961 (2002).

68. Wang, Z., Saifee, O., Nonet, M. L. & Salkoff, L. SLO-1 Potassium Channels Control Quantal Content of Neurotransmitter Release at the C. elegans Neuromuscular Junction. Neuron 32, 867–881 (2001).

69. Wei, A. D., Butler, A. & Salkoff, L. KCNQ-like potassium channels in Caenorhabditis elegans. Conserved properties and modulation. The Journal of biological chemistry 280, 21337–21345 (2005).

70. Risley, M. G., Kelly, S. P., Minnerly, J., Jia, K. & Dawson-Scully, K. egl-4 modulates electroconvulsive seizure duration in C. elegans. Invert Neurosci 18, 8–6 (2018).

71. Ikenaka, K., et al. A behavior-based drug screening system using a Caenorhabditis elegans model of motor neuron disease. Scientific Reports 9, 10104–10 (2019).

72. Liu, H., et al. Spontaneous Vesicle Fusion Is Differentially Regulated at Cholinergic and GABAergic Synapses. Cell Reports 22, 2334–2345 (2018).

73. Jospin, M., Jacquemond, V., Mariol, M., Ségalat, L. & Allard, B. The L-Type Voltage-Dependent Ca2+ Channel EGL-19 Controls Body Wall Muscle Function in Caenorhabditis elegans. The Journal of Cell Biology 159, 337–347 (2002).

74. Yoshii, M., et al. Neuronal Ca2+ Channels and Nicotinic ACh Receptors as Functional Targets of the Nootropic Nefiracetam. Psychogeriatrics 1, 39–49 (2001).

75. Yoshii, M. & Watabe, S. Enhancement of neuronal calcium channel currents by the nootropic agent, nefiracetam (DM-9384), in NG108-15 cells. Brain research 642, 123–131 (1994).

76. Yoshii, M., Watabe, S., Murashima, Y. L., Nukada, T. & Shiotani, T. Cellular Mechanism of Action of Cognitive Enhancers: Effects of Nefiracetam on Neuronal Ca2+ Channels Ser. 14, Lippincott Williams & Wilkins, Hagerstown, MD, 2000).

77. Nimmrich, V. & Gross, G. P/Q-type calcium channel modulators. British journal of pharmacology 167, 741–759 (2012).

78. Davies, A. G., et al. A Central Role of the BK Potassium Channel in Behavioral Responses to Ethanol in C. elegans. Cell 115, 655–666 (2003).

79. Bhattacharjee, A., et al. Slick (Slo2.1), a Rapidly-Gating Sodium-Activated Potassium Channel Inhibited by ATP. The Journal of neuroscience 23, 11681–11691 (2003).

80. De los Angeles Tejada, M., Stolpe, K., Meinild, A. & Klaerke, D.A. Clofilium inhibits Slick and Slack potassium channels. Biologics 6, 465–470 (2012).

81. Scholz, A. Mechanisms of (local) anaesthetics on voltage-gated sodium and other ion channels. Br. J. Anaesth 89, 52–61 (2002).

82. Roy, P. J., et al. A small-molecule screen in C. elegans yields a new calcium channel antagonist. Nature 441, 91–95 (2006).

83. Wong, S. Q., et al. α-Methyl-α-phenylsuccinimide ameliorates neurodegeneration in a C. elegans model of TDP-43 proteinopathy. Neurobiology of Disease 118, 40–54 (2018).

84. Chen, X., et al. Ethosuximide ameliorates neurodegenerative disease phenotypes by modulating DAF-16/FOXO target gene expression. Molecular Neurodegeneration 10, 51 (2015).

85. Tauffenberger, A., Julien, C. & Parker, J. A. Evaluation of longevity enhancing compounds against transactive response DNA-binding protein-43 neuronal toxicity. Neurobiology of aging 34, 2175–2182 (2013).

86. Tiwari, S. K., et al. Ethosuximide Induces Hippocampal Neurogenesis and Reverses Cognitive Deficits in an Amyloid-β Toxin-induced Alzheimer Rat Model via the Phosphatidylinositol 3-Kinase (PI3K)/Akt/Wnt/β-Catenin Pathway. The Journal of biological chemistry 290, 28540–28558 (2015).

87. Rand, J. B. & Johnson, C. D. Genetic pharmacology: interactions between drugs and gene products in Caenorhabditis elegans. Methods in Cell Biology 48, 187–204 (1995).

88. Roy, P. J., et al. A predictive model for drug bioaccumulation and bioactivity in Caenorhabditis elegans. Nature Chemical Biology 6, 549–557 (2010).

89. Partridge, F. A., Tearle, A. W., Gravato-Nobre, M. J., Schafer, W. R. & Hodgkin, J. The C. elegans glycosyltransferase BUS-8 has two distinct and essential roles in epidermal morphogenesis. Developmental Biology 317, 549–559 (2008).

90. Zheng, S., Ding, A., Li, G., Wu, G. & Luo, H. Drug absorption efficiency in Caenorhabditis elegans delivered by different methods. PLoS ONE 8, e56877 (2013).

91. Li, G., et al. Genetic and pharmacological interventions in the aging motor nervous system slow motor aging and extend life span in C. elegans. Science Advances 5, eaau5041 (2019).

92. Laine, V., Zhan, H. & Segor, J. R. Hyperactivation of L-type voltage-gated Ca.sup.2+ channels in Caenorhabditis elegans striated muscle can result from point mutations in the IS6 or the IIIS4 segment of the alpha1 subunit. Journal of experimental biology 217, 3805 (2014).

93. Liu, Q., Kidd, P. B., Dobosiewicz, M. & Bargmann, C. I. C. elegans AWA Olfactory Neurons Fire Calcium-Mediated All-or-None Action Potentials. Cell 175, 57–70.e17 (2018).

94. Chen, B., Liu, P., Zhan, H. & Wang, Z. -. Dystrobrevin Controls Neurotransmitter Release and Muscle Ca2+ Transients by Localizing BK Channels in Caenorhabditis elegans. The Journal of neuroscience 31, 17338–17347 (2011).

95. Tong, X., et al. Retrograde Synaptic Inhibition Is Mediated by α-Neurexin Binding to the α2δ Subunits of N-Type Calcium Channels. Neuron 95, 326–340.e5 (2017).

96. Husson, S. J., Gottschalk, A. & Leifer, A. M. Optogenetic manipulation of neural activity in C. elegans: From synapse to circuits and behaviour. Biology of the cell 105, 235–250 (2013).

97. Liewald, J. F., et al. Optogenetic analysis of synaptic function. Nature methods 5, 895–902 (2008).

98. Zhang, F., et al. Multimodal fast optical interrogation of neural circuitry. Nature 446, 633–639 (2007).

99. Fleming, J. T., et al. Caenorhabditis elegans Levamisole Resistance Genes lev-1, unc-29, and unc-38 Encode Functional Nicotinic Acetylcholine Receptor Subunits. The Journal of neuroscience 17, 5843–5857 (1997).

100. Culetto, E., et al. The Caenorhabditis elegans unc-63 gene encodes a levamisole-sensitive nicotinic acetylcholine receptor alpha subunit. The Journal of biological chemistry 279, 42476–42483 (2004).

101. Touroutine, D., et al. acr-16 Encodes an Essential Subunit of the Levamisole-resistant Nicotinic Receptor at the Caenorhabditis elegans Neuromuscular Junction. The Journal of biological chemistry 280, 27013–27021 (2005).

102. Maryon, E. B., Coronado, R. & Anderson, P. unc-68 Encodes a Ryanodine Receptor Involved in Regulating C. elegans Body-Wall Muscle Contraction. The Journal of Cell Biology 134, 885–893 (1996).

103. Lee, R. Y., Lobel, L., Hengartner, M., Horvitz, H. R. & Avery, L. Mutations in the alpha1 subunit of an L-type voltage-activated Ca2+ channel cause myotonia in Caenorhabditis elegans. The EMBO journal 16, 6066–6076 (1997).

104. Sun, L., et al. Neuronal Regeneration in C. elegans Requires Subcellular Calcium Release by Ryanodine Receptor Channels and Can Be Enhanced by Optogenetic Stimulation. The Journal of neuroscience 34, 15947–15956 (2014).

105. Hutter, H., Ng, M. & Chen, N. GExplore: a web server for integrated queries of protein domains, gene expression and mutant phenotypes. BMC Genomics 10, 529 (2009).

106. Jiang, G., et al. A Na+/Cl- -coupled GABA transporter, GAT-1, from Caenorhabditis elegans: structural and functional features, specific expression in GABA-ergic neurons, and involvement in muscle function. The Journal of biological chemistry 280, 2065–2077 (2005).

107. Esposito, G., Amoroso, M. R., Bergamasco, C., Di Schiavi, E. & Bazzicalupo, P. The G Protein regulators EGL-10 and EAT-16, the Giα GOA-1 and the Gqα EGL-30 modulate the response of the C. elegans ASH polymodal nociceptive sensory neurons to repellents. BMC Biology 8, 138 (2010).

108. McIntire, S. L., Jorgensen, E., Kaplan, J. & Horvitz, H. R. The GABAergic nervous system of Caenorhabditis elegans. Nature 364, 337–41 (1993).

109. Deng, L., et al. Inhibition Underlies Fast Undulatory Locomotion in Caenorhabditis elegans. eNeuro 8, ENEURO.0241-20.2020 (2021).

110. Nijhout, H. F., Best, J. & Reed, M. C. Escape from homeostasis. Mathematical biosciences 257, 104–110 (2014).

111. Brownstone, R. M. & Lancelin, C. Escape from homeostasis: spinal microcircuits and progression of amyotrophic lateral sclerosis. Journal of neurophysiology 119, 1782–1794 (2018).

112. Masrori, P. & Van Damme, P. Amyotrophic lateral sclerosis: a clinical review. European journal of neurology 27, 1918–1929 (2020).

113. Chiò, A., et al. Prognostic factors in ALS: A critical review. Amyotrophic Lateral Sclerosis and Other Motor Neuron Disorders 10, 310–323 (2009).

114. Riva, N., Agosta, F., Lunetta, C., Filippi, M. & Quattrini, A. Recent advances in amyotrophic lateral sclerosis. J Neurol 263, 1241–1254 (2016).

115. Schultz, J. Disease-modifying treatment of amyotrophic lateral sclerosis. The American journal of managed care 24, S327–S335 (2018).

116. Miller, R. G., Mitchell, J. D. & Moore, D. H. Riluzole for amyotrophic lateral sclerosis (ALS)/motor neuron disease (MND). Cochrane database of systematic reviews 2012, CD001447 (2012).

117. Bensimon, G., Lacomblez, L. & Meininger, V. A controlled trial of riluzole in amyotrophic lateral sclerosis. ALS/Riluzole Study Group. The New England journal of medicine 330, 585–591 (1994).

118. Turner, M. R., PhD et al. Controversies and priorities in amyotrophic lateral sclerosis. Lancet Neurology 12, 310–322 (2013).

119. Mitsumoto, H., Dr, Brooks, B. R., MD & Silani, V., MD. Clinical trials in amyotrophic lateral sclerosis: why so many negative trials and how can trials be improved? Lancet neurology 13, 1127–1138 (2014).

120. Geser, F., et al. Evidence of Multisystem Disorder in Whole-Brain Map of Pathological TDP-43 in Amyotrophic Lateral Sclerosis. Archives of neurology (Chicago) 65, 636–641 (2008).

121. van Es, M. A., Goedee, H. S., Westeneng, H., Nijboer, T. C. W. & van den Berg, Leonard H. Is it accurate to classify ALS as a neuromuscular disorder? Expert review of neurotherapeutics 20, 895–906 (2020).

122. Braak, H., et al. Amyotrophic lateral sclerosis—a model of corticofugal axonal spread. Nature reviews. Neurology 9, 708–714 (2013).

123. Lalancette-Hebert, M., Sharma, A., Lyashchenko, A. K. & Shneider, N. A. Gamma motor neurons survive and exacerbate alpha motor neuron degeneration in ALS. Proceedings of the National Academy of Sciences - PNAS 113, E8316–E8325 (2016).

124. Seki, S., et al. Circuit-Specific Early Impairment of Proprioceptive Sensory Neurons in the SOD1G93A Mouse Model for ALS. The Journal of neuroscience 39, 8798–8815 (2019).

125. Van den Bos, M., et al. Imbalance of cortical facilitatory and inhibitory circuits underlies hyperexcitability in ALS. Neurology 91, e1669–e1676 (2018).

126. Kuo, S., Binder, M. D. & Heckman, C. J. Excessive Homeostatic Gain in Spinal Motoneurons in a Mouse Model of Amyotrophic Lateral Sclerosis. Scientific reports 10, 9049 (2020).

127. Landoni, L. M., Myles, J. R., Wells, T. L., Mayer, W. P. & Akay, T. Cholinergic modulation of motor neurons through the C-boutons are necessary for the locomotor compensation for severe motor neuron loss during amyotrophic lateral sclerosis disease progression. Behavioural brain research 369, 111914 (2019).

128. Perry, S., Han, Y., Das, A. & Dickman, D. Homeostatic plasticity can be induced and expressed to restore synaptic strength at neuromuscular junctions undergoing ALS-related degeneration. Human molecular genetics 26, 4153–4167 (2017).

129. Saxena, S., et al. Neuroprotection through Excitability and mTOR Required in ALS Motoneurons to Delay Disease and Extend Survival. Neuron 80, 80–96 (2013).

130. Delestrée, N., et al. Adult spinal motoneurones are not hyperexcitable in a mouse model of inherited amyotrophic lateral sclerosis. The Journal of physiology 592, 1687–1703 (2014).

131. Brettschneider, J., et al. Stages of pTDP-43 pathology in amyotrophic lateral sclerosis. Annals of neurology 74, 20–38 (2013).

132. Sonobe, Y., et al. A C. elegans model of C9orf72-associated ALS/FTD uncovers a conserved role for eIF2D in RAN translation. Nature communications 12, 6025 (2021).

133. Liachko, N. F., et al. CDC7 inhibition blocks pathological TDP-43 phosphorylation and neurodegeneration. Annals of neurology 74, 39–52 (2013).

134. Lynne Greenup et al. The Tau Tubulin Kinases TTBK1/2 Promote Accumulation of Pathological TDP-43. PLoS genetics 10 (2014).

135. Jablonski, A. M., et al. Loss of RAD-23 Protects Against Models of Motor Neuron Disease by Enhancing Mutant Protein Clearance. The Journal of neuroscience 35, 14286–14306 (2015).

136. Liachko, N. F., et al. The phosphatase calcineurin regulates pathological TDP-43 phosphorylation. Acta Neuropathol 132, 545–561 (2016).

137. Vaccaro, A., et al. Methylene blue protects against TDP-43 and FUS neuronal toxicity in C. elegans and D. rerio. PLoS ONE 7, e42117 (2012).

138. Bose, P., et al. The Novel Small Molecule TRVA242 Stabilizes Neuromuscular Junction Defects in Multiple Animal Models of Amyotrophic Lateral Sclerosis. Neurotherapeutics 16, 1149–1166 (2019).

139. Liachko, N. F., et al. Genome wide analysis reveals heparan sulfate epimerase modulates TDP-43 proteinopathy. PLoS Genetics 15, e1008526 (2019).

140. Wen, Q., et al. Proprioceptive Coupling within Motor Neurons Drives C. elegans Forward Locomotion. *Neuron (Cambridge*, Mass.) 76, 750–761 (2012).

141. Cornelia Bargmann, Mei Zhen & Shangbang Gao. Action potentials drive body wall muscle contractions in Caenorhabditis elegans. Proceedings of the National Academy of Sciences - PNAS 108, 2557–2562 (2011).

142. Menon, P., Kiernan, M. C. & Vucic, S. Cortical hyperexcitability precedes lower motor neuron dysfunction in ALS. Clinical neurophysiology 126, 803–809 (2015).

143. Zhang, W., et al. Hyperactive somatostatin interneurons contribute to excitotoxicity in neurodegenerative disorders. Nature neuroscience 19, 557–559 (2016).

144. D Thielsen, K., et al. The Wobbler Mouse Model of Amyotrophic Lateral Sclerosis (ALS) Displays Hippocampal Hyperexcitability, and Reduced Number of Interneurons, but No Presynaptic Vesicle Release Impairments. (2016).

145. Nihei, K., Mckee, A. C. & Kowall, N. W. Patterns of neuronal degeneration in the motor cortex of amyotrophic lateral sclerosis patients. Acta neuropathologica 86, 55–64 (1993).

146. Nieto-Gonzalez, J. L., Moser, J., Lauritzen, M., Schmitt-John, T. & Jensen, K. Reduced GABAergic Inhibition Explains Cortical Hyperexcitability in the Wobbler Mouse Model of ALS. Cerebral cortex 21, 625–635 (2011).

147. Khademullah, C. S., et al. Cortical interneuron-mediated inhibition delays the onset of amyotrophic lateral sclerosis. Brain 143, 800–810 (2020).

148. Boussicault, L., et al. Combination of acamprosate and baclofen (PXT864) as a potential new therapy for amyotrophic lateral sclerosis. Journal of neuroscience research 98, 2435–2450 (2020).

149. Huang, X., et al. Human amyotrophic lateral sclerosis excitability phenotype screen: Target discovery and validation. Cell reports 35, 109224 (2021).

150. Jorgensen, E. M. & Mango, S. E. The art and design of genetic screens: Caenorhabditis elegans. Nature reviews. Genetics 3, 356–369 (2002).

151. Yu, G., Wang, L., Han, Y. & He, Q. clusterProfiler: an R Package for Comparing Biological Themes Among Gene Clusters. Omics 16, 284–287 (2012).

152. Carlson, M. org.Ce.eg.db: Genome wide annotation for Worm. (2020).

153. Hahsler, M., Hornik, K. & Buchta, C. Getting Things in Order: An Introduction to the R Package seriation. Journal of Statistical Software 25 (2008).

154. Hahsler, M. An experimental comparison of seriation methods for one-mode two-way data. European journal of operational research 257, 133–143 (2017).

155. Koopman, M., et al. Assessing motor-related phenotypes of Caenorhabditis elegans with the wide field-of-view nematode tracking platform. Nature protocols 15, 2071–2106 (2020).

156. Cohen, E., Bieschke, J., Perciavalle, R. M., Kelly, J. W. & Dillin, A. Opposing Activities Protect Against Age-Onset Proteotoxicity. Science 313, 1604–1610 (2006).

157. Mulcahy, B., Holden-Dye, L. & O’Connor, V. Pharmacological assays reveal age-related changes in synaptic transmission at the Caenorhabditis elegans neuromuscular junction that are modified by reduced insulin signalling. Journal of experimental biology 216, 492–501 (2013).

158. Ahringer, J., et al. Functional genomic analysis of C. elegans chromosome I by systematic RNA interference. Nature (London) 408, 325–330 (2000).

159. Ahringer, J., et al. Systematic functional analysis of the Caenorhabditis elegans genome using RNAi. Nature (London) 421, 231–237 (2003).

160. Koopman, M., Janssen, L. & Nollen, E. A. A. An economical and highly adaptable optogenetics system for individual and population-level manipulation of Caenorhabditis elegans. BMC biology 19, 170 (2021).

161. Moser, B. K. & Stevens, G. R. Homogeneity of Variance in the Two-Sample Means Test. The American Statistician 46, 19 (1992).

162. Hayes, A. F. & Cai, L. Further evaluating the conditional decision rule for comparing two independent means. The British journal of mathematical and statistical psychology 60, 217–44 (2007).

