## Supplemental Table 2(2) for "Rebalancing the motor circuit restores movement in a *Caenorhabditis elegans* model for TDP-43-toxicity"

**Supplementary table 1: General information and $R^2$ of the compounds used in our screen.**

| Compound | Target | $R^2$ at 20 °C | | | $R^2$ at 25 °C | | | | | Remarks |
|---|---|---|---|---|---|---|---|---|---|---|
| | | 30 minutes | 60 minutes | 120 minutes | 30 minutes | 60 minutes | 120 minutes | HIT* | Optimal concentration | |
| **Ethoxusimide** | unknown | - | 0.53 | 0.46 | 0.33 | 0 | 1 | n.a. | 10 mM* | Ethosuximide has been shown to confer neuroprotection and life-span extension in several neurodegenerative *C. elegans* models, via a molecular target that still needs to be determined (Wong et al., 2018; Tauffenberger et al., 2013; Tiwari et al., 2015). |
| **Arecoline** | gar-3 | 0.38 | 0.8 | 0.3 | 0.66 | 0.76 | 0.71 | n.a. | 5 mM* | Higher concentrations increase the trashing frequency but also increase the coiling propensity. Lower concentrations limit the coiling behaviour but als induce a smaller increase in trashing frequency. |
| **Nefiracetam** | L-type/N-type $Ca^{2+}$-channel agonist | 0.21 | 0.75 | 0.42 | 0.6 | 0.18 | 0.47 | Yes | **100 uM** | |
| **Bay K 8644** | L-type $Ca^{2+}$-channel agonist | 0.66 | 0.39 | 0.42 | 0.78 | 0.62 | 0.68 | Possible | **100 uM** | Used as a control compound: Bay K 8644 is unable to potentiate current amplitude in *C. elegans* (Jospin et al., 2002) |
| **Nemadipine A** | egl-19 antagonist | 0.71 | 0.52 | 0.59 | 0.37 | 0.37 | 0.37 | No | 300 uM* | Nemadinpine-A elongates hTDP-43 worms, while barely affecting control worms. |
| **Nifidipine** | L-type $Ca^{2+}$-channel antagonist | 0.74 | 0.75 | 0.09 | 0.36 | 0.11 | 0.23 | No | 300 uM* | |
| **Cilnidipine** | L-type/N-type $Ca^{2+}$-channel antagonist | 0.05 | 0.43 | 0.39 | 0.76 | 0.52 | 0.52 | No | - | |
| **Iberiotoxin** | Antagonist of slo-1 | 0.09 | - | - | 0.47 | - | - | Yes | **10 uM** | In the primary screen only 30-min treatment intervals were used. The effect can be enhanced with longer treatment. |
| **Linopirdine** | Antagonist of kqt-1 and kqt-3 | 0.04 | 0.82 | 0.13 | 0.42 | 0.64 | 0.75 | Possible | **100 uM** | |
| **4-AP** | General K+ antagonist and Ca2+ potentiator | 0.52 | 0.05 | 0.24 | 0.5 | 0.68 | 0.57 | Yes | **300 uM** | Found as modifier in a smn-1 screen (Sleigh et al., 2011) |
| **Quinidine** | General $K^+$-channel and $Na^+$-channel blocker | 0 | 0.06 | 0.02 | 0.81 | 0.44 | 0.36 | No | - | |
| **Lidocaine** | General $K^+$-channel and $Na^+$-channel blocker | 0.3 | 0.36 | 0.28 | 0.07 | 0 | 0.02 | No | - | |
| **Clofilium** | $K^+$-channel antagonist | 0.8 | 0.88 | 0.86 | 0.78 | 0.91 | 0.87 | Yes | **30 uM** | At higher concentrations, hTDP-43 worms coil extremely and cannot make appropriate bends anymore |
| **Ezogabine** | $K^+$-channel agonist | 0.17 | 0.16 | 0.4 | 0 | 0.36 | 0.11 | No | 100 uM* | |
| **Carbamazepine** | Voltage-gated Na+ antagonist | 0.12 | 0.03 | 1 | 0 | 0.08 | 1 | No | - | *C. elegans* neurons can signal effectively without classic voltage-gated sodium channels (Goodman et al., 1998; Pierce-Shimomura et al., 2001). In fact, the *C. elegans* genome does not encode any of such a channel gene (Bargmann, 1998). *Used as control.* |
| **Veratridine** | Voltage-gated Na+ modulator | 0.02 | 0.11 | 0.12 | 1 | 0.07 | 0.08 | No | - | *C. elegans* neurons can signal effectively without classic voltage-gated sodium channels (Goodman et al., 1998; Pierce-Shimomura et al., 2001). In fact, the *C. elegans* genome does not encode any of such a channel gene (Bargmann, 1998). *Used as control.* |

*** A compound was considered a hit when it increased the thrashing frequency to a level that exceeded the 95% bootstrapped confidence interval of the untreated worms and had a dose-response relationship of R2>0.3.**


### References:

Wong S.Q., Pontifex M.G., Phelan M.M., Pidathala C., Kraemer B.C., Barclay J.W., Berry N.G., O'Neill P.M., Burgoyne R.D., Morgan A. 2018. α-Methyl-α-phenylsuccinimide ameliorates neurodegeneration in a C. elegans model of TDP-43 proteinopathy. *Neurobiology of Disease.* **118**: 40-54.

Tauffenberger A., Julien C., Parker J.A. 2013. Evaluation of longevity enhancing compounds against transactive response DNA-binding protein-43 neurona toxicity. *Neurobiol. Aging.* **34**: 2175-2182.

Tiwari S.K., Seth B., Agarwal S., Yadav A., Karmakar M., Gupta S.K., Choubey V., Sharma A., Chaturvedi R.K. 2015. Ethosuximide induces hippocampal neurogenesis and reverses cognitive deficits in amyloid- toxin induced Alzheimer's rat model via PI3K/Akt/Wnt/-catenin pathway. J. Biol. Chem. 290: 28540-28558.

Jospin M., Jacquemond V., Mariol M-C., Ségalat L., Allard B. 2002. The L-type voltage-dependent $Ca^{2+}$ channel EGL-19 controls body wall muscle function in *Caenorhabditis elegans*. *J Cell Biol.* **159**(2): 337-348.

Goodmann M.B., Hall D.H., Avery L., Lockery S.R. 1998. Active currents regulate sensitivity and dynamic range in *C. elegans* neurons. *Neuron,* **20**: 763-772.

Pierce-Shimomura J.T., Faumont S., Gaston M.R., Pearson B.J., Lockery S.R. 2001. The homeobox gene lim-6 is required for distinct chemosensory representations in C. elegans. *Nature*, **410**: 694-698.

Bargmann C.I. 1998. Neurobiology of the *Caenorhabditis elegans* genome. *Science*, **282**: 2028-2033.

Sleigh J.N., Buckingham S.D., Esmaeili B., Viswanathan M., Cuppen E., Westlund B.M., Satelle D.B. 2011. A novel Caenorhabditis elegans allele, smn-1(cb131), mimicking a mild form of spinal muscular atrophy, provides a convenient drug screening platform highlighting new and pre-approved compounds. *Hum Mol Genet.* **20**(2): 245-260.
