## Supplemental Table 1(2) for "Rebalancing the motor circuit restores movement in a *Caenorhabditis elegans* model for TDP-43-toxicity"

| Genes_all_mutants | Genes_hTDP-43_hits |
|---|---|
| acr-10 | asic-1 |
| acr-11 | clh-3 |
| acr-14 | egl-11 |
| acr-15 | flp-1 |
| acr-18 | gly-2 |
| acr-19 | hcf-1 |
| acr-2 | ins-27 |
| acr-21 | ins-28 |
| acr-23 | ins-3 |
| acr-3 | ins-4 |
| acr-6 | jnk-1 |
| acr-7 | lite-1 |
| acr-9 | mec-10 |
| asic-1 | mec-14 |
| bas-1 | nca-2 |
| C24G7.1 | nlp-1 |
| cat-2 | nlp-3 |
| cat-4 | osm-9 |
| ced-1 | rab-3 |
| clh-3 | rig-6 |
| daf-3 | sng-1 |
| daf-5 | srp-8 |
| daf-7 | unc-1 |
| dat-1 | unc-10 |
| del-1 | unc-122 |
| dnc-1 | unc-2 |
| dop-1 | unc-3 |
| dop-2 | unc-30 |
| dop-3 | unc-32 |
| dpy-20 | unc-37 |
| eat-16 | unc-4 |
| egl-1 | unc-55 |
| egl-10 | unc-7 |
| egl-11 | unc-75 |
| egl-12 | unc-79 |
| egl-13 | unc-8 |
| egl-14 | unc-80 |
| egl-15 | unc-89 |
| egl-17 | unc-9 |
| egl-18 | vab-7 |
| egl-19 | unc-77 |
| egl-2 | |
| egl-20 | |
| egl-21 | |
| egl-23 | |
| egl-24 | |
| egl-27 | |
| egl-28 | |
| egl-30 | |

egl-31
egl-32
egl-33
egl-36
egl-37
egl-40
egl-42
egl-43
egl-44
egl-46
egl-47
egl-49
egl-5
egl-50
egl-6
egl-7
egl-8
egl-9
flp-1
flp-10
flp-11
flp-12
flp-13
flp-16
flp-17
flp-18
flp-19
flp-20
flp-21
flp-25
flp-28
flp-3
flp-33
flp-6
flp-7
flp-9
flr-1
ftn-2
gar-2
gld-1
gly-2
goa-1
gon-2
gpa-1
gpa-11
gpa-12
gpa-13
gpa-14
gpa-15
gpa-16

gpa-17
gpa-2
gpa-3
gpa-4
gpa-5
gpa-6
gpa-7
gpa-8
gpa-9
gpb-2
gpc-1
gsa-1
hcf-1
him-5
ins-11
ins-15
ins-16
ins-18
ins-22
ins-25
ins-27
ins-28
ins-3
ins-30
ins-31
ins-35
ins-4
jnk-1
lev-1
lev-8
lig-4
lin-39
lite-1
lon-2
lov-1
mec-10
mec-12
mec-14
mec-18
mec-4
mec-7
mir-124
mod-1
mod-5
nca-2
nhr-95
nlp-1
nlp-12
nlp-14
nlp-15

nlp-17
nlp-18
nlp-2
nlp-20
nlp-3
nlp-5
nlp-8
npr-1
npr-10
npr-11
npr-12
npr-13
npr-2
npr-3
npr-4
npr-5
npr-7
npr-8
npr-9
ocr-1
ocr-2
ocr-3
ocr-4
octr-1
odr-3
osm-9
pdl-1
pkc-1
pkg-1
pmk-1
pqn-66
rab-3
rgs-6
ric-19
rig-6
sem-4
ser-1
ser-2
ser-4
ser-5
ser-6
ser-7
sma-2
sma-3
snf-1
snf-10
snf-11
snf-2
snf-4
snf-5

snf-6
snf-7
snf-8
snf-9
sng-1
spe-41
srp-8
sup-9
syg-1
syg-2
tag-336
tbh-1
tdc-1
tmc-1
tom-1
tph-1
trp-1
trp-2
trp-4
trpa-1
tyra-2
tyra-3
unc-1
unc-10
unc-101
unc-103
unc-104
unc-105
unc-108
unc-115
unc-116
unc-118
unc-122
unc-127
unc-14
unc-16
unc-17
unc-18
unc-2
unc-26
unc-29
unc-3
unc-30
unc-31
unc-32
unc-34
unc-37
unc-38
unc-4
unc-40

unc-42
unc-43
unc-44
unc-55
unc-60
unc-63
unc-69
unc-7
unc-75
unc-76
unc-77
unc-79
unc-8
unc-80
unc-86
unc-89
unc-9
unc-98
vab-7
zyg-9
