## Supplemental Table 1(4) for "Rebalancing the motor circuit restores movement in a *Caenorhabditis elegans* model for TDP-43-toxicity"

| ID | Description | GeneRatio | BgRatio | pvalue | p.adjust | qvalue | geneID | Count |
|---|---|---|---|---|---|---|---|---|
| GO:0023052 | signaling | 150/243 | 1936/9281 | 1.05E-44 | 1.00E-41 | 6.11E-42 | acr-2/acr-3/acr-6/acr-7/acr-9/acr-10/[illegible] | 150 |
| GO:0007154 | cell communication | 151/243 | 1964/9281 | 1.06E-44 | 1.00E-41 | 6.11E-42 | acr-2/acr-3/acr-6/acr-7/acr-9/acr-10/[illegible] | 151 |
| GO:0007187 | G protein-coupled receptor signaling pathway, coupled to cyclic nucleotide second messenger | 33/243 | 57/9281 | 2.98E-38 | 1.88E-35 | 1.14E-35 | dop-1/dop-2/egl-30/gar-2/goa-1/gpa-1/[illegible] | 33 |
| GO:0050896 | response to stimulus | 172/243 | 2896/9281 | 1.06E-37 | 5.00E-35 | 3.04E-35 | acr-2/acr-3/acr-6/acr-7/acr-9/acr-10/[illegible] | 172 |
| GO:0065007 | biological regulation | 206/243 | 4273/9281 | 6.08E-37 | 2.30E-34 | 1.40E-34 | acr-2/acr-3/acr-6/acr-7/acr-9/acr-10/[illegible] | 206 |
| GO:0050789 | regulation of biological process | 197/243 | 3937/9281 | 6.90E-36 | 2.17E-33 | 1.32E-33 | acr-2/acr-3/acr-6/acr-7/acr-9/acr-10/[illegible] | 197 |
| GO:0007165 | signal transduction | 132/243 | 1827/9281 | 5.52E-34 | 1.49E-31 | 9.05E-32 | acr-2/acr-3/acr-6/acr-7/acr-9/acr-10/[illegible] | 132 |
| GO:0050795 | regulation of behavior | 43/243 | 152/9281 | 2.88E-33 | 6.80E-31 | 4.13E-31 | daf-3/daf-7/eat-16/egl-6/egl-8/egl-10/[illegible] | 43 |
| GO:0007268 | chemical synaptic transmission | 52/243 | 252/9281 | 8.88E-33 | 1.68E-30 | 1.02E-30 | acr-2/acr-3/acr-6/acr-7/acr-9/acr-10/[illegible] | 52 |
| GO:0098916 | anterograde trans-synaptic signaling | 52/243 | 252/9281 | 8.88E-33 | 1.68E-30 | 1.02E-30 | acr-2/acr-3/acr-6/acr-7/acr-9/acr-10/[illegible] | 52 |
| GO:0099537 | trans-synaptic signaling | 52/243 | 253/9281 | 1.09E-32 | 1.88E-30 | 1.14E-30 | acr-2/acr-3/acr-6/acr-7/acr-9/acr-10/[illegible] | 52 |
| GO:0099536 | synaptic signaling | 52/243 | 254/9281 | 1.35E-32 | 2.12E-30 | 1.29E-30 | acr-2/acr-3/acr-6/acr-7/acr-9/acr-10/[illegible] | 52 |
| GO:0007186 | G protein-coupled receptor signaling pathway | 78/243 | 693/9281 | 2.14E-30 | 3.11E-28 | 1.89E-28 | dop-1/dop-2/eat-16/egl-6/egl-8/egl-10/[illegible] | 78 |
| GO:0051716 | cellular response to stimulus | 141/243 | 2238/9281 | 2.47E-30 | 3.34E-28 | 2.03E-28 | acr-2/acr-3/acr-6/acr-7/acr-9/acr-10/[illegible] | 141 |
| GO:0007267 | cell-cell signaling | 56/243 | 336/9281 | 3.33E-30 | 4.20E-28 | 2.55E-28 | acr-2/acr-3/acr-6/acr-7/acr-9/acr-10/[illegible] | 56 |
| GO:0007610 | behavior | 71/243 | 580/9281 | 9.12E-30 | 1.08E-27 | 6.55E-28 | bas-1/cat-2/daf-3/daf-7/dop-1/eat-16/[illegible] | 71 |
| GO:0007188 | adenylate cyclase-modulating G protein-coupled receptor signaling pathway | 25/243 | 43/9281 | 3.27E-29 | 3.63E-27 | 2.21E-27 | dop-1/egl-30/goa-1/gpa-1/gpa-2/gpa-3/[illegible] | 25 |
| GO:0051239 | regulation of multicellular organismal process | 68/243 | 558/9281 | 2.64E-28 | 2.77E-26 | 1.68E-26 | acr-2/daf-3/daf-5/daf-7/dnc-1/eat-16/[illegible] | 68 |
| GO:0050794 | regulation of cellular process | 175/243 | 3542/9281 | 2.18E-27 | 2.17E-25 | 1.32E-25 | acr-2/acr-3/acr-6/acr-7/acr-9/acr-10/[illegible] | 175 |
| GO:0032501 | multicellular organismal process | 137/243 | 2359/9281 | 2.61E-25 | 2.46E-23 | 1.50E-23 | acr-2/acr-3/acr-6/acr-7/acr-9/acr-10/[illegible] | 137 |
| GO:0040011 | locomotion | 62/243 | 523/9281 | 5.14E-25 | 4.62E-23 | 2.81E-23 | acr-2/bas-1/cat-2/ced-1/dop-1/eat-16/[illegible] | 62 |
| GO:0044057 | regulation of system process | 28/243 | 79/9281 | 6.21E-25 | 5.33E-23 | 3.24E-23 | acr-2/egl-19/egl-21/egl-30/flp-1/flp-16/[illegible] | 28 |
| GO:0007218 | neuropeptide signaling pathway | 33/243 | 129/9281 | 4.56E-24 | 3.75E-22 | 2.28E-22 | flp-1/flp-3/flp-6/flp-7/flp-9/flp-11/flp-13/[illegible] | 33 |
| GO:0034220 | ion transmembrane transport | 61/243 | 565/9281 | 2.02E-22 | 1.59E-20 | 9.68E-21 | acr-2/acr-3/acr-6/acr-7/acr-9/acr-10/[illegible] | 61 |
| GO:0003008 | system process | 65/243 | 647/9281 | 3.25E-22 | 2.46E-20 | 1.49E-20 | acr-2/acr-3/acr-6/acr-7/acr-9/acr-10/[illegible] | 65 |
| GO:0019098 | reproductive behavior | 36/243 | 181/9281 | 3.81E-22 | 2.77E-20 | 1.68E-20 | bas-1/cat-2/eat-16/egl-1/egl-2/egl-6/[illegible] | 36 |
| GO:0006811 | ion transport | 69/243 | 735/9281 | 5.97E-22 | 4.18E-20 | 2.54E-20 | acr-2/acr-3/acr-6/acr-7/acr-9/acr-10/[illegible] | 69 |
| GO:0040012 | regulation of locomotion | 42/243 | 262/9281 | 6.31E-22 | 4.26E-20 | 2.59E-20 | acr-2/cat-2/ced-1/dop-1/eat-16/egl-8/[illegible] | 42 |
| GO:0043051 | regulation of pharyngeal pumping | 21/243 | 46/9281 | 9.81E-22 | 6.39E-20 | 3.89E-20 | daf-3/daf-7/eat-16/egl-8/egl-19/egl-21/[illegible] | 21 |
| GO:1903998 | regulation of eating behavior | 22/243 | 53/9281 | 1.37E-21 | 8.60E-20 | 5.23E-20 | daf-3/daf-7/eat-16/egl-8/egl-19/egl-21/[illegible] | 22 |
| GO:0007631 | feeding behavior | 25/243 | 81/9281 | 1.18E-20 | 7.19E-19 | 4.37E-19 | daf-3/daf-7/eat-16/egl-8/egl-10/egl-19/[illegible] | 25 |
| GO:0060259 | regulation of feeding behavior | 22/243 | 58/9281 | 1.48E-20 | 8.73E-19 | 5.31E-19 | daf-3/daf-7/eat-16/egl-8/egl-19/egl-21/[illegible] | 22 |
| GO:0006812 | cation transport | 48/243 | 401/9281 | 1.90E-19 | 1.09E-17 | 6.61E-18 | acr-2/acr-3/dat-1/del-1/egl-2/egl-8/[illegible] | 48 |
| GO:0006937 | regulation of muscle contraction | 20/243 | 52/9281 | 6.28E-19 | 3.39E-17 | 2.06E-17 | acr-2/egl-19/egl-21/flp-1/flp-16/flp-18/[illegible] | 20 |
| GO:0090257 | regulation of muscle system process | 20/243 | 52/9281 | 6.28E-19 | 3.39E-17 | 2.06E-17 | acr-2/egl-19/egl-21/flp-1/flp-16/flp-18/[illegible] | 20 |
| GO:0042755 | eating behavior | 22/243 | 69/9281 | 1.20E-18 | 6.20E-17 | 3.77E-17 | daf-3/daf-7/eat-16/egl-8/egl-19/egl-21/[illegible] | 22 |
| GO:0043050 | pharyngeal pumping | 21/243 | 61/9281 | 1.22E-18 | 6.20E-17 | 3.77E-17 | daf-3/daf-7/eat-16/egl-8/egl-19/egl-21/[illegible] | 21 |
| GO:0010646 | regulation of cell communication | 49/243 | 442/9281 | 2.00E-18 | 9.93E-17 | 6.04E-17 | daf-5/daf-7/eat-16/egl-8/egl-10/egl-17/[illegible] | 49 |
| GO:0060079 | excitatory postsynaptic potential | 15/243 | 25/9281 | 3.14E-18 | 1.52E-16 | 9.24E-17 | acr-2/acr-3/acr-6/acr-9/acr-10/acr-11/[illegible] | 15 |
| GO:0099565 | chemical synaptic transmission, postsynaptic | 15/243 | 26/9281 | 7.25E-18 | 3.42E-16 | 2.08E-16 | acr-2/acr-3/acr-6/acr-9/acr-10/acr-11/[illegible] | 15 |
| GO:0003012 | muscle system process | 21/243 | 67/9281 | 1.13E-17 | 5.07E-16 | 3.08E-16 | acr-2/egl-2/egl-19/egl-21/flp-1/flp-16/[illegible] | 21 |
| GO:0006936 | muscle contraction | 21/243 | 67/9281 | 1.13E-17 | 5.07E-16 | 3.08E-16 | acr-2/egl-2/egl-19/egl-21/flp-1/flp-16/[illegible] | 21 |
| GO:0009605 | response to external stimulus | 52/243 | 519/9281 | 1.31E-17 | 5.73E-16 | 3.48E-16 | cat-2/ced-1/daf-3/daf-7/dop-1/dop-2/[illegible] | 52 |
| GO:0023051 | regulation of signaling | 48/243 | 444/9281 | 1.33E-17 | 5.73E-16 | 3.48E-16 | daf-5/daf-7/eat-16/egl-8/egl-10/egl-17/[illegible] | 48 |
| GO:0018991 | oviposition | 29/243 | 158/9281 | 5.46E-17 | 2.29E-15 | 1.39E-15 | bas-1/eat-16/egl-1/egl-2/egl-6/egl-8/[illegible] | 29 |
| GO:0065008 | regulation of biological quality | 65/243 | 826/9281 | 1.34E-16 | 5.50E-15 | 3.34E-15 | acr-2/acr-3/acr-6/acr-7/acr-9/acr-10/[illegible] | 65 |
| GO:0007271 | synaptic transmission, cholinergic | 17/243 | 43/9281 | 1.68E-16 | 6.75E-15 | 4.10E-15 | egl-8/egl-21/egl-30/gar-2/goa-1/nca-2/[illegible] | 17 |
| GO:0048583 | regulation of response to stimulus | 51/243 | 536/9281 | 2.54E-16 | 9.99E-15 | 6.07E-15 | daf-5/daf-7/dat-1/eat-16/egl-8/egl-10/[illegible] | 51 |
| GO:0001505 | regulation of neurotransmitter levels | 22/243 | 87/9281 | 3.06E-16 | 1.15E-14 | 7.02E-15 | bas-1/cat-2/dat-1/egl-8/egl-21/egl-30/[illegible] | 22 |
| GO:0006836 | neurotransmitter transport | 22/243 | 87/9281 | 3.06E-16 | 1.15E-14 | 7.02E-15 | dat-1/egl-8/egl-21/egl-30/mod-5/pkc-1/[illegible] | 22 |
| GO:0098655 | cation transmembrane transport | 40/243 | 335/9281 | 3.19E-16 | 1.18E-14 | 7.18E-15 | acr-2/acr-3/del-1/egl-2/egl-19/egl-23/[illegible] | 40 |
| GO:0060078 | regulation of postsynaptic membrane potential | 15/243 | 33/9281 | 8.26E-16 | 3.00E-14 | 1.82E-14 | acr-2/acr-3/acr-6/acr-9/acr-10/acr-11/[illegible] | 15 |
| GO:0048521 | negative regulation of behavior | 12/243 | 18/9281 | 1.28E-15 | 4.56E-14 | 2.77E-14 | egl-6/egl-21/flp-10/flp-12/flp-16/flp-17/[illegible] | 12 |
| GO:0055085 | transmembrane transport | 68/243 | 949/9281 | 2.71E-15 | 9.49E-14 | 5.77E-14 | acr-2/acr-3/acr-6/acr-7/acr-9/acr-10/[illegible] | 68 |
| GO:0007212 | dopamine receptor signaling pathway | 11/243 | 15/9281 | 3.94E-15 | 1.35E-13 | 8.23E-14 | dop-1/dop-2/eat-16/egl-8/egl-10/egl-30/[illegible] | 11 |
| GO:0007210 | serotonin receptor signaling pathway | 12/243 | 20/9281 | 8.29E-15 | 2.80E-13 | 1.70E-13 | dop-1/dop-2/egl-30/gar-2/gpa-11/mod-1/[illegible] | 12 |
| GO:0070588 | calcium ion transmembrane transport | 16/243 | 48/9281 | 3.19E-14 | 1.06E-12 | 6.42E-13 | acr-2/acr-3/egl-19/lov-1/ocr-1/ocr-2/[illegible] | 16 |
| GO:0046662 | regulation of oviposition | 19/243 | 76/9281 | 4.40E-14 | 1.43E-12 | 8.70E-13 | eat-16/egl-6/egl-8/egl-10/egl-17/egl-30/[illegible] | 19 |
| GO:0007626 | locomotory behavior | 18/243 | 67/9281 | 5.33E-14 | 1.71E-12 | 1.04E-12 | bas-1/cat-2/egl-21/flp-1/flp-9/flp-10/[illegible] | 18 |
| GO:0006816 | calcium ion transport | 17/243 | 58/9281 | 5.55E-14 | 1.75E-12 | 1.06E-12 | acr-2/acr-3/egl-19/lov-1/ocr-1/ocr-2/[illegible] | 17 |
| GO:0098662 | inorganic cation transmembrane transport | 34/243 | 289/9281 | 1.06E-13 | 3.27E-12 | 1.99E-12 | acr-2/acr-3/del-1/egl-2/egl-19/egl-23/[illegible] | 34 |
| GO:0030001 | metal ion transport | 32/243 | 262/9281 | 2.11E-13 | 6.43E-12 | 3.91E-12 | acr-2/acr-3/del-1/egl-2/egl-19/egl-23/[illegible] | 32 |
| GO:0050804 | modulation of chemical synaptic transmission | 18/243 | 73/9281 | 2.72E-13 | 8.03E-12 | 4.88E-12 | eat-16/egl-8/egl-21/egl-30/gar-2/goa-1/[illegible] | 18 |
| GO:0099177 | regulation of trans-synaptic signaling | 18/243 | 73/9281 | 2.72E-13 | 8.03E-12 | 4.88E-12 | eat-16/egl-8/egl-21/egl-30/gar-2/goa-1/[illegible] | 18 |
| GO:0042391 | regulation of membrane potential | 26/243 | 172/9281 | 3.02E-13 | 8.76E-12 | 5.33E-12 | acr-2/acr-3/acr-6/acr-7/acr-9/acr-10/[illegible] | 26 |
| GO:0040017 | positive regulation of locomotion | 22/243 | 119/9281 | 3.30E-13 | 9.44E-12 | 5.74E-12 | ced-1/egl-8/egl-10/egl-19/egl-20/egl-21/[illegible] | 22 |
| GO:0030182 | neuron differentiation | 32/243 | 268/9281 | 3.98E-13 | 1.12E-11 | 6.82E-12 | daf-5/dnc-1/egl-8/egl-13/egl-20/egl-30/[illegible] | 32 |
| GO:0007399 | nervous system development | 37/243 | 362/9281 | 6.42E-13 | 1.78E-11 | 1.08E-11 | daf-5/dnc-1/egl-8/egl-13/egl-19/egl-20/[illegible] | 37 |
| GO:0048699 | generation of neurons | 33/243 | 292/9281 | 7.92E-13 | 2.17E-11 | 1.32E-11 | daf-5/dnc-1/egl-8/egl-13/egl-20/egl-30/[illegible] | 33 |
| GO:0098664 | G protein-coupled serotonin receptor signaling pathway | 10/243 | 16/9281 | 8.79E-13 | 2.37E-11 | 1.44E-11 | dop-1/dop-2/gar-2/mod-1/ser-1/ser-4/[illegible] | 10 |
| GO:0022008 | neurogenesis | 33/243 | 296/9281 | 1.16E-12 | 3.10E-11 | 1.88E-11 | daf-5/dnc-1/egl-8/egl-13/egl-20/egl-30/[illegible] | 33 |
| GO:0072347 | response to anesthetic | 8/243 | 9/9281 | 1.73E-12 | 4.51E-11 | 2.74E-11 | egl-21/flp-16/nlp-1/nlp-14/unc-7/unc-31/[illegible] | 8 |
| GO:0098660 | inorganic ion transmembrane transport | 37/243 | 374/9281 | 1.74E-12 | 4.51E-11 | 2.74E-11 | acr-2/acr-3/clh-3/dat-1/del-1/egl-2/[illegible] | 37 |
| GO:2000241 | regulation of reproductive process | 25/243 | 172/9281 | 2.19E-12 | 5.60E-11 | 3.40E-11 | eat-16/egl-6/egl-8/egl-10/egl-17/egl-30/[illegible] | 25 |
| GO:0032103 | positive regulation of response to external stimulus | 16/243 | 64/9281 | 4.75E-12 | 1.20E-10 | 7.28E-11 | egl-8/egl-20/egl-30/gpa-12/mec-4/mec-7/[illegible] | 16 |
| GO:0051179 | localization | 107/243 | 2252/9281 | 6.37E-12 | 1.58E-10 | 9.63E-11 | acr-2/acr-3/acr-6/acr-7/acr-9/acr-10/[illegible] | 107 |
| GO:0048609 | multicellular organismal reproductive process | 39/243 | 435/9281 | 9.03E-12 | 2.22E-10 | 1.35E-10 | bas-1/cat-2/eat-16/egl-1/egl-2/egl-6/[illegible] | 39 |
| GO:0048518 | positive regulation of biological process | 70/243 | 1176/9281 | 9.67E-12 | 2.34E-10 | 1.42E-10 | ced-1/daf-3/daf-5/daf-7/dat-1/egl-1/[illegible] | 70 |
| GO:0032222 | regulation of synaptic transmission, cholinergic | 11/243 | 25/9281 | 1.02E-11 | 2.44E-10 | 1.48E-10 | egl-8/egl-21/egl-30/gar-2/goa-1/nca-2/[illegible] | 11 |
| GO:1905114 | cell surface receptor signaling pathway involved in cell-cell signaling | 18/243 | 91/9281 | 1.54E-11 | 3.63E-10 | 2.21E-10 | acr-2/acr-3/acr-6/acr-9/acr-10/acr-11/[illegible] | 18 |
| GO:0006810 | transport | 91/243 | 1795/9281 | 2.04E-11 | 4.76E-10 | 2.89E-10 | acr-2/acr-3/acr-6/acr-7/acr-9/acr-10/[illegible] | 91 |
| GO:0070838 | divalent metal ion transport | 17/243 | 81/9281 | 2.11E-11 | 4.80E-10 | 2.92E-10 | acr-2/acr-3/egl-19/lov-1/ocr-1/ocr-2/[illegible] | 17 |
| GO:0072511 | divalent inorganic cation transport | 17/243 | 81/9281 | 2.11E-11 | 4.80E-10 | 2.92E-10 | acr-2/acr-3/egl-19/lov-1/ocr-1/ocr-2/[illegible] | 17 |
| GO:0032504 | multicellular organism reproduction | 39/243 | 454/9281 | 3.36E-11 | 7.56E-10 | 4.60E-10 | bas-1/cat-2/eat-16/egl-1/egl-2/egl-6/[illegible] | 39 |
| GO:0051234 | establishment of localization | 93/243 | 1872/9281 | 3.43E-11 | 7.62E-10 | 4.63E-10 | acr-2/acr-3/acr-6/acr-7/acr-9/acr-10/[illegible] | 93 |
| GO:0048666 | neuron development | 27/243 | 233/9281 | 6.32E-11 | 1.39E-09 | 8.44E-10 | daf-5/dnc-1/egl-8/egl-20/egl-30/egl-46/[illegible] | 27 |
| GO:0009612 | response to mechanical stimulus | 12/243 | 37/9281 | 8.21E-11 | 1.78E-09 | 1.08E-09 | cat-2/dop-1/flp-1/lov-1/mec-4/mec-7/[illegible] | 12 |
| GO:0032101 | regulation of response to external stimulus | 19/243 | 115/9281 | 1.12E-10 | 2.41E-09 | 1.47E-09 | egl-8/egl-20/egl-30/goa-1/gpa-12/jnk-1/[illegible] | 19 |
| GO:0045664 | regulation of neuron differentiation | 19/243 | 116/9281 | 1.31E-10 | 2.78E-09 | 1.69E-09 | dnc-1/egl-8/egl-20/egl-30/egl-43/goa-1/[illegible] | 19 |
| GO:0051960 | regulation of nervous system development | 21/243 | 146/9281 | 1.69E-10 | 3.55E-09 | 2.16E-09 | dnc-1/egl-8/egl-20/egl-21/egl-30/egl-43/[illegible] | 21 |
| GO:0009628 | response to abiotic stimulus | 27/243 | 244/9281 | 1.83E-10 | 3.80E-09 | 2.31E-09 | cat-2/daf-7/dop-1/egl-8/egl-9/flp-1/[illegible] | 27 |
| GO:0045595 | regulation of cell differentiation | 27/243 | 245/9281 | 2.01E-10 | 4.13E-09 | 2.51E-09 | daf-5/dnc-1/egl-8/egl-20/egl-30/egl-43/[illegible] | 27 |
| GO:0042493 | response to drug | 23/243 | 179/9281 | 2.26E-10 | 4.58E-09 | 2.79E-09 | dat-1/egl-8/egl-21/egl-30/flp-16/gar-2/[illegible] | 23 |
| GO:0031175 | neuron projection development | 25/243 | 214/9281 | 2.84E-10 | 5.71E-09 | 3.47E-09 | dnc-1/egl-8/egl-20/egl-30/egl-46/goa-1/[illegible] | 25 |
| GO:0060284 | regulation of cell development | 22/243 | 167/9281 | 3.48E-10 | 6.93E-09 | 4.21E-09 | dnc-1/egl-8/egl-20/egl-30/egl-43/gld-1/[illegible] | 22 |
| GO:0007617 | mating behavior | 11/243 | 33/9281 | 3.68E-10 | 7.21E-09 | 4.39E-09 | bas-1/cat-2/flp-10/flp-12/flp-20/lov-1/[illegible] | 11 |
| GO:0007166 | cell surface receptor signaling pathway | 32/243 | 346/9281 | 3.72E-10 | 7.21E-09 | 4.39E-09 | acr-2/acr-3/acr-6/acr-9/acr-10/acr-11/[illegible] | 32 |
| GO:0048731 | system development | 48/243 | 699/9281 | 3.77E-10 | 7.21E-09 | 4.39E-09 | daf-5/dnc-1/egl-8/egl-13/egl-15/egl-19/[illegible] | 48 |
| GO:0051240 | positive regulation of multicellular organismal process | 28/243 | 270/9281 | 3.78E-10 | 7.21E-09 | 4.39E-09 | daf-3/daf-5/egl-8/egl-19/egl-20/egl-21/[illegible] | 28 |
| GO:0022414 | reproductive process | 51/243 | 780/9281 | 5.44E-10 | 1.03E-08 | 6.25E-09 | bas-1/cat-2/dnc-1/eat-16/egl-1/egl-2/[illegible] | 51 |
| GO:0031344 | regulation of cell projection organization | 19/243 | 126/9281 | 5.70E-10 | 1.05E-08 | 6.41E-09 | dnc-1/egl-8/egl-15/egl-20/egl-30/goa-1/[illegible] | 19 |
| GO:0050767 | regulation of neurogenesis | 19/243 | 126/9281 | 5.70E-10 | 1.05E-08 | 6.41E-09 | dnc-1/egl-8/egl-20/egl-30/egl-43/goa-1/[illegible] | 19 |
| GO:0050877 | nervous system process | 41/243 | 547/9281 | 6.84E-10 | 1.25E-08 | 7.62E-09 | acr-2/acr-3/acr-6/acr-7/acr-9/acr-10/[illegible] | 41 |
| GO:0050793 | regulation of developmental process | 39/243 | 503/9281 | 7.15E-10 | 1.30E-08 | 7.89E-09 | ced-1/daf-3/daf-5/daf-7/dnc-1/egl-1/[illegible] | 39 |
| GO:0007618 | mating | 11/243 | 35/9281 | 7.58E-10 | 1.36E-08 | 8.29E-09 | bas-1/cat-2/flp-10/flp-12/flp-20/lov-1/[illegible] | 11 |
| GO:0032094 | response to food | 7/243 | 10/9281 | 8.69E-10 | 1.55E-08 | 9.42E-09 | cat-2/dop-1/dop-2/eat-16/goa-1/gpb-2/[illegible] | 7 |
| GO:2000026 | regulation of multicellular organismal development | 32/243 | 358/9281 | 8.84E-10 | 1.56E-08 | 9.49E-09 | daf-3/daf-5/daf-7/dnc-1/egl-8/egl-20/[illegible] | 32 |
| GO:0007638 | mechanosensory behavior | 10/243 | 28/9281 | 1.09E-09 | 1.88E-08 | 1.14E-08 | cat-2/dop-1/flp-1/mec-4/mec-7/mec-10/[illegible] | 10 |
| GO:0046928 | regulation of neurotransmitter secretion | 10/243 | 28/9281 | 1.09E-09 | 1.88E-08 | 1.14E-08 | egl-8/egl-21/egl-30/ric-19/tom-1/unc-10/[illegible] | 10 |
| GO:0051588 | regulation of neurotransmitter transport | 10/243 | 28/9281 | 1.09E-09 | 1.88E-08 | 1.14E-08 | egl-8/egl-21/egl-30/ric-19/tom-1/unc-10/[illegible] | 10 |
| GO:0061564 | axon development | 22/243 | 178/9281 | 1.20E-09 | 2.05E-08 | 1.25E-08 | egl-8/egl-20/egl-30/egl-46/goa-1/jnk-1/[illegible] | 22 |
| GO:0010975 | regulation of neuron projection development | 17/243 | 104/9281 | 1.32E-09 | 2.22E-08 | 1.35E-08 | dnc-1/egl-8/egl-20/egl-30/goa-1/jnk-1/[illegible] | 17 |
| GO:0010469 | regulation of signaling receptor activity | 13/243 | 56/9281 | 1.37E-09 | 2.29E-08 | 1.39E-08 | daf-7/egl-17/egl-20/ins-3/ins-4/ins-15/[illegible] | 13 |
| GO:0070848 | response to growth factor | 12/243 | 46/9281 | 1.40E-09 | 2.30E-08 | 1.40E-08 | daf-3/daf-5/daf-7/egl-15/egl-17/lon-2/[illegible] | 12 |
| GO:0071363 | cellular response to growth factor stimulus | 12/243 | 46/9281 | 1.40E-09 | 2.30E-08 | 1.40E-08 | daf-3/daf-5/daf-7/egl-15/egl-17/lon-2/[illegible] | 12 |
| GO:0051046 | regulation of secretion | 14/243 | 68/9281 | 1.72E-09 | 2.79E-08 | 1.70E-08 | egl-8/egl-21/egl-30/gsa-1/rab-3/ric-19/[illegible] | 14 |
| GO:0120035 | regulation of plasma membrane bounded cell projection organization | 18/243 | 121/9281 | 2.06E-09 | 3.33E-08 | 2.02E-08 | dnc-1/egl-8/egl-20/egl-30/goa-1/jnk-1/[illegible] | 18 |
| GO:0000003 | reproduction | 52/243 | 848/9281 | 3.44E-09 | 5.51E-08 | 3.35E-08 | bas-1/cat-2/dnc-1/eat-16/egl-1/egl-2/[illegible] | 52 |
| GO:0050806 | positive regulation of synaptic transmission | 10/243 | 32/9281 | 4.90E-09 | 7.77E-08 | 4.72E-08 | egl-8/egl-21/egl-30/nca-2/unc-10/unc-18/[illegible] | 10 |
| GO:0042221 | response to chemical | 61/243 | 1101/9281 | 5.70E-09 | 8.98E-08 | 5.46E-08 | cat-2/ced-1/daf-3/daf-5/daf-7/dat-1/[illegible] | 61 |
| GO:0032224 | positive regulation of synaptic transmission, cholinergic | 8/243 | 18/9281 | 6.87E-09 | 1.06E-07 | 6.47E-08 | egl-8/egl-21/egl-30/nca-2/unc-10/unc-29/[illegible] | 8 |
| GO:0035690 | cellular response to drug | 8/243 | 18/9281 | 6.87E-09 | 1.06E-07 | 6.47E-08 | egl-21/flp-16/gar-2/goa-1/gpb-2/nlp-1/[illegible] | 8 |
| GO:0071495 | cellular response to endogenous stimulus | 19/243 | 146/9281 | 7.20E-09 | 1.11E-07 | 6.72E-08 | daf-3/daf-5/daf-7/dat-1/egl-15/egl-17/[illegible] | 19 |
| GO:0009719 | response to endogenous stimulus | 19/243 | 147/9281 | 8.08E-09 | 1.23E-07 | 7.48E-08 | daf-3/daf-5/daf-7/dat-1/egl-15/egl-17/[illegible] | 19 |
| GO:1903530 | regulation of secretion by cell | 12/243 | 53/9281 | 8.16E-09 | 1.23E-07 | 7.49E-08 | egl-8/egl-21/egl-30/rab-3/ric-19/sng-1/[illegible] | 12 |
| GO:0048584 | positive regulation of response to stimulus | 24/243 | 234/9281 | 9.22E-09 | 1.38E-07 | 8.40E-08 | daf-7/egl-8/egl-20/egl-21/egl-30/gpa-12/[illegible] | 24 |
| GO:0009966 | regulation of signal transduction | 31/243 | 373/9281 | 9.61E-09 | 1.43E-07 | 8.69E-08 | daf-5/daf-7/eat-16/egl-10/egl-17/egl-20/[illegible] | 31 |
| GO:0032879 | regulation of localization | 34/243 | 438/9281 | 9.85E-09 | 1.45E-07 | 8.84E-08 | ced-1/egl-2/egl-8/egl-9/egl-17/egl-19/[illegible] | 34 |
| GO:0007269 | neurotransmitter secretion | 12/243 | 54/9281 | 1.02E-08 | 1.48E-07 | 8.98E-08 | egl-8/egl-21/egl-30/pkc-1/ric-19/tom-1/[illegible] | 12 |
| GO:0097479 | synaptic vesicle localization | 12/243 | 54/9281 | 1.02E-08 | 1.48E-07 | 8.98E-08 | pkc-1/sng-1/tom-1/unc-4/unc-10/unc-14/[illegible] | 12 |
| GO:0099643 | signal release from synapse | 12/243 | 54/9281 | 1.02E-08 | 1.48E-07 | 8.98E-08 | egl-8/egl-21/egl-30/pkc-1/ric-19/tom-1/[illegible] | 12 |
| GO:0051241 | negative regulation of multicellular organismal process | 17/243 | 120/9281 | 1.25E-08 | 1.79E-07 | 1.09E-07 | daf-7/egl-6/egl-20/flp-10/flp-12/flp-16/[illegible] | 17 |
| GO:0007179 | transforming growth factor beta receptor signaling pathway | 10/243 | 35/9281 | 1.30E-08 | 1.79E-07 | 1.09E-07 | daf-3/daf-5/daf-7/lon-2/sma-2/sma-3/[illegible] | 10 |
| GO:0009581 | detection of external stimulus | 10/243 | 35/9281 | 1.30E-08 | 1.79E-07 | 1.09E-07 | gpa-3/lite-1/lov-1/mec-4/mec-7/mec-10/[illegible] | 10 |
| GO:0009582 | detection of abiotic stimulus | 10/243 | 35/9281 | 1.30E-08 | 1.79E-07 | 1.09E-07 | gpa-3/lite-1/lov-1/mec-4/mec-7/mec-10/[illegible] | 10 |
| GO:0071559 | response to transforming growth factor beta | 10/243 | 35/9281 | 1.30E-08 | 1.79E-07 | 1.09E-07 | daf-3/daf-5/daf-7/lon-2/sma-2/sma-3/[illegible] | 10 |
| GO:0071560 | cellular response to transforming growth factor beta stimulus | 10/243 | 35/9281 | 1.30E-08 | 1.79E-07 | 1.09E-07 | daf-3/daf-5/daf-7/lon-2/sma-2/sma-3/[illegible] | 10 |
| GO:0030030 | cell projection organization | 28/243 | 316/9281 | 1.33E-08 | 1.82E-07 | 1.11E-07 | dnc-1/egl-8/egl-15/egl-20/egl-30/egl-46/[illegible] | 28 |
| GO:0120036 | plasma membrane bounded cell projection organization | 27/243 | 303/9281 | 2.19E-08 | 2.97E-07 | 1.81E-07 | dnc-1/egl-8/egl-20/egl-30/egl-46/goa-1/[illegible] | 27 |
| GO:0050975 | sensory perception of touch | 7/243 | 14/9281 | 2.27E-08 | 3.07E-07 | 1.86E-07 | mec-4/mec-7/mec-10/mec-12/mec-14/[illegible] | 7 |
| GO:0023061 | signal release | 12/243 | 58/9281 | 2.43E-08 | 3.25E-07 | 1.98E-07 | egl-8/egl-21/egl-30/pkc-1/ric-19/tom-1/[illegible] | 12 |
| GO:1901700 | response to oxygen-containing compound | 16/243 | 111/9281 | 2.65E-08 | 3.52E-07 | 2.14E-07 | daf-7/dat-1/egl-21/flp-16/gar-2/goa-1/[illegible] | 16 |
| GO:1901701 | cellular response to oxygen-containing compound | 13/243 | 71/9281 | 2.97E-08 | 3.93E-07 | 2.39E-07 | daf-7/dat-1/egl-21/flp-16/gar-2/goa-1/[illegible] | 13 |
| GO:0050982 | detection of mechanical stimulus | 7/243 | 15/9281 | 4.16E-08 | 5.46E-07 | 3.32E-07 | lov-1/mec-4/mec-7/mec-10/mec-12/[illegible] | 7 |
| GO:0048468 | cell development | 35/243 | 488/9281 | 4.40E-08 | 5.73E-07 | 3.48E-07 | daf-5/daf-7/dnc-1/egl-8/egl-20/egl-30/[illegible] | 35 |
| GO:0042330 | taxis | 22/243 | 216/9281 | 4.49E-08 | 5.81E-07 | 3.53E-07 | egl-8/egl-20/egl-30/egl-46/flp-10/goa-1/[illegible] | 22 |
| GO:0051480 | regulation of cytosolic calcium ion concentration | 8/243 | 22/9281 | 4.58E-08 | 5.89E-07 | 3.58E-07 | acr-2/acr-3/spe-41/trp-1/trp-2/unc-38/[illegible] | 8 |
| GO:0048489 | synaptic vesicle transport | 11/243 | 50/9281 | 4.79E-08 | 6.08E-07 | 3.70E-07 | pkc-1/sng-1/tom-1/unc-4/unc-10/unc-16/[illegible] | 11 |
| GO:0097480 | establishment of synaptic vesicle localization | 11/243 | 50/9281 | 4.79E-08 | 6.08E-07 | 3.70E-07 | pkc-1/sng-1/tom-1/unc-4/unc-10/unc-16/[illegible] | 11 |
| GO:0015696 | ammonium transport | 10/243 | 41/9281 | 6.91E-08 | 8.51E-07 | 5.17E-07 | dat-1/egl-8/egl-21/egl-30/mod-5/snf-6/[illegible] | 10 |
| GO:0010037 | response to carbon dioxide | 5/243 | 6/9281 | 6.94E-08 | 8.51E-07 | 5.17E-07 | egl-21/flp-16/nlp-1/nlp-14/unc-31 | 5 |
| GO:0071244 | cellular response to carbon dioxide | 5/243 | 6/9281 | 6.94E-08 | 8.51E-07 | 5.17E-07 | egl-21/flp-16/nlp-1/nlp-14/unc-31 | 5 |
| GO:1903745 | negative regulation of pharyngeal pumping | 5/243 | 6/9281 | 6.94E-08 | 8.51E-07 | 5.17E-07 | egl-21/flp-16/nlp-1/nlp-14/unc-31 | 5 |
| GO:1903999 | negative regulation of eating behavior | 5/243 | 6/9281 | 6.94E-08 | 8.51E-07 | 5.17E-07 | egl-21/flp-16/nlp-1/nlp-14/unc-31 | 5 |
| GO:0051648 | vesicle localization | 13/243 | 77/9281 | 8.18E-08 | 9.97E-07 | 6.06E-07 | jnk-1/pkc-1/sng-1/tom-1/unc-4/unc-10/[illegible] | 13 |
| GO:0007178 | transmembrane receptor protein serine/threonine kinase signaling pathway | 10/243 | 43/9281 | 1.13E-07 | 1.37E-06 | 8.31E-07 | daf-3/daf-5/daf-7/lon-2/sma-2/sma-3/[illegible] | 10 |
| GO:0046903 | secretion | 17/243 | 139/9281 | 1.16E-07 | 1.40E-06 | 8.49E-07 | egl-8/egl-21/egl-30/gsa-1/pkc-1/rab-3/[illegible] | 17 |
| GO:0050954 | sensory perception of mechanical stimulus | 7/243 | 17/9281 | 1.20E-07 | 1.44E-06 | 8.75E-07 | mec-4/mec-7/mec-10/mec-12/mec-14/[illegible] | 7 |
| GO:0046189 | phenol-containing compound biosynthetic process | 6/243 | 11/9281 | 1.25E-07 | 1.48E-06 | 9.00E-07 | bas-1/cat-2/cat-4/tbh-1/tdc-1/tph-1 | 6 |
| GO:0070509 | calcium ion import | 6/243 | 11/9281 | 1.25E-07 | 1.48E-06 | 9.00E-07 | acr-2/acr-3/egl-19/unc-2/unc-38/unc-63 | 6 |
| GO:0048812 | neuron projection morphogenesis | 19/243 | 176/9281 | 1.55E-07 | 1.80E-06 | 1.10E-06 | dnc-1/egl-20/egl-46/rab-3/syg-1/unc-4/[illegible] | 19 |
| GO:0120039 | plasma membrane bounded cell projection morphogenesis | 19/243 | 176/9281 | 1.55E-07 | 1.80E-06 | 1.10E-06 | dnc-1/egl-20/egl-46/rab-3/syg-1/unc-4/[illegible] | 19 |
| GO:0099003 | vesicle-mediated transport in synapse | 11/243 | 56/9281 | 1.66E-07 | 1.93E-06 | 1.17E-06 | pkc-1/sng-1/tom-1/unc-4/unc-10/unc-16/[illegible] | 11 |
| GO:0048858 | cell projection morphogenesis | 19/243 | 178/9281 | 1.85E-07 | 2.12E-06 | 1.29E-06 | dnc-1/egl-20/egl-46/rab-3/syg-1/unc-4/[illegible] | 19 |
| GO:0048522 | positive regulation of cellular process | 51/243 | 928/9281 | 1.85E-07 | 2.12E-06 | 1.29E-06 | ced-1/daf-7/dat-1/egl-1/egl-5/egl-8/[illegible] | 51 |
| GO:0060179 | male mating behavior | 8/243 | 26/9281 | 2.05E-07 | 2.33E-06 | 1.42E-06 | bas-1/cat-2/flp-10/flp-12/flp-20/lov-1/[illegible] | 8 |
| GO:0051049 | regulation of transport | 22/243 | 236/9281 | 2.18E-07 | 2.46E-06 | 1.50E-06 | ced-1/egl-2/egl-8/egl-19/egl-21/egl-30/[illegible] | 22 |
| GO:0034607 | turning behavior involved in mating | 6/243 | 12/9281 | 2.45E-07 | 2.71E-06 | 1.65E-06 | bas-1/cat-2/flp-10/flp-12/flp-20/mec-4 | 6 |
| GO:0035178 | turning behavior | 6/243 | 12/9281 | 2.45E-07 | 2.71E-06 | 1.65E-06 | bas-1/cat-2/flp-10/flp-12/flp-20/mec-4 | 6 |
| GO:0050974 | detection of mechanical stimulus involved in sensory perception | 6/243 | 12/9281 | 2.45E-07 | 2.71E-06 | 1.65E-06 | mec-4/mec-7/mec-10/mec-12/mec-14/[illegible] | 6 |
| GO:0050976 | detection of mechanical stimulus involved in sensory perception of touch | 6/243 | 12/9281 | 2.45E-07 | 2.71E-06 | 1.65E-06 | mec-4/mec-7/mec-10/mec-12/mec-14/[illegible] | 6 |
| GO:0032990 | cell part morphogenesis | 19/243 | 182/9281 | 2.63E-07 | 2.89E-06 | 1.76E-06 | dnc-1/egl-20/egl-46/rab-3/syg-1/unc-4/[illegible] | 19 |
| GO:0030154 | cell differentiation | 47/243 | 833/9281 | 2.91E-07 | 3.18E-06 | 1.93E-06 | daf-5/daf-7/dnc-1/egl-8/egl-13/egl-18/[illegible] | 47 |
| GO:0015870 | acetylcholine transport | 7/243 | 19/9281 | 2.98E-07 | 3.22E-06 | 1.96E-06 | egl-8/egl-21/egl-30/snf-6/unc-17/unc-26/[illegible] | 7 |
| GO:1901374 | acetate ester transport | 7/243 | 19/9281 | 2.98E-07 | 3.22E-06 | 1.96E-06 | egl-8/egl-21/egl-30/snf-6/unc-17/unc-26/[illegible] | 7 |
| GO:0010033 | response to organic substance | 31/243 | 434/9281 | 3.09E-07 | 3.32E-06 | 2.02E-06 | ced-1/daf-3/daf-5/daf-7/dat-1/egl-15/[illegible] | 31 |
| GO:0051094 | positive regulation of developmental process | 22/243 | 241/9281 | 3.14E-07 | 3.35E-06 | 2.04E-06 | ced-1/daf-3/daf-5/egl-1/egl-8/egl-20/[illegible] | 22 |
| GO:0032940 | secretion by cell | 15/243 | 116/9281 | 3.16E-07 | 3.35E-06 | 2.04E-06 | egl-8/egl-21/egl-30/pkc-1/rab-3/ric-19/[illegible] | 15 |
| GO:0007275 | multicellular organism development | 73/243 | 1587/9281 | 3.28E-07 | 3.47E-06 | 2.11E-06 | ced-1/daf-3/daf-5/daf-7/dnc-1/eat-16/[illegible] | 73 |
| GO:0051650 | establishment of vesicle localization | 12/243 | 73/9281 | 3.53E-07 | 3.71E-06 | 2.25E-06 | jnk-1/pkc-1/sng-1/tom-1/unc-4/unc-10/[illegible] | 12 |
| GO:0090092 | regulation of transmembrane receptor protein serine/threonine kinase signaling pathway | 8/243 | 28/9281 | 3.89E-07 | 4.05E-06 | 2.46E-06 | daf-5/daf-7/lon-2/sma-3/tph-1/unc-2/[illegible] | 8 |
| GO:0022603 | regulation of anatomical structure morphogenesis | 17/243 | 151/9281 | 3.90E-07 | 4.05E-06 | 2.46E-06 | dnc-1/egl-20/lin-39/lon-2/rab-3/sma-2/[illegible] | 17 |
| GO:0048519 | negative regulation of biological process | 52/243 | 981/9281 | 4.29E-07 | 4.43E-06 | 2.69E-06 | daf-3/daf-5/daf-7/dat-1/eat-16/egl-5/[illegible] | 52 |
| GO:0048667 | cell morphogenesis involved in neuron differentiation | 18/243 | 170/9281 | 4.44E-07 | 4.55E-06 | 2.77E-06 | dnc-1/egl-20/egl-46/rab-3/syg-1/unc-4/[illegible] | 18 |
| GO:0018958 | phenol-containing compound metabolic process | 6/243 | 13/9281 | 4.46E-07 | 4.55E-06 | 2.77E-06 | bas-1/cat-2/cat-4/tbh-1/tdc-1/tph-1 | 6 |
| GO:0048869 | cellular developmental process | 47/243 | 853/9281 | 5.82E-07 | 5.91E-06 | 3.60E-06 | daf-5/daf-7/dnc-1/egl-8/egl-13/egl-18/[illegible] | 47 |
| GO:2000252 | negative regulation of feeding behavior | 5/243 | 8/9281 | 6.20E-07 | 6.27E-06 | 3.81E-06 | egl-21/flp-16/nlp-1/nlp-14/unc-31 | 5 |
| GO:0009914 | hormone transport | 7/243 | 21/9281 | 6.58E-07 | 6.61E-06 | 4.02E-06 | egl-8/egl-21/egl-30/snf-6/unc-17/unc-26/[illegible] | 7 |
| GO:0007409 | axonogenesis | 17/243 | 159/9281 | 8.17E-07 | 8.12E-06 | 4.94E-06 | egl-20/egl-46/rab-3/syg-1/unc-4/unc-30/[illegible] | 17 |
| GO:0010647 | positive regulation of cell communication | 17/243 | 159/9281 | 8.17E-07 | 8.12E-06 | 4.94E-06 | daf-7/egl-8/egl-21/egl-30/nca-2/sma-3/[illegible] | 17 |
| GO:0032989 | cellular component morphogenesis | 22/243 | 258/9281 | 1.01E-06 | 1.00E-05 | 6.08E-06 | dnc-1/egl-20/egl-46/rab-3/syg-1/unc-4/[illegible] | 22 |
| GO:0000904 | cell morphogenesis involved in differentiation | 18/243 | 182/9281 | 1.22E-06 | 1.20E-05 | 7.30E-06 | dnc-1/egl-20/egl-46/rab-3/syg-1/unc-4/[illegible] | 18 |
| GO:0001956 | positive regulation of neurotransmitter secretion | 6/243 | 15/9281 | 1.24E-06 | 1.21E-05 | 7.36E-06 | egl-8/egl-21/egl-30/unc-18/unc-26/unc-29 | 6 |
| GO:0051590 | positive regulation of neurotransmitter transport | 6/243 | 15/9281 | 1.24E-06 | 1.21E-05 | 7.36E-06 | egl-8/egl-21/egl-30/unc-18/unc-26/unc-29 | 6 |
| GO:0015695 | organic cation transport | 7/243 | 24/9281 | 1.83E-06 | 1.77E-05 | 1.08E-05 | egl-8/egl-21/egl-30/snf-6/unc-17/unc-26/[illegible] | 7 |
| GO:0051704 | multi-organism process | 33/243 | 523/9281 | 2.07E-06 | 2.00E-05 | 1.21E-05 | bas-1/cat-2/ced-1/daf-3/daf-7/dnc-1/[illegible] | 33 |
| GO:0010769 | regulation of cell morphogenesis involved in differentiation | 12/243 | 86/9281 | 2.17E-06 | 2.08E-05 | 1.27E-05 | dnc-1/egl-20/rab-3/unc-4/unc-31/unc-37/[illegible] | 12 |
| GO:0071875 | adrenergic receptor signaling pathway | 4/243 | 5/9281 | 2.25E-06 | 2.11E-05 | 1.28E-05 | ser-2/ser-5/tyra-2/dop-3 | 4 |
| GO:0071880 | adenylate cyclase-activating adrenergic receptor signaling pathway | 4/243 | 5/9281 | 2.25E-06 | 2.11E-05 | 1.28E-05 | ser-2/ser-5/tyra-2/dop-3 | 4 |
| GO:0098703 | calcium ion import across plasma membrane | 4/243 | 5/9281 | 2.25E-06 | 2.11E-05 | 1.28E-05 | acr-2/acr-3/unc-38/unc-63 | 4 |
| GO:1902656 | calcium ion import into cytosol | 4/243 | 5/9281 | 2.25E-06 | 2.11E-05 | 1.28E-05 | acr-2/acr-3/unc-38/unc-63 | 4 |
| GO:0048856 | anatomical structure development | 75/243 | 1735/9281 | 2.53E-06 | 2.37E-05 | 1.44E-05 | ced-1/daf-3/daf-5/daf-7/dnc-1/eat-16/[illegible] | 75 |
| GO:0051931 | regulation of sensory perception | 5/243 | 10/9281 | 2.67E-06 | 2.43E-05 | 1.48E-05 | mec-4/mec-7/mec-10/mec-12/mec-14 | 5 |
| GO:1905787 | regulation of detection of mechanical stimulus involved in sensory perception of touch | 5/243 | 10/9281 | 2.67E-06 | 2.43E-05 | 1.48E-05 | mec-4/mec-7/mec-10/mec-12/mec-14 | 5 |
| GO:1905789 | positive regulation of detection of mechanical stimulus involved in sensory perception of touch | 5/243 | 10/9281 | 2.67E-06 | 2.43E-05 | 1.48E-05 | mec-4/mec-7/mec-10/mec-12/mec-14 | 5 |
| GO:1905790 | regulation of mechanosensory behavior | 5/243 | 10/9281 | 2.67E-06 | 2.43E-05 | 1.48E-05 | mec-4/mec-7/mec-10/mec-12/mec-14 | 5 |
| GO:1905792 | positive regulation of mechanosensory behavior | 5/243 | 10/9281 | 2.67E-06 | 2.43E-05 | 1.48E-05 | mec-4/mec-7/mec-10/mec-12/mec-14 | 5 |
| GO:0032502 | developmental process | 78/243 | 1833/9281 | 2.68E-06 | 2.43E-05 | 1.48E-05 | ced-1/daf-3/daf-5/daf-7/dnc-1/eat-16/[illegible] | 78 |
| GO:0007189 | adenylate cyclase-activating G protein-coupled receptor signaling pathway | 6/243 | 17/9281 | 2.94E-06 | 2.66E-05 | 1.62E-05 | dop-1/gsa-1/ser-2/ser-5/tyra-2/dop-3 | 6 |
| GO:0030421 | defecation | 9/243 | 48/9281 | 3.43E-06 | 3.08E-05 | 1.88E-05 | egl-21/egl-30/flr-1/gon-2/gsa-1/unc-10/[illegible] | 9 |
| GO:0023056 | positive regulation of signaling | 16/243 | 158/9281 | 3.56E-06 | 3.19E-05 | 1.94E-05 | daf-7/egl-8/egl-21/egl-30/nca-2/sma-3/[illegible] | 16 |
| GO:0000902 | cell morphogenesis | 19/243 | 217/9281 | 3.86E-06 | 3.44E-05 | 2.09E-05 | dnc-1/egl-20/egl-46/rab-3/syg-1/unc-4/[illegible] | 19 |
| GO:0090287 | regulation of cellular response to growth factor stimulus | 7/243 | 27/9281 | 4.39E-06 | 3.89E-05 | 2.37E-05 | daf-5/lon-2/sma-3/tph-1/unc-2/unc-40/[illegible] | 7 |
| GO:0031646 | positive regulation of neurological system process | 5/243 | 11/9281 | 4.80E-06 | 4.23E-05 | 2.57E-05 | mec-4/mec-7/mec-10/mec-12/mec-14 | 5 |
| GO:0007588 | excretion | 9/243 | 50/9281 | 4.89E-06 | 4.30E-05 | 2.61E-05 | egl-21/egl-30/flr-1/gon-2/gsa-1/unc-10/[illegible] | 9 |
| GO:0048585 | negative regulation of response to stimulus | 16/243 | 163/9281 | 5.35E-06 | 4.68E-05 | 2.84E-05 | daf-5/dat-1/egl-10/goa-1/gsa-1/ins-18/[illegible] | 16 |
| GO:0019932 | second-messenger-mediated signaling | 10/243 | 64/9281 | 5.50E-06 | 4.79E-05 | 2.91E-05 | dop-1/flp-7/gsa-1/ser-1/ser-2/unc-89/[illegible] | 10 |
| GO:0050921 | positive regulation of chemotaxis | 7/243 | 28/9281 | 5.72E-06 | 4.96E-05 | 3.02E-05 | egl-20/pkc-1/rab-3/unc-31/unc-104/rig-6/[illegible] | 7 |
| GO:0006584 | catecholamine metabolic process | 4/243 | 6/9281 | 6.60E-06 | 5.59E-05 | 3.40E-05 | bas-1/cat-2/cat-4/tbh-1 | 4 |
| GO:0009712 | catechol-containing compound metabolic process | 4/243 | 6/9281 | 6.60E-06 | 5.59E-05 | 3.40E-05 | bas-1/cat-2/cat-4/tbh-1 | 4 |
| GO:0009713 | catechol-containing compound biosynthetic process | 4/243 | 6/9281 | 6.60E-06 | 5.59E-05 | 3.40E-05 | bas-1/cat-2/cat-4/tbh-1 | 4 |
| GO:0042417 | dopamine metabolic process | 4/243 | 6/9281 | 6.60E-06 | 5.59E-05 | 3.40E-05 | bas-1/cat-2/cat-4/tbh-1 | 4 |
| GO:0042423 | catecholamine biosynthetic process | 4/243 | 6/9281 | 6.60E-06 | 5.59E-05 | 3.40E-05 | bas-1/cat-2/cat-4/tbh-1 | 4 |
| GO:0035725 | sodium ion transmembrane transport | 10/243 | 66/9281 | 7.32E-06 | 6.15E-05 | 3.74E-05 | del-1/flr-1/mec-4/mec-10/unc-8/unc-105/[illegible] | 10 |
| GO:0006874 | cellular calcium ion homeostasis | 8/243 | 40/9281 | 7.33E-06 | 6.15E-05 | 3.74E-05 | acr-2/acr-3/spe-41/trp-1/trp-2/unc-38/[illegible] | 8 |
| GO:0050803 | regulation of synapse structure or activity | 9/243 | 53/9281 | 8.08E-06 | 6.76E-05 | 4.11E-05 | egl-1/egl-21/ins-4/unc-4/unc-37/unc-40/[illegible] | 9 |
| GO:0017015 | regulation of transforming growth factor beta receptor signaling pathway | 6/243 | 20/9281 | 8.63E-06 | 7.12E-05 | 4.33E-05 | daf-5/lon-2/sma-3/tph-1/unc-2/egl-4 | 6 |
| GO:0019933 | cAMP-mediated signaling | 6/243 | 20/9281 | 8.63E-06 | 7.12E-05 | 4.33E-05 | dop-1/gsa-1/ser-2/ser-5/tyra-2/dop-3 | 6 |
| GO:1903844 | regulation of cellular response to transforming growth factor beta stimulus | 6/243 | 20/9281 | 8.63E-06 | 7.12E-05 | 4.33E-05 | daf-5/lon-2/sma-3/tph-1/unc-2/egl-4 | 6 |
| GO:0055074 | calcium ion homeostasis | 8/243 | 41/9281 | 8.90E-06 | 7.31E-05 | 4.45E-05 | acr-2/acr-3/spe-41/trp-1/trp-2/unc-38/[illegible] | 8 |
| GO:0051128 | regulation of cellular component organization | 30/243 | 484/9281 | 8.99E-06 | 7.35E-05 | 4.47E-05 | ced-1/dnc-1/egl-1/egl-8/egl-15/egl-20/[illegible] | 30 |
| GO:0022604 | regulation of cell morphogenesis | 12/243 | 100/9281 | 1.08E-05 | 8.79E-05 | 5.35E-05 | dnc-1/egl-20/rab-3/unc-4/unc-31/unc-37/[illegible] | 12 |
| GO:0031644 | regulation of neurological system process | 5/243 | 13/9281 | 1.28E-05 | 1.04E-04 | 6.31E-05 | mec-4/mec-7/mec-10/mec-12/mec-14 | 5 |
| GO:0048520 | positive regulation of behavior | 9/243 | 56/9281 | 1.29E-05 | 1.04E-04 | 6.33E-05 | egl-21/mec-4/mec-7/mec-10/mec-12/[illegible] | 9 |
| GO:0006942 | regulation of striated muscle contraction | 4/243 | 7/9281 | 1.51E-05 | 1.20E-04 | 7.28E-05 | egl-19/flp-16/unc-89/unc-104 | 4 |
| GO:0061094 | regulation of turning behavior involved in mating | 4/243 | 7/9281 | 1.51E-05 | 1.20E-04 | 7.28E-05 | flp-10/flp-12/flp-20/mec-4 | 4 |
| GO:0061096 | negative regulation of turning behavior involved in mating | 4/243 | 7/9281 | 1.51E-05 | 1.20E-04 | 7.28E-05 | flp-10/flp-12/flp-20/mec-4 | 4 |
| GO:1902436 | negative regulation of male mating behavior | 4/243 | 7/9281 | 1.51E-05 | 1.20E-04 | 7.28E-05 | flp-10/flp-12/flp-20/mec-4 | 4 |
| GO:0050770 | regulation of axonogenesis | 10/243 | 72/9281 | 1.62E-05 | 1.28E-04 | 7.79E-05 | egl-20/rab-3/unc-4/unc-31/unc-37/unc-40/[illegible] | 10 |
| GO:0006585 | dopamine biosynthetic process from tyrosine | 3/243 | 3/9281 | 1.77E-05 | 1.38E-04 | 8.42E-05 | bas-1/cat-2/cat-4 | 3 |
| GO:0008039 | synaptic target recognition | 3/243 | 3/9281 | 1.77E-05 | 1.38E-04 | 8.42E-05 | syg-1/unc-4/syg-2 | 3 |
| GO:1901045 | negative regulation of oviposition | 3/243 | 3/9281 | 1.77E-05 | 1.38E-04 | 8.42E-05 | egl-6/flp-10/flp-17 | 3 |
| GO:0019935 | cyclic-nucleotide-mediated signaling | 7/243 | 33/9281 | 1.85E-05 | 1.42E-04 | 8.66E-05 | dop-1/gsa-1/ser-2/ser-5/tyra-2/dop-3/[illegible] | 7 |
| GO:0071241 | cellular response to inorganic substance | 7/243 | 33/9281 | 1.85E-05 | 1.42E-04 | 8.66E-05 | dat-1/egl-21/flp-16/jnk-1/nlp-1/nlp-14/[illegible] | 7 |
| GO:2000242 | negative regulation of reproductive process | 7/243 | 33/9281 | 1.85E-05 | 1.42E-04 | 8.66E-05 | egl-6/flp-10/flp-12/flp-17/flp-20/mec-4/[illegible] | 7 |
| GO:0007204 | positive regulation of cytosolic calcium ion concentration | 5/243 | 14/9281 | 1.95E-05 | 1.50E-04 | 9.10E-05 | acr-2/acr-3/unc-38/unc-63/ser-5 | 5 |
| GO:0070887 | cellular response to chemical stimulus | 30/243 | 504/9281 | 1.97E-05 | 1.51E-04 | 9.18E-05 | ced-1/daf-3/daf-5/daf-7/dat-1/egl-15/[illegible] | 30 |
| GO:0010942 | positive regulation of cell death | 9/243 | 59/9281 | 2.00E-05 | 1.52E-04 | 9.26E-05 | ced-1/dat-1/egl-1/jnk-1/unc-3/unc-26/[illegible] | 9 |
| GO:1903532 | positive regulation of secretion by cell | 6/243 | 23/9281 | 2.10E-05 | 1.60E-04 | 9.70E-05 | egl-8/egl-21/egl-30/unc-18/unc-26/unc-29 | 6 |
| GO:0006935 | chemotaxis | 17/243 | 202/9281 | 2.13E-05 | 1.61E-04 | 9.76E-05 | egl-8/egl-20/egl-30/egl-46/goa-1/odr-3/[illegible] | 17 |
| GO:0008037 | cell recognition | 7/243 | 34/9281 | 2.27E-05 | 1.71E-04 | 1.04E-04 | ced-1/syg-1/unc-4/unc-40/unc-76/syg-2/[illegible] | 7 |
| GO:0051656 | establishment of organelle localization | 16/243 | 184/9281 | 2.49E-05 | 1.87E-04 | 1.14E-04 | ced-1/dnc-1/egl-20/gpa-16/jnk-1/pkc-1/[illegible] | 16 |
| GO:0048638 | regulation of developmental growth | 14/243 | 145/9281 | 2.52E-05 | 1.88E-04 | 1.15E-04 | egl-21/ins-4/lon-2/sma-2/sma-3/unc-10/[illegible] | 14 |

| | | | | | | | | |
|---|---|---|---|---|---|---|---|---|
| GO:0009791 | post-embryonic development | 29/243 | 486/9281 | 2.62E-05 | 1.95E-04 | 1.19E-04 | daf-3/daf-5/daf-7/eat-16/egl-13/egl-15/egl-18/egl-20/egl-27/egl-30/gpa-3/hcf-1/ins-3/ins-4/ins-18/lin-39/pkc-1/sma-2/sma-3/tph-1/unc-3/unc-17/unc-32/unc-63/unc-98/unc-101/unc-104/unc-108/egl-4 | 29 |
| GO:0043269 | regulation of ion transport | 10/243 | 76/9281 | 2.63E-05 | 1.95E-04 | 1.19E-04 | egl-2/egl-8/egl-19/egl-21/egl-30/egl-36/unc-2/unc-29/unc-103/egl-4 | 10 |
| GO:0048589 | developmental growth | 16/243 | 185/9281 | 2.67E-05 | 1.97E-04 | 1.20E-04 | egl-21/ins-4/lon-2/sma-2/sma-3/syg-1/unc-10/unc-26/unc-40/unc-44/unc-69/unc-76/unc-104/syg-2/rig-6/egl-4 | 16 |
| GO:0050909 | sensory perception of taste | 5/243 | 15/9281 | 2.86E-05 | 2.10E-04 | 1.28E-04 | egl-47/gpa-3/gpc-1/lite-1/egl-4 | 5 |
| GO:0030512 | negative regulation of transforming growth factor beta receptor signaling pathway | 4/243 | 8/9281 | 2.95E-05 | 2.15E-04 | 1.31E-04 | daf-5/lon-2/unc-2/egl-4 | 4 |
| GO:1902435 | regulation of male mating behavior | 4/243 | 8/9281 | 2.95E-05 | 2.15E-04 | 1.31E-04 | flp-10/flp-12/flp-20/mec-4 | 4 |
| GO:1903845 | negative regulation of cellular response to transforming growth factor beta stimulus | 4/243 | 8/9281 | 2.95E-05 | 2.15E-04 | 1.31E-04 | daf-5/lon-2/unc-2/egl-4 | 4 |
| GO:0072503 | cellular divalent inorganic cation homeostasis | 8/243 | 48/9281 | 3.00E-05 | 2.17E-04 | 1.32E-04 | acr-2/acr-3/spe-41/trp-1/trp-2/unc-38/unc-63/ser-5 | 8 |
| GO:0071310 | cellular response to organic substance | 24/243 | 367/9281 | 3.20E-05 | 2.31E-04 | 1.40E-04 | ced-1/daf-3/daf-5/daf-7/dat-1/egl-15/egl-17/egl-20/gar-2/goa-1/gpa-3/gpb-2/ins-18/jnk-1/lon-2/pmk-1/sma-2/sma-3/octr-1/tph-1/unc-2/unc-40/unc-55/egl-4 | 24 |
| GO:0010817 | regulation of hormone levels | 7/243 | 36/9281 | 3.38E-05 | 2.42E-04 | 1.47E-04 | egl-8/egl-21/egl-30/snf-6/unc-17/unc-26/unc-29 | 7 |
| GO:0045597 | positive regulation of cell differentiation | 11/243 | 95/9281 | 3.55E-05 | 2.54E-04 | 1.54E-04 | egl-8/egl-20/egl-30/gld-1/gsa-1/lin-39/rab-3/unc-31/unc-89/unc-104/rig-6 | 11 |
| GO:1901699 | cellular response to nitrogen compound | 8/243 | 50/9281 | 4.08E-05 | 2.91E-04 | 1.77E-04 | daf-7/dat-1/gar-2/goa-1/gpa-3/gpb-2/ins-18/egl-4 | 8 |
| GO:0040024 | dauer larval development | 12/243 | 114/9281 | 4.11E-05 | 2.92E-04 | 1.77E-04 | daf-3/daf-5/daf-7/gpa-3/hcf-1/ins-3/ins-4/ins-18/pkc-1/tph-1/unc-3/egl-4 | 12 |
| GO:0006814 | sodium ion transport | 10/243 | 80/9281 | 4.14E-05 | 2.92E-04 | 1.78E-04 | del-1/flr-1/mec-4/mec-10/unc-8/unc-105/tmc-1/asic-1/delm-2/egas-2 | 10 |
| GO:0010720 | positive regulation of cell development | 10/243 | 80/9281 | 4.14E-05 | 2.92E-04 | 1.78E-04 | egl-8/egl-20/egl-30/gld-1/gsa-1/rab-3/unc-31/unc-89/unc-104/rig-6 | 10 |
| GO:0031987 | locomotion involved in locomotory behavior | 6/243 | 26/9281 | 4.49E-05 | 3.15E-04 | 1.92E-04 | bas-1/egl-21/ric-19/unc-31/unc-108/rig-6 | 6 |
| GO:0050808 | synapse organization | 10/243 | 81/9281 | 4.62E-05 | 3.23E-04 | 1.97E-04 | egl-1/egl-21/ins-4/syg-1/unc-4/unc-40/unc-69/unc-76/unc-104/syg-2 | 10 |
| GO:0061065 | regulation of dauer larval development | 7/243 | 38/9281 | 4.88E-05 | 3.40E-04 | 2.07E-04 | daf-3/daf-5/daf-7/ins-3/ins-4/ins-18/egl-4 | 7 |
| GO:0051962 | positive regulation of nervous system development | 9/243 | 66/9281 | 5.02E-05 | 3.49E-04 | 2.12E-04 | egl-8/egl-20/egl-21/egl-30/rab-3/unc-31/unc-40/unc-104/rig-6 | 9 |
| GO:0043900 | regulation of multi-organism process | 10/243 | 82/9281 | 5.14E-05 | 3.56E-04 | 2.16E-04 | egl-30/flp-10/flp-12/flp-20/gld-1/gpa-12/gsa-1/mec-4/pmk-1/rig-6 | 10 |
| GO:0072507 | divalent inorganic cation homeostasis | 8/243 | 52/9281 | 5.47E-05 | 3.77E-04 | 2.29E-04 | acr-2/acr-3/spe-41/trp-1/trp-2/unc-38/unc-63/ser-5 | 8 |
| GO:0042136 | neurotransmitter biosynthetic process | 5/243 | 17/9281 | 5.65E-05 | 3.88E-04 | 2.36E-04 | bas-1/cat-2/tbh-1/tdc-1/tph-1 | 5 |
| GO:0031667 | response to nutrient levels | 9/243 | 68/9281 | 6.39E-05 | 4.36E-04 | 2.65E-04 | cat-2/dop-1/dop-2/eat-16/goa-1/gpb-2/mod-1/unc-2/dop-3 | 9 |
| GO:0050920 | regulation of chemotaxis | 9/243 | 68/9281 | 6.39E-05 | 4.36E-04 | 2.65E-04 | egl-20/egl-30/pkc-1/rab-3/unc-31/unc-40/unc-104/rig-6/egl-4 | 9 |
| GO:0007167 | enzyme linked receptor protein signaling pathway | 14/243 | 158/9281 | 6.56E-05 | 4.46E-04 | 2.71E-04 | daf-3/daf-5/daf-7/egl-15/egl-17/ins-18/lon-2/pkc-1/sma-2/sma-3/tph-1/unc-2/unc-40/egl-4 | 14 |
| GO:0001504 | neurotransmitter uptake | 3/243 | 4/9281 | 6.96E-05 | 4.68E-04 | 2.84E-04 | dat-1/mod-5/snf-6 | 3 |
| GO:0007211 | octopamine or tyramine signaling pathway | 3/243 | 4/9281 | 6.96E-05 | 4.68E-04 | 2.84E-04 | ser-2/octr-1/ser-6 | 3 |
| GO:0032230 | positive regulation of synaptic transmission, GABAergic | 3/243 | 4/9281 | 6.96E-05 | 4.68E-04 | 2.84E-04 | nca-2/unc-75/unc-77 | 3 |
| GO:1902669 | positive regulation of axon guidance | 5/243 | 18/9281 | 7.65E-05 | 5.13E-04 | 3.12E-04 | egl-20/rab-3/unc-31/unc-104/rig-6 | 5 |
| GO:0010648 | negative regulation of cell communication | 13/243 | 141/9281 | 7.96E-05 | 5.30E-04 | 3.22E-04 | daf-5/egl-10/gar-2/goa-1/gsa-1/ins-18/lon-2/osm-9/rgs-6/unc-2/unc-37/unc-122/egl-4 | 13 |
| GO:0023057 | negative regulation of signaling | 13/243 | 141/9281 | 7.96E-05 | 5.30E-04 | 3.22E-04 | daf-5/egl-10/gar-2/goa-1/gsa-1/ins-18/lon-2/osm-9/rgs-6/unc-2/unc-37/unc-122/egl-4 | 13 |
| GO:0051640 | organelle localization | 18/243 | 247/9281 | 8.25E-05 | 5.47E-04 | 3.32E-04 | ced-1/dnc-1/egl-20/gpa-16/jnk-1/pkc-1/rab-3/sng-1/tom-1/unc-4/unc-10/unc-14/unc-16/unc-18/unc-26/unc-31/unc-104/unc-116 | 18 |
| GO:0090101 | negative regulation of transmembrane receptor protein serine/threonine kinase signaling pathway | 4/243 | 10/9281 | 8.50E-05 | 5.62E-04 | 3.41E-04 | daf-5/lon-2/unc-2/egl-4 | 4 |
| GO:0040013 | negative regulation of locomotion | 6/243 | 29/9281 | 8.67E-05 | 5.69E-04 | 3.46E-04 | eat-16/egl-20/flp-7/unc-32/unc-40/rig-6 | 6 |
| GO:0051047 | positive regulation of secretion | 6/243 | 29/9281 | 8.67E-05 | 5.69E-04 | 3.46E-04 | egl-8/egl-21/egl-30/unc-18/unc-26/unc-29 | 6 |
| GO:0009991 | response to extracellular stimulus | 9/243 | 71/9281 | 9.02E-05 | 5.89E-04 | 3.58E-04 | cat-2/dop-1/dop-2/eat-16/goa-1/gpb-2/mod-1/unc-2/dop-3 | 9 |
| GO:0040014 | regulation of multicellular organism growth | 10/243 | 88/9281 | 9.46E-05 | 6.16E-04 | 3.75E-04 | lon-2/sma-2/sma-3/unc-10/unc-26/unc-40/unc-44/unc-104/rig-6/egl-4 | 10 |
| GO:0048639 | positive regulation of developmental growth | 9/243 | 72/9281 | 1.01E-04 | 6.54E-04 | 3.98E-04 | egl-21/sma-2/sma-3/unc-10/unc-26/unc-40/unc-44/unc-104/rig-6 | 9 |
| GO:0031099 | regeneration | 6/243 | 30/9281 | 1.06E-04 | 6.82E-04 | 4.14E-04 | egl-8/egl-30/goa-1/jnk-1/tph-1/unc-34 | 6 |
| GO:0031102 | neuron projection regeneration | 6/243 | 30/9281 | 1.06E-04 | 6.82E-04 | 4.14E-04 | egl-8/egl-30/goa-1/jnk-1/tph-1/unc-34 | 6 |
| GO:0031103 | axon regeneration | 6/243 | 30/9281 | 1.06E-04 | 6.82E-04 | 4.14E-04 | egl-8/egl-30/goa-1/jnk-1/tph-1/unc-34 | 6 |
| GO:0008038 | neuron recognition | 6/243 | 31/9281 | 1.29E-04 | 8.19E-04 | 4.98E-04 | syg-1/unc-4/unc-40/unc-76/syg-2/rig-6 | 6 |
| GO:0060341 | regulation of cellular localization | 13/243 | 148/9281 | 1.31E-04 | 8.19E-04 | 4.98E-04 | egl-8/egl-21/egl-30/pmk-1/ric-19/tom-1/unc-10/unc-17/unc-18/unc-26/unc-29/unc-89/unc-104 | 13 |
| GO:0014057 | positive regulation of acetylcholine secretion, neurotransmission | 4/243 | 11/9281 | 1.31E-04 | 8.19E-04 | 4.98E-04 | egl-8/egl-21/egl-30/unc-29 | 4 |
| GO:0030536 | larval feeding behavior | 4/243 | 11/9281 | 1.31E-04 | 8.19E-04 | 4.98E-04 | daf-7/egl-21/unc-31/egl-4 | 4 |
| GO:0035545 | determination of left/right asymmetry in nervous system | 4/243 | 11/9281 | 1.31E-04 | 8.19E-04 | 4.98E-04 | egl-19/unc-2/unc-40/rig-6 | 4 |
| GO:0051954 | positive regulation of amine transport | 4/243 | 11/9281 | 1.31E-04 | 8.19E-04 | 4.98E-04 | egl-8/egl-21/egl-30/unc-29 | 4 |
| GO:0090288 | negative regulation of cellular response to growth factor stimulus | 4/243 | 11/9281 | 1.31E-04 | 8.19E-04 | 4.98E-04 | daf-5/lon-2/unc-2/egl-4 | 4 |
| GO:2001025 | positive regulation of response to drug | 4/243 | 11/9281 | 1.31E-04 | 8.19E-04 | 4.98E-04 | egl-8/egl-21/egl-30/unc-29 | 4 |
| GO:2001023 | regulation of response to drug | 5/243 | 20/9281 | 1.33E-04 | 8.27E-04 | 5.03E-04 | egl-8/egl-21/egl-30/pkc-1/unc-29 | 5 |
| GO:0040007 | growth | 16/243 | 212/9281 | 1.37E-04 | 8.46E-04 | 5.15E-04 | egl-21/ins-4/lon-2/sma-2/sma-3/syg-1/unc-10/unc-26/unc-40/unc-44/unc-69/unc-76/unc-104/syg-2/rig-6/egl-4 | 16 |
| GO:0006873 | cellular ion homeostasis | 11/243 | 110/9281 | 1.37E-04 | 8.46E-04 | 5.15E-04 | acr-2/acr-3/ftn-2/spe-41/trp-1/trp-2/unc-32/unc-38/unc-63/unc-108/ser-5 | 11 |
| GO:0030003 | cellular cation homeostasis | 11/243 | 110/9281 | 1.37E-04 | 8.46E-04 | 5.15E-04 | acr-2/acr-3/ftn-2/spe-41/trp-1/trp-2/unc-32/unc-38/unc-63/unc-108/ser-5 | 11 |
| GO:0055080 | cation homeostasis | 12/243 | 129/9281 | 1.38E-04 | 8.47E-04 | 5.15E-04 | acr-2/acr-3/ftn-2/spe-41/trp-1/trp-2/unc-32/unc-38/unc-63/unc-80/unc-108/ser-5 | 12 |
| GO:0009653 | anatomical structure morphogenesis | 31/243 | 588/9281 | 1.42E-04 | 8.69E-04 | 5.28E-04 | dnc-1/egl-5/egl-13/egl-15/egl-20/egl-27/egl-46/lin-39/lon-2/rab-3/sem-4/sma-2/sma-3/syg-1/unc-4/unc-30/unc-31/unc-34/unc-37/unc-40/unc-44/unc-60/unc-69/unc-76/unc-89/unc-98/unc-104/unc-115/unc-116/syg-2/rig-6 | 31 |
| GO:0007416 | synapse assembly | 7/243 | 45/9281 | 1.50E-04 | 9.15E-04 | 5.56E-04 | egl-21/ins-4/syg-1/unc-4/unc-40/unc-104/syg-2 | 7 |
| GO:0010976 | positive regulation of neuron projection development | 7/243 | 45/9281 | 1.50E-04 | 9.15E-04 | 5.56E-04 | egl-8/egl-20/egl-30/rab-3/unc-31/unc-104/rig-6 | 7 |
| GO:0002119 | nematode larval development | 26/243 | 457/9281 | 1.53E-04 | 9.32E-04 | 5.67E-04 | daf-3/daf-5/daf-7/eat-16/egl-13/egl-15/egl-18/egl-20/egl-27/egl-30/gpa-3/hcf-1/ins-3/ins-4/ins-18/lin-39/pkc-1/sma-3/tph-1/unc-3/unc-17/unc-32/unc-101/unc-104/unc-108/egl-4 | 26 |
| GO:0048678 | response to axon injury | 6/243 | 32/9281 | 1.55E-04 | 9.38E-04 | 5.70E-04 | egl-8/egl-30/goa-1/jnk-1/tph-1/unc-34 | 6 |
| GO:0050801 | ion homeostasis | 12/243 | 131/9281 | 1.59E-04 | 9.59E-04 | 5.83E-04 | acr-2/acr-3/ftn-2/spe-41/trp-1/trp-2/unc-32/unc-38/unc-63/unc-80/unc-108/ser-5 | 12 |
| GO:0051270 | regulation of cellular component movement | 12/243 | 131/9281 | 1.59E-04 | 9.59E-04 | 5.83E-04 | ced-1/egl-17/egl-20/egl-27/egl-43/lon-2/rab-3/snf-10/unc-31/unc-40/unc-104/rig-6 | 12 |
| GO:0040008 | regulation of growth | 14/243 | 172/9281 | 1.63E-04 | 9.80E-04 | 5.96E-04 | egl-21/ins-4/lon-2/sma-2/sma-3/unc-10/unc-26/unc-40/unc-44/unc-69/unc-76/unc-104/rig-6/egl-4 | 14 |
| GO:0006875 | cellular metal ion homeostasis | 9/243 | 77/9281 | 1.70E-04 | 1.01E-03 | 6.12E-04 | acr-2/acr-3/ftn-2/spe-41/trp-1/trp-2/unc-38/unc-63/ser-5 | 9 |
| GO:0015837 | amine transport | 5/243 | 21/9281 | 1.70E-04 | 1.01E-03 | 6.12E-04 | dat-1/egl-8/egl-21/egl-30/unc-29 | 5 |
| GO:0090325 | regulation of locomotion involved in locomotory behavior | 5/243 | 21/9281 | 1.70E-04 | 1.01E-03 | 6.12E-04 | egl-21/ric-19/unc-31/unc-108/rig-6 | 5 |
| GO:0032223 | negative regulation of synaptic transmission, cholinergic | 3/243 | 5/9281 | 1.71E-04 | 1.01E-03 | 6.12E-04 | gar-2/goa-1/unc-122 | 3 |
| GO:0042416 | dopamine biosynthetic process | 3/243 | 5/9281 | 1.71E-04 | 1.01E-03 | 6.12E-04 | bas-1/cat-2/cat-4 | 3 |
| GO:0045666 | positive regulation of neuron differentiation | 7/243 | 46/9281 | 1.73E-04 | 1.02E-03 | 6.18E-04 | egl-8/egl-20/egl-30/rab-3/unc-31/unc-104/rig-6 | 7 |
| GO:0050807 | regulation of synapse organization | 7/243 | 46/9281 | 1.73E-04 | 1.02E-03 | 6.18E-04 | egl-1/egl-21/ins-4/unc-40/unc-69/unc-76/unc-104 | 7 |
| GO:0002164 | larval development | 26/243 | 462/9281 | 1.83E-04 | 1.07E-03 | 6.49E-04 | daf-3/daf-5/daf-7/eat-16/egl-13/egl-15/egl-18/egl-20/egl-27/egl-30/gpa-3/hcf-1/ins-3/ins-4/ins-18/lin-39/pkc-1/sma-3/tph-1/unc-3/unc-17/unc-32/unc-101/unc-104/unc-108/egl-4 | 26 |
| GO:0014056 | regulation of acetylcholine secretion, neurotransmission | 4/243 | 12/9281 | 1.92E-04 | 1.11E-03 | 6.73E-04 | egl-8/egl-21/egl-30/unc-29 | 4 |
| GO:0032228 | regulation of synaptic transmission, GABAergic | 4/243 | 12/9281 | 1.92E-04 | 1.11E-03 | 6.73E-04 | nca-2/rab-3/unc-75/unc-77 | 4 |
| GO:0046887 | positive regulation of hormone secretion | 4/243 | 12/9281 | 1.92E-04 | 1.11E-03 | 6.73E-04 | egl-8/egl-21/egl-30/unc-29 | 4 |
| GO:0060402 | calcium ion transport into cytosol | 4/243 | 12/9281 | 1.92E-04 | 1.11E-03 | 6.73E-04 | acr-2/acr-3/unc-38/unc-63 | 4 |
| GO:0097553 | calcium ion transmembrane import into cytosol | 4/243 | 12/9281 | 1.92E-04 | 1.11E-03 | 6.73E-04 | acr-2/acr-3/unc-38/unc-63 | 4 |
| GO:0051130 | positive regulation of cellular component organization | 15/243 | 197/9281 | 2.01E-04 | 1.16E-03 | 7.02E-04 | ced-1/egl-1/egl-8/egl-20/egl-21/egl-30/egl-44/egl-46/rab-3/unc-31/unc-40/unc-60/unc-89/unc-104/rig-6 | 15 |
| GO:0055082 | cellular chemical homeostasis | 11/243 | 117/9281 | 2.38E-04 | 1.36E-03 | 8.27E-04 | acr-2/acr-3/ftn-2/spe-41/trp-1/trp-2/unc-32/unc-38/unc-63/unc-108/ser-5 | 11 |
| GO:0035264 | multicellular organism growth | 10/243 | 99/9281 | 2.54E-04 | 1.45E-03 | 8.81E-04 | lon-2/sma-2/sma-3/unc-10/unc-26/unc-40/unc-44/unc-104/rig-6/egl-4 | 10 |
| GO:0040018 | positive regulation of multicellular organism growth | 7/243 | 49/9281 | 2.60E-04 | 1.47E-03 | 8.97E-04 | sma-2/sma-3/unc-10/unc-26/unc-40/unc-44/unc-104 | 7 |
| GO:0051272 | positive regulation of cellular component movement | 7/243 | 49/9281 | 2.60E-04 | 1.47E-03 | 8.97E-04 | ced-1/egl-20/rab-3/snf-10/unc-31/unc-104/rig-6 | 7 |
| GO:0006941 | striated muscle contraction | 4/243 | 13/9281 | 2.72E-04 | 1.53E-03 | 9.30E-04 | egl-19/flp-16/unc-89/unc-104 | 4 |
| GO:0014055 | acetylcholine secretion, neurotransmission | 4/243 | 13/9281 | 2.72E-04 | 1.53E-03 | 9.30E-04 | egl-8/egl-21/egl-30/unc-29 | 4 |
| GO:0061526 | acetylcholine secretion | 4/243 | 13/9281 | 2.72E-04 | 1.53E-03 | 9.30E-04 | egl-8/egl-21/egl-30/unc-29 | 4 |
| GO:0010941 | regulation of cell death | 11/243 | 119/9281 | 2.76E-04 | 1.55E-03 | 9.40E-04 | ced-1/daf-7/dat-1/egl-1/jnk-1/pmk-1/unc-3/unc-26/unc-32/unc-104/unc-116 | 11 |
| GO:0015672 | monovalent inorganic cation transport | 16/243 | 228/9281 | 3.15E-04 | 1.76E-03 | 1.07E-03 | del-1/egl-2/egl-23/egl-36/flr-1/mec-4/mec-10/sup-9/unc-8/unc-32/unc-103/unc-105/tmc-1/asic-1/delm-2/egas-2 | 16 |
| GO:0010035 | response to inorganic substance | 11/243 | 121/9281 | 3.19E-04 | 1.78E-03 | 1.08E-03 | dat-1/egl-21/flp-16/ftn-2/hcf-1/jnk-1/nlp-1/nlp-14/pmk-1/unc-31/egl-4 | 11 |
| GO:0016049 | cell growth | 9/243 | 84/9281 | 3.32E-04 | 1.84E-03 | 1.12E-03 | sma-2/syg-1/unc-40/unc-69/unc-76/unc-104/syg-2/rig-6/egl-4 | 9 |
| GO:0033563 | dorsal/ventral axon guidance | 5/243 | 24/9281 | 3.34E-04 | 1.84E-03 | 1.12E-03 | unc-30/unc-34/unc-40/unc-44/rig-6 | 5 |
| GO:0008277 | regulation of G protein-coupled receptor signaling pathway | 3/243 | 6/9281 | 3.34E-04 | 1.84E-03 | 1.12E-03 | eat-16/egl-10/gpb-2 | 3 |
| GO:0031346 | positive regulation of cell projection organization | 7/243 | 51/9281 | 3.35E-04 | 1.84E-03 | 1.12E-03 | egl-8/egl-20/egl-30/rab-3/unc-31/unc-104/rig-6 | 7 |
| GO:1901617 | organic hydroxy compound biosynthetic process | 7/243 | 51/9281 | 3.35E-04 | 1.84E-03 | 1.12E-03 | bas-1/cat-2/cat-4/egl-8/tbh-1/tdc-1/tph-1 | 7 |
| GO:0051093 | negative regulation of developmental process | 11/243 | 123/9281 | 3.68E-04 | 2.01E-03 | 1.22E-03 | daf-5/daf-7/egl-20/goa-1/ins-3/ins-4/jnk-1/lon-2/unc-40/rig-6/egl-4 | 11 |
| GO:0007274 | neuromuscular synaptic transmission | 4/243 | 14/9281 | 3.73E-04 | 2.01E-03 | 1.22E-03 | acr-2/snf-6/unc-29/unc-38 | 4 |
| GO:0007368 | determination of left/right symmetry | 4/243 | 14/9281 | 3.73E-04 | 2.01E-03 | 1.22E-03 | egl-19/unc-2/unc-40/rig-6 | 4 |
| GO:0009799 | specification of symmetry | 4/243 | 14/9281 | 3.73E-04 | 2.01E-03 | 1.22E-03 | egl-19/unc-2/unc-40/rig-6 | 4 |
| GO:0009855 | determination of bilateral symmetry | 4/243 | 14/9281 | 3.73E-04 | 2.01E-03 | 1.22E-03 | egl-19/unc-2/unc-40/rig-6 | 4 |
| GO:0046883 | regulation of hormone secretion | 4/243 | 14/9281 | 3.73E-04 | 2.01E-03 | 1.22E-03 | egl-8/egl-21/egl-30/unc-29 | 4 |
| GO:0060401 | cytosolic calcium ion transport | 4/243 | 14/9281 | 3.73E-04 | 2.01E-03 | 1.22E-03 | acr-2/acr-3/unc-38/unc-63 | 4 |
| GO:0050769 | positive regulation of neurogenesis | 7/243 | 52/9281 | 3.79E-04 | 2.03E-03 | 1.24E-03 | egl-8/egl-20/egl-30/rab-3/unc-31/unc-104/rig-6 | 7 |
| GO:0007612 | learning | 6/243 | 38/9281 | 4.14E-04 | 2.21E-03 | 1.35E-03 | cat-2/dop-1/gpa-2/osm-9/pkc-1/egl-4 | 6 |
| GO:0001707 | mesoderm formation | 6/243 | 39/9281 | 4.79E-04 | 2.54E-03 | 1.54E-03 | lin-39/lon-2/sem-4/sma-2/sma-3/unc-40 | 6 |
| GO:0048332 | mesoderm morphogenesis | 6/243 | 39/9281 | 4.79E-04 | 2.54E-03 | 1.54E-03 | lin-39/lon-2/sem-4/sma-2/sma-3/unc-40 | 6 |
| GO:0048333 | mesodermal cell differentiation | 6/243 | 39/9281 | 4.79E-04 | 2.54E-03 | 1.54E-03 | lin-39/lon-2/sem-4/sma-2/sma-3/unc-40 | 6 |
| GO:0030537 | larval behavior | 4/243 | 15/9281 | 4.98E-04 | 2.61E-03 | 1.59E-03 | daf-7/egl-21/unc-31/egl-4 | 4 |
| GO:0045933 | positive regulation of muscle contraction | 4/243 | 15/9281 | 4.98E-04 | 2.61E-03 | 1.59E-03 | egl-19/unc-26/unc-89/ser-5 | 4 |
| GO:0046879 | hormone secretion | 4/243 | 15/9281 | 4.98E-04 | 2.61E-03 | 1.59E-03 | egl-8/egl-21/egl-30/unc-29 | 4 |
| GO:1905486 | regulation of anterior/posterior axon guidance | 4/243 | 15/9281 | 4.98E-04 | 2.61E-03 | 1.59E-03 | egl-20/rab-3/unc-31/unc-104 | 4 |
| GO:1905488 | positive regulation of anterior/posterior axon guidance | 4/243 | 15/9281 | 4.98E-04 | 2.61E-03 | 1.59E-03 | egl-20/rab-3/unc-31/unc-104 | 4 |
| GO:0006887 | exocytosis | 8/243 | 71/9281 | 5.04E-04 | 2.63E-03 | 1.60E-03 | pkc-1/rab-3/sng-1/tom-1/unc-10/unc-18/unc-31/unc-104 | 8 |
| GO:0051649 | establishment of localization in cell | 31/243 | 633/9281 | 5.16E-04 | 2.69E-03 | 1.63E-03 | acr-2/acr-3/ced-1/dat-1/dnc-1/egl-8/egl-20/egl-21/egl-30/gpa-16/jnk-1/mod-5/pkc-1/rab-3/ric-19/sng-1/tom-1/unc-4/unc-10/unc-16/unc-17/unc-18/unc-26/unc-29/unc-31/unc-38/unc-63/unc-101/unc-104/unc-108/unc-116 | 31 |
| GO:0098771 | inorganic ion homeostasis | 11/243 | 128/9281 | 5.18E-04 | 2.69E-03 | 1.63E-03 | acr-2/acr-3/ftn-2/spe-41/trp-1/trp-2/unc-32/unc-38/unc-63/unc-108/ser-5 | 11 |
| GO:0007611 | learning or memory | 7/243 | 55/9281 | 5.37E-04 | 2.78E-03 | 1.69E-03 | cat-2/dop-1/gpa-2/osm-9/pkc-1/unc-43/egl-4 | 7 |
| GO:0009611 | response to wounding | 6/243 | 40/9281 | 5.51E-04 | 2.84E-03 | 1.73E-03 | egl-8/egl-30/goa-1/jnk-1/tph-1/unc-34 | 6 |
| GO:0006828 | manganese ion transport | 3/243 | 7/9281 | 5.74E-04 | 2.95E-03 | 1.79E-03 | spe-41/trp-1/trp-2 | 3 |
| GO:1990504 | dense core granule exocytosis | 3/243 | 7/9281 | 5.74E-04 | 2.95E-03 | 1.79E-03 | pkc-1/tom-1/unc-31 | 3 |
| GO:0006972 | hyperosmotic response | 5/243 | 27/9281 | 5.95E-04 | 3.02E-03 | 1.84E-03 | flp-1/jnk-1/odr-3/osm-9/pmk-1 | 5 |
| GO:0016079 | synaptic vesicle exocytosis | 5/243 | 27/9281 | 5.95E-04 | 3.02E-03 | 1.84E-03 | pkc-1/tom-1/unc-10/unc-18/unc-31 | 5 |
| GO:0042133 | neurotransmitter metabolic process | 5/243 | 27/9281 | 5.95E-04 | 3.02E-03 | 1.84E-03 | bas-1/cat-2/tbh-1/tdc-1/tph-1 | 5 |
| GO:0043901 | negative regulation of multi-organism process | 5/243 | 27/9281 | 5.95E-04 | 3.02E-03 | 1.84E-03 | flp-10/flp-12/flp-20/mec-4/rig-6 | 5 |
| GO:0045165 | cell fate commitment | 15/243 | 219/9281 | 6.25E-04 | 3.16E-03 | 1.92E-03 | egl-13/egl-18/egl-20/egl-27/egl-44/egl-46/gld-1/lin-39/lon-2/sma-2/sma-3/unc-4/unc-40/unc-42/unc-86 | 15 |
| GO:0007411 | axon guidance | 11/243 | 131/9281 | 6.30E-04 | 3.18E-03 | 1.93E-03 | egl-20/egl-46/rab-3/unc-30/unc-31/unc-34/unc-40/unc-44/unc-104/unc-116/rig-6 | 11 |
| GO:0045055 | regulated exocytosis | 6/243 | 41/9281 | 6.31E-04 | 3.18E-03 | 1.93E-03 | pkc-1/tom-1/unc-10/unc-18/unc-31/unc-104 | 6 |
| GO:0097485 | neuron projection guidance | 11/243 | 132/9281 | 6.72E-04 | 3.38E-03 | 2.05E-03 | egl-20/egl-46/rab-3/unc-30/unc-31/unc-34/unc-40/unc-44/unc-104/unc-116/rig-6 | 11 |
| GO:0006589 | octopamine biosynthetic process | 2/243 | 2/9281 | 6.83E-04 | 3.39E-03 | 2.06E-03 | tbh-1/tdc-1 | 2 |
| GO:0031536 | positive regulation of exit from mitosis | 2/243 | 2/9281 | 6.83E-04 | 3.39E-03 | 2.06E-03 | egl-44/egl-46 | 2 |
| GO:0046333 | octopamine metabolic process | 2/243 | 2/9281 | 6.83E-04 | 3.39E-03 | 2.06E-03 | tbh-1/tdc-1 | 2 |
| GO:0071927 | octopamine signaling pathway | 2/243 | 2/9281 | 6.83E-04 | 3.39E-03 | 2.06E-03 | octr-1/ser-6 | 2 |
| GO:1903554 | G protein-coupled receptor signaling pathway involved in defense response to Gram-negative bacterium | 2/243 | 2/9281 | 6.83E-04 | 3.39E-03 | 2.06E-03 | gpa-2/gpa-3 | 2 |
| GO:0045927 | positive regulation of growth | 9/243 | 93/9281 | 7.07E-04 | 3.50E-03 | 2.13E-03 | egl-21/sma-2/sma-3/unc-10/unc-26/unc-40/unc-44/unc-104/rig-6 | 9 |
| GO:0055065 | metal ion homeostasis | 9/243 | 94/9281 | 7.64E-04 | 3.77E-03 | 2.29E-03 | acr-2/acr-3/ftn-2/spe-41/trp-1/trp-2/unc-38/unc-63/ser-5 | 9 |
| GO:0006928 | movement of cell or subcellular component | 20/243 | 349/9281 | 8.26E-04 | 4.04E-03 | 2.46E-03 | ced-1/dnc-1/egl-15/egl-17/egl-20/egl-27/egl-43/egl-46/jnk-1/lon-2/rab-3/snf-10/unc-30/unc-31/unc-34/unc-40/unc-44/unc-104/unc-116/rig-6 | 20 |
| GO:0007498 | mesoderm development | 7/243 | 59/9281 | 8.27E-04 | 4.04E-03 | 2.46E-03 | egl-27/lin-39/lon-2/sem-4/sma-2/sma-3/unc-40 | 7 |
| GO:0050890 | cognition | 7/243 | 59/9281 | 8.27E-04 | 4.04E-03 | 2.46E-03 | cat-2/dop-1/gpa-2/osm-9/pkc-1/unc-43/egl-4 | 7 |
| GO:0090100 | positive regulation of transmembrane receptor protein serine/threonine kinase signaling pathway | 4/243 | 17/9281 | 8.34E-04 | 4.04E-03 | 2.46E-03 | daf-7/sma-3/tph-1/unc-40 | 4 |
| GO:0035418 | protein localization to synapse | 5/243 | 29/9281 | 8.38E-04 | 4.04E-03 | 2.46E-03 | egl-8/egl-30/unc-26/unc-104/syg-2 | 5 |
| GO:0042661 | regulation of mesodermal cell fate specification | 5/243 | 29/9281 | 8.38E-04 | 4.04E-03 | 2.46E-03 | lin-39/lon-2/sma-2/sma-3/unc-40 | 5 |
| GO:1905770 | regulation of mesodermal cell differentiation | 5/243 | 29/9281 | 8.38E-04 | 4.04E-03 | 2.46E-03 | lin-39/lon-2/sma-2/sma-3/unc-40 | 5 |
| GO:1905902 | regulation of mesoderm formation | 5/243 | 29/9281 | 8.38E-04 | 4.04E-03 | 2.46E-03 | lin-39/lon-2/sma-2/sma-3/unc-40 | 5 |
| GO:2000380 | regulation of mesoderm development | 5/243 | 29/9281 | 8.38E-04 | 4.04E-03 | 2.46E-03 | lin-39/lon-2/sma-2/sma-3/unc-40 | 5 |
| GO:0001966 | thigmotaxis | 3/243 | 8/9281 | 9.01E-04 | 4.24E-03 | 2.58E-03 | flp-10/mec-12/nlp-1 | 3 |
| GO:0007213 | G protein-coupled acetylcholine receptor signaling pathway | 3/243 | 8/9281 | 9.01E-04 | 4.24E-03 | 2.58E-03 | gar-2/goa-1/gpb-2 | 3 |
| GO:0050805 | negative regulation of synaptic transmission | 3/243 | 8/9281 | 9.01E-04 | 4.24E-03 | 2.58E-03 | gar-2/goa-1/unc-122 | 3 |
| GO:0071242 | cellular response to ammonium ion | 3/243 | 8/9281 | 9.01E-04 | 4.24E-03 | 2.58E-03 | gar-2/goa-1/gpb-2 | 3 |
| GO:0095500 | acetylcholine receptor signaling pathway | 3/243 | 8/9281 | 9.01E-04 | 4.24E-03 | 2.58E-03 | gar-2/goa-1/gpb-2 | 3 |
| GO:0097305 | response to alcohol | 3/243 | 8/9281 | 9.01E-04 | 4.24E-03 | 2.58E-03 | npr-1/unc-79/egl-4 | 3 |
| GO:1903831 | signal transduction involved in cellular response to ammonium ion | 3/243 | 8/9281 | 9.01E-04 | 4.24E-03 | 2.58E-03 | gar-2/goa-1/gpb-2 | 3 |
| GO:1905144 | response to acetylcholine | 3/243 | 8/9281 | 9.01E-04 | 4.24E-03 | 2.58E-03 | gar-2/goa-1/gpb-2 | 3 |
| GO:1905145 | cellular response to acetylcholine | 3/243 | 8/9281 | 9.01E-04 | 4.24E-03 | 2.58E-03 | gar-2/goa-1/gpb-2 | 3 |
| GO:0001558 | regulation of cell growth | 7/243 | 60/9281 | 9.16E-04 | 4.30E-03 | 2.62E-03 | sma-2/unc-40/unc-69/unc-76/unc-104/rig-6/egl-4 | 7 |
| GO:0034762 | regulation of transmembrane transport | 6/243 | 44/9281 | 9.28E-04 | 4.33E-03 | 2.63E-03 | egl-2/egl-19/egl-36/unc-2/unc-103/egl-4 | 6 |
| GO:0034765 | regulation of ion transmembrane transport | 6/243 | 44/9281 | 9.28E-04 | 4.33E-03 | 2.63E-03 | egl-2/egl-19/egl-36/unc-2/unc-103/egl-4 | 6 |
| GO:1902667 | regulation of axon guidance | 6/243 | 44/9281 | 9.28E-04 | 4.33E-03 | 2.63E-03 | egl-20/rab-3/unc-31/unc-40/unc-104/rig-6 | 6 |
| GO:2000145 | regulation of cell motility | 9/243 | 97/9281 | 9.60E-04 | 4.46E-03 | 2.71E-03 | ced-1/egl-17/egl-20/egl-27/egl-43/lon-2/snf-10/unc-40/rig-6 | 9 |
| GO:0050772 | positive regulation of axonogenesis | 5/243 | 30/9281 | 9.85E-04 | 4.57E-03 | 2.78E-03 | egl-20/rab-3/unc-31/unc-104/rig-6 | 5 |
| GO:0015893 | drug transport | 8/243 | 79/9281 | 1.03E-03 | 4.78E-03 | 2.91E-03 | dat-1/egl-8/egl-21/egl-30/snf-6/unc-17/unc-26/unc-29 | 8 |
| GO:0043270 | positive regulation of ion transport | 4/243 | 18/9281 | 1.05E-03 | 4.85E-03 | 2.95E-03 | egl-8/egl-21/egl-30/unc-29 | 4 |
| GO:2000027 | regulation of animal organ morphogenesis | 5/243 | 31/9281 | 1.15E-03 | 5.30E-03 | 3.22E-03 | lin-39/lon-2/sma-2/sma-3/unc-40 | 5 |
| GO:0099504 | synaptic vesicle cycle | 6/243 | 46/9281 | 1.18E-03 | 5.42E-03 | 3.30E-03 | pkc-1/tom-1/unc-10/unc-18/unc-26/unc-31 | 6 |
| GO:0048878 | chemical homeostasis | 12/243 | 165/9281 | 1.30E-03 | 5.92E-03 | 3.60E-03 | acr-2/acr-3/ftn-2/spe-41/trp-1/trp-2/unc-32/unc-38/unc-63/unc-80/unc-108/ser-5 | 12 |
| GO:0044062 | regulation of excretion | 4/243 | 19/9281 | 1.30E-03 | 5.92E-03 | 3.60E-03 | egl-21/egl-30/gsa-1/rig-6 | 4 |
| GO:0051932 | synaptic transmission, GABAergic | 4/243 | 19/9281 | 1.30E-03 | 5.92E-03 | 3.60E-03 | nca-2/rab-3/unc-75/unc-77 | 4 |
| GO:0051952 | regulation of amine transport | 4/243 | 19/9281 | 1.30E-03 | 5.92E-03 | 3.60E-03 | egl-8/egl-21/egl-30/unc-29 | 4 |
| GO:2000292 | regulation of defecation | 4/243 | 19/9281 | 1.30E-03 | 5.92E-03 | 3.60E-03 | egl-21/egl-30/gsa-1/rig-6 | 4 |
| GO:0060359 | response to ammonium ion | 3/243 | 9/9281 | 1.32E-03 | 6.00E-03 | 3.65E-03 | gar-2/goa-1/gpb-2 | 3 |
| GO:0019953 | sexual reproduction | 18/243 | 314/9281 | 1.49E-03 | 6.74E-03 | 4.10E-03 | bas-1/cat-2/dnc-1/flp-10/flp-12/flp-20/gld-1/gsa-1/lov-1/mec-4/rab-3/ser-1/spe-41/trp-1/unc-10/unc-103/unc-116/rig-6 | 18 |
| GO:0033564 | anterior/posterior axon guidance | 4/243 | 20/9281 | 1.60E-03 | 7.18E-03 | 4.36E-03 | egl-20/rab-3/unc-31/unc-104 | 4 |
| GO:0061067 | negative regulation of dauer larval development | 4/243 | 20/9281 | 1.60E-03 | 7.18E-03 | 4.36E-03 | daf-7/ins-3/ins-4/egl-4 | 4 |
| GO:0010470 | regulation of gastrulation | 5/243 | 34/9281 | 1.77E-03 | 7.92E-03 | 4.82E-03 | lin-39/lon-2/sma-2/sma-3/unc-40 | 5 |
| GO:0006570 | tyrosine metabolic process | 3/243 | 10/9281 | 1.86E-03 | 8.21E-03 | 4.99E-03 | bas-1/cat-2/cat-4 | 3 |
| GO:0007419 | ventral cord development | 3/243 | 10/9281 | 1.86E-03 | 8.21E-03 | 4.99E-03 | rab-3/unc-31/unc-104 | 3 |
| GO:0030509 | BMP signaling pathway | 3/243 | 10/9281 | 1.86E-03 | 8.21E-03 | 4.99E-03 | daf-5/sma-3/unc-40 | 3 |
| GO:0071772 | response to BMP | 3/243 | 10/9281 | 1.86E-03 | 8.21E-03 | 4.99E-03 | daf-5/sma-3/unc-40 | 3 |
| GO:0071773 | cellular response to BMP stimulus | 3/243 | 10/9281 | 1.86E-03 | 8.21E-03 | 4.99E-03 | daf-5/sma-3/unc-40 | 3 |
| GO:1901048 | transforming growth factor beta receptor signaling pathway involved in regulation of multicellular organism growth | 3/243 | 10/9281 | 1.86E-03 | 8.21E-03 | 4.99E-03 | lon-2/sma-3/unc-40 | 3 |
| GO:0098542 | defense response to other organism | 13/243 | 196/9281 | 1.90E-03 | 8.40E-03 | 5.11E-03 | ced-1/daf-3/daf-7/egl-1/egl-30/ftn-2/goa-1/gpa-2/gpa-3/gpa-12/pmk-1/sma-3/unc-32 | 13 |
| GO:0060560 | developmental growth involved in morphogenesis | 7/243 | 68/9281 | 1.93E-03 | 8.48E-03 | 5.15E-03 | syg-1/unc-40/unc-69/unc-76/unc-104/syg-2/rig-6 | 7 |
| GO:0017157 | regulation of exocytosis | 4/243 | 21/9281 | 1.93E-03 | 8.48E-03 | 5.16E-03 | rab-3/sng-1/tom-1/unc-10 | 4 |
| GO:0007191 | adenylate cyclase-activating dopamine receptor signaling pathway | 2/243 | 3/9281 | 2.01E-03 | 8.54E-03 | 5.19E-03 | dop-1/gsa-1 | 2 |
| GO:0031998 | regulation of fatty acid beta-oxidation | 2/243 | 3/9281 | 2.01E-03 | 8.54E-03 | 5.19E-03 | mod-1/ser-6 | 2 |
| GO:0032000 | positive regulation of fatty acid beta-oxidation | 2/243 | 3/9281 | 2.01E-03 | 8.54E-03 | 5.19E-03 | mod-1/ser-6 | 2 |
| GO:0042427 | serotonin biosynthetic process | 2/243 | 3/9281 | 2.01E-03 | 8.54E-03 | 5.19E-03 | bas-1/tph-1 | 2 |
| GO:0042435 | indole-containing compound biosynthetic process | 2/243 | 3/9281 | 2.01E-03 | 8.54E-03 | 5.19E-03 | bas-1/tph-1 | 2 |
| GO:0050994 | regulation of lipid catabolic process | 2/243 | 3/9281 | 2.01E-03 | 8.54E-03 | 5.19E-03 | mod-1/ser-6 | 2 |
| GO:0050996 | positive regulation of lipid catabolic process | 2/243 | 3/9281 | 2.01E-03 | 8.54E-03 | 5.19E-03 | mod-1/ser-6 | 2 |
| GO:0070305 | response to cGMP | 2/243 | 3/9281 | 2.01E-03 | 8.54E-03 | 5.19E-03 | daf-7/gpa-3 | 2 |
| GO:0071321 | cellular response to cGMP | 2/243 | 3/9281 | 2.01E-03 | 8.54E-03 | 5.19E-03 | daf-7/gpa-3 | 2 |
| GO:0071444 | cellular response to pheromone | 2/243 | 3/9281 | 2.01E-03 | 8.54E-03 | 5.19E-03 | daf-3/daf-7 | 2 |
| GO:0098810 | neurotransmitter reuptake | 2/243 | 3/9281 | 2.01E-03 | 8.54E-03 | 5.19E-03 | dat-1/mod-5 | 2 |
| GO:1901162 | primary amino compound biosynthetic process | 2/243 | 3/9281 | 2.01E-03 | 8.54E-03 | 5.19E-03 | bas-1/tph-1 | 2 |
| GO:1904068 | G protein-coupled receptor signaling pathway involved in social behavior | 2/243 | 3/9281 | 2.01E-03 | 8.54E-03 | 5.19E-03 | flp-21/npr-1 | 2 |
| GO:0098659 | inorganic cation import across plasma membrane | 5/243 | 35/9281 | 2.02E-03 | 8.54E-03 | 5.19E-03 | acr-2/acr-3/ftn-2/unc-38/unc-63 | 5 |
| GO:0098739 | import across plasma membrane | 5/243 | 35/9281 | 2.02E-03 | 8.54E-03 | 5.19E-03 | acr-2/acr-3/ftn-2/unc-38/unc-63 | 5 |
| GO:0099587 | inorganic ion import across plasma membrane | 5/243 | 35/9281 | 2.02E-03 | 8.54E-03 | 5.19E-03 | acr-2/acr-3/ftn-2/unc-38/unc-63 | 5 |
| GO:0009607 | response to biotic stimulus | 13/243 | 198/9281 | 2.08E-03 | 8.77E-03 | 5.33E-03 | ced-1/daf-3/daf-7/egl-1/egl-30/ftn-2/goa-1/gpa-2/gpa-3/gpa-12/pmk-1/sma-3/unc-32 | 13 |
| GO:0043207 | response to external biotic stimulus | 13/243 | 198/9281 | 2.08E-03 | 8.77E-03 | 5.33E-03 | ced-1/daf-3/daf-7/egl-1/egl-30/ftn-2/goa-1/gpa-2/gpa-3/gpa-12/pmk-1/sma-3/unc-32 | 13 |
| GO:0051707 | response to other organism | 13/243 | 198/9281 | 2.08E-03 | 8.77E-03 | 5.33E-03 | ced-1/daf-3/daf-7/egl-1/egl-30/ftn-2/goa-1/gpa-2/gpa-3/gpa-12/pmk-1/sma-3/unc-32 | 13 |
| GO:1901615 | organic hydroxy compound metabolic process | 8/243 | 89/9281 | 2.24E-03 | 9.39E-03 | 5.71E-03 | bas-1/cat-2/cat-4/egl-8/tbh-1/tdc-1/tph-1/unc-26 | 8 |
| GO:0002682 | regulation of immune system process | 4/243 | 22/9281 | 2.31E-03 | 9.64E-03 | 5.86E-03 | egl-30/gpa-12/pmk-1/octr-1 | 4 |
| GO:0050776 | regulation of immune response | 4/243 | 22/9281 | 2.31E-03 | 9.64E-03 | 5.86E-03 | egl-30/gpa-12/pmk-1/octr-1 | 4 |
| GO:1902074 | response to salt | 4/243 | 22/9281 | 2.31E-03 | 9.64E-03 | 5.86E-03 | egl-30/pkc-1/tom-1/unc-31 | 4 |
| GO:0051641 | cellular localization | 36/243 | 848/9281 | 2.43E-03 | 1.01E-02 | 6.16E-03 | acr-2/acr-3/ced-1/dat-1/dnc-1/egl-8/egl-20/egl-21/egl-30/ftn-2/gpa-16/hcf-1/jnk-1/mod-5/pkc-1/pmk-1/rab-3/ric-19/sng-1/tom-1/unc-4/unc-10/unc-14/unc-16/unc-17/unc-18/unc-26/unc-29/unc-31/unc-38/unc-63/unc-89/unc-101/unc-104/unc-108/unc-116 | 36 |
| GO:0044703 | multi-organism reproductive process | 18/243 | 329/9281 | 2.49E-03 | 1.03E-02 | 6.29E-03 | bas-1/cat-2/dnc-1/flp-10/flp-12/flp-20/gld-1/gsa-1/lov-1/mec-4/rab-3/ser-1/spe-41/trp-1/unc-10/unc-103/unc-116/rig-6 | 18 |
| GO:0010940 | positive regulation of necrotic cell death | 3/243 | 11/9281 | 2.50E-03 | 1.03E-02 | 6.29E-03 | unc-26/unc-104/unc-116 | 3 |
| GO:0040015 | negative regulation of multicellular organism growth | 3/243 | 11/9281 | 2.50E-03 | 1.03E-02 | 6.29E-03 | lon-2/rig-6/egl-4 | 3 |
| GO:0009968 | negative regulation of signal transduction | 10/243 | 133/9281 | 2.54E-03 | 1.05E-02 | 6.36E-03 | daf-5/egl-10/gsa-1/ins-18/lon-2/osm-9/rgs-6/unc-2/unc-37/egl-4 | 10 |
| GO:0010770 | positive regulation of cell morphogenesis involved in differentiation | 5/243 | 37/9281 | 2.60E-03 | 1.07E-02 | 6.49E-03 | egl-20/rab-3/unc-31/unc-104/rig-6 | 5 |
| GO:0048588 | developmental cell growth | 7/243 | 72/9281 | 2.68E-03 | 1.10E-02 | 6.69E-03 | syg-1/unc-40/unc-69/unc-76/unc-104/syg-2/rig-6 | 7 |
| GO:0001710 | mesodermal cell fate commitment | 5/243 | 38/9281 | 2.93E-03 | 1.20E-02 | 7.27E-03 | lin-39/lon-2/sma-2/sma-3/unc-40 | 5 |
| GO:0007501 | mesodermal cell fate specification | 5/243 | 38/9281 | 2.93E-03 | 1.20E-02 | 7.27E-03 | lin-39/lon-2/sma-2/sma-3/unc-40 | 5 |
| GO:0097164 | ammonium ion metabolic process | 6/243 | 55/9281 | 3.00E-03 | 1.22E-02 | 7.45E-03 | bas-1/cat-2/cat-4/tbh-1/tdc-1/tph-1 | 6 |
| GO:0042742 | defense response to bacterium | 11/243 | 159/9281 | 3.02E-03 | 1.23E-02 | 7.48E-03 | ced-1/daf-3/daf-7/egl-1/egl-30/ftn-2/goa-1/gpa-2/gpa-3/pmk-1/unc-32 | 11 |
| GO:0030334 | regulation of cell migration | 8/243 | 94/9281 | 3.15E-03 | 1.28E-02 | 7.79E-03 | ced-1/egl-17/egl-20/egl-27/egl-43/lon-2/unc-40/rig-6 | 8 |
| GO:0009617 | response to bacterium | 11/243 | 160/9281 | 3.17E-03 | 1.29E-02 | 7.82E-03 | ced-1/daf-3/daf-7/egl-1/egl-30/ftn-2/goa-1/gpa-2/gpa-3/pmk-1/unc-32 | 11 |
| GO:0048679 | regulation of axon regeneration | 4/243 | 24/9281 | 3.22E-03 | 1.30E-02 | 7.90E-03 | egl-8/egl-30/goa-1/jnk-1 | 4 |
| GO:0050768 | negative regulation of neurogenesis | 4/243 | 24/9281 | 3.22E-03 | 1.30E-02 | 7.90E-03 | egl-20/goa-1/jnk-1/unc-40 | 4 |
| GO:0070570 | regulation of neuron projection regeneration | 4/243 | 24/9281 | 3.22E-03 | 1.30E-02 | 7.90E-03 | egl-8/egl-30/goa-1/jnk-1 | 4 |
| GO:0010939 | regulation of necrotic cell death | 3/243 | 12/9281 | 3.27E-03 | 1.31E-02 | 7.98E-03 | unc-26/unc-104/unc-116 | 3 |
| GO:1905909 | regulation of dauer entry | 3/243 | 12/9281 | 3.27E-03 | 1.31E-02 | 7.98E-03 | ins-3/ins-4/ins-18 | 3 |
| GO:0017156 | calcium ion regulated exocytosis | 5/243 | 39/9281 | 3.29E-03 | 1.31E-02 | 7.98E-03 | pkc-1/tom-1/unc-10/unc-18/unc-31 | 5 |
| GO:0022611 | dormancy process | 5/243 | 39/9281 | 3.29E-03 | 1.31E-02 | 7.98E-03 | daf-7/hcf-1/ins-3/ins-4/ins-18 | 5 |
| GO:0080135 | regulation of cellular response to stress | 7/243 | 75/9281 | 3.38E-03 | 1.35E-02 | 8.19E-03 | egl-8/egl-30/goa-1/jnk-1/nlp-2/octr-1/unc-16 | 7 |
| GO:0009072 | aromatic amino acid family metabolic process | 4/243 | 25/9281 | 3.76E-03 | 1.50E-02 | 9.09E-03 | bas-1/cat-2/cat-4/tph-1 | 4 |
| GO:0006837 | serotonin transport | 2/243 | 4/9281 | 3.96E-03 | 1.55E-02 | 9.43E-03 | mod-5/unc-31 | 2 |
| GO:0015812 | gamma-aminobutyric acid transport | 2/243 | 4/9281 | 3.96E-03 | 1.55E-02 | 9.43E-03 | snf-11/unc-26 | 2 |
| GO:0042428 | serotonin metabolic process | 2/243 | 4/9281 | 3.96E-03 | 1.55E-02 | 9.43E-03 | bas-1/tph-1 | 2 |
| GO:0042439 | ethanolamine-containing compound metabolic process | 2/243 | 4/9281 | 3.96E-03 | 1.55E-02 | 9.43E-03 | tbh-1/tdc-1 | 2 |
| GO:0046958 | nonassociative learning | 2/243 | 4/9281 | 3.96E-03 | 1.55E-02 | 9.43E-03 | cat-2/dop-1 | 2 |
| GO:0046959 | habituation | 2/243 | 4/9281 | 3.96E-03 | 1.55E-02 | 9.43E-03 | cat-2/dop-1 | 2 |
| GO:2000300 | regulation of synaptic vesicle exocytosis | 2/243 | 4/9281 | 3.96E-03 | 1.55E-02 | 9.43E-03 | tom-1/unc-10 | 2 |
| GO:0006970 | response to osmotic stress | 5/243 | 41/9281 | 4.10E-03 | 1.60E-02 | 9.72E-03 | flp-1/jnk-1/odr-3/osm-9/pmk-1 | 5 |
| GO:0071417 | cellular response to organonitrogen compound | 5/243 | 41/9281 | 4.10E-03 | 1.60E-02 | 9.72E-03 | daf-7/dat-1/gpa-3/ins-18/egl-4 | 5 |
| GO:0007405 | neuroblast proliferation | 3/243 | 13/9281 | 4.17E-03 | 1.60E-02 | 9.72E-03 | egl-13/egl-44/egl-46 | 3 |
| GO:0007602 | phototransduction | 3/243 | 13/9281 | 4.17E-03 | 1.60E-02 | 9.72E-03 | gpa-3/lite-1/npr-3 | 3 |
| GO:0009583 | detection of light stimulus | 3/243 | 13/9281 | 4.17E-03 | 1.60E-02 | 9.72E-03 | gpa-3/lite-1/npr-3 | 3 |
| GO:0015844 | monoamine transport | 3/243 | 13/9281 | 4.17E-03 | 1.60E-02 | 9.72E-03 | dat-1/mod-5/unc-31 | 3 |
| GO:0036445 | neuronal stem cell division | 3/243 | 13/9281 | 4.17E-03 | 1.60E-02 | 9.72E-03 | egl-13/egl-44/egl-46 | 3 |
| GO:0048103 | somatic stem cell division | 3/243 | 13/9281 | 4.17E-03 | 1.60E-02 | 9.72E-03 | egl-13/egl-44/egl-46 | 3 |
| GO:0048663 | neuron fate commitment | 3/243 | 13/9281 | 4.17E-03 | 1.60E-02 | 9.72E-03 | egl-13/unc-42/unc-86 | 3 |
| GO:0055057 | neuroblast division | 3/243 | 13/9281 | 4.17E-03 | 1.60E-02 | 9.72E-03 | egl-13/egl-44/egl-46 | 3 |
| GO:0061351 | neural precursor cell proliferation | 3/243 | 13/9281 | 4.17E-03 | 1.60E-02 | 9.72E-03 | egl-13/egl-44/egl-46 | 3 |
| GO:0001704 | formation of primary germ layer | 6/243 | 59/9281 | 4.28E-03 | 1.64E-02 | 9.95E-03 | lin-39/lon-2/sem-4/sma-2/sma-3/unc-40 | 6 |
| GO:1901698 | response to nitrogen compound | 8/243 | 99/9281 | 4.34E-03 | 1.66E-02 | 1.01E-02 | daf-7/dat-1/gar-2/goa-1/gpa-3/gpb-2/ins-18/egl-4 | 8 |
| GO:0048640 | negative regulation of developmental growth | 4/243 | 26/9281 | 4.35E-03 | 1.66E-02 | 1.01E-02 | lon-2/unc-40/rig-6/egl-4 | 4 |
| GO:0007389 | pattern specification process | 9/243 | 121/9281 | 4.42E-03 | 1.68E-02 | 1.02E-02 | ced-1/egl-5/egl-19/egl-20/gpa-16/lin-39/unc-2/unc-40/rig-6 | 9 |
| GO:0030516 | regulation of axon extension | 5/243 | 42/9281 | 4.56E-03 | 1.73E-02 | 1.05E-02 | unc-40/unc-69/unc-76/unc-104/rig-6 | 5 |
| GO:0019725 | cellular homeostasis | 12/243 | 192/9281 | 4.56E-03 | 1.73E-02 | 1.05E-02 | acr-2/acr-3/egl-15/ftn-2/spe-41/trp-1/trp-2/unc-32/unc-38/unc-63/unc-108/ser-5 | 12 |
| GO:0051961 | negative regulation of nervous system development | 4/243 | 27/9281 | 5.01E-03 | 1.89E-02 | 1.15E-02 | egl-20/goa-1/jnk-1/unc-40 | 4 |
| GO:0061387 | regulation of extent of cell growth | 5/243 | 43/9281 | 5.05E-03 | 1.90E-02 | 1.16E-02 | unc-40/unc-69/unc-76/unc-104/rig-6 | 5 |
| GO:1990834 | response to odorant | 3/243 | 14/9281 | 5.21E-03 | 1.96E-02 | 1.19E-02 | odr-3/dop-3/egl-4 | 3 |
| GO:0045596 | negative regulation of cell differentiation | 5/243 | 44/9281 | 5.58E-03 | 2.09E-02 | 1.27E-02 | daf-5/egl-20/goa-1/jnk-1/unc-40 | 5 |
| GO:0045926 | negative regulation of growth | 4/243 | 28/9281 | 5.73E-03 | 2.15E-02 | 1.30E-02 | lon-2/unc-40/rig-6/egl-4 | 4 |
| GO:0070265 | necrotic cell death | 3/243 | 15/9281 | 6.39E-03 | 2.35E-02 | 1.43E-02 | unc-26/unc-104/unc-116 | 3 |
| GO:1903356 | positive regulation of distal tip cell migration | 3/243 | 15/9281 | 6.39E-03 | 2.35E-02 | 1.43E-02 | ced-1/egl-20/rig-6 | 3 |
| GO:0014074 | response to purine-containing compound | 2/243 | 5/9281 | 6.48E-03 | 2.35E-02 | 1.43E-02 | daf-7/gpa-3 | 2 |

| | | | | | | | | |
|---|---|---|---|---|---|---|---|---|
| GO:0030510 | regulation of BMP signaling pathway | 2/243 | 5/9281 | 6.48E-03 | 2.35E-02 | 1.43E-02 | daf-5/unc-40 | 2 |
| GO:0030951 | establishment or maintenance of microtubule cytoskeleton polarity | 2/243 | 5/9281 | 6.48E-03 | 2.35E-02 | 1.43E-02 | unc-116/zyg-9 | 2 |
| GO:0033363 | secretory granule organization | 2/243 | 5/9281 | 6.48E-03 | 2.35E-02 | 1.43E-02 | ric-19/unc-108 | 2 |
| GO:0035176 | social behavior | 2/243 | 5/9281 | 6.48E-03 | 2.35E-02 | 1.43E-02 | flp-21/npr-1 | 2 |
| GO:0045989 | positive regulation of striated muscle contraction | 2/243 | 5/9281 | 6.48E-03 | 2.35E-02 | 1.43E-02 | egl-19/unc-89 | 2 |
| GO:0046683 | response to organophosphorus | 2/243 | 5/9281 | 6.48E-03 | 2.35E-02 | 1.43E-02 | daf-7/gpa-3 | 2 |
| GO:0051703 | intraspecies interaction between organisms | 2/243 | 5/9281 | 6.48E-03 | 2.35E-02 | 1.43E-02 | flp-21/npr-1 | 2 |
| GO:0051705 | multi-organism behavior | 2/243 | 5/9281 | 6.48E-03 | 2.35E-02 | 1.43E-02 | flp-21/npr-1 | 2 |
| GO:0061792 | secretory granule maturation | 2/243 | 5/9281 | 6.48E-03 | 2.35E-02 | 1.43E-02 | ric-19/unc-108 | 2 |
| GO:1901160 | primary amino compound metabolic process | 2/243 | 5/9281 | 6.48E-03 | 2.35E-02 | 1.43E-02 | bas-1/tph-1 | 2 |
| GO:1990502 | dense core granule maturation | 2/243 | 5/9281 | 6.48E-03 | 2.35E-02 | 1.43E-02 | ric-19/unc-108 | 2 |
| GO:0043053 | dauer entry | 4/243 | 29/9281 | 6.51E-03 | 2.35E-02 | 1.43E-02 | daf-7/ins-3/ins-4/ins-18 | 4 |
| GO:0055115 | entry into diapause | 4/243 | 29/9281 | 6.51E-03 | 2.35E-02 | 1.43E-02 | daf-7/ins-3/ins-4/ins-18 | 4 |
| GO:0140029 | exocytic process | 4/243 | 29/9281 | 6.51E-03 | 2.35E-02 | 1.43E-02 | rab-3/tom-1/unc-10/unc-18 | 4 |
| GO:1903034 | regulation of response to wounding | 4/243 | 29/9281 | 6.51E-03 | 2.35E-02 | 1.43E-02 | egl-8/egl-30/goa-1/jnk-1 | 4 |
| GO:0008361 | regulation of cell size | 6/243 | 65/9281 | 6.89E-03 | 2.49E-02 | 1.51E-02 | sma-3/unc-40/unc-69/unc-76/unc-104/rig-6 | 6 |
| GO:0007163 | establishment or maintenance of cell polarity | 8/243 | 108/9281 | 7.30E-03 | 2.63E-02 | 1.60E-02 | ced-1/dnc-1/egl-20/egl-27/gpa-16/gsa-1/unc-116/zyg-9 | 8 |
| GO:0009987 | cellular process | 207/243 | 7322/9281 | 7.46E-03 | 2.68E-02 | 1.63E-02 | acr-2/acr-3/acr-6/acr-7/acr-9/acr-10/acr-11/acr-14/acr-15/acr-18/acr-19/acr-21/acr-23/bas-1/cat-2/cat-4/ced-1/daf-3/daf-5/daf-7/dat-1/dnc-1/dop-1/dop-2/eat-16/egl-1/egl-2/egl-5/egl-6/egl-8/egl-9/egl-10/egl-13/egl-15/egl-17/egl-18/egl-20/egl-21/egl-23/egl-27/egl-30/egl-36/egl-43/egl-44/egl-46/egl-47/flp-1/flp-3/flp-6/f | 207 |
| GO:0035556 | intracellular signal transduction | 23/243 | 511/9281 | 7.71E-03 | 2.77E-02 | 1.68E-02 | daf-7/dop-1/eat-16/egl-8/egl-10/egl-44/flp-7/gpa-12/gsa-1/jnk-1/pkc-1/pmk-1/rab-3/ser-1/ser-2/unc-16/unc-43/unc-89/unc-108/ser-5/tyra-2/dop-3/egl-4 | 23 |
| GO:0070482 | response to oxygen levels | 4/243 | 31/9281 | 8.28E-03 | 2.97E-02 | 1.80E-02 | egl-9/npr-1/unc-32/egl-4 | 4 |
| GO:0048646 | anatomical structure formation involved in morphogenesis | 9/243 | 134/9281 | 8.52E-03 | 3.05E-02 | 1.85E-02 | lin-39/lon-2/sem-4/sma-2/sma-3/unc-40/unc-60/unc-89/unc-98 | 9 |
| GO:0051050 | positive regulation of transport | 8/243 | 112/9281 | 9.03E-03 | 3.22E-02 | 1.96E-02 | ced-1/egl-8/egl-21/egl-30/unc-18/unc-26/unc-29/unc-104 | 8 |
| GO:0051965 | positive regulation of synapse assembly | 3/243 | 17/9281 | 9.19E-03 | 3.27E-02 | 1.99E-02 | egl-21/unc-40/unc-104 | 3 |
| GO:0072089 | stem cell proliferation | 3/243 | 17/9281 | 9.19E-03 | 3.27E-02 | 1.99E-02 | egl-13/egl-44/egl-46 | 3 |
| GO:0010628 | positive regulation of gene expression | 19/243 | 402/9281 | 9.20E-03 | 3.27E-02 | 1.99E-02 | daf-7/egl-5/egl-13/egl-44/gld-1/gpa-2/gpa-3/gpa-12/lin-39/pmk-1/sma-3/unc-3/unc-30/unc-42/unc-43/unc-55/unc-86/unc-89/dop-3 | 19 |
| GO:0051963 | regulation of synapse assembly | 4/243 | 32/9281 | 9.27E-03 | 3.28E-02 | 2.00E-02 | egl-21/ins-4/unc-40/unc-104 | 4 |
| GO:0048870 | cell motility | 11/243 | 185/9281 | 9.32E-03 | 3.29E-02 | 2.00E-02 | ced-1/egl-15/egl-17/egl-20/egl-27/egl-43/egl-46/lon-2/snf-10/unc-40/rig-6 | 11 |
| GO:0051674 | localization of cell | 11/243 | 185/9281 | 9.32E-03 | 3.29E-02 | 2.00E-02 | ced-1/egl-15/egl-17/egl-20/egl-27/egl-43/egl-46/lon-2/snf-10/unc-40/rig-6 | 11 |
| GO:0007096 | regulation of exit from mitosis | 2/243 | 6/9281 | 9.55E-03 | 3.30E-02 | 2.01E-02 | egl-44/egl-46 | 2 |
| GO:0009309 | amine biosynthetic process | 2/243 | 6/9281 | 9.55E-03 | 3.30E-02 | 2.01E-02 | tbh-1/tdc-1 | 2 |
| GO:0010458 | exit from mitosis | 2/243 | 6/9281 | 9.55E-03 | 3.30E-02 | 2.01E-02 | egl-44/egl-46 | 2 |
| GO:0016082 | synaptic vesicle priming | 2/243 | 6/9281 | 9.55E-03 | 3.30E-02 | 2.01E-02 | tom-1/unc-10 | 2 |
| GO:0035265 | organ growth | 2/243 | 6/9281 | 9.55E-03 | 3.30E-02 | 2.01E-02 | sma-3/egl-4 | 2 |
| GO:0042401 | cellular biogenic amine biosynthetic process | 2/243 | 6/9281 | 9.55E-03 | 3.30E-02 | 2.01E-02 | tbh-1/tdc-1 | 2 |
| GO:0046320 | regulation of fatty acid oxidation | 2/243 | 6/9281 | 9.55E-03 | 3.30E-02 | 2.01E-02 | mod-1/ser-6 | 2 |
| GO:0046321 | positive regulation of fatty acid oxidation | 2/243 | 6/9281 | 9.55E-03 | 3.30E-02 | 2.01E-02 | mod-1/ser-6 | 2 |
| GO:0046620 | regulation of organ growth | 2/243 | 6/9281 | 9.55E-03 | 3.30E-02 | 2.01E-02 | sma-3/egl-4 | 2 |
| GO:0048668 | collateral sprouting | 2/243 | 6/9281 | 9.55E-03 | 3.30E-02 | 2.01E-02 | syg-1/syg-2 | 2 |
| GO:0098693 | regulation of synaptic vesicle cycle | 2/243 | 6/9281 | 9.55E-03 | 3.30E-02 | 2.01E-02 | tom-1/unc-10 | 2 |
| GO:0098657 | import into cell | 11/243 | 186/9281 | 9.68E-03 | 3.34E-02 | 2.03E-02 | acr-2/acr-3/ced-1/dat-1/ftn-2/mod-5/snf-6/unc-26/unc-38/unc-63/unc-108 | 11 |
| GO:0010721 | negative regulation of cell development | 4/243 | 33/9281 | 1.03E-02 | 3.56E-02 | 2.16E-02 | egl-20/goa-1/jnk-1/unc-40 | 4 |
| GO:0046165 | alcohol biosynthetic process | 4/243 | 33/9281 | 1.03E-02 | 3.56E-02 | 2.16E-02 | cat-4/egl-8/tbh-1/tdc-1 | 4 |
| GO:0002684 | positive regulation of immune system process | 3/243 | 18/9281 | 1.08E-02 | 3.68E-02 | 2.24E-02 | egl-30/gpa-12/pmk-1 | 3 |
| GO:0008582 | regulation of synaptic growth at neuromuscular junction | 3/243 | 18/9281 | 1.08E-02 | 3.68E-02 | 2.24E-02 | egl-21/ins-4/unc-104 | 3 |
| GO:0032102 | negative regulation of response to external stimulus | 3/243 | 18/9281 | 1.08E-02 | 3.68E-02 | 2.24E-02 | goa-1/jnk-1/unc-40 | 3 |
| GO:0050778 | positive regulation of immune response | 3/243 | 18/9281 | 1.08E-02 | 3.68E-02 | 2.24E-02 | egl-30/gpa-12/pmk-1 | 3 |
| GO:0051124 | synaptic growth at neuromuscular junction | 3/243 | 18/9281 | 1.08E-02 | 3.68E-02 | 2.24E-02 | egl-21/ins-4/unc-104 | 3 |
| GO:1904396 | regulation of neuromuscular junction development | 3/243 | 18/9281 | 1.08E-02 | 3.68E-02 | 2.24E-02 | egl-21/ins-4/unc-104 | 3 |
| GO:0051294 | establishment of spindle orientation | 5/243 | 52/9281 | 1.13E-02 | 3.83E-02 | 2.33E-02 | ced-1/dnc-1/egl-20/gpa-16/unc-116 | 5 |
| GO:0008306 | associative learning | 4/243 | 34/9281 | 1.15E-02 | 3.89E-02 | 2.36E-02 | gpa-2/osm-9/pkc-1/egl-4 | 4 |
| GO:2000147 | positive regulation of cell motility | 4/243 | 34/9281 | 1.15E-02 | 3.89E-02 | 2.36E-02 | ced-1/egl-20/snf-10/rig-6 | 4 |
| GO:0090066 | regulation of anatomical structure size | 8/243 | 117/9281 | 1.16E-02 | 3.92E-02 | 2.38E-02 | sma-3/unc-40/unc-60/unc-69/unc-76/unc-104/ser-5/rig-6 | 8 |
| GO:0008219 | cell death | 11/243 | 191/9281 | 1.17E-02 | 3.94E-02 | 2.39E-02 | ced-1/daf-7/dat-1/egl-1/jnk-1/pmk-1/unc-3/unc-26/unc-32/unc-104/unc-116 | 11 |
| GO:0002831 | regulation of response to biotic stimulus | 3/243 | 19/9281 | 1.26E-02 | 4.23E-02 | 2.57E-02 | egl-30/gpa-12/pmk-1 | 3 |
| GO:0006576 | cellular biogenic amine metabolic process | 3/243 | 19/9281 | 1.26E-02 | 4.23E-02 | 2.57E-02 | tbh-1/tdc-1/tph-1 | 3 |
| GO:0048511 | rhythmic process | 3/243 | 19/9281 | 1.26E-02 | 4.23E-02 | 2.57E-02 | egl-10/flr-1/rig-6 | 3 |
| GO:0048729 | tissue morphogenesis | 7/243 | 96/9281 | 1.28E-02 | 4.29E-02 | 2.61E-02 | lin-39/lon-2/sem-4/sma-2/sma-3/unc-40/rig-6 | 7 |
| GO:0045471 | response to ethanol | 2/243 | 7/9281 | 1.31E-02 | 4.36E-02 | 2.65E-02 | npr-1/unc-79 | 2 |
| GO:0048681 | negative regulation of axon regeneration | 2/243 | 7/9281 | 1.31E-02 | 4.36E-02 | 2.65E-02 | goa-1/jnk-1 | 2 |
| GO:0070571 | negative regulation of neuron projection regeneration | 2/243 | 7/9281 | 1.31E-02 | 4.36E-02 | 2.65E-02 | goa-1/jnk-1 | 2 |
| GO:1902075 | cellular response to salt | 2/243 | 7/9281 | 1.31E-02 | 4.36E-02 | 2.65E-02 | tom-1/unc-31 | 2 |
| GO:1905803 | negative regulation of cellular response to manganese ion | 2/243 | 7/9281 | 1.31E-02 | 4.36E-02 | 2.65E-02 | dat-1/jnk-1 | 2 |
| GO:0030010 | establishment of cell polarity | 6/243 | 75/9281 | 1.35E-02 | 4.49E-02 | 2.73E-02 | ced-1/dnc-1/egl-20/gpa-16/gsa-1/unc-116 | 6 |
| GO:0045944 | positive regulation of transcription by RNA polymerase II | 14/243 | 278/9281 | 1.45E-02 | 4.78E-02 | 2.91E-02 | daf-7/egl-5/egl-13/egl-44/gpa-12/lin-39/pmk-1/sma-3/unc-3/unc-30/unc-42/unc-55/unc-86/dop-3 | 14 |
| GO:0097628 | distal tip cell migration | 3/243 | 20/9281 | 1.45E-02 | 4.78E-02 | 2.91E-02 | ced-1/egl-20/rig-6 | 3 |
| GO:1901214 | regulation of neuron death | 3/243 | 20/9281 | 1.45E-02 | 4.78E-02 | 2.91E-02 | dat-1/jnk-1/pmk-1 | 3 |
| GO:1903354 | regulation of distal tip cell migration | 3/243 | 20/9281 | 1.45E-02 | 4.78E-02 | 2.91E-02 | ced-1/egl-20/rig-6 | 3 |
| GO:0050829 | defense response to Gram-negative bacterium | 7/243 | 99/9281 | 1.50E-02 | 4.92E-02 | 2.99E-02 | daf-3/daf-7/egl-1/goa-1/gpa-2/gpa-3/pmk-1 | 7 |
| GO:0051606 | detection of stimulus | 12/243 | 225/9281 | 1.52E-02 | 4.98E-02 | 3.03E-02 | daf-7/gpa-3/lite-1/lov-1/mec-4/mec-7/mec-10/mec-12/mec-14/mec-18/npr-3/tmc-1 | 12 |
| GO:0060795 | cell fate commitment involved in formation of primary germ layer | 5/243 | 56/9281 | 1.52E-02 | 4.99E-02 | 3.03E-02 | lin-39/lon-2/sma-2/sma-3/unc-40 | 5 |
| GO:0007600 | sensory perception | 18/243 | 396/9281 | 1.60E-02 | 5.22E-02 | 3.17E-02 | bas-1/egl-47/gpa-3/gpc-1/lite-1/mec-4/mec-7/mec-10/mec-12/mec-14/mec-18/odr-3/osm-9/pdl-1/pkc-1/unc-86/tmc-1/egl-4 | 18 |
| GO:0048513 | animal organ development | 18/243 | 396/9281 | 1.60E-02 | 5.22E-02 | 3.17E-02 | egl-13/egl-15/egl-20/gld-1/gon-2/lin-39/lon-2/rab-3/sma-2/sma-3/unc-31/unc-34/unc-40/unc-89/unc-98/unc-101/unc-104/unc-105 | 18 |
| GO:0001708 | cell fate specification | 10/243 | 174/9281 | 1.60E-02 | 5.22E-02 | 3.17E-02 | egl-13/egl-18/egl-27/lin-39/lon-2/sma-2/sma-3/unc-40/unc-42/unc-86 | 10 |
| GO:0016477 | cell migration | 10/243 | 174/9281 | 1.60E-02 | 5.22E-02 | 3.17E-02 | ced-1/egl-15/egl-17/egl-20/egl-27/egl-43/egl-46/lon-2/unc-40/rig-6 | 10 |
| GO:0014070 | response to organic cyclic compound | 5/243 | 57/9281 | 1.64E-02 | 5.30E-02 | 3.22E-02 | daf-7/dat-1/gpa-3/unc-55/unc-79 | 5 |
| GO:0048675 | axon extension | 5/243 | 57/9281 | 1.64E-02 | 5.30E-02 | 3.22E-02 | unc-40/unc-69/unc-76/unc-104/rig-6 | 5 |
| GO:0006904 | vesicle docking involved in exocytosis | 3/243 | 21/9281 | 1.66E-02 | 5.32E-02 | 3.23E-02 | rab-3/tom-1/unc-18 | 3 |
| GO:0044106 | cellular amine metabolic process | 3/243 | 21/9281 | 1.66E-02 | 5.32E-02 | 3.23E-02 | tbh-1/tdc-1/tph-1 | 3 |
| GO:0001508 | action potential | 2/243 | 8/9281 | 1.72E-02 | 5.32E-02 | 3.23E-02 | flp-3/flp-9 | 2 |
| GO:0002225 | positive regulation of antimicrobial peptide production | 2/243 | 8/9281 | 1.72E-02 | 5.32E-02 | 3.23E-02 | gpa-12/pmk-1 | 2 |
| GO:0002440 | production of molecular mediator of immune response | 2/243 | 8/9281 | 1.72E-02 | 5.32E-02 | 3.23E-02 | gpa-12/pmk-1 | 2 |
| GO:0002699 | positive regulation of immune effector process | 2/243 | 8/9281 | 1.72E-02 | 5.32E-02 | 3.23E-02 | gpa-12/pmk-1 | 2 |
| GO:0002700 | regulation of production of molecular mediator of immune response | 2/243 | 8/9281 | 1.72E-02 | 5.32E-02 | 3.23E-02 | gpa-12/pmk-1 | 2 |
| GO:0002702 | positive regulation of production of molecular mediator of immune response | 2/243 | 8/9281 | 1.72E-02 | 5.32E-02 | 3.23E-02 | gpa-12/pmk-1 | 2 |
| GO:0002759 | regulation of antimicrobial humoral response | 2/243 | 8/9281 | 1.72E-02 | 5.32E-02 | 3.23E-02 | gpa-12/pmk-1 | 2 |
| GO:0002760 | positive regulation of antimicrobial humoral response | 2/243 | 8/9281 | 1.72E-02 | 5.32E-02 | 3.23E-02 | gpa-12/pmk-1 | 2 |
| GO:0002775 | antimicrobial peptide production | 2/243 | 8/9281 | 1.72E-02 | 5.32E-02 | 3.23E-02 | gpa-12/pmk-1 | 2 |
| GO:0002784 | regulation of antimicrobial peptide production | 2/243 | 8/9281 | 1.72E-02 | 5.32E-02 | 3.23E-02 | gpa-12/pmk-1 | 2 |
| GO:0002920 | regulation of humoral immune response | 2/243 | 8/9281 | 1.72E-02 | 5.32E-02 | 3.23E-02 | gpa-12/pmk-1 | 2 |
| GO:0002922 | positive regulation of humoral immune response | 2/243 | 8/9281 | 1.72E-02 | 5.32E-02 | 3.23E-02 | gpa-12/pmk-1 | 2 |
| GO:0007624 | ultradian rhythm | 2/243 | 8/9281 | 1.72E-02 | 5.32E-02 | 3.23E-02 | flr-1/rig-6 | 2 |
| GO:0008543 | fibroblast growth factor receptor signaling pathway | 2/243 | 8/9281 | 1.72E-02 | 5.32E-02 | 3.23E-02 | egl-15/egl-17 | 2 |
| GO:0014016 | neuroblast differentiation | 2/243 | 8/9281 | 1.72E-02 | 5.32E-02 | 3.23E-02 | sem-4/unc-4 | 2 |
| GO:0014866 | skeletal myofibril assembly | 2/243 | 8/9281 | 1.72E-02 | 5.32E-02 | 3.23E-02 | unc-60/unc-89 | 2 |
| GO:0016081 | synaptic vesicle docking | 2/243 | 8/9281 | 1.72E-02 | 5.32E-02 | 3.23E-02 | tom-1/unc-18 | 2 |
| GO:0019730 | antimicrobial humoral response | 2/243 | 8/9281 | 1.72E-02 | 5.32E-02 | 3.23E-02 | gpa-12/pmk-1 | 2 |
| GO:0030952 | establishment or maintenance of cytoskeleton polarity | 2/243 | 8/9281 | 1.72E-02 | 5.32E-02 | 3.23E-02 | unc-116/zyg-9 | 2 |
| GO:0044344 | cellular response to fibroblast growth factor stimulus | 2/243 | 8/9281 | 1.72E-02 | 5.32E-02 | 3.23E-02 | egl-15/egl-17 | 2 |
| GO:0060756 | foraging behavior | 2/243 | 8/9281 | 1.72E-02 | 5.32E-02 | 3.23E-02 | nlp-12/npr-9 | 2 |
| GO:0061844 | antimicrobial humoral immune response mediated by antimicrobial peptide | 2/243 | 8/9281 | 1.72E-02 | 5.32E-02 | 3.23E-02 | gpa-12/pmk-1 | 2 |
| GO:0071287 | cellular response to manganese ion | 2/243 | 8/9281 | 1.72E-02 | 5.32E-02 | 3.23E-02 | dat-1/jnk-1 | 2 |
| GO:0071774 | response to fibroblast growth factor | 2/243 | 8/9281 | 1.72E-02 | 5.32E-02 | 3.23E-02 | egl-15/egl-17 | 2 |
| GO:1903035 | negative regulation of response to wounding | 2/243 | 8/9281 | 1.72E-02 | 5.32E-02 | 3.23E-02 | goa-1/jnk-1 | 2 |
| GO:1905802 | regulation of cellular response to manganese ion | 2/243 | 8/9281 | 1.72E-02 | 5.32E-02 | 3.23E-02 | dat-1/jnk-1 | 2 |
| GO:1905910 | negative regulation of dauer entry | 2/243 | 8/9281 | 1.72E-02 | 5.32E-02 | 3.23E-02 | ins-3/ins-4 | 2 |
| GO:0009308 | amine metabolic process | 3/243 | 22/9281 | 1.89E-02 | 5.80E-02 | 3.52E-02 | tbh-1/tdc-1/tph-1 | 3 |
| GO:0010977 | negative regulation of neuron projection development | 3/243 | 22/9281 | 1.89E-02 | 5.80E-02 | 3.52E-02 | goa-1/jnk-1/unc-40 | 3 |
| GO:0045665 | negative regulation of neuron differentiation | 3/243 | 22/9281 | 1.89E-02 | 5.80E-02 | 3.52E-02 | goa-1/jnk-1/unc-40 | 3 |
| GO:0051403 | stress-activated MAPK cascade | 3/243 | 22/9281 | 1.89E-02 | 5.80E-02 | 3.52E-02 | jnk-1/pmk-1/unc-16 | 3 |
| GO:0006357 | regulation of transcription by RNA polymerase II | 22/243 | 526/9281 | 1.98E-02 | 6.07E-02 | 3.69E-02 | daf-3/daf-5/daf-7/egl-5/egl-13/egl-18/egl-27/egl-44/egl-46/gpa-12/lin-39/pmk-1/sem-4/sma-3/unc-3/unc-4/unc-30/unc-40/unc-42/unc-55/unc-86/dop-3 | 22 |
| GO:0042659 | regulation of cell fate specification | 5/243 | 60/9281 | 2.00E-02 | 6.13E-02 | 3.72E-02 | lin-39/lon-2/sma-2/sma-3/unc-40 | 5 |
| GO:0007417 | central nervous system development | 3/243 | 23/9281 | 2.13E-02 | 6.50E-02 | 3.95E-02 | rab-3/unc-31/unc-104 | 3 |
| GO:0070997 | neuron death | 3/243 | 23/9281 | 2.13E-02 | 6.50E-02 | 3.95E-02 | dat-1/jnk-1/pmk-1 | 3 |
| GO:0009566 | fertilization | 4/243 | 41/9281 | 2.18E-02 | 6.51E-02 | 3.96E-02 | dnc-1/spe-41/trp-1/rig-6 | 4 |
| GO:0001845 | phagolysosome assembly | 2/243 | 9/9281 | 2.18E-02 | 6.51E-02 | 3.96E-02 | ced-1/unc-108 | 2 |
| GO:0002697 | regulation of immune effector process | 2/243 | 9/9281 | 2.18E-02 | 6.51E-02 | 3.96E-02 | gpa-12/pmk-1 | 2 |
| GO:0006959 | humoral immune response | 2/243 | 9/9281 | 2.18E-02 | 6.51E-02 | 3.96E-02 | gpa-12/pmk-1 | 2 |
| GO:0019236 | response to pheromone | 2/243 | 9/9281 | 2.18E-02 | 6.51E-02 | 3.96E-02 | daf-3/daf-7 | 2 |
| GO:0030431 | sleep | 2/243 | 9/9281 | 2.18E-02 | 6.51E-02 | 3.96E-02 | flp-11/egl-4 | 2 |
| GO:0045887 | positive regulation of synaptic growth at neuromuscular junction | 2/243 | 9/9281 | 2.18E-02 | 6.51E-02 | 3.96E-02 | egl-21/unc-104 | 2 |
| GO:0045923 | positive regulation of fatty acid metabolic process | 2/243 | 9/9281 | 2.18E-02 | 6.51E-02 | 3.96E-02 | mod-1/ser-6 | 2 |
| GO:0071482 | cellular response to light stimulus | 2/243 | 9/9281 | 2.18E-02 | 6.51E-02 | 3.96E-02 | lite-1/npr-3 | 2 |
| GO:1901244 | positive regulation of transcription from RNA polymerase II promoter involved in defense response to fungus | 2/243 | 9/9281 | 2.18E-02 | 6.51E-02 | 3.96E-02 | gpa-12/pmk-1 | 2 |
| GO:1902803 | regulation of synaptic vesicle transport | 2/243 | 9/9281 | 2.18E-02 | 6.51E-02 | 3.96E-02 | tom-1/unc-10 | 2 |
| GO:1904398 | positive regulation of neuromuscular junction development | 2/243 | 9/9281 | 2.18E-02 | 6.51E-02 | 3.96E-02 | egl-21/unc-104 | 2 |
| GO:0032535 | regulation of cellular component size | 7/243 | 107/9281 | 2.20E-02 | 6.58E-02 | 4.00E-02 | sma-3/unc-40/unc-60/unc-69/unc-76/unc-104/rig-6 | 7 |
| GO:0006066 | alcohol metabolic process | 5/243 | 62/9281 | 2.28E-02 | 6.73E-02 | 4.09E-02 | cat-4/egl-8/tbh-1/tdc-1/unc-26 | 5 |
| GO:0042592 | homeostatic process | 13/243 | 267/9281 | 2.32E-02 | 6.73E-02 | 4.09E-02 | acr-2/acr-3/egl-15/ftn-2/spe-41/trp-1/trp-2/unc-32/unc-38/unc-63/unc-80/unc-108/ser-5 | 13 |
| GO:0048523 | negative regulation of cellular process | 28/243 | 723/9281 | 2.33E-02 | 6.73E-02 | 4.09E-02 | daf-3/daf-5/dat-1/egl-5/egl-10/egl-13/egl-20/egl-27/egl-46/gar-2/gld-1/goa-1/gsa-1/hcf-1/ins-18/jnk-1/lon-2/osm-9/pmk-1/rgs-6/sem-4/srp-8/unc-2/unc-37/unc-40/unc-55/unc-122/egl-4 | 28 |
| GO:0008355 | olfactory learning | 3/243 | 24/9281 | 2.39E-02 | 6.73E-02 | 4.09E-02 | gpa-2/osm-9/egl-4 | 3 |
| GO:0017145 | stem cell division | 3/243 | 24/9281 | 2.39E-02 | 6.73E-02 | 4.09E-02 | egl-13/egl-44/egl-46 | 3 |
| GO:0010453 | regulation of cell fate commitment | 5/243 | 63/9281 | 2.42E-02 | 6.73E-02 | 4.09E-02 | lin-39/lon-2/sma-2/sma-3/unc-40 | 5 |
| GO:1990138 | neuron projection extension | 5/243 | 63/9281 | 2.42E-02 | 6.73E-02 | 4.09E-02 | unc-40/unc-69/unc-76/unc-104/rig-6 | 5 |
| GO:0080134 | regulation of response to stress | 8/243 | 135/9281 | 2.53E-02 | 6.73E-02 | 4.09E-02 | egl-8/egl-30/goa-1/jnk-1/nlp-2/pmk-1/octr-1/unc-16 | 8 |
| GO:0043068 | positive regulation of programmed cell death | 4/243 | 43/9281 | 2.55E-02 | 6.73E-02 | 4.09E-02 | ced-1/egl-1/unc-3/unc-32 | 4 |
| GO:0043902 | positive regulation of multi-organism process | 4/243 | 43/9281 | 2.55E-02 | 6.73E-02 | 4.09E-02 | gld-1/gpa-12/gsa-1/pmk-1 | 4 |
| GO:0048580 | regulation of post-embryonic development | 9/243 | 161/9281 | 2.55E-02 | 6.73E-02 | 4.09E-02 | daf-3/daf-5/daf-7/egl-20/ins-3/ins-4/ins-18/lin-39/egl-4 | 9 |
| GO:0061062 | regulation of nematode larval development | 9/243 | 161/9281 | 2.55E-02 | 6.73E-02 | 4.09E-02 | daf-3/daf-5/daf-7/egl-20/ins-3/ins-4/ins-18/lin-39/egl-4 | 9 |
| GO:0001993 | regulation of systemic arterial blood pressure by norepinephrine-epinephrine | 1/243 | 1/9281 | 2.62E-02 | 6.73E-02 | 4.09E-02 | ser-5 | 1 |
| GO:0001994 | norepinephrine-epinephrine vasoconstriction involved in regulation of systemic arterial blood pressure | 1/243 | 1/9281 | 2.62E-02 | 6.73E-02 | 4.09E-02 | ser-5 | 1 |
| GO:0003018 | vascular process in circulatory system | 1/243 | 1/9281 | 2.62E-02 | 6.73E-02 | 4.09E-02 | ser-5 | 1 |
| GO:0003321 | positive regulation of blood pressure by epinephrine-norepinephrine | 1/243 | 1/9281 | 2.62E-02 | 6.73E-02 | 4.09E-02 | ser-5 | 1 |
| GO:0006587 | serotonin biosynthetic process from tryptophan | 1/243 | 1/9281 | 2.62E-02 | 6.73E-02 | 4.09E-02 | tph-1 | 1 |
| GO:0007195 | adenylate cyclase-inhibiting dopamine receptor signaling pathway | 1/243 | 1/9281 | 2.62E-02 | 6.73E-02 | 4.09E-02 | dop-3 | 1 |
| GO:0007400 | neuroblast fate determination | 1/243 | 1/9281 | 2.62E-02 | 6.73E-02 | 4.09E-02 | unc-4 | 1 |
| GO:0009606 | tropism | 1/243 | 1/9281 | 2.62E-02 | 6.73E-02 | 4.09E-02 | egl-4 | 1 |
| GO:0009637 | response to blue light | 1/243 | 1/9281 | 2.62E-02 | 6.73E-02 | 4.09E-02 | lite-1 | 1 |
| GO:0009785 | blue light signaling pathway | 1/243 | 1/9281 | 2.62E-02 | 6.73E-02 | 4.09E-02 | lite-1 | 1 |
| GO:0010621 | negative regulation of transcription by transcription factor localization | 1/243 | 1/9281 | 2.62E-02 | 6.73E-02 | 4.09E-02 | hcf-1 | 1 |
| GO:0010649 | regulation of cell communication by electrical coupling | 1/243 | 1/9281 | 2.62E-02 | 6.73E-02 | 4.09E-02 | unc-1 | 1 |
| GO:0010650 | positive regulation of cell communication by electrical coupling | 1/243 | 1/9281 | 2.62E-02 | 6.73E-02 | 4.09E-02 | unc-1 | 1 |
| GO:0010807 | regulation of synaptic vesicle priming | 1/243 | 1/9281 | 2.62E-02 | 6.73E-02 | 4.09E-02 | tom-1 | 1 |
| GO:0014722 | regulation of skeletal muscle contraction by calcium ion signaling | 1/243 | 1/9281 | 2.62E-02 | 6.73E-02 | 4.09E-02 | unc-89 | 1 |
| GO:0014819 | regulation of skeletal muscle contraction | 1/243 | 1/9281 | 2.62E-02 | 6.73E-02 | 4.09E-02 | unc-89 | 1 |
| GO:0015874 | norepinephrine transport | 1/243 | 1/9281 | 2.62E-02 | 6.73E-02 | 4.09E-02 | dat-1 | 1 |
| GO:0016191 | synaptic vesicle uncoating | 1/243 | 1/9281 | 2.62E-02 | 6.73E-02 | 4.09E-02 | unc-26 | 1 |
| GO:0016264 | gap junction assembly | 1/243 | 1/9281 | 2.62E-02 | 6.73E-02 | 4.09E-02 | unc-7 | 1 |
| GO:0019229 | regulation of vasoconstriction | 1/243 | 1/9281 | 2.62E-02 | 6.73E-02 | 4.09E-02 | ser-5 | 1 |
| GO:0019614 | catechol-containing compound catabolic process | 1/243 | 1/9281 | 2.62E-02 | 6.73E-02 | 4.09E-02 | tbh-1 | 1 |
| GO:0030050 | vesicle transport along actin filament | 1/243 | 1/9281 | 2.62E-02 | 6.73E-02 | 4.09E-02 | jnk-1 | 1 |
| GO:0030845 | phospholipase C-inhibiting G protein-coupled receptor signaling pathway | 1/243 | 1/9281 | 2.62E-02 | 6.73E-02 | 4.09E-02 | eat-16 | 1 |
| GO:0032490 | detection of molecule of bacterial origin | 1/243 | 1/9281 | 2.62E-02 | 6.73E-02 | 4.09E-02 | daf-7 | 1 |
| GO:0035296 | regulation of tube diameter | 1/243 | 1/9281 | 2.62E-02 | 6.73E-02 | 4.09E-02 | ser-5 | 1 |
| GO:0035998 | 7,8-dihydroneopterin 3'-triphosphate biosynthetic process | 1/243 | 1/9281 | 2.62E-02 | 6.73E-02 | 4.09E-02 | cat-4 | 1 |
| GO:0042310 | vasoconstriction | 1/243 | 1/9281 | 2.62E-02 | 6.73E-02 | 4.09E-02 | ser-5 | 1 |
| GO:0042331 | phototaxis | 1/243 | 1/9281 | 2.62E-02 | 6.73E-02 | 4.09E-02 | lite-1 | 1 |
| GO:0042415 | norepinephrine metabolic process | 1/243 | 1/9281 | 2.62E-02 | 6.73E-02 | 4.09E-02 | tbh-1 | 1 |
| GO:0042420 | dopamine catabolic process | 1/243 | 1/9281 | 2.62E-02 | 6.73E-02 | 4.09E-02 | tbh-1 | 1 |
| GO:0042421 | norepinephrine biosynthetic process | 1/243 | 1/9281 | 2.62E-02 | 6.73E-02 | 4.09E-02 | tbh-1 | 1 |
| GO:0042424 | catecholamine catabolic process | 1/243 | 1/9281 | 2.62E-02 | 6.73E-02 | 4.09E-02 | tbh-1 | 1 |
| GO:0043577 | chemotropism | 1/243 | 1/9281 | 2.62E-02 | 6.73E-02 | 4.09E-02 | egl-4 | 1 |
| GO:0045759 | negative regulation of action potential | 1/243 | 1/9281 | 2.62E-02 | 6.73E-02 | 4.09E-02 | flp-3 | 1 |
| GO:0045777 | positive regulation of blood pressure | 1/243 | 1/9281 | 2.62E-02 | 6.73E-02 | 4.09E-02 | ser-5 | 1 |
| GO:0045907 | positive regulation of vasoconstriction | 1/243 | 1/9281 | 2.62E-02 | 6.73E-02 | 4.09E-02 | ser-5 | 1 |
| GO:0045988 | negative regulation of striated muscle contraction | 1/243 | 1/9281 | 2.62E-02 | 6.73E-02 | 4.09E-02 | flp-16 | 1 |
| GO:0046621 | negative regulation of organ growth | 1/243 | 1/9281 | 2.62E-02 | 6.73E-02 | 4.09E-02 | egl-4 | 1 |
| GO:0046957 | negative phototaxis | 1/243 | 1/9281 | 2.62E-02 | 6.73E-02 | 4.09E-02 | lite-1 | 1 |
| GO:0048212 | Golgi vesicle uncoating | 1/243 | 1/9281 | 2.62E-02 | 6.73E-02 | 4.09E-02 | unc-26 | 1 |
| GO:0048664 | neuron fate determination | 1/243 | 1/9281 | 2.62E-02 | 6.73E-02 | 4.09E-02 | egl-13 | 1 |
| GO:0048867 | stem cell fate determination | 1/243 | 1/9281 | 2.62E-02 | 6.73E-02 | 4.09E-02 | unc-4 | 1 |
| GO:0050849 | negative regulation of calcium-mediated signaling | 1/243 | 1/9281 | 2.62E-02 | 6.73E-02 | 4.09E-02 | egl-4 | 1 |
| GO:0050880 | regulation of blood vessel size | 1/243 | 1/9281 | 2.62E-02 | 6.73E-02 | 4.09E-02 | ser-5 | 1 |
| GO:0051066 | dihydrobiopterin metabolic process | 1/243 | 1/9281 | 2.62E-02 | 6.73E-02 | 4.09E-02 | cat-4 | 1 |
| GO:0051103 | DNA ligation involved in DNA repair | 1/243 | 1/9281 | 2.62E-02 | 6.73E-02 | 4.09E-02 | lig-4 | 1 |
| GO:0051342 | regulation of cyclic-nucleotide phosphodiesterase activity | 1/243 | 1/9281 | 2.62E-02 | 6.73E-02 | 4.09E-02 | egl-4 | 1 |
| GO:0051343 | positive regulation of cyclic-nucleotide phosphodiesterase activity | 1/243 | 1/9281 | 2.62E-02 | 6.73E-02 | 4.09E-02 | egl-4 | 1 |
| GO:0051583 | dopamine uptake involved in synaptic transmission | 1/243 | 1/9281 | 2.62E-02 | 6.73E-02 | 4.09E-02 | dat-1 | 1 |
| GO:0051934 | catecholamine uptake involved in synaptic transmission | 1/243 | 1/9281 | 2.62E-02 | 6.73E-02 | 4.09E-02 | dat-1 | 1 |
| GO:0052129 | negative energy taxis | 1/243 | 1/9281 | 2.62E-02 | 6.73E-02 | 4.09E-02 | lite-1 | 1 |
| GO:0060850 | regulation of transcription involved in cell fate commitment | 1/243 | 1/9281 | 2.62E-02 | 6.73E-02 | 4.09E-02 | unc-86 | 1 |
| GO:0070141 | response to UV-A | 1/243 | 1/9281 | 2.62E-02 | 6.73E-02 | 4.09E-02 | lite-1 | 1 |
| GO:0071483 | cellular response to blue light | 1/243 | 1/9281 | 2.62E-02 | 6.73E-02 | 4.09E-02 | lite-1 | 1 |
| GO:0071868 | cellular response to monoamine stimulus | 1/243 | 1/9281 | 2.62E-02 | 6.73E-02 | 4.09E-02 | dat-1 | 1 |
| GO:0071869 | response to catecholamine | 1/243 | 1/9281 | 2.62E-02 | 6.73E-02 | 4.09E-02 | dat-1 | 1 |
| GO:0071870 | cellular response to catecholamine stimulus | 1/243 | 1/9281 | 2.62E-02 | 6.73E-02 | 4.09E-02 | dat-1 | 1 |
| GO:0090327 | negative regulation of locomotion involved in locomotory behavior | 1/243 | 1/9281 | 2.62E-02 | 6.73E-02 | 4.09E-02 | rig-6 | 1 |
| GO:0097154 | GABAergic neuron differentiation | 1/243 | 1/9281 | 2.62E-02 | 6.73E-02 | 4.09E-02 | unc-30 | 1 |
| GO:0097306 | cellular response to alcohol | 1/243 | 1/9281 | 2.62E-02 | 6.73E-02 | 4.09E-02 | egl-4 | 1 |
| GO:0097746 | regulation of blood vessel diameter | 1/243 | 1/9281 | 2.62E-02 | 6.73E-02 | 4.09E-02 | ser-5 | 1 |
| GO:0097756 | negative regulation of blood vessel diameter | 1/243 | 1/9281 | 2.62E-02 | 6.73E-02 | 4.09E-02 | ser-5 | 1 |
| GO:0099148 | regulation of synaptic vesicle docking | 1/243 | 1/9281 | 2.62E-02 | 6.73E-02 | 4.09E-02 | tom-1 | 1 |
| GO:0099161 | regulation of presynaptic dense core granule exocytosis | 1/243 | 1/9281 | 2.62E-02 | 6.73E-02 | 4.09E-02 | tom-1 | 1 |
| GO:0099515 | actin filament-based transport | 1/243 | 1/9281 | 2.62E-02 | 6.73E-02 | 4.09E-02 | jnk-1 | 1 |
| GO:0106020 | regulation of vesicle docking | 1/243 | 1/9281 | 2.62E-02 | 6.73E-02 | 4.09E-02 | tom-1 | 1 |
| GO:1901419 | regulation of response to alcohol | 1/243 | 1/9281 | 2.62E-02 | 6.73E-02 | 4.09E-02 | egl-4 | 1 |
| GO:1901421 | positive regulation of response to alcohol | 1/243 | 1/9281 | 2.62E-02 | 6.73E-02 | 4.09E-02 | egl-4 | 1 |
| GO:1901424 | response to toluene | 1/243 | 1/9281 | 2.62E-02 | 6.73E-02 | 4.09E-02 | unc-79 | 1 |
| GO:1901951 | regulation of anterograde dense core granule transport | 1/243 | 1/9281 | 2.62E-02 | 6.73E-02 | 4.09E-02 | unc-104 | 1 |
| GO:1901953 | positive regulation of anterograde dense core granule transport | 1/243 | 1/9281 | 2.62E-02 | 6.73E-02 | 4.09E-02 | unc-104 | 1 |
| GO:1903522 | regulation of blood circulation | 1/243 | 1/9281 | 2.62E-02 | 6.73E-02 | 4.09E-02 | ser-5 | 1 |
| GO:1903524 | positive regulation of blood circulation | 1/243 | 1/9281 | 2.62E-02 | 6.73E-02 | 4.09E-02 | ser-5 | 1 |
| GO:1904080 | positive regulation of transcription from RNA polymerase II promoter involved in neuron fate specification | 1/243 | 1/9281 | 2.62E-02 | 6.73E-02 | 4.09E-02 | unc-86 | 1 |
| GO:1904122 | positive regulation of fatty acid beta-oxidation by octopamine signaling pathway | 1/243 | 1/9281 | 2.62E-02 | 6.73E-02 | 4.09E-02 | ser-6 | 1 |
| GO:1904123 | positive regulation of fatty acid beta-oxidation by serotonin receptor signaling pathway | 1/243 | 1/9281 | 2.62E-02 | 6.73E-02 | 4.09E-02 | mod-1 | 1 |
| GO:1905550 | regulation of protein localization to endoplasmic reticulum | 1/243 | 1/9281 | 2.62E-02 | 6.73E-02 | 4.09E-02 | unc-89 | 1 |
| GO:1905552 | positive regulation of protein localization to endoplasmic reticulum | 1/243 | 1/9281 | 2.62E-02 | 6.73E-02 | 4.09E-02 | unc-89 | 1 |
| GO:1905841 | response to oxidopamine | 1/243 | 1/9281 | 2.62E-02 | 6.73E-02 | 4.09E-02 | dat-1 | 1 |
| GO:1905842 | cellular response to oxidopamine | 1/243 | 1/9281 | 2.62E-02 | 6.73E-02 | 4.09E-02 | dat-1 | 1 |
| GO:1905846 | regulation of cellular response to oxidopamine | 1/243 | 1/9281 | 2.62E-02 | 6.73E-02 | 4.09E-02 | dat-1 | 1 |
| GO:1905847 | negative regulation of cellular response to oxidopamine | 1/243 | 1/9281 | 2.62E-02 | 6.73E-02 | 4.09E-02 | dat-1 | 1 |
| GO:1905957 | regulation of cellular response to alcohol | 1/243 | 1/9281 | 2.62E-02 | 6.73E-02 | 4.09E-02 | egl-4 | 1 |
| GO:1905959 | positive regulation of cellular response to alcohol | 1/243 | 1/9281 | 2.62E-02 | 6.73E-02 | 4.09E-02 | egl-4 | 1 |
| GO:1990335 | process resulting in tolerance to alcohol | 1/243 | 1/9281 | 2.62E-02 | 6.73E-02 | 4.09E-02 | egl-4 | 1 |
| GO:2000293 | negative regulation of defecation | 1/243 | 1/9281 | 2.62E-02 | 6.73E-02 | 4.09E-02 | rig-6 | 1 |
| GO:2000622 | regulation of nuclear-transcribed mRNA catabolic process, nonsense-mediated decay | 1/243 | 1/9281 | 2.62E-02 | 6.73E-02 | 4.09E-02 | gld-1 | 1 |
| GO:2000623 | negative regulation of nuclear-transcribed mRNA catabolic process, nonsense-mediated decay | 1/243 | 1/9281 | 2.62E-02 | 6.73E-02 | 4.09E-02 | gld-1 | 1 |
| GO:2000747 | negative regulation of defecation rhythm | 1/243 | 1/9281 | 2.62E-02 | 6.73E-02 | 4.09E-02 | rig-6 | 1 |
| GO:2001024 | negative regulation of response to drug | 1/243 | 1/9281 | 2.62E-02 | 6.73E-02 | 4.09E-02 | pkc-1 | 1 |
| GO:0031347 | regulation of defense response | 3/243 | 25/9281 | 2.67E-02 | 6.81E-02 | 4.14E-02 | egl-30/pmk-1/octr-1 | 3 |
| GO:0002252 | immune effector process | 2/243 | 10/9281 | 2.67E-02 | 6.81E-02 | 4.14E-02 | gpa-12/pmk-1 | 2 |
| GO:0050913 | sensory perception of bitter taste | 2/243 | 10/9281 | 2.67E-02 | 6.81E-02 | 4.14E-02 | gpa-3/egl-4 | 2 |
| GO:0060290 | transdifferentiation | 2/243 | 10/9281 | 2.67E-02 | 6.81E-02 | 4.14E-02 | egl-27/sem-4 | 2 |
| GO:1900424 | regulation of defense response to bacterium | 2/243 | 10/9281 | 2.67E-02 | 6.81E-02 | 4.14E-02 | egl-30/pmk-1 | 2 |
| GO:1901216 | positive regulation of neuron death | 2/243 | 10/9281 | 2.67E-02 | 6.81E-02 | 4.14E-02 | dat-1/jnk-1 | 2 |
| GO:1902473 | regulation of protein localization to synapse | 2/243 | 10/9281 | 2.67E-02 | 6.81E-02 | 4.14E-02 | egl-30/unc-104 | 2 |
| GO:0000041 | transition metal ion transport | 4/243 | 44/9281 | 2.74E-02 | 6.98E-02 | 4.24E-02 | ftn-2/spe-41/trp-1/trp-2 | 4 |
| GO:0007369 | gastrulation | 6/243 | 88/9281 | 2.76E-02 | 7.00E-02 | 4.26E-02 | lin-39/lon-2/sem-4/sma-2/sma-3/unc-40 | 6 |
| GO:0032870 | cellular response to hormone stimulus | 6/243 | 88/9281 | 2.76E-02 | 7.00E-02 | 4.26E-02 | gar-2/goa-1/gpb-2/ins-18/unc-55/egl-4 | 6 |
| GO:0009725 | response to hormone | 6/243 | 89/9281 | 2.90E-02 | 7.34E-02 | 4.46E-02 | gar-2/goa-1/gpb-2/ins-18/unc-55/egl-4 | 6 |
| GO:0006865 | amino acid transport | 4/243 | 45/9281 | 2.95E-02 | 7.47E-02 | 4.54E-02 | snf-1/snf-5/snf-11/unc-26 | 4 |
| GO:0007528 | neuromuscular junction development | 3/243 | 26/9281 | 2.96E-02 | 7.47E-02 | 4.54E-02 | egl-21/ins-4/unc-104 | 3 |
| GO:0000165 | MAPK cascade | 6/243 | 90/9281 | 3.04E-02 | 7.67E-02 | 4.66E-02 | daf-7/jnk-1/pkc-1/pmk-1/unc-16/unc-43 | 6 |
| GO:0000132 | establishment of mitotic spindle orientation | 4/243 | 46/9281 | 3.17E-02 | 7.96E-02 | 4.84E-02 | ced-1/dnc-1/egl-20/gpa-16 | 4 |
| GO:0008356 | asymmetric cell division | 4/243 | 46/9281 | 3.17E-02 | 7.96E-02 | 4.84E-02 | egl-13/egl-20/egl-27/egl-44 | 4 |
| GO:1903829 | positive regulation of cellular protein localization | 4/243 | 46/9281 | 3.17E-02 | 7.96E-02 | 4.84E-02 | egl-30/pmk-1/unc-89/unc-104 | 4 |
| GO:0061061 | muscle structure development | 6/243 | 91/9281 | 3.19E-02 | 7.97E-02 | 4.84E-02 | egl-15/sem-4/unc-60/unc-89/unc-98/unc-105 | 6 |
| GO:0002833 | positive regulation of response to biotic stimulus | 2/243 | 11/9281 | 3.21E-02 | 7.97E-02 | 4.84E-02 | gpa-12/pmk-1 | 2 |
| GO:0010042 | response to manganese ion | 2/243 | 11/9281 | 3.21E-02 | 7.97E-02 | 4.84E-02 | dat-1/jnk-1 | 2 |
| GO:0017158 | regulation of calcium ion-dependent exocytosis | 2/243 | 11/9281 | 3.21E-02 | 7.97E-02 | 4.84E-02 | tom-1/unc-10 | 2 |
| GO:0030511 | positive regulation of transforming growth factor beta receptor signaling pathway | 2/243 | 11/9281 | 3.21E-02 | 7.97E-02 | 4.84E-02 | sma-3/tph-1 | 2 |
| GO:0043052 | thermotaxis | 2/243 | 11/9281 | 3.21E-02 | 7.97E-02 | 4.84E-02 | egl-8/npr-1 | 2 |
| GO:0062013 | positive regulation of small molecule metabolic process | 2/243 | 11/9281 | 3.21E-02 | 7.97E-02 | 4.84E-02 | mod-1/ser-6 | 2 |
| GO:1903305 | regulation of regulated secretory pathway | 2/243 | 11/9281 | 3.21E-02 | 7.97E-02 | 4.84E-02 | tom-1/unc-10 | 2 |
| GO:1903846 | positive regulation of cellular response to transforming growth factor beta stimulus | 2/243 | 11/9281 | 3.21E-02 | 7.97E-02 | 4.84E-02 | sma-3/tph-1 | 2 |

| | | | | | | | | |
|---|---|---|---|---|---|---|---|---|
| GO:1905489 | regulation of sensory neuron axon guidance | 2/243 | 11/9281 | 3.21E-02 | 7.97E-02 | 4.84E-02 | egl-20/unc-40 | 2 |
| GO:0031345 | negative regulation of cell projection organization | 3/243 | 27/9281 | 3.26E-02 | 8.08E-02 | 4.91E-02 | goa-1/jnk-1/unc-40 | 3 |
