## Supplemental Table 1(5) for "Rebalancing the motor circuit restores movement in a *Caenorhabditis elegans* model for TDP-43-toxicity"

| GO-cluster | GO term | GO term name | GO-cluster name |
|---|---|---|---|
| 1 | GO:0007187 | G protein-coupled receptor signaling pathway, coupled to cyclic nucleotide second messenger | G protein-coupled receptor signaling pathway |
| 1 | GO:0007188 | adenylate cyclase-modulating G protein-coupled receptor signaling pathway | |
| 2 | GO:2000241 | regulation of reproductive process | regulation of behavior |
| 2 | GO:0046662 | regulation of oviposition | |
| 2 | GO:0018991 | oviposition | |
| 2 | GO:0019098 | reproductive behavior | |
| 2 | GO:0050795 | regulation of behavior | |
| 2 | GO:0007631 | feeding behavior | |
| 2 | GO:0043050 | pharyngeal pumping | |
| 2 | GO:0042755 | eating behavior | |
| 2 | GO:0060259 | regulation of feeding behavior | |
| 2 | GO:0043051 | regulation of pharyngeal pumping | |
| 2 | GO:1903998 | regulation of eating behavior | |
| 2 | GO:0042493 | response to drug | |
| 3 | GO:0006836 | neurotransmitter transport | regulation of synaptic transmission (cholinergic) |
| 3 | GO:0001505 | regulation of neurotransmitter levels | |
| 3 | GO:0007269 | neurotransmitter secretion | |
| 3 | GO:0099643 | signal release from synapse | |
| 3 | GO:0023061 | signal release | |
| 3 | GO:0032940 | secretion by cell | |
| 3 | GO:0046903 | secretion | |
| 3 | GO:0051046 | regulation of secretion | |
| 3 | GO:1903530 | regulation of secretion by cell | |
| 3 | GO:0046928 | regulation of neurotransmitter secretion | |
| 3 | GO:0051588 | regulation of neurotransmitter transport | |
| 3 | GO:0060341 | regulation of cellular localization | |
| 3 | GO:0050804 | modulation of chemical synaptic transmission | |
| 3 | GO:0099177 | regulation of trans-synaptic signaling | |
| 3 | GO:0007271 | synaptic transmission, cholinergic | |
| 3 | GO:0032222 | regulation of synaptic transmission, cholinergic | |
| 3 | GO:0032224 | positive regulation of synaptic transmission, cholinergic | |
| 3 | GO:0050806 | positive regulation of synaptic transmission | |
| 3 | GO:0023056 | positive regulation of signaling | |
| 3 | GO:0010647 | positive regulation of cell communication | |
| 4 | GO:0044057 | regulation of system process | regulation of muscle system process |
| 4 | GO:0090257 | regulation of muscle system process | |
| 4 | GO:0006937 | regulation of muscle contraction | |
| 4 | GO:0003012 | muscle system process | |
| 4 | GO:0006936 | muscle contraction | |
| 5 | GO:0042391 | regulation of membrane potential | calcium ion homeostasis |
| 5 | GO:0060079 | excitatory postsynaptic potential | |
| 5 | GO:0099565 | chemical synaptic transmission, postsynaptic | |
| 5 | GO:0060078 | regulation of postsynaptic membrane potential | |
| 5 | GO:1905114 | cell surface receptor signaling pathway involved in cell-cell signaling | |
| 5 | GO:0070509 | calcium ion import | |
| 5 | GO:0098703 | calcium ion import across plasma membrane | |
| 5 | GO:1902656 | calcium ion import into cytosol | |
| 5 | GO:0060402 | calcium ion transport into cytosol | |
| 5 | GO:0097553 | calcium ion transmembrane import into cytosol | |
| 5 | GO:0007204 | positive regulation of cytosolic calcium ion concentration | |
| 5 | GO:0072507 | divalent inorganic cation homeostasis | |
| 5 | GO:0072503 | cellular divalent inorganic cation homeostasis | |
| 5 | GO:0055074 | calcium ion homeostasis | |
| 5 | GO:0006874 | cellular calcium ion homeostasis | |
| 5 | GO:0051480 | regulation of cytosolic calcium ion concentration | |
| 5 | GO:0006875 | cellular metal ion homeostasis | |
| 5 | GO:0006873 | cellular ion homeostasis | |
| 5 | GO:0030003 | cellular cation homeostasis | |
| 5 | GO:0055082 | cellular chemical homeostasis | |
| 5 | GO:0055080 | cation homeostasis | |
| 5 | GO:0050801 | ion homeostasis | |
| 5 | GO:0070588 | calcium ion transmembrane transport | |
| 5 | GO:0072511 | divalent inorganic cation transport | |
| 5 | GO:0006816 | calcium ion transport | |
| 5 | GO:0070838 | divalent metal ion transport | |
| 6 | GO:1901700 | response to oxygen-containing compound | |
| 6 | GO:1901701 | cellular response to oxygen-containing compound | response to endogenous stimulus |
| 6 | GO:1901699 | cellular response to nitrogen compound | |
| 6 | GO:0032223 | negative regulation of synaptic transmission, cholinergic | |
| 6 | GO:0010648 | negative regulation of cell communication | |
| 6 | GO:0023057 | negative regulation of signaling | |
| 6 | GO:0048585 | negative regulation of response to stimulus | |
| 6 | GO:0009719 | response to endogenous stimulus | |
| 6 | GO:0071495 | cellular response to endogenous stimulus | |
| 6 | GO:0007167 | enzyme linked receptor protein signaling pathway | |
| 6 | GO:0071363 | cellular response to growth factor stimulus | |
| 6 | GO:0070848 | response to growth factor | |
| 6 | GO:0007178 | transmembrane receptor protein serine/threonine kinase signaling pathway | |
| 6 | GO:0007179 | transforming growth factor beta receptor signaling pathway | |
| 6 | GO:0071559 | response to transforming growth factor beta | |
| 6 | GO:0071560 | cellular response to transforming growth factor beta stimulus | |
| 6 | GO:0040024 | dauer larval development | |
| 6 | GO:0061065 | regulation of dauer larval development | |
| 7 | GO:0010469 | regulation of signaling receptor activity | regulation of signaling receptor activity |
| 8 | GO:0051241 | negative regulation of multicellular organismal process | negative regulation of behavior |
| 8 | GO:0048521 | negative regulation of behavior | |
| 8 | GO:1901045 | negative regulation of oviposition | |
| 8 | GO:2000242 | negative regulation of reproductive process | |
| 8 | GO:1902435 | regulation of male mating behavior | |
| 8 | GO:1902436 | negative regulation of male mating behavior | |
| 8 | GO:0061094 | regulation of turning behavior involved in mating | |
| 8 | GO:0061096 | negative regulation of turning behavior involved in mating | |
| 8 | GO:0043900 | regulation of multi-organism process | |
| 8 | GO:0035178 | turning behavior | |
| 8 | GO:0034607 | turning behavior involved in mating | |
| 8 | GO:0060179 | male mating behavior | |
| 8 | GO:0007618 | mating | |
| 8 | GO:0007617 | mating behavior | |
| 8 | GO:0007626 | locomotory behavior | |
| 8 | GO:0031987 | locomotion involved in locomotory behavior | |
| 8 | GO:0090325 | regulation of locomotion involved in locomotory behavior | |
| 8 | GO:0030536 | larval feeding behavior | |

| | | | |
|---|---|---|---|
| 8 | GO:0035690 | cellular response to drug | |
| 8 | GO:0072347 | response to anesthetic | |
| 8 | GO:0010037 | response to carbon dioxide | |
| 8 | GO:0071244 | cellular response to carbon dioxide | |
| 8 | GO:1903745 | negative regulation of pharyngeal pumping | |
| 8 | GO:1903999 | negative regulation of eating behavior | |
| 8 | GO:2000252 | negative regulation of feeding behavior | |
| 8 | GO:0071241 | cellular response to inorganic substance | |
| 9 | GO:0015696 | ammonium transport | positive regulation of secretion |
| 9 | GO:0010817 | regulation of hormone levels | |
| 9 | GO:1901374 | acetate ester transport | |
| 9 | GO:0015870 | acetylcholine transport | |
| 9 | GO:0009914 | hormone transport | |
| 9 | GO:0015695 | organic cation transport | |
| 9 | GO:0051590 | positive regulation of neurotransmitter transport | |
| 9 | GO:0001956 | positive regulation of neurotransmitter secretion | |
| 9 | GO:1903532 | positive regulation of secretion by cell | |
| 9 | GO:0051047 | positive regulation of secretion | |
| 9 | GO:0015837 | amine transport | |
| 9 | GO:0014055 | acetylcholine secretion, neurotransmission | |
| 9 | GO:0046887 | positive regulation of hormone secretion | |
| 9 | GO:0051954 | positive regulation of amine transport | |
| 9 | GO:0014057 | positive regulation of acetylcholine secretion, neurotransmission | |
| 9 | GO:2001025 | positive regulation of response to drug | |
| 9 | GO:0014056 | regulation of acetylcholine secretion, neurotransmission | |
| 9 | GO:0061526 | acetylcholine secretion | |
| 9 | GO:2001023 | regulation of response to drug | |
| 9 | GO:0043269 | regulation of ion transport | |
| 10 | GO:0030421 | defecation | defecation/excretion |
| 10 | GO:0007588 | excretion | |
| 11 | GO:0045597 | positive regulation of cell differentiation | positive regulation of nervous system development |
| 11 | GO:0010720 | positive regulation of cell development | |
| 11 | GO:0045666 | positive regulation of neuron differentiation | |
| 11 | GO:0010976 | positive regulation of neuron projection development | |
| 11 | GO:0051962 | positive regulation of nervous system development | |
| 11 | GO:0050920 | regulation of chemotaxis | |
| 11 | GO:0050921 | positive regulation of chemotaxis | |
| 11 | GO:1902669 | positive regulation of axon guidance | |
| 12 | GO:0040013 | negative regulation of locomotion | negative regulation of locomotion |
| 13 | GO:0008037 | cell recognition | neuron recognition |
| 13 | GO:0008038 | neuron recognition | |
| 13 | GO:0008039 | synaptic target recognition | |
| 14 | GO:0090101 | negative regulation of transmembrane receptor protein serine/threonine kinase signaling pathway | regulation of cellular response to growth factor stimulus |
| 14 | GO:0030512 | negative regulation of transforming growth factor beta receptor signaling pathway | |
| 14 | GO:1903845 | negative regulation of cellular response to transforming growth factor beta stimulus | |
| 14 | GO:0090288 | negative regulation of cellular response to growth factor stimulus | |
| 14 | GO:0017015 | regulation of transforming growth factor beta receptor signaling pathway | |
| 14 | GO:1903844 | regulation of cellular response to transforming growth factor beta stimulus | |
| 14 | GO:0090287 | regulation of cellular response to growth factor stimulus | |
| 14 | GO:0090092 | regulation of transmembrane receptor protein serine/threonine kinase signaling pathway | |
| 14 | GO:0035545 | determination of left/right asymmetry in nervous system | |
| 15 | GO:0006942 | regulation of striated muscle contraction | regulation of striated muscle contraction |
| 15 | GO:0006941 | striated muscle contraction | |
| 16 | GO:0007211 | octopamine or tyramine signaling pathway | octopamine or tyramine signaling pathway |
| 17 | GO:0001504 | neurotransmitter uptake | neurotransmitter uptake |
| 18 | GO:0050909 | sensory perception of taste | sensory perception of taste |
| 19 | GO:0032228 | regulation of synaptic transmission, GABAergic | regulation of synaptic transmission, GABAergic |
| 19 | GO:0032230 | positive regulation of synaptic transmission, GABAergic | |
| 20 | GO:0006585 | dopamine biosynthetic process from tyrosine | dopamine biosynthetic process |
| 20 | GO:0042416 | dopamine biosynthetic process | |
| 20 | GO:0042417 | dopamine metabolic process | |
| 20 | GO:0009713 | catechol-containing compound biosynthetic process | |
| 20 | GO:0006584 | catecholamine metabolic process | |
| 20 | GO:0009712 | catechol-containing compound metabolic process | |
| 20 | GO:0042423 | catecholamine biosynthetic process | |
| 20 | GO:0046189 | phenol-containing compound biosynthetic process | |
| 20 | GO:0018958 | phenol-containing compound metabolic process | |
| 20 | GO:0042136 | neurotransmitter biosynthetic process | |
| 21 | GO:0048678 | response to axon injury | neuron projection regeneration |
| 21 | GO:0031103 | axon regeneration | |
| 21 | GO:0031102 | neuron projection regeneration | |
| 21 | GO:0031099 | regeneration | |
| 22 | GO:0035725 | sodium ion transmembrane transport | sodium ion transport |
| 22 | GO:0006814 | sodium ion transport | |
| 23 | GO:0071875 | adrenergic receptor signaling pathway | adenylate cyclase-activating G protein-coupled receptor signaling pathway |
| 23 | GO:0071880 | adenylate cyclase-activating adrenergic receptor signaling pathway | |
| 23 | GO:0019935 | cyclic-nucleotide-mediated signaling | |
| 23 | GO:0007189 | adenylate cyclase-activating G protein-coupled receptor signaling pathway | |
| 23 | GO:0019933 | cAMP-mediated signaling | |
| 23 | GO:0019932 | second-messenger-mediated signaling | |
| 24 | GO:0007212 | dopamine receptor signaling pathway | dopamine receptor signaling pathway |
| 24 | GO:0032094 | response to food | |
| 24 | GO:0031667 | response to nutrient levels | |
| 24 | GO:0009991 | response to extracellular stimulus | |
| 25 | GO:0098664 | G protein-coupled serotonin receptor signaling pathway | serotonin receptor signaling pathway |
| 25 | GO:0007210 | serotonin receptor signaling pathway | |
| 26 | GO:0007638 | mechanosensory behavior | regulation of mechanosensory behavior |
| 26 | GO:0009612 | response to mechanical stimulus | |
| 26 | GO:0009581 | detection of external stimulus | |
| 26 | GO:0009582 | detection of abiotic stimulus | |
| 26 | GO:0050982 | detection of mechanical stimulus | |
| 26 | GO:0050976 | detection of mechanical stimulus involved in sensory perception of touch | |
| 26 | GO:0050974 | detection of mechanical stimulus involved in sensory perception | |
| 26 | GO:0050975 | sensory perception of touch | |
| 26 | GO:0050954 | sensory perception of mechanical stimulus | |
| 26 | GO:0031644 | regulation of neurological system process | |
| 26 | GO:0031646 | positive regulation of neurological system process | |
| 26 | GO:1905792 | positive regulation of mechanosensory behavior | |
| 26 | GO:1905790 | regulation of mechanosensory behavior | |
| 26 | GO:1905789 | positive regulation of detection of mechanical stimulus involved in sensory perception of touch | |
| 26 | GO:0051931 | regulation of sensory perception | |
| 26 | GO:1905787 | regulation of detection of mechanical stimulus involved in sensory perception of touch | |
| 26 | GO:0048520 | positive regulation of behavior | |

| | | | |
|---|---|---|---|
| 26 | GO:0032103 | positive regulation of response to external stimulus | |
| 26 | GO:0032101 | regulation of response to external stimulus | |
| 27 | GO:0010941 | regulation of cell death | synaptic vesicle transport |
| 27 | GO:0010942 | positive regulation of cell death | |
| 27 | GO:0051656 | establishment of organelle localization | |
| 27 | GO:0051650 | establishment of vesicle localization | |
| 27 | GO:0051648 | vesicle localization | |
| 27 | GO:0097479 | synaptic vesicle localization | |
| 27 | GO:0099003 | vesicle-mediated transport in synapse | |
| 27 | GO:0097480 | establishment of synaptic vesicle localization | |
| 27 | GO:0048489 | synaptic vesicle transport | |
| 28 | GO:0040018 | positive regulation of multicellular organism growth | regulation of synapse organization |
| 28 | GO:0048639 | positive regulation of developmental growth | |
| 28 | GO:0035264 | multicellular organism growth | |
| 28 | GO:0040014 | regulation of multicellular organism growth | |
| 28 | GO:0048638 | regulation of developmental growth | |
| 28 | GO:0040008 | regulation of growth | |
| 28 | GO:0048589 | developmental growth | |
| 28 | GO:0007416 | synapse assembly | |
| 28 | GO:0050808 | synapse organization | |
| 28 | GO:0050807 | regulation of synapse organization | |
| 28 | GO:0050803 | regulation of synapse structure or activity | |
| 28 | GO:0050770 | regulation of axonogenesis | |
| 28 | GO:0022604 | regulation of cell morphogenesis | |
| 28 | GO:0010769 | regulation of cell morphogenesis involved in differentiation | |
| 28 | GO:0022603 | regulation of anatomical structure morphogenesis | |
| 28 | GO:0060284 | regulation of cell development | |
| 28 | GO:0051960 | regulation of nervous system development | |
| 28 | GO:0050767 | regulation of neurogenesis | |
| 28 | GO:0045664 | regulation of neuron differentiation | |
| 28 | GO:0010975 | regulation of neuron projection development | |
| 28 | GO:0120035 | regulation of plasma membrane bounded cell projection organization | |
| 28 | GO:0031344 | regulation of cell projection organization | |
| 28 | GO:0061564 | axon development | |
| 28 | GO:0007409 | axonogenesis | |
| 28 | GO:0048667 | cell morphogenesis involved in neuron differentiation | |
| 28 | GO:0000904 | cell morphogenesis involved in differentiation | |
| 28 | GO:0032990 | cell part morphogenesis | |
| 28 | GO:0048858 | cell projection morphogenesis | |
| 28 | GO:0120039 | plasma membrane bounded cell projection morphogenesis | |
| 28 | GO:0048812 | neuron projection morphogenesis | |
| 28 | GO:0051270 | regulation of cellular component movement | |
| 28 | GO:0051272 | positive regulation of cellular component movement | |
| 28 | GO:0040017 | positive regulation of locomotion | |
| 29 | GO:0007218 | neuropeptide signaling pathway | neuropeptide signaling pathway |
