## Supplemental Table 1(6) for "Rebalancing the motor circuit restores movement in a *Caenorhabditis elegans* model for TDP-43-toxicity"

| id | description | gene ratio | bg ratio | pvalue | p adjust | qvalue | gene id | count | bg count | observed | expected | hits coun | hits list (hTDP-43 hits) | hits ratio | OLO orde | cluster id |
|---|---|---|---|---|---|---|---|---|---|---|---|---|---|---|---|---|
| GO:00071 | G protein-coupled receptor signaling pathway, coupled to cyclic nucleotide | 33/243 | 57/9281 | 2.97964E-38 | 1.87618E-35 | 1.14063E-35 | dop-1/dop-2/egl-30/gar-2/goa-1/gpa-1/gpa-2/gpa-3/gpa-4/gpa-5/gpa-6/gpa-7/gpa-8/gpa-9/gpa-11/gpa-12/gpa- | 33 | 57 | 0.135802 | 0.006142 | 0 | | 0 | 1 | 1 |
| GO:00071 | adenylate cyclase-modulating G protein-coupled receptor signaling pathwa | 25/243 | 43/9281 | 3.26715E-29 | 3.63039E-27 | 2.2071E-27 | dop-1/egl-30/goa-1/gpa-1/gpa-2/gpa-3/gpa-4/gpa-5/gpa-6/gpa-7/gpa-8/gpa-9/gpa-11/gpa-12/gpa-13/gpa-14/gp | 25 | 43 | 0.102881 | 0.004633 | 0 | | 0 | 2 | 1 |
| GO:20002 | regulation of reproductive process | 25/243 | 172/9281 | 2.19227E-12 | 5.59621E-11 | 3.40223E-11 | eat-16/egl-6/egl-8/egl-10/egl-17/egl-30/flp-1/flp-10/flp-12/flp-17/flp-20/gld-1/goa-1/gpb-2/gsa-1/lev-1/mec-4/tp | 25 | 172 | 0.102881 | 0.018532 | 2 | flp-1/rig-6 | 0.08 | 3 | 2 |
| GO:00466 | regulation of oviposition | 19/243 | 76/9281 | 4.39612E-14 | 1.43177E-12 | 8.70448E-13 | eat-16/egl-6/egl-8/egl-10/egl-17/egl-30/flp-1/flp-10/flp-17/goa-1/gpb-2/lev-1/tph-1/unc-29/unc-38/unc-63/unc- | 19 | 76 | 0.078189 | 0.008189 | 1 | flp-1 | 0.052632 | 4 | 2 |
| GO:00189 | oviposition | 29/243 | 158/9281 | 5.45754E-17 | 2.29095E-15 | 1.39279E-15 | bas-1/eat-16/egl-1/egl-2/egl-6/egl-8/egl-10/egl-15/egl-17/egl-27/egl-30/egl-47/flp-1/flp-10/flp-17/goa-1/gpb-2/l | 29 | 158 | 0.119342 | 0.017024 | 2 | flp-1/unc-7 | 0.068966 | 5 | 2 |
| GO:00190 | reproductive behavior | 36/243 | 181/9281 | 3.81239E-22 | 2.76985E-20 | 1.68393E-20 | bas-1/cat-2/eat-16/egl-1/egl-2/egl-6/egl-8/egl-10/egl-15/egl-17/egl-27/egl-30/egl-47/flp-1/flp-10/flp-12/flp-17/fl | 36 | 181 | 0.148148 | 0.019502 | 4 | flp-1/rab-3/unc-10/unc-7 | 0.111111 | 6 | 2 |
| GO:00507 | regulation of behavior | 43/243 | 152/9281 | 2.87844E-33 | 6.79671E-31 | 4.13207E-31 | daf-3/daf-7/eat-16/egl-6/egl-8/egl-10/egl-17/egl-19/egl-21/egl-30/flp-1/flp-3/flp-10/flp-12/flp-16/flp-17/flp-20/g | 43 | 152 | 0.176955 | 0.016378 | 8 | flp-1/mec-10/mec-14/nlp-1/rab-3/rig-6/unc-10/unc-7 | 0.186047 | 7 | 2 |
| GO:00076 | feeding behavior | 25/243 | 81/9281 | 1.17955E-20 | 7.18764E-19 | 4.36974E-19 | daf-3/daf-7/eat-16/egl-8/egl-10/egl-19/egl-21/egl-30/flp-3/flp-16/flp-21/goa-1/gpb-2/nlp-1/nlp-14/npr-1/rab-3/t | 25 | 81 | 0.102881 | 0.008728 | 4 | nlp-1/rab-3/unc-10/unc-7 | 0.16 | 8 | 2 |
| GO:00430 | pharyngeal pumping | 21/243 | 61/9281 | 1.21525E-18 | 6.20436E-17 | 3.77195E-17 | daf-3/daf-7/eat-16/egl-8/egl-19/egl-21/egl-30/flp-3/flp-16/goa-1/gpb-2/nlp-1/nlp-14/rab-3/tph-1/unc-7/unc-10/ | 21 | 61 | 0.08642 | 0.006573 | 4 | nlp-1/rab-3/unc-10/unc-7 | 0.190476 | 9 | 2 |
| GO:00427 | eating behavior | 22/243 | 69/9281 | 1.20031E-18 | 6.20436E-17 | 3.77195E-17 | daf-3/daf-7/eat-16/egl-8/egl-19/egl-21/egl-30/flp-3/flp-16/goa-1/gpb-2/nlp-1/nlp-14/rab-3/tph-1/unc-7/unc-10/ | 22 | 69 | 0.090535 | 0.007435 | 4 | nlp-1/rab-3/unc-10/unc-7 | 0.181818 | 10 | 2 |
| GO:00602 | regulation of feeding behavior | 22/243 | 58/9281 | 1.47872E-20 | 8.72908E-19 | 5.30686E-19 | daf-3/daf-7/eat-16/egl-8/egl-19/egl-21/egl-30/flp-3/flp-16/goa-1/gpb-2/nlp-1/nlp-14/rab-3/tph-1/unc-7/unc-10/ | 22 | 58 | 0.090535 | 0.006249 | 4 | nlp-1/rab-3/unc-10/unc-7 | 0.181818 | 11 | 2 |
| GO:00430 | regulation of pharyngeal pumping | 21/243 | 46/9281 | 9.81298E-22 | 6.39197E-20 | 3.88601E-20 | daf-3/daf-7/eat-16/egl-8/egl-19/egl-21/egl-30/flp-3/flp-16/goa-1/gpb-2/nlp-1/nlp-14/rab-3/tph-1/unc-7/unc-10/ | 21 | 46 | 0.08642 | 0.004956 | 4 | nlp-1/rab-3/unc-10/unc-7 | 0.190476 | 12 | 2 |
| GO:19039 | regulation of eating behavior | 22/243 | 53/9281 | 1.36584E-21 | 8.60025E-20 | 5.22854E-20 | daf-3/daf-7/eat-16/egl-8/egl-19/egl-21/egl-30/flp-3/flp-16/goa-1/gpb-2/nlp-1/nlp-14/rab-3/tph-1/unc-7/unc-10/ | 22 | 53 | 0.090535 | 0.005711 | 4 | nlp-1/rab-3/unc-10/unc-7 | 0.181818 | 13 | 2 |
| GO:00424 | response to drug | 23/243 | 179/9281 | 2.25537E-10 | 4.58108E-09 | 2.78507E-09 | dat-1/egl-8/egl-21/egl-30/flp-16/gar-2/goa-1/gpb-2/nlp-1/nlp-14/npr-1/pkc-1/pmk-1/snf-6/unc-1/unc-7/unc-17/ | 23 | 179 | 0.09465 | 0.019287 | 5 | nlp-1/unc-1/unc-7/unc-79/unc-80 | 0.217391 | 14 | 2 |
| GO:00068 | neurotransmitter transport | 22/243 | 87/9281 | 3.0551E-16 | 1.15422E-14 | 7.01708E-15 | dat-1/egl-8/egl-21/egl-30/mod-5/pkc-1/ric-19/snf-1/snf-2/snf-5/snf-6/snf-7/snf-9/snf-10/snf-11/tom-1/unc-10/ur | 22 | 87 | 0.090535 | 0.009374 | 1 | unc-10 | 0.045455 | 15 | 3 |
| GO:00015 | regulation of neurotransmitter levels | 22/243 | 87/9281 | 3.0551E-16 | 1.15422E-14 | 7.01708E-15 | bas-1/cat-2/dat-1/egl-8/egl-21/egl-30/mod-5/pkc-1/ric-19/snf-6/tbh-1/tdc-1/tom-1/tph-1/unc-2/unc-10/unc-17/ | 22 | 87 | 0.090535 | 0.009374 | 3 | unc-10/unc-2/unc-37 | 0.136364 | 16 | 3 |
| GO:00072 | neurotransmitter secretion | 12/243 | 54/9281 | 1.02462E-08 | 1.47748E-07 | 8.98238E-08 | egl-8/egl-21/egl-30/pkc-1/ric-19/tom-1/unc-10/unc-17/unc-18/unc-26/unc-29/unc-31 | 12 | 54 | 0.049383 | 0.005818 | 1 | unc-10 | 0.083333 | 17 | 3 |
| GO:00996 | signal release from synapse | 12/243 | 54/9281 | 1.02462E-08 | 1.47748E-07 | 8.98238E-08 | egl-8/egl-21/egl-30/pkc-1/ric-19/tom-1/unc-10/unc-17/unc-18/unc-26/unc-29/unc-31 | 12 | 54 | 0.049383 | 0.005818 | 1 | unc-10 | 0.083333 | 18 | 3 |
| GO:00230 | signal release | 12/243 | 58/9281 | 2.42764E-08 | 3.25234E-07 | 1.97727E-07 | egl-8/egl-21/egl-30/pkc-1/ric-19/tom-1/unc-10/unc-17/unc-18/unc-26/unc-29/unc-31 | 12 | 58 | 0.049383 | 0.006249 | 1 | unc-10 | 0.083333 | 19 | 3 |
| GO:00329 | secretion by cell | 15/243 | 116/9281 | 3.15992E-07 | 3.35342E-06 | 2.03872E-06 | egl-8/egl-21/egl-30/pkc-1/rab-3/ric-19/sng-1/tom-1/unc-10/unc-17/unc-18/unc-26/unc-29/unc-31/unc-104 | 15 | 116 | 0.061728 | 0.012499 | 3 | rab-3/sng-1/unc-10 | 0.2 | 20 | 3 |
| GO:00469 | secretion | 17/243 | 139/9281 | 1.16027E-07 | 1.39602E-06 | 8.48712E-07 | egl-8/egl-21/egl-30/gsa-1/pkc-1/rab-3/ric-19/sng-1/tom-1/unc-10/unc-17/unc-18/unc-26/unc-29/unc-31/unc-10 | 17 | 139 | 0.069959 | 0.014977 | 4 | rab-3/rig-6/sng-1/unc-10 | 0.235294 | 21 | 3 |
| GO:00510 | regulation of secretion | 14/243 | 68/9281 | 1.71529E-09 | 2.79326E-08 | 1.69817E-08 | egl-8/egl-21/egl-30/gsa-1/rab-3/ric-19/sng-1/tom-1/unc-10/unc-17/unc-18/unc-26/unc-29/rig-6 | 14 | 68 | 0.057613 | 0.007327 | 4 | rab-3/rig-6/sng-1/unc-10 | 0.285714 | 22 | 3 |
| GO:19035 | regulation of secretion by cell | 12/243 | 53/9281 | 8.1565E-09 | 1.23261E-07 | 7.49368E-08 | egl-8/egl-21/egl-30/rab-3/ric-19/sng-1/tom-1/unc-10/unc-17/unc-18/unc-26/unc-29 | 12 | 53 | 0.049383 | 0.005711 | 3 | rab-3/sng-1/unc-10 | 0.25 | 23 | 3 |
| GO:00469 | regulation of neurotransmitter secretion | 10/243 | 28/9281 | 1.09205E-09 | 1.87535E-08 | 1.14012E-08 | egl-8/egl-21/egl-30/ric-19/tom-1/unc-10/unc-17/unc-18/unc-26/unc-29 | 10 | 28 | 0.041152 | 0.003017 | 1 | unc-10 | 0.1 | 24 | 3 |
| GO:00515 | regulation of neurotransmitter transport | 10/243 | 28/9281 | 1.09205E-09 | 1.87535E-08 | 1.14012E-08 | egl-8/egl-21/egl-30/ric-19/tom-1/unc-10/unc-17/unc-18/unc-26/unc-29 | 10 | 28 | 0.041152 | 0.003017 | 1 | unc-10 | 0.1 | 25 | 3 |
| GO:00603 | regulation of cellular localization | 13/243 | 148/9281 | 0.000130694 | 0.000818714 | 0.000497739 | egl-8/egl-21/egl-30/pmk-1/ric-19/tom-1/unc-10/unc-17/unc-18/unc-26/unc-29/unc-89/unc-104 | 13 | 148 | 0.053498 | 0.015947 | 2 | unc-10/unc-89 | 0.153846 | 26 | 3 |
| GO:00322 | positive regulation of synaptic transmission, cholinergic | 8/243 | 18/9281 | 6.87119E-09 | 1.06391E-07 | 6.46805E-08 | egl-8/egl-21/egl-30/nca-2/unc-10/unc-29/unc-75/unc-77 | 8 | 18 | 0.032922 | 0.001939 | 4 | nca-2/unc-10/unc-75/unc-77 | 0.5 | 31 | 3 |
| GO:00322 | regulation of synaptic transmission, cholinergic | 11/243 | 25/9281 | 1.02091E-11 | 2.44115E-10 | 1.4841E-10 | egl-8/egl-21/egl-30/gar-2/goa-1/nca-2/unc-10/unc-29/unc-75/unc-77/unc-122 | 11 | 25 | 0.045267 | 0.002694 | 5 | nca-2/unc-10/unc-122/unc-75/unc-77 | 0.454545 | 30 | 3 |
| GO:00072 | synaptic transmission, cholinergic | 17/243 | 43/9281 | 1.67867E-16 | 6.74684E-15 | 4.10176E-15 | egl-8/egl-21/egl-30/gar-2/goa-1/nca-2/rab-3/unc-10/unc-17/unc-18/unc-29/unc-32/unc-63/unc-75/unc-77/unc-1 | 17 | 43 | 0.069959 | 0.004633 | 7 | nca-2/rab-3/unc-10/unc-122/unc-32/unc-75/unc-77 | 0.411765 | 29 | 3 |
| GO:00508 | positive regulation of synaptic transmission | 10/243 | 32/9281 | 4.8953E-09 | 7.77078E-08 | 4.72426E-08 | egl-8/egl-21/egl-30/nca-2/unc-10/unc-18/unc-26/unc-29/unc-75/unc-77 | 10 | 32 | 0.041152 | 0.003448 | 4 | nca-2/unc-10/unc-75/unc-77 | 0.4 | 32 | 3 |
| GO:00508 | modulation of chemical synaptic transmission | 18/243 | 73/9281 | 2.7215E-13 | 8.03267E-12 | 4.88347E-12 | eat-16/egl-8/egl-21/egl-30/gar-2/goa-1/nca-2/rab-3/ric-19/tom-1/unc-10/unc-17/unc-18/unc-26/unc-29/unc-75/ | 18 | 73 | 0.074074 | 0.007866 | 6 | nca-2/rab-3/unc-10/unc-122/unc-75/unc-77 | 0.333333 | 27 | 3 |
| GO:00991 | regulation of trans-synaptic signaling | 18/243 | 73/9281 | 2.7215E-13 | 8.03267E-12 | 4.88347E-12 | eat-16/egl-8/egl-21/egl-30/gar-2/goa-1/nca-2/rab-3/ric-19/tom-1/unc-10/unc-17/unc-18/unc-26/unc-29/unc-75/ | 18 | 73 | 0.074074 | 0.007866 | 6 | nca-2/rab-3/unc-10/unc-122/unc-75/unc-77 | 0.333333 | 28 | 3 |
| GO:00230 | positive regulation of signaling | 16/243 | 158/9281 | 3.5625E-06 | 3.18936E-05 | 1.93898E-05 | daf-7/egl-8/egl-21/egl-30/nca-2/sma-3/tph-1/unc-10/unc-16/unc-18/unc-26/unc-29/unc-40/unc-75/unc-77/egl-4 | 16 | 158 | 0.065844 | 0.017024 | 4 | nca-2/unc-10/unc-75/unc-77 | 0.25 | 33 | 3 |
| GO:00106 | positive regulation of cell communication | 17/243 | 159/9281 | 8.16521E-07 | 8.11793E-06 | 4.93531E-06 | daf-7/egl-8/egl-21/egl-30/nca-2/sma-3/tph-1/unc-1/unc-10/unc-16/unc-18/unc-26/unc-29/unc-40/unc-75/unc-7 | 17 | 159 | 0.069959 | 0.017132 | 5 | nca-2/unc-1/unc-10/unc-75/unc-77 | 0.294118 | 34 | 3 |
| GO:00440 | regulation of system process | 28/243 | 79/9281 | 6.20506E-25 | 5.32789E-23 | 3.2391E-23 | acr-2/egl-19/egl-21/egl-30/flp-1/flp-16/flp-18/gsa-1/mec-4/mec-7/mec-10/mec-12/mec-14/npr-1/sup-9/unc-17/u | 28 | 79 | 0.115226 | 0.008512 | 5 | flp-1/mec-10/mec-14/rig-6/unc-89 | 0.178571 | 35 | 4 |
| GO:00902 | regulation of muscle system process | 20/243 | 52/9281 | 6.27737E-19 | 3.38799E-17 | 2.05973E-17 | acr-2/egl-19/egl-21/flp-1/flp-16/flp-18/npr-1/sup-9/unc-17/unc-26/unc-29/unc-31/unc-38/unc-63/unc-89/unc-1 | 20 | 52 | 0.082305 | 0.005603 | 2 | flp-1/unc-89 | 0.1 | 36 | 4 |
| GO:00069 | regulation of muscle contraction | 20/243 | 52/9281 | 6.27737E-19 | 3.38799E-17 | 2.05973E-17 | acr-2/egl-19/egl-21/flp-1/flp-16/flp-18/npr-1/sup-9/unc-17/unc-26/unc-29/unc-31/unc-38/unc-63/unc-89/unc-1 | 20 | 52 | 0.082305 | 0.005603 | 2 | flp-1/unc-89 | 0.1 | 37 | 4 |
| GO:00030 | muscle system process | 21/243 | 67/9281 | 1.12649E-17 | 5.06651E-16 | 3.08019E-16 | acr-2/egl-2/egl-19/egl-21/flp-1/flp-16/flp-18/npr-1/sup-9/unc-17/unc-26/unc-29/unc-31/unc-38/unc-63/unc-89/ | 21 | 67 | 0.08642 | 0.007219 | 2 | flp-1/unc-89 | 0.095238 | 38 | 4 |
| GO:00069 | muscle contraction | 21/243 | 67/9281 | 1.12649E-17 | 5.06651E-16 | 3.08019E-16 | acr-2/egl-2/egl-19/egl-21/flp-1/flp-16/flp-18/npr-1/sup-9/unc-17/unc-26/unc-29/unc-31/unc-38/unc-63/unc-89/ | 21 | 67 | 0.08642 | 0.007219 | 2 | flp-1/unc-89 | 0.095238 | 39 | 4 |
| GO:00423 | regulation of membrane potential | 26/243 | 172/9281 | 3.01589E-13 | 8.76464E-12 | 5.32848E-12 | acr-2/acr-3/acr-6/acr-7/acr-9/acr-10/acr-11/acr-14/acr-15/acr-18/acr-19/acr-21/acr-23/egl-2/egl-23/flp-3/flp-9/le | 26 | 172 | 0.106996 | 0.018532 | 1 | unc-10 | 0.038462 | 40 | 5 |
| GO:00600 | excitatory postsynaptic potential | 15/243 | 25/9281 | 3.13765E-18 | 1.51975E-16 | 9.23933E-17 | acr-2/acr-3/acr-6/acr-9/acr-10/acr-11/acr-14/acr-15/acr-19/acr-21/lev-1/lev-8/unc-29/unc-38/unc-63 | 15 | 25 | 0.061728 | 0.002694 | 0 | | 0 | 41 | 5 |
| GO:00995 | chemical synaptic transmission, postsynaptic | 15/243 | 26/9281 | 7.24516E-18 | 3.42153E-16 | 2.08012E-16 | acr-2/acr-3/acr-6/acr-9/acr-10/acr-11/acr-14/acr-15/acr-19/acr-21/lev-1/lev-8/unc-29/unc-38/unc-63 | 15 | 26 | 0.061728 | 0.002801 | 0 | | 0 | 42 | 5 |
| GO:00600 | regulation of postsynaptic membrane potential | 15/243 | 33/9281 | 8.25985E-16 | 3.00055E-14 | 1.82419E-14 | acr-2/acr-3/acr-6/acr-9/acr-10/acr-11/acr-14/acr-15/acr-19/acr-21/lev-1/lev-8/unc-29/unc-38/unc-63 | 15 | 33 | 0.061728 | 0.003556 | 0 | | 0 | 43 | 5 |
| GO:19051 | cell surface receptor signaling pathway involved in cell-cell signaling | 18/243 | 91/9281 | 1.53626E-11 | 3.6275E-10 | 2.20535E-10 | acr-2/acr-3/acr-6/acr-9/acr-10/acr-11/acr-14/acr-15/acr-19/acr-21/egl-20/egl-27/lev-1/lev-8/unc-29/unc-37/unc- | 18 | 91 | 0.074074 | 0.009805 | 1 | unc-37 | 0.055556 | 44 | 5 |
| GO:00705 | calcium ion import | 6/243 | 11/9281 | 1.25415E-07 | 1.48068E-06 | 9.0018E-07 | acr-2/acr-3/egl-19/unc-2/unc-38/unc-63 | 6 | 11 | 0.024691 | 0.001185 | 1 | unc-2 | 0.166667 | 45 | 5 |
| GO:00987 | calcium ion import across plasma membrane | 4/243 | 5/9281 | 2.24635E-06 | 2.11112E-05 | 1.28346E-05 | acr-2/acr-3/unc-38/unc-63 | 4 | 5 | 0.016461 | 0.000539 | 0 | | 0 | 46 | 5 |
| GO:19026 | calcium ion import into cytosol | 4/243 | 5/9281 | 2.24635E-06 | 2.11112E-05 | 1.28346E-05 | acr-2/acr-3/unc-38/unc-63 | 4 | 5 | 0.016461 | 0.000539 | 0 | | 0 | 47 | 5 |
| GO:00604 | calcium ion transport into cytosol | 4/243 | 12/9281 | 0.000192312 | 0.001107551 | 0.000673337 | acr-2/acr-3/unc-38/unc-63 | 4 | 12 | 0.016461 | 0.001293 | 0 | | 0 | 48 | 5 |
| GO:00975 | calcium ion transmembrane import into cytosol | 4/243 | 12/9281 | 0.000192312 | 0.001107551 | 0.000673337 | acr-2/acr-3/unc-38/unc-63 | 4 | 12 | 0.016461 | 0.001293 | 0 | | 0 | 49 | 5 |
| GO:00072 | positive regulation of cytosolic calcium ion concentration | 5/243 | 14/9281 | 1.94855E-05 | 0.000149627 | 9.09657E-05 | acr-2/acr-3/unc-38/unc-63/ser-5 | 5 | 14 | 0.020576 | 0.001508 | 0 | | 0 | 50 | 5 |
| GO:00725 | divalent inorganic cation homeostasis | 8/243 | 52/9281 | 5.46966E-05 | 0.000377087 | 0.000229251 | acr-2/acr-3/spe-41/trp-1/trp-2/unc-38/unc-63/ser-5 | 8 | 52 | 0.032922 | 0.005603 | 0 | | 0 | 51 | 5 |
| GO:00725 | cellular divalent inorganic cation homeostasis | 8/243 | 48/9281 | 3.00189E-05 | 0.000217263 | 0.000132086 | acr-2/acr-3/spe-41/trp-1/trp-2/unc-38/unc-63/ser-5 | 8 | 48 | 0.032922 | 0.005172 | 0 | | 0 | 52 | 5 |
| GO:00550 | calcium ion homeostasis | 8/243 | 41/9281 | 8.90434E-06 | 7.31318E-05 | 4.44606E-05 | acr-2/acr-3/spe-41/trp-1/trp-2/unc-38/unc-63/ser-5 | 8 | 41 | 0.032922 | 0.004418 | 0 | | 0 | 53 | 5 |
| GO:00068 | cellular calcium ion homeostasis | 8/243 | 40/9281 | 7.33094E-06 | 6.15473E-05 | 3.74178E-05 | acr-2/acr-3/spe-41/trp-1/trp-2/unc-38/unc-63/ser-5 | 8 | 40 | 0.032922 | 0.00431 | 0 | | 0 | 54 | 5 |
| GO:00514 | regulation of cytosolic calcium ion concentration | 8/243 | 22/9281 | 4.58483E-08 | 5.89166E-07 | 3.58185E-07 | acr-2/acr-3/spe-41/trp-1/trp-2/unc-38/unc-63/ser-5 | 8 | 22 | 0.032922 | 0.00237 | 0 | | 0 | 55 | 5 |
| GO:00068 | cellular metal ion homeostasis | 9/243 | 77/9281 | 0.000170418 | 0.001006631 | 0.000611983 | acr-2/acr-3/ftn-2/spe-41/trp-1/trp-2/unc-38/unc-63/ser-5 | 9 | 77 | 0.037037 | 0.008297 | 0 | | 0 | 56 | 5 |
| GO:00068 | cellular ion homeostasis | 11/243 | 110/9281 | 0.000137113 | 0.000846427 | 0.000514587 | acr-2/acr-3/ftn-2/spe-41/trp-1/trp-2/unc-32/unc-38/unc-63/unc-108/ser-5 | 11 | 110 | 0.045267 | 0.011852 | 1 | unc-32 | 0.090909 | 57 | 5 |
| GO:00300 | cellular cation homeostasis | 11/243 | 110/9281 | 0.000137113 | 0.000846427 | 0.000514587 | acr-2/acr-3/ftn-2/spe-41/trp-1/trp-2/unc-32/unc-38/unc-63/unc-108/ser-5 | 11 | 110 | 0.045267 | 0.011852 | 1 | unc-32 | 0.090909 | 58 | 5 |
| GO:00550 | cellular chemical homeostasis | 11/243 | 117/9281 | 0.000237584 | 0.001359989 | 0.000826808 | acr-2/acr-3/ftn-2/spe-41/trp-1/trp-2/unc-32/unc-38/unc-63/unc-108/ser-5 | 11 | 117 | 0.045267 | 0.012606 | 1 | unc-32 | 0.090909 | 59 | 5 |
| GO:00550 | cation homeostasis | 12/243 | 129/9281 | 0.000137591 | 0.000846608 | 0.000514697 | acr-2/acr-3/ftn-2/spe-41/trp-1/trp-2/unc-32/unc-38/unc-63/unc-80/unc-108/ser-5 | 12 | 129 | 0.049383 | 0.013899 | 2 | unc-32/unc-80 | 0.166667 | 60 | 5 |
| GO:00508 | ion homeostasis | 12/243 | 131/9281 | 0.000159343 | 0.000958594 | 0.000582779 | acr-2/acr-3/ftn-2/spe-41/trp-1/trp-2/unc-32/unc-38/unc-63/unc-80/unc-108/ser-5 | 12 | 131 | 0.049383 | 0.014115 | 2 | unc-32/unc-80 | 0.166667 | 61 | 5 |
| GO:00705 | calcium ion transmembrane transport | 16/243 | 48/9281 | 3.18736E-14 | 1.0563E-12 | 6.42181E-13 | acr-2/acr-3/egl-19/lov-1/ocr-1/ocr-2/ocr-3/ocr-4/osm-9/spe-41/trp-1/trp-2/trp-4/unc-2/unc-38/unc-63 | 16 | 48 | 0.065844 | 0.005172 | 2 | osm-9/unc-2 | 0.125 | 62 | 5 |
| GO:00725 | divalent inorganic cation transport | 17/243 | 81/9281 | 2.10923E-11 | 4.80039E-10 | 2.91841E-10 | acr-2/acr-3/egl-19/lov-1/ocr-1/ocr-2/ocr-3/ocr-4/osm-9/spe-41/trp-1/trp-2/trp-4/unc-2/unc-38/unc-63/unc-105 | 17 | 81 | 0.069959 | 0.008728 | 2 | osm-9/unc-2 | 0.117647 | 63 | 5 |
| GO:00068 | calcium ion transport | 17/243 | 58/9281 | 5.54881E-14 | 1.74695E-12 | 1.06206E-12 | acr-2/acr-3/egl-19/lov-1/ocr-1/ocr-2/ocr-3/ocr-4/osm-9/spe-41/trp-1/trp-2/trp-4/unc-2/unc-38/unc-63/unc-105 | 17 | 58 | 0.069959 | 0.006249 | 2 | osm-9/unc-2 | 0.117647 | 64 | 5 |
| GO:00708 | divalent metal ion transport | 17/243 | 81/9281 | 2.10923E-11 | 4.80039E-10 | 2.91841E-10 | acr-2/acr-3/egl-19/lov-1/ocr-1/ocr-2/ocr-3/ocr-4/osm-9/spe-41/trp-1/trp-2/trp-4/unc-2/unc-38/unc-63/unc-105 | 17 | 81 | 0.069959 | 0.008728 | 2 | osm-9/unc-2 | 0.117647 | 65 | 5 |
| GO:19017 | response to oxygen-containing compound | 16/243 | 111/9281 | 2.64877E-08 | 3.52361E-07 | 2.14218E-07 | daf-7/dat-1/egl-21/flp-16/gar-2/goa-1/gpa-3/gpb-2/ins-18/nlp-1/nlp-14/npr-1/pmk-1/unc-31/unc-79/egl-4 | 16 | 111 | 0.065844 | 0.01196 | 2 | nlp-1/unc-79 | 0.125 | 66 | 6 |
| GO:19017 | cellular response to oxygen-containing compound | 13/243 | 71/9281 | 2.97426E-08 | 3.92894E-07 | 2.38861E-07 | daf-7/dat-1/egl-21/flp-16/gar-2/goa-1/gpa-3/gpb-2/ins-18/nlp-1/nlp-14/unc-31/egl-4 | 13 | 71 | 0.053498 | 0.00765 | 1 | nlp-1 | 0.076923 | 67 | 6 |
| GO:19016 | cellular response to nitrogen compound | 8/243 | 50/9281 | 4.08238E-05 | 0.000291004 | 0.000176916 | daf-7/dat-1/gar-2/goa-1/gpa-3/gpb-2/ins-18/egl-4 | 8 | 50 | 0.032922 | 0.005387 | 0 | | 0 | 68 | 6 |
| GO:00322 | negative regulation of synaptic transmission, cholinergic | 3/243 | 5/9281 | 0.000170525 | 0.001006631 | 0.000611983 | gar-2/goa-1/unc-122 | 3 | 5 | 0.012346 | 0.000539 | 1 | unc-122 | 0.333333 | 69 | 6 |
| GO:00400 | dauer larval development | 12/243 | 114/9281 | 4.10765E-05 | 0.000291705 | 0.000177343 | daf-3/daf-5/daf-7/gpa-3/hcf-1/ins-3/ins-4/ins-18/pkc-1/tph-1/unc-3/egl-4 | 12 | 114 | 0.049383 | 0.012283 | 4 | hcf-1/ins-3/ins-4/unc-3 | 0.333333 | 82 | 6 |
| GO:00106 | negative regulation of cell communication | 13/243 | 141/9281 | 7.96238E-05 | 0.00052961 | 0.000321978 | daf-5/egl-10/gar-2/goa-1/gsa-1/ins-18/lon-2/osm-9/rgs-6/unc-2/unc-37/unc-122/egl-4 | 13 | 141 | 0.053498 | 0.015192 | 4 | osm-9/unc-122/unc-2/unc-37 | 0.307692 | 70 | 6 |
| GO:00485 | negative regulation of response to stimulus | 16/243 | 163/9281 | 5.35052E-06 | 4.67923E-05 | 2.84475E-05 | daf-5/dat-1/egl-10/goa-1/gsa-1/ins-18/jnk-1/lon-2/osm-9/pkc-1/rgs-6/octr-1/unc-2/unc-37/unc-40/egl-4 | 16 | 163 | 0.065844 | 0.017563 | 4 | jnk-1/osm-9/unc-2/unc-37 | 0.25 | 72 | 6 |
| GO:00097 | response to endogenous stimulus | 19/243 | 147/9281 | 8.08027E-09 | 1.23094E-07 | 7.48351E-08 | daf-3/daf-5/daf-7/dat-1/egl-15/egl-17/gar-2/goa-1/gpa-3/gpb-2/ins-18/lon-2/sma-2/sma-3/tph-1/unc-2/unc-40/ | 19 | 147 | 0.078189 | 0.015839 | 2 | unc-2/unc-55 | 0.105263 | 73 | 6 |
| GO:00714 | cellular response to endogenous stimulus | 19/243 | 146/9281 | 7.20207E-09 | 1.10607E-07 | 6.7244E-08 | daf-3/daf-5/daf-7/dat-1/egl-15/egl-17/gar-2/goa-1/gpa-3/gpb-2/ins-18/lon-2/sma-2/sma-3/tph-1/unc-2/unc-40/ | 19 | 146 | 0.078189 | 0.015731 | 2 | unc-2/unc-55 | 0.105263 | 74 | 6 |
| GO:00071 | enzyme linked receptor protein signaling pathway | 14/243 | 158/9281 | 6.55982E-05 | 0.000445737 | 0.000270987 | daf-3/daf-5/daf-7/egl-15/egl-17/ins-18/lon-2/pkc-1/sma-2/sma-3/tph-1/unc-2/unc-40/egl-4 | 14 | 158 | 0.057613 | 0.017024 | 1 | unc-2 | 0.071429 | 75 | 6 |
| GO:00713 | cellular response to growth factor stimulus | 12/243 | 46/9281 | 1.39971E-09 | 2.29917E-08 | 1.39778E-08 | daf-3/daf-5/daf-7/egl-15/egl-17/lon-2/sma-2/sma-3/tph-1/unc-2/unc-40/egl-4 | 12 | 46 | 0.049383 | 0.004956 | 1 | unc-2 | 0.083333 | 76 | 6 |
| GO:00708 | response to growth factor | 12/243 | 46/9281 | 1.39971E-09 | 2.29917E-08 | 1.39778E-08 | daf-3/daf-5/daf-7/egl-15/egl-17/lon-2/sma-2/sma-3/tph-1/unc-2/unc-40/egl-4 | 12 | 46 | 0.049383 | 0.004956 | 1 | unc-2 | 0.083333 | 77 | 6 |
| GO:00071 | transmembrane receptor protein serine/threonine kinase signaling pathwa | 10/243 | 43/9281 | 1.12912E-07 | 1.36725E-06 | 8.31224E-07 | daf-3/daf-5/daf-7/lon-2/sma-2/sma-3/tph-1/unc-2/unc-40/egl-4 | 10 | 43 | 0.041152 | 0.004633 | 1 | unc-2 | 0.1 | 78 | 6 |
| GO:00071 | transforming growth factor beta receptor signaling pathway | 10/243 | 35/9281 | 1.29993E-08 | 1.79239E-07 | 1.08969E-07 | daf-3/daf-5/daf-7/lon-2/sma-2/sma-3/tph-1/unc-2/unc-40/egl-4 | 10 | 35 | 0.041152 | 0.003771 | 1 | unc-2 | 0.1 | 79 | 6 |
| GO:00715 | response to transforming growth factor beta | 10/243 | 35/9281 | 1.29993E-08 | 1.79239E-07 | 1.08969E-07 | daf-3/daf-5/daf-7/lon-2/sma-2/sma-3/tph-1/unc-2/unc-40/egl-4 | 10 | 35 | 0.041152 | 0.003771 | 1 | unc-2 | 0.1 | 80 | 6 |
| GO:00715 | cellular response to transforming growth factor beta stimulus | 10/243 | 35/9281 | 1.29993E-08 | 1.79239E-07 | 1.08969E-07 | daf-3/daf-5/daf-7/lon-2/sma-2/sma-3/tph-1/unc-2/unc-40/egl-4 | 10 | 35 | 0.041152 | 0.003771 | 1 | unc-2 | 0.1 | 81 | 6 |
| GO:00230 | negative regulation of signaling | 13/243 | 141/9281 | 7.96238E-05 | 0.00052961 | 0.000321978 | daf-5/egl-10/gar-2/goa-1/gsa-1/ins-18/lon-2/osm-9/rgs-6/unc-2/unc-37/unc-122/egl-4 | 13 | 141 | 0.053498 | 0.015192 | 4 | osm-9/unc-122/unc-2/unc-37 | 0.307692 | 71 | 6 |
| GO:00610 | regulation of dauer larval development | 7/243 | 38/9281 | 4.87967E-05 | 0.000340137 | 0.000206787 | daf-3/daf-5/daf-7/ins-3/ins-4/ins-18/egl-4 | 7 | 38 | 0.028807 | 0.004094 | 2 | ins-3/ins-4 | 0.285714 | 83 | 6 |
| GO:00104 | regulation of signaling receptor activity | 13/243 | 56/9281 | 1.37167E-09 | 2.293E-08 | 1.39403E-08 | daf-7/egl-17/egl-20/ins-3/ins-4/ins-15/ins-25/ins-27/ins-28/ins-30/ins-31/nlp-14/nlp-15 | 13 | 56 | 0.053498 | 0.006034 | 4 | ins-27/ins-28/ins-3/ins-4 | 0.307692 | 84 | 7 |
| GO:00512 | negative regulation of multicellular organismal process | 17/243 | 120/9281 | 1.25449E-08 | 1.79239E-07 | 1.08969E-07 | daf-7/egl-6/egl-20/flp-10/flp-12/flp-16/flp-17/flp-20/goa-1/ins-3/ins-4/jnk-1/lon-2/mec-4/unc-40/rig-6/egl-4 | 17 | 120 | 0.069959 | 0.01293 | 4 | ins-3/ins-4/jnk-1/rig-6 | 0.235294 | 85 | 8 |
| GO:00485 | negative regulation of behavior | 12/243 | 18/9281 | 1.28055E-15 | 4.56409E-14 | 2.77474E-14 | egl-6/egl-21/flp-10/flp-12/flp-16/flp-17/flp-20/mec-4/nlp-1/nlp-14/unc-31/rig-6 | 12 | 18 | 0.049383 | 0.001939 | 2 | nlp-1/rig-6 | 0.166667 | 86 | 8 |
| GO:19010 | negative regulation of oviposition | 3/243 | 3/9281 | 1.77335E-05 | 0.000138424 | 8.41551E-05 | egl-6/flp-10/flp-17 | 3 | 3 | 0.012346 | 0.000323 | 0 | | 0 | 87 | 8 |
| GO:20002 | negative regulation of reproductive process | 7/243 | 33/9281 | 1.84696E-05 | 0.000142404 | 8.6575E-05 | egl-6/flp-10/flp-12/flp-17/flp-20/mec-4/rig-6 | 7 | 33 | 0.028807 | 0.003556 | 1 | rig-6 | 0.142857 | 88 | 8 |
| GO:19024 | regulation of male mating behavior | 4/243 | 8/9281 | 2.95471E-05 | 0.000214671 | 0.00013051 | flp-10/flp-12/flp-20/mec-4 | 4 | 8 | 0.016461 | 0.000862 | 0 | | 0 | 89 | 8 |
| GO:19024 | negative regulation of male mating behavior | 4/243 | 7/9281 | 1.50837E-05 | 0.000119719 | 7.27834E-05 | flp-10/flp-12/flp-20/mec-4 | 4 | 7 | 0.016461 | 0.000754 | 0 | | 0 | 90 | 8 |
| GO:00610 | regulation of turning behavior involved in mating | 4/243 | 7/9281 | 1.50837E-05 | 0.000119719 | 7.27834E-05 | flp-10/flp-12/flp-20/mec-4 | 4 | 7 | 0.016461 | 0.000754 | 0 | | 0 | 91 | 8 |
| GO:00610 | negative regulation of turning behavior involved in mating | 4/243 | 7/9281 | 1.50837E-05 | 0.000119719 | 7.27834E-05 | flp-10/flp-12/flp-20/mec-4 | 4 | 7 | 0.016461 | 0.000754 | 0 | | 0 | 92 | 8 |
| GO:00439 | regulation of multi-organism process | 10/243 | 82/9281 | 5.14344E-05 | 0.000355896 | 0.000216367 | egl-30/flp-10/flp-12/flp-20/gld-1/gpa-12/gsa-1/mec-4/pmk-1/rig-6 | 10 | 82 | 0.041152 | 0.008835 | 1 | rig-6 | 0.1 | 93 | 8 |
| GO:00351 | turning behavior | 6/243 | 12/9281 | 2.45346E-07 | 2.71028E-06 | 1.64772E-06 | bas-1/cat-2/flp-10/flp-12/flp-20/mec-4 | 6 | 12 | 0.024691 | 0.001293 | 0 | | 0 | 94 | 8 |
| GO:00346 | turning behavior involved in mating | 6/243 | 12/9281 | 2.45346E-07 | 2.71028E-06 | 1.64772E-06 | bas-1/cat-2/flp-10/flp-12/flp-20/mec-4 | 6 | 12 | 0.024691 | 0.001293 | 0 | | 0 | 95 | 8 |
| GO:00601 | male mating behavior | 8/243 | 26/9281 | 2.04544E-07 | 2.32761E-06 | 1.41508E-06 | bas-1/cat-2/flp-10/flp-12/flp-20/lov-1/mec-4/ser-1 | 8 | 26 | 0.032922 | 0.002801 | 0 | | 0 | 96 | 8 |
| GO:00076 | mating | 11/243 | 35/9281 | 7.5791E-10 | 1.36352E-08 | 8.28952E-09 | bas-1/cat-2/flp-10/flp-12/flp-20/lov-1/mec-4/rab-3/ser-1/unc-10/unc-103 | 11 | 35 | 0.045267 | 0.003771 | 2 | rab-3/unc-10 | 0.181818 | 97 | 8 |
| GO:00076 | mating behavior | 11/243 | 33/9281 | 3.68244E-10 | 7.21421E-09 | 4.38589E-09 | bas-1/cat-2/flp-10/flp-12/flp-20/lov-1/mec-4/rab-3/ser-1/unc-10/unc-103 | 11 | 33 | 0.045267 | 0.003556 | 2 | rab-3/unc-10 | 0.181818 | 98 | 8 |
| GO:00076 | locomotory behavior | 18/243 | 67/9281 | 5.33067E-14 | 1.70672E-12 | 1.0376E-12 | bas-1/cat-2/egl-21/flp-1/flp-9/flp-10/flp-12/flp-20/lite-1/mec-4/ric-19/unc-2/unc-10/unc-31/unc-98/unc-108/rig- | 18 | 67 | 0.074074 | 0.007219 | 5 | flp-1/lite-1/rig-6/unc-10/unc-2 | 0.277778 | 99 | 8 |
| GO:00319 | locomotion involved in locomotory behavior | 6/243 | 26/9281 | 4.49033E-05 | 0.000315325 | 0.000191702 | bas-1/egl-21/ric-19/unc-31/unc-108/rig-6 | 6 | 26 | 0.024691 | 0.002801 | 1 | rig-6 | 0.166667 | 100 | 8 |
| GO:00903 | regulation of locomotion involved in locomotory behavior | 5/243 | 21/9281 | 0.000170421 | 0.001006631 | 0.000611983 | egl-21/ric-19/unc-31/unc-108/rig-6 | 5 | 21 | 0.020576 | 0.002263 | 1 | rig-6 | 0.2 | 101 | 8 |
| GO:00305 | larval feeding behavior | 4/243 | 11/9281 | 0.00013089 | 0.000818714 | 0.000497739 | daf-7/egl-21/unc-31/egl-4 | 4 | 11 | 0.016461 | 0.001185 | 0 | | 0 | 102 | 8 |
| GO:00356 | cellular response to drug | 8/243 | 18/9281 | 6.87119E-09 | 1.06391E-07 | 6.46805E-08 | egl-21/flp-16/gar-2/goa-1/gpb-2/nlp-1/nlp-14/unc-31 | 8 | 18 | 0.032922 | 0.001939 | 1 | nlp-1 | 0.125 | 103 | 8 |
| GO:00723 | response to anesthetic | 8/243 | 9/9281 | 1.73454E-12 | 4.51011E-11 | 2.74193E-11 | egl-21/flp-16/nlp-1/nlp-14/unc-7/unc-31/unc-79/unc-80 | 8 | 9 | 0.032922 | 0.00097 | 4 | nlp-1/unc-7/unc-79/unc-80 | 0.5 | 104 | 8 |
| GO:00100 | response to carbon dioxide | 5/243 | 6/9281 | 6.93916E-08 | 8.51173E-07 | 5.17472E-07 | egl-21/flp-16/nlp-1/nlp-14/unc-31 | 5 | 6 | 0.020576 | 0.000646 | 1 | nlp-1 | 0.2 | 105 | 8 |
| GO:00712 | cellular response to carbon dioxide | 5/243 | 6/9281 | 6.93916E-08 | 8.51173E-07 | 5.17472E-07 | egl-21/flp-16/nlp-1/nlp-14/unc-31 | 5 | 6 | 0.020576 | 0.000646 | 1 | nlp-1 | 0.2 | 106 | 8 |
| GO:19037 | negative regulation of pharyngeal pumping | 5/243 | 6/9281 | 6.93916E-08 | 8.51173E-07 | 5.17472E-07 | egl-21/flp-16/nlp-1/nlp-14/unc-31 | 5 | 6 | 0.020576 | 0.000646 | 1 | nlp-1 | 0.2 | 107 | 8 |
| GO:19039 | negative regulation of eating behavior | 5/243 | 6/9281 | 6.93916E-08 | 8.51173E-07 | 5.17472E-07 | egl-21/flp-16/nlp-1/nlp-14/unc-31 | 5 | 6 | 0.020576 | 0.000646 | 1 | nlp-1 | 0.2 | 108 | 8 |
| GO:20002 | negative regulation of feeding behavior | 5/243 | 8/9281 | 6.20277E-07 | 6.26579E-06 | 3.8093E-06 | egl-21/flp-16/nlp-1/nlp-14/unc-31 | 5 | 8 | 0.020576 | 0.000862 | 1 | nlp-1 | 0.2 | 109 | 8 |
| GO:00712 | cellular response to inorganic substance | 7/243 | 33/9281 | 1.84696E-05 | 0.000142404 | 8.6575E-05 | dat-1/egl-21/flp-16/jnk-1/nlp-1/nlp-14/unc-31 | 7 | 33 | 0.028807 | 0.003556 | 2 | jnk-1/nlp-1 | 0.285714 | 110 | 8 |
| GO:00156 | ammonium transport | 10/243 | 41/9281 | 6.91344E-08 | 8.51173E-07 | 5.17472E-07 | dat-1/egl-8/egl-21/egl-30/mod-5/snf-6/unc-17/unc-26/unc-29/unc-31 | 10 | 41 | 0.041152 | 0.004418 | 0 | | 0 | 111 | 9 |
| GO:00108 | regulation of hormone levels | 7/243 | 36/9281 | 3.37504E-05 | 0.000242412 | 0.000147375 | egl-8/egl-21/egl-30/snf-6/unc-17/unc-26/unc-29 | 7 | 36 | 0.028807 | 0.003879 | 0 | | 0 | 112 | 9 |
| GO:19013 | acetate ester transport | 7/243 | 19/9281 | 2.98087E-07 | 3.21763E-06 | 1.95617E-06 | egl-8/egl-21/egl-30/snf-6/unc-17/unc-26/unc-29 | 7 | 19 | 0.028807 | 0.002047 | 0 | | 0 | 113 | 9 |
| GO:00158 | acetylcholine transport | 7/243 | 19/9281 | 2.98087E-07 | 3.21763E-06 | 1.95617E-06 | egl-8/egl-21/egl-30/snf-6/unc-17/unc-26/unc-29 | 7 | 19 | 0.028807 | 0.002047 | 0 | | 0 | 114 | 9 |
| GO:00099 | hormone transport | 7/243 | 21/9281 | 6.57702E-07 | 6.60851E-06 | 4.01766E-06 | egl-8/egl-21/egl-30/snf-6/unc-17/unc-26/unc-29 | 7 | 21 | 0.028807 | 0.002263 | 0 | | 0 | 115 | 9 |
| GO:00156 | organic cation transport | 7/243 | 24/9281 | 1.83027E-06 | 1.77302E-05 | 1.07791E-05 | egl-8/egl-21/egl-30/snf-6/unc-17/unc-26/unc-29 | 7 | 24 | 0.028807 | 0.002586 | 0 | | 0 | 116 | 9 |
| GO:00515 | positive regulation of neurotransmitter transport | 6/243 | 15/9281 | 1.24376E-06 | 1.21106E-05 | 7.36265E-06 | egl-8/egl-21/egl-30/unc-18/unc-26/unc-29 | 6 | 15 | 0.024691 | 0.001616 | 0 | | 0 | 117 | 9 |
| GO:00019 | positive regulation of neurotransmitter secretion | 6/243 | 15/9281 | 1.24376E-06 | 1.21106E-05 | 7.36265E-06 | egl-8/egl-21/egl-30/unc-18/unc-26/unc-29 | 6 | 15 | 0.024691 | 0.001616 | 0 | | 0 | 118 | 9 |
| GO:19035 | positive regulation of secretion by cell | 6/243 | 23/9281 | 2.10309E-05 | 0.000159548 | 9.69972E-05 | egl-8/egl-21/egl-30/unc-18/unc-26/unc-29 | 6 | 23 | 0.024691 | 0.002478 | 0 | | 0 | 119 | 9 |
| GO:00510 | positive regulation of secretion | 6/243 | 29/9281 | 8.67395E-05 | 0.000568927 | 0.00034588 | egl-8/egl-21/egl-30/unc-18/unc-26/unc-29 | 6 | 29 | 0.024691 | 0.003125 | 0 | | 0 | 120 | 9 |
| GO:00158 | amine transport | 5/243 | 21/9281 | 0.000170421 | 0.001006631 | 0.000611983 | dat-1/egl-8/egl-21/egl-30/unc-29 | 5 | 21 | 0.020576 | 0.002263 | 0 | | 0 | 121 | 9 |
| GO:00140 | acetylcholine secretion, neurotransmission | 4/243 | 13/9281 | 0.000272095 | 0.001529725 | 0.000929999 | egl-8/egl-21/egl-30/unc-29 | 4 | 13 | 0.016461 | 0.001401 | 0 | | 0 | 122 | 9 |
| GO:00468 | positive regulation of hormone secretion | 4/243 | 12/9281 | 0.000192312 | 0.001107551 | 0.000673337 | egl-8/egl-21/egl-30/unc-29 | 4 | 12 | 0.016461 | 0.001293 | 0 | | 0 | 123 | 9 |
| GO:00519 | positive regulation of amine transport | 4/243 | 11/9281 | 0.00013089 | 0.000818714 | 0.000497739 | egl-8/egl-21/egl-30/unc-29 | 4 | 11 | 0.016461 | 0.001185 | 0 | | 0 | 124 | 9 |
| GO:00140 | positive regulation of acetylcholine secretion, neurotransmission | 4/243 | 11/9281 | 0.00013089 | 0.000818714 | 0.000497739 | egl-8/egl-21/egl-30/unc-29 | 4 | 11 | 0.016461 | 0.001185 | 0 | | 0 | 125 | 9 |
| GO:20010 | positive regulation of response to drug | 4/243 | 11/9281 | 0.00013089 | 0.000818714 | 0.000497739 | egl-8/egl-21/egl-30/unc-29 | 4 | 11 | 0.016461 | 0.001185 | 0 | | 0 | 126 | 9 |
| GO:00140 | regulation of acetylcholine secretion, neurotransmission | 4/243 | 12/9281 | 0.000192312 | 0.001107551 | 0.000673337 | egl-8/egl-21/egl-30/unc-29 | 4 | 12 | 0.016461 | 0.001293 | 0 | | 0 | 127 | 9 |
| GO:00615 | acetylcholine secretion | 4/243 | 13/9281 | 0.000272095 | 0.001529725 | 0.000929999 | egl-8/egl-21/egl-30/unc-29 | 4 | 13 | 0.016461 | 0.001401 | 0 | | 0 | 128 | 9 |
| GO:20010 | regulation of response to drug | 5/243 | 20/9281 | 0.000132657 | 0.000827029 | 0.000502794 | egl-8/egl-21/egl-30/pkc-1/unc-29 | 5 | 20 | 0.020576 | 0.002155 | 0 | | 0 | 129 | 9 |
| GO:00432 | regulation of ion transport | 10/243 | 76/9281 | 2.63322E-05 | 0.000195065 | 0.00011859 | egl-2/egl-8/egl-19/egl-21/egl-30/egl-36/unc-2/unc-29/unc-103/egl-4 | 10 | 76 | 0.041152 | 0.008189 | 1 | unc-2 | 0.1 | 130 | 9 |
| GO:00304 | defecation | 9/243 | 48/9281 | 3.42935E-06 | 3.08478E-05 | 1.8754E-05 | egl-21/egl-30/flr-1/gon-2/gsa-1/unc-10/unc-16/unc-104/rig-6 | 9 | 48 | 0.037037 | 0.005172 | 2 | rig-6/unc-10 | 0.222222 | 131 | 10 |
| GO:00075 | excretion | 9/243 | 50/9281 | 4.89442E-06 | 4.30026E-05 | 2.61435E-05 | egl-21/egl-30/flr-1/gon-2/gsa-1/unc-10/unc-16/unc-104/rig-6 | 9 | 50 | 0.037037 | 0.005387 | 2 | rig-6/unc-10 | 0.222222 | 132 | 10 |
| GO:00455 | positive regulation of cell differentiation | 11/243 | 95/9281 | 3.54617E-05 | 0.000253739 | 0.000154261 | egl-8/egl-20/egl-30/gld-1/gsa-1/lin-39/rab-3/unc-31/unc-89/unc-104/rig-6 | 11 | 95 | 0.045267 | 0.010236 | 3 | rab-3/rig-6/unc-89 | 0.272727 | 133 | 11 |
| GO:00107 | positive regulation of cell development | 10/243 | 80/9281 | 4.14423E-05 | 0.000292106 | 0.000177586 | egl-8/egl-20/egl-30/gld-1/gsa-1/rab-3/unc-31/unc-89/unc-104/rig-6 | 10 | 80 | 0.041152 | 0.00862 | 3 | rab-3/rig-6/unc-89 | 0.3 | 134 | 11 |
| GO:00456 | positive regulation of neuron differentiation | 7/243 | 46/9281 | 0.000173158 | 0.001015824 | 0.000617572 | egl-8/egl-20/egl-30/rab-3/unc-31/unc-104/rig-6 | 7 | 46 | 0.028807 | 0.004956 | 2 | rab-3/rig-6 | 0.285714 | 135 | 11 |
| GO:00109 | positive regulation of neuron projection development | 7/243 | 45/9281 | 0.000150113 | 0.000914719 | 0.000556105 | egl-8/egl-20/egl-30/rab-3/unc-31/unc-104/rig-6 | 7 | 45 | 0.028807 | 0.004849 | 2 | rab-3/rig-6 | 0.285714 | 136 | 11 |
| GO:00519 | positive regulation of nervous system development | 9/243 | 66/9281 | 5.02103E-05 | 0.000348703 | 0.000211995 | egl-8/egl-20/egl-21/egl-30/rab-3/unc-31/unc-40/unc-104/rig-6 | 9 | 66 | 0.037037 | 0.007111 | 2 | rab-3/rig-6 | 0.222222 | 137 | 11 |
| GO:00509 | regulation of chemotaxis | 9/243 | 68/9281 | 6.38752E-05 | 0.000435597 | 0.000264822 | egl-20/egl-30/pkc-1/rab-3/unc-31/unc-40/unc-104/rig-6/egl-4 | 9 | 68 | 0.037037 | 0.007327 | 2 | rab-3/rig-6 | 0.222222 | 138 | 11 |
| GO:00509 | positive regulation of chemotaxis | 7/243 | 28/9281 | 5.72483E-06 | 4.96064E-05 | 3.01583E-05 | egl-20/pkc-1/rab-3/unc-31/unc-104/rig-6/egl-4 | 7 | 28 | 0.028807 | 0.003017 | 2 | rab-3/rig-6 | 0.285714 | 139 | 11 |
| GO:19026 | positive regulation of axon guidance | 5/243 | 18/9281 | 7.65242E-05 | 0.000512604 | 0.000311638 | egl-20/rab-3/unc-31/unc-104/rig-6 | 5 | 18 | 0.020576 | 0.001939 | 2 | rab-3/rig-6 | 0.4 | 140 | 11 |
| GO:00400 | negative regulation of locomotion | 6/243 | 29/9281 | 8.67395E-05 | 0.000568927 | 0.00034588 | eat-16/egl-20/flp-7/unc-32/unc-40/rig-6 | 6 | 29 | 0.024691 | 0.003125 | 2 | rig-6/unc-32 | 0.333333 | 141 | 12 |
| GO:00080 | cell recognition | 7/243 | 34/9281 | 2.2744E-05 | 0.000171169 | 0.000104062 | ced-1/syg-1/unc-4/unc-40/unc-76/syg-2/rig-6 | 7 | 34 | 0.028807 | 0.003663 | 2 | rig-6/unc-4 | 0.285714 | 142 | 13 |
| GO:00080 | neuron recognition | 6/243 | 31/9281 | 0.000128675 | 0.000818714 | 0.000497739 | syg-1/unc-4/unc-40/unc-76/syg-2/rig-6 | 6 | 31 | 0.024691 | 0.00334 | 2 | rig-6/unc-4 | 0.333333 | 143 | 13 |
| GO:00080 | synaptic target recognition | 3/243 | 3/9281 | 1.77335E-05 | 0.000138424 | 8.41551E-05 | syg-1/unc-4/syg-2 | 3 | 3 | 0.012346 | 0.000323 | 1 | unc-4 | 0.333333 | 144 | 13 |
| GO:00901 | negative regulation of transmembrane receptor protein serine/threonine k | 4/243 | 10/9281 | 8.50381E-05 | 0.000561667 | 0.000341467 | daf-5/lon-2/unc-2/egl-4 | 4 | 10 | 0.016461 | 0.001077 | 1 | unc-2 | 0.25 | 145 | 14 |
| GO:00305 | negative regulation of transforming growth factor beta receptor signaling p | 4/243 | 8/9281 | 2.95471E-05 | 0.000214671 | 0.00013051 | daf-5/lon-2/unc-2/egl-4 | 4 | 8 | 0.016461 | 0.000862 | 1 | unc-2 | 0.25 | 146 | 14 |
| GO:19038 | negative regulation of cellular response to transforming growth factor beta | 4/243 | 8/9281 | 2.95471E-05 | 0.000214671 | 0.00013051 | daf-5/lon-2/unc-2/egl-4 | 4 | 8 | 0.016461 | 0.000862 | 1 | unc-2 | 0.25 | 147 | 14 |
| GO:00902 | negative regulation of cellular response to growth factor stimulus | 4/243 | 11/9281 | 0.00013089 | 0.000818714 | 0.000497739 | daf-5/lon-2/unc-2/egl-4 | 4 | 11 | 0.016461 | 0.001185 | 1 | unc-2 | 0.25 | 148 | 14 |
| GO:00170 | regulation of transforming growth factor beta receptor signaling pathway | 6/243 | 20/9281 | 8.62643E-06 | 7.11587E-05 | 4.3261E-05 | daf-5/lon-2/sma-3/tph-1/unc-2/egl-4 | 6 | 20 | 0.024691 | 0.002155 | 1 | unc-2 | 0.166667 | 149 | 14 |
| GO:19038 | regulation of cellular response to transforming growth factor beta stimulus | 6/243 | 20/9281 | 8.62643E-06 | 7.11587E-05 | 4.3261E-05 | daf-5/lon-2/sma-3/tph-1/unc-2/egl-4 | 6 | 20 | 0.024691 | 0.002155 | 1 | unc-2 | 0.166667 | 150 | 14 |
| GO:00902 | regulation of cellular response to growth factor stimulus | 7/243 | 27/9281 | 4.39084E-06 | 3.89404E-05 | 2.36739E-05 | daf-5/lon-2/sma-3/tph-1/unc-2/unc-40/egl-4 | 7 | 27 | 0.028807 | 0.002909 | 1 | unc-2 | 0.142857 | 151 | 14 |
| GO:00900 | regulation of transmembrane receptor protein serine/threonine kinase sig | 8/243 | 28/9281 | 3.88875E-07 | 4.04511E-06 | 2.45923E-06 | daf-5/daf-7/lon-2/sma-3/tph-1/unc-2/unc-40/egl-4 | 8 | 28 | 0.032922 | 0.003017 | 1 | unc-2 | 0.125 | 152 | 14 |
| GO:00355 | determination of left/right asymmetry in nervous system | 4/243 | 11/9281 | 0.00013089 | 0.000818714 | 0.000497739 | egl-19/unc-2/unc-40/rig-6 | 4 | 11 | 0.016461 | 0.001185 | 2 | rig-6/unc-2 | 0.5 | 153 | 14 |
| GO:00069 | regulation of striated muscle contraction | 4/243 | 7/9281 | 1.50837E-05 | 0.000119719 | 7.27834E-05 | egl-19/flp-16/unc-89/unc-104 | 4 | 7 | 0.016461 | 0.000754 | 1 | unc-89 | 0.25 | 154 | 15 |
| GO:00069 | striated muscle contraction | 4/243 | 13/9281 | 0.000272095 | 0.001529725 | 0.000929999 | egl-19/flp-16/unc-89/unc-104 | 4 | 13 | 0.016461 | 0.001401 | 1 | unc-89 | 0.25 | 155 | 15 |
| GO:00072 | octopamine or tyramine signaling pathway | 3/243 | 4/9281 | 6.95579E-05 | 0.000467597 | 0.000284277 | ser-2/octr-1/ser-6 | 3 | 4 | 0.012346 | 0.000431 | 0 | | 0 | 156 | 16 |
| GO:00015 | neurotransmitter uptake | 3/243 | 4/9281 | 6.95579E-05 | 0.000467597 | 0.000284277 | dat-1/mod-5/snf-6 | 3 | 4 | 0.012346 | 0.000431 | 0 | | 0 | 157 | 17 |
| GO:00509 | sensory perception of taste | 5/243 | 15/9281 | 2.86064E-05 | 0.000210262 | 0.000127829 | egl-47/gpa-3/gpc-1/lite-1/egl-4 | 5 | 15 | 0.020576 | 0.001616 | 1 | lite-1 | 0.2 | 158 | 18 |
| GO:00322 | regulation of synaptic transmission, GABAergic | 4/243 | 12/9281 | 0.000192312 | 0.001107551 | 0.000673337 | nca-2/rab-3/unc-75/unc-77 | 4 | 12 | 0.016461 | 0.001293 | 4 | nca-2/rab-3/unc-75/unc-77 | 1 | 159 | 19 |
| GO:00322 | positive regulation of synaptic transmission, GABAergic | 3/243 | 4/9281 | 6.95579E-05 | 0.000467597 | 0.000284277 | nca-2/unc-75/unc-77 | 3 | 4 | 0.012346 | 0.000431 | 3 | nca-2/unc-75/unc-77 | 1 | 160 | 19 |
| GO:00065 | dopamine biosynthetic process from tyrosine | 3/243 | 3/9281 | 1.77335E-05 | 0.000138424 | 8.41551E-05 | bas-1/cat-2/cat-4 | 3 | 3 | 0.012346 | 0.000323 | 0 | | 0 | 161 | 20 |
| GO:00424 | dopamine biosynthetic process | 3/243 | 5/9281 | 0.000170525 | 0.001006631 | 0.000611983 | bas-1/cat-2/cat-4 | 3 | 5 | 0.012346 | 0.000539 | 0 | | 0 | 162 | 20 |
| GO:00424 | dopamine metabolic process | 4/243 | 6/9281 | 6.60026E-06 | 5.59099E-05 | 3.39905E-05 | bas-1/cat-2/cat-4/tbh-1 | 4 | 6 | 0.016461 | 0.000646 | 0 | | 0 | 163 | 20 |
| GO:00097 | catechol-containing compound biosynthetic process | 4/243 | 6/9281 | 6.60026E-06 | 5.59099E-05 | 3.39905E-05 | bas-1/cat-2/cat-4/tbh-1 | 4 | 6 | 0.016461 | 0.000646 | 0 | | 0 | 164 | 20 |
| GO:00065 | catecholamine metabolic process | 4/243 | 6/9281 | 6.60026E-06 | 5.59099E-05 | 3.39905E-05 | bas-1/cat-2/cat-4/tbh-1 | 4 | 6 | 0.016461 | 0.000646 | 0 | | 0 | 165 | 20 |
| GO:00097 | catechol-containing compound metabolic process | 4/243 | 6/9281 | 6.60026E-06 | 5.59099E-05 | 3.39905E-05 | bas-1/cat-2/cat-4/tbh-1 | 4 | 6 | 0.016461 | 0.000646 | 0 | | 0 | 166 | 20 |
| GO:00424 | catecholamine biosynthetic process | 4/243 | 6/9281 | 6.60026E-06 | 5.59099E-05 | 3.39905E-05 | bas-1/cat-2/cat-4/tbh-1 | 4 | 6 | 0.016461 | 0.000646 | 0 | | 0 | 167 | 20 |
| GO:00461 | phenol-containing compound biosynthetic process | 6/243 | 11/9281 | 1.25415E-07 | 1.48068E-06 | 9.0018E-07 | bas-1/cat-2/cat-4/tbh-1/tdc-1/tph-1 | 6 | 11 | 0.024691 | 0.001185 | 0 | | 0 | 168 | 20 |
| GO:00189 | phenol-containing compound metabolic process | 6/243 | 13/9281 | 4.45685E-07 | 4.55081E-06 | 2.76667E-06 | bas-1/cat-2/cat-4/tbh-1/tdc-1/tph-1 | 6 | 13 | 0.024691 | 0.001401 | 0 | | 0 | 169 | 20 |
| GO:00421 | neurotransmitter biosynthetic process | 5/243 | 17/9281 | 5.64669E-05 | 0.000387876 | 0.00023581 | bas-1/cat-2/tbh-1/tdc-1/tph-1 | 5 | 17 | 0.020576 | 0.001832 | 0 | | 0 | 170 | 20 |
| GO:00486 | response to axon injury | 6/243 | 32/9281 | 0.000154935 | 0.000938055 | 0.000570292 | egl-8/egl-30/goa-1/jnk-1/tph-1/unc-34 | 6 | 32 | 0.024691 | 0.003448 | 1 | jnk-1 | 0.166667 | 171 | 21 |
| GO:00311 | axon regeneration | 6/243 | 30/9281 | 0.000106071 | 0.000681526 | 0.000414335 | egl-8/egl-30/goa-1/jnk-1/tph-1/unc-34 | 6 | 30 | 0.024691 | 0.003232 | 1 | jnk-1 | 0.166667 | 172 | 21 |
| GO:00311 | neuron projection regeneration | 6/243 | 30/9281 | 0.000106071 | 0.000681526 | 0.000414335 | egl-8/egl-30/goa-1/jnk-1/tph-1/unc-34 | 6 | 30 | 0.024691 | 0.003232 | 1 | jnk-1 | 0.166667 | 173 | 21 |
| GO:00310 | regeneration | 6/243 | 30/9281 | 0.000106071 | 0.000681526 | 0.000414335 | egl-8/egl-30/goa-1/jnk-1/tph-1/unc-34 | 6 | 30 | 0.024691 | 0.003232 | 1 | jnk-1 | 0.166667 | 174 | 21 |
| GO:00357 | sodium ion transmembrane transport | 10/243 | 66/9281 | 7.3212E-06 | 6.15473E-05 | 3.74178E-05 | del-1/flr-1/mec-4/mec-10/unc-8/unc-105/tmc-1/asic-1/delm-2/egas-2 | 10 | 66 | 0.041152 | 0.007111 | 3 | asic-1/mec-10/unc-8 | 0.3 | 175 | 22 |
| GO:00068 | sodium ion transport | 10/243 | 80/9281 | 4.14423E-05 | 0.000292106 | 0.000177586 | del-1/flr-1/mec-4/mec-10/unc-8/unc-105/tmc-1/asic-1/delm-2/egas-2 | 10 | 80 | 0.041152 | 0.00862 | 3 | asic-1/mec-10/unc-8 | 0.3 | 176 | 22 |
| GO:00718 | adrenergic receptor signaling pathway | 4/243 | 5/9281 | 2.24635E-06 | 2.11112E-05 | 1.28346E-05 | ser-2/ser-5/tyra-2/dop-3 | 4 | 5 | 0.016461 | 0.000539 | 0 | | 0 | 177 | 23 |
| GO:00718 | adenylate cyclase-activating adrenergic receptor signaling pathway | 4/243 | 5/9281 | 2.24635E-06 | 2.11112E-05 | 1.28346E-05 | ser-2/ser-5/tyra-2/dop-3 | 4 | 5 | 0.016461 | 0.000539 | 0 | | 0 | 178 | 23 |
| GO:00199 | cyclic-nucleotide-mediated signaling | 7/243 | 33/9281 | 1.84696E-05 | 0.000142404 | 8.6575E-05 | dop-1/gsa-1/ser-2/ser-5/tyra-2/dop-3/egl-4 | 7 | 33 | 0.028807 | 0.003556 | 0 | | 0 | 179 | 23 |
| GO:00071 | adenylate cyclase-activating G protein-coupled receptor signaling pathway | 6/243 | 17/9281 | 2.9427E-06 | 2.6597E-05 | 1.61697E-05 | dop-1/gsa-1/ser-2/ser-5/tyra-2/dop-3 | 6 | 17 | 0.024691 | 0.001832 | 0 | | 0 | 180 | 23 |
| GO:00199 | cAMP-mediated signaling | 6/243 | 20/9281 | 8.62643E-06 | 7.11587E-05 | 4.3261E-05 | dop-1/gsa-1/ser-2/ser-5/tyra-2/dop-3 | 6 | 20 | 0.024691 | 0.002155 | 0 | | 0 | 181 | 23 |
| GO:00199 | second-messenger-mediated signaling | 10/243 | 64/9281 | 5.50328E-06 | 4.79064E-05 | 2.91248E-05 | dop-1/flp-7/gsa-1/ser-1/ser-2/unc-89/ser-5/tyra-2/dop-3/egl-4 | 10 | 64 | 0.041152 | 0.006896 | 1 | unc-89 | 0.1 | 182 | 23 |
| GO:00072 | dopamine receptor signaling pathway | 11/243 | 15/9281 | 3.94254E-15 | 1.35408E-13 | 8.23217E-14 | dop-1/dop-2/eat-16/egl-8/egl-10/egl-30/goa-1/gpb-2/gsa-1/ser-2/dop-3 | 11 | 15 | 0.045267 | 0.001616 | 0 | | 0 | 183 | 24 |
| GO:00320 | response to food | 7/243 | 10/9281 | 8.69047E-10 | 1.54871E-08 | 9.41539E-09 | cat-2/dop-1/dop-2/eat-16/goa-1/gpb-2/dop-3 | 7 | 10 | 0.028807 | 0.001077 | 0 | | 0 | 184 | 24 |
| GO:00316 | response to nutrient levels | 9/243 | 68/9281 | 6.38752E-05 | 0.000435597 | 0.000264822 | cat-2/dop-1/dop-2/eat-16/goa-1/gpb-2/mod-1/unc-2/dop-3 | 9 | 68 | 0.037037 | 0.007327 | 1 | unc-2 | 0.111111 | 185 | 24 |
| GO:00099 | response to extracellular stimulus | 9/243 | 71/9281 | 9.01639E-05 | 0.000589342 | 0.000358291 | cat-2/dop-1/dop-2/eat-16/goa-1/gpb-2/mod-1/unc-2/dop-3 | 9 | 71 | 0.037037 | 0.00765 | 1 | unc-2 | 0.111111 | 186 | 24 |
| GO:00986 | G protein-coupled serotonin receptor signaling pathway | 10/243 | 16/9281 | 8.79105E-13 | 2.37233E-11 | 1.44226E-11 | dop-1/dop-2/gar-2/mod-1/ser-1/ser-4/ser-7/octr-1/tyra-3/ser-6 | 10 | 16 | 0.041152 | 0.001724 | 0 | | 0 | 187 | 25 |
| GO:00072 | serotonin receptor signaling pathway | 12/243 | 20/9281 | 8.29438E-15 | 2.79787E-13 | 1.70097E-13 | dop-1/dop-2/egl-30/gar-2/gpa-11/mod-1/ser-1/ser-4/ser-7/octr-1/tyra-3/ser-6 | 12 | 20 | 0.049383 | 0.002155 | 0 | | 0 | 188 | 25 |
| GO:00076 | mechanosensory behavior | 10/243 | 28/9281 | 1.09205E-09 | 1.87535E-08 | 1.14012E-08 | cat-2/dop-1/flp-1/mec-4/mec-7/mec-10/mec-12/mec-14/mec-18/unc-86 | 10 | 28 | 0.041152 | 0.003017 | 3 | flp-1/mec-10/mec-14 | 0.3 | 189 | 26 |
| GO:00096 | response to mechanical stimulus | 12/243 | 37/9281 | 8.2142E-11 | 1.78352E-09 | 1.08429E-09 | cat-2/dop-1/flp-1/lov-1/mec-4/mec-7/mec-10/mec-12/mec-14/mec-18/osm-9/unc-86 | 12 | 37 | 0.049383 | 0.003987 | 4 | flp-1/mec-10/mec-14/osm-9 | 0.333333 | 190 | 26 |
| GO:00095 | detection of external stimulus | 10/243 | 35/9281 | 1.29993E-08 | 1.79239E-07 | 1.08969E-07 | gpa-3/lite-1/lov-1/mec-4/mec-7/mec-10/mec-12/mec-14/mec-18/npr-3 | 10 | 35 | 0.041152 | 0.003771 | 3 | lite-1/mec-10/mec-14 | 0.3 | 191 | 26 |
| GO:00095 | detection of abiotic stimulus | 10/243 | 35/9281 | 1.29993E-08 | 1.79239E-07 | 1.08969E-07 | gpa-3/lite-1/lov-1/mec-4/mec-7/mec-10/mec-12/mec-14/mec-18/npr-3 | 10 | 35 | 0.041152 | 0.003771 | 3 | lite-1/mec-10/mec-14 | 0.3 | 192 | 26 |
| GO:00509 | detection of mechanical stimulus | 7/243 | 15/9281 | 4.16466E-08 | 5.46322E-07 | 3.32138E-07 | lov-1/mec-4/mec-7/mec-10/mec-12/mec-14/mec-18 | 7 | 15 | 0.028807 | 0.001616 | 2 | mec-10/mec-14 | 0.285714 | 193 | 26 |
| GO:00509 | detection of mechanical stimulus involved in sensory perception of touch | 6/243 | 12/9281 | 2.45346E-07 | 2.71028E-06 | 1.64772E-06 | mec-4/mec-7/mec-10/mec-12/mec-14/mec-18 | 6 | 12 | 0.024691 | 0.001293 | 2 | mec-10/mec-14 | 0.333333 | 194 | 26 |
| GO:00509 | detection of mechanical stimulus involved in sensory perception | 6/243 | 12/9281 | 2.45346E-07 | 2.71028E-06 | 1.64772E-06 | mec-4/mec-7/mec-10/mec-12/mec-14/mec-18 | 6 | 12 | 0.024691 | 0.001293 | 2 | mec-10/mec-14 | 0.333333 | 195 | 26 |
| GO:00509 | sensory perception of touch | 7/243 | 14/9281 | 2.27164E-08 | 3.06509E-07 | 1.86343E-07 | mec-4/mec-7/mec-10/mec-12/mec-14/mec-18/pkc-1 | 7 | 14 | 0.028807 | 0.001508 | 2 | mec-10/mec-14 | 0.285714 | 196 | 26 |

| | | | | | | | | | | | | | | | | |
|---|---|---|---|---|---|---|---|---|---|---|---|---|---|---|---|---|
| GO:00509 | sensory perception of mechanical stimulus | 7/243 | 17/9281 | 1.20335E-07 | 1.43869E-06 | 8.74655E-07 | mec-4/mec-7/mec-10/mec-12/mec-14/mec-18/pkc-1 | 7 | 17 | 0.028807 | 0.001832 | 2 | mec-10/mec-14 | 0.285714 | 197 | 26 |
| GO:00316 | regulation of neurological system process | 5/243 | 13/9281 | 1.27989E-05 | 0.000103764 | 6.30836E-05 | mec-4/mec-7/mec-10/mec-12/mec-14 | 5 | 13 | 0.020576 | 0.001401 | 2 | mec-10/mec-14 | 0.4 | 198 | 26 |
| GO:00316 | positive regulation of neurological system process | 5/243 | 11/9281 | 4.79669E-06 | 4.23408E-05 | 2.57412E-05 | mec-4/mec-7/mec-10/mec-12/mec-14 | 5 | 11 | 0.020576 | 0.001185 | 2 | mec-10/mec-14 | 0.4 | 199 | 26 |
| GO:19057 | positive regulation of mechanosensory behavior | 5/243 | 10/9281 | 2.67338E-06 | 2.4317E-05 | 1.47836E-05 | mec-4/mec-7/mec-10/mec-12/mec-14 | 5 | 10 | 0.020576 | 0.001077 | 2 | mec-10/mec-14 | 0.4 | 200 | 26 |
| GO:19057 | regulation of mechanosensory behavior | 5/243 | 10/9281 | 2.67338E-06 | 2.4317E-05 | 1.47836E-05 | mec-4/mec-7/mec-10/mec-12/mec-14 | 5 | 10 | 0.020576 | 0.001077 | 2 | mec-10/mec-14 | 0.4 | 201 | 26 |
| GO:19057 | positive regulation of detection of mechanical stimulus involved in sensory | 5/243 | 10/9281 | 2.67338E-06 | 2.4317E-05 | 1.47836E-05 | mec-4/mec-7/mec-10/mec-12/mec-14 | 5 | 10 | 0.020576 | 0.001077 | 2 | mec-10/mec-14 | 0.4 | 202 | 26 |
| GO:00519 | regulation of sensory perception | 5/243 | 10/9281 | 2.67338E-06 | 2.4317E-05 | 1.47836E-05 | mec-4/mec-7/mec-10/mec-12/mec-14 | 5 | 10 | 0.020576 | 0.001077 | 2 | mec-10/mec-14 | 0.4 | 203 | 26 |
| GO:19057 | regulation of detection of mechanical stimulus involved in sensory percept | 5/243 | 10/9281 | 2.67338E-06 | 2.4317E-05 | 1.47836E-05 | mec-4/mec-7/mec-10/mec-12/mec-14 | 5 | 10 | 0.020576 | 0.001077 | 2 | mec-10/mec-14 | 0.4 | 204 | 26 |
| GO:00485 | positive regulation of behavior | 9/243 | 56/9281 | 1.29063E-05 | 0.000104188 | 6.33415E-05 | egl-21/mec-4/mec-7/mec-10/mec-12/mec-14/unc-104/ser-5/egl-4 | 9 | 56 | 0.037037 | 0.006034 | 2 | mec-10/mec-14 | 0.222222 | 205 | 26 |
| GO:00321 | positive regulation of response to external stimulus | 16/243 | 64/9281 | 4.75327E-12 | 1.19719E-10 | 7.27833E-11 | egl-8/egl-20/egl-30/gpa-12/mec-4/mec-7/mec-10/mec-12/mec-14/pkc-1/pmk-1/rab-3/unc-31/unc-104/rig-6/egl- | 16 | 64 | 0.065844 | 0.006896 | 4 | mec-10/mec-14/rab-3/rig-6 | 0.25 | 206 | 26 |
| GO:00321 | regulation of response to external stimulus | 19/243 | 115/9281 | 1.12283E-10 | 2.41025E-09 | 1.46532E-09 | egl-8/egl-20/egl-30/goa-1/gpa-12/jnk-1/mec-4/mec-7/mec-10/mec-12/mec-14/pkc-1/pmk-1/rab-3/unc-31/unc-4 | 19 | 115 | 0.078189 | 0.012391 | 5 | jnk-1/mec-10/mec-14/rab-3/rig-6 | 0.263158 | 207 | 26 |
| GO:00109 | regulation of cell death | 11/243 | 119/9281 | 0.000275738 | 0.001545608 | 0.000939655 | ced-1/daf-7/dat-1/egl-1/jnk-1/pmk-1/unc-3/unc-26/unc-32/unc-104/unc-116 | 11 | 119 | 0.045267 | 0.012822 | 3 | jnk-1/unc-3/unc-32 | 0.272727 | 208 | 27 |
| GO:00109 | positive regulation of cell death | 9/243 | 59/9281 | 1.99917E-05 | 0.000152275 | 9.2576E-05 | ced-1/dat-1/egl-1/jnk-1/unc-3/unc-26/unc-32/unc-104/unc-116 | 9 | 59 | 0.037037 | 0.006357 | 3 | jnk-1/unc-3/unc-32 | 0.333333 | 209 | 27 |
| GO:00516 | establishment of organelle localization | 16/243 | 184/9281 | 2.49216E-05 | 0.000186813 | 0.000113573 | ced-1/dnc-1/egl-20/gpa-16/jnk-1/pkc-1/sng-1/tom-1/unc-4/unc-10/unc-16/unc-18/unc-26/unc-31/unc-104/unc- | 16 | 184 | 0.065844 | 0.019825 | 4 | jnk-1/sng-1/unc-10/unc-4 | 0.25 | 210 | 27 |
| GO:00516 | establishment of vesicle localization | 12/243 | 73/9281 | 3.53358E-07 | 3.7083E-06 | 2.25447E-06 | jnk-1/pkc-1/sng-1/tom-1/unc-4/unc-10/unc-16/unc-18/unc-26/unc-31/unc-104/unc-116 | 12 | 73 | 0.049383 | 0.007866 | 4 | jnk-1/sng-1/unc-10/unc-4 | 0.333333 | 211 | 27 |
| GO:00516 | vesicle localization | 13/243 | 77/9281 | 8.18009E-08 | 9.96915E-07 | 6.06077E-07 | jnk-1/pkc-1/sng-1/tom-1/unc-4/unc-10/unc-14/unc-16/unc-18/unc-26/unc-31/unc-104/unc-116 | 13 | 77 | 0.053498 | 0.008297 | 4 | jnk-1/sng-1/unc-10/unc-4 | 0.307692 | 212 | 27 |
| GO:00974 | synaptic vesicle localization | 12/243 | 54/9281 | 1.02462E-08 | 1.47748E-07 | 8.98238E-08 | pkc-1/sng-1/tom-1/unc-4/unc-10/unc-14/unc-16/unc-18/unc-26/unc-31/unc-104/unc-116 | 12 | 54 | 0.049383 | 0.005818 | 3 | sng-1/unc-10/unc-4 | 0.25 | 213 | 27 |
| GO:00990 | vesicle-mediated transport in synapse | 11/243 | 56/9281 | 1.66352E-07 | 1.92784E-06 | 1.17203E-06 | pkc-1/sng-1/tom-1/unc-4/unc-10/unc-16/unc-18/unc-26/unc-31/unc-104/unc-116 | 11 | 56 | 0.045267 | 0.006034 | 3 | sng-1/unc-10/unc-4 | 0.272727 | 214 | 27 |
| GO:00974 | establishment of synaptic vesicle localization | 11/243 | 50/9281 | 4.79447E-08 | 6.07835E-07 | 3.69535E-07 | pkc-1/sng-1/tom-1/unc-4/unc-10/unc-16/unc-18/unc-26/unc-31/unc-104/unc-116 | 11 | 50 | 0.045267 | 0.005387 | 3 | sng-1/unc-10/unc-4 | 0.272727 | 215 | 27 |
| GO:00484 | synaptic vesicle transport | 11/243 | 50/9281 | 4.79447E-08 | 6.07835E-07 | 3.69535E-07 | pkc-1/sng-1/tom-1/unc-4/unc-10/unc-16/unc-18/unc-26/unc-31/unc-104/unc-116 | 11 | 50 | 0.045267 | 0.005387 | 3 | sng-1/unc-10/unc-4 | 0.272727 | 216 | 27 |
| GO:00400 | positive regulation of multicellular organism growth | 7/243 | 49/9281 | 0.000259956 | 0.001474646 | 0.000896514 | sma-2/sma-3/unc-10/unc-26/unc-40/unc-44/unc-104 | 7 | 49 | 0.028807 | 0.00528 | 1 | unc-10 | 0.142857 | 217 | 28 |
| GO:00486 | positive regulation of developmental growth | 9/243 | 72/9281 | 0.000100728 | 0.000653869 | 0.000397521 | egl-21/sma-2/sma-3/unc-10/unc-26/unc-40/unc-44/unc-104/rig-6 | 9 | 72 | 0.037037 | 0.007758 | 2 | rig-6/unc-10 | 0.222222 | 218 | 28 |
| GO:00352 | multicellular organism growth | 10/243 | 99/9281 | 0.000253901 | 0.001448998 | 0.000880921 | lon-2/sma-2/sma-3/unc-10/unc-26/unc-40/unc-44/unc-104/rig-6/egl-4 | 10 | 99 | 0.041152 | 0.010667 | 2 | rig-6/unc-10 | 0.2 | 219 | 28 |
| GO:00400 | regulation of multicellular organism growth | 10/243 | 88/9281 | 9.45932E-05 | 0.00061616 | 0.000374596 | lon-2/sma-2/sma-3/unc-10/unc-26/unc-40/unc-44/unc-104/rig-6/egl-4 | 10 | 88 | 0.041152 | 0.009482 | 2 | rig-6/unc-10 | 0.2 | 220 | 28 |
| GO:00486 | regulation of developmental growth | 14/243 | 145/9281 | 2.52453E-05 | 0.000188492 | 0.000114594 | egl-21/ins-4/lon-2/sma-2/sma-3/unc-10/unc-26/unc-40/unc-44/unc-69/unc-76/unc-104/rig-6/egl-4 | 14 | 145 | 0.057613 | 0.015623 | 3 | ins-4/rig-6/unc-10 | 0.214286 | 221 | 28 |
| GO:00400 | regulation of growth | 14/243 | 172/9281 | 0.000163484 | 0.000980387 | 0.000596028 | egl-21/ins-4/lon-2/sma-2/sma-3/unc-10/unc-26/unc-40/unc-44/unc-69/unc-76/unc-104/rig-6/egl-4 | 14 | 172 | 0.057613 | 0.018532 | 3 | ins-4/rig-6/unc-10 | 0.214286 | 222 | 28 |
| GO:00485 | developmental growth | 16/243 | 185/9281 | 2.66522E-05 | 0.000196664 | 0.000119562 | egl-21/ins-4/lon-2/sma-2/sma-3/syg-1/unc-10/unc-26/unc-40/unc-44/unc-69/unc-76/unc-104/syg-2/rig-6/egl-4 | 16 | 185 | 0.065844 | 0.019933 | 3 | ins-4/rig-6/unc-10 | 0.1875 | 223 | 28 |
| GO:00074 | synapse assembly | 7/243 | 45/9281 | 0.000150113 | 0.000914719 | 0.000556105 | egl-21/ins-4/syg-1/unc-4/unc-40/unc-104/syg-2 | 7 | 45 | 0.028807 | 0.004849 | 2 | ins-4/unc-4 | 0.285714 | 224 | 28 |
| GO:00508 | synapse organization | 10/243 | 81/9281 | 4.62083E-05 | 0.000323287 | 0.000196543 | egl-1/egl-21/ins-4/syg-1/unc-4/unc-40/unc-69/unc-76/unc-104/syg-2 | 10 | 81 | 0.041152 | 0.008728 | 2 | ins-4/unc-4 | 0.2 | 225 | 28 |
| GO:00508 | regulation of synapse organization | 7/243 | 46/9281 | 0.000173158 | 0.001015824 | 0.000617572 | egl-1/egl-21/ins-4/unc-40/unc-69/unc-76/unc-104 | 7 | 46 | 0.028807 | 0.004956 | 1 | ins-4 | 0.142857 | 226 | 28 |
| GO:00507 | regulation of axonogenesis | 10/243 | 72/9281 | 1.62134E-05 | 0.000128147 | 7.79071E-05 | egl-20/rab-3/unc-4/unc-31/unc-37/unc-40/unc-69/unc-76/unc-104/rig-6 | 10 | 72 | 0.041152 | 0.007758 | 4 | rab-3/rig-6/unc-37/unc-4 | 0.4 | 228 | 28 |
| GO:00519 | regulation of nervous system development | 21/243 | 146/9281 | 1.69021E-10 | 3.54756E-09 | 2.15675E-09 | dnc-1/egl-8/egl-20/egl-21/egl-30/egl-43/goa-1/ins-4/jnk-1/rab-3/unc-3/unc-4/unc-31/unc-37/unc-40/unc-69/unc | 21 | 146 | 0.08642 | 0.015731 | 8 | ins-4/jnk-1/rab-3/rig-6/unc-3/unc-37/unc-4/unc-75 | 0.380952 | 233 | 28 |
| GO:00507 | regulation of neurogenesis | 19/243 | 126/9281 | 5.69592E-10 | 1.05486E-08 | 6.41306E-09 | dnc-1/egl-8/egl-20/egl-30/egl-43/goa-1/jnk-1/rab-3/unc-3/unc-4/unc-31/unc-37/unc-40/unc-69/unc-75/unc-76/ | 19 | 126 | 0.078189 | 0.013576 | 7 | jnk-1/rab-3/rig-6/unc-3/unc-37/unc-4/unc-75 | 0.368421 | 234 | 28 |
| GO:00456 | regulation of neuron differentiation | 19/243 | 116/9281 | 1.3118E-10 | 2.78427E-09 | 1.6927E-09 | dnc-1/egl-8/egl-20/egl-30/egl-43/goa-1/jnk-1/rab-3/unc-3/unc-4/unc-31/unc-37/unc-40/unc-69/unc-75/unc-76/ | 19 | 116 | 0.078189 | 0.012499 | 7 | jnk-1/rab-3/rig-6/unc-3/unc-37/unc-4/unc-75 | 0.368421 | 235 | 28 |
| GO:00226 | regulation of anatomical structure morphogenesis | 17/243 | 151/9281 | 3.89735E-07 | 4.04511E-06 | 2.45923E-06 | dnc-1/egl-20/lin-39/lon-2/rab-3/sma-2/sma-3/unc-4/unc-31/unc-37/unc-40/unc-69/unc-76/unc-89/unc-104/unc | 17 | 151 | 0.069959 | 0.01627 | 5 | rab-3/rig-6/unc-37/unc-4/unc-89 | 0.294118 | 231 | 28 |
| GO:00602 | regulation of cell development | 22/243 | 167/9281 | 3.48331E-10 | 6.92628E-09 | 4.21085E-09 | dnc-1/egl-8/egl-20/egl-30/egl-43/gld-1/goa-1/gsa-1/jnk-1/rab-3/unc-3/unc-4/unc-31/unc-37/unc-40/unc-69/unc | 22 | 167 | 0.090535 | 0.017994 | 8 | jnk-1/rab-3/rig-6/unc-3/unc-37/unc-4/unc-75/unc-89 | 0.363636 | 232 | 28 |
| GO:00109 | regulation of neuron projection development | 17/243 | 104/9281 | 1.31851E-09 | 2.22382E-08 | 1.35197E-08 | dnc-1/egl-8/egl-20/egl-30/goa-1/jnk-1/rab-3/unc-3/unc-4/unc-31/unc-37/unc-40/unc-69/unc-76/unc-104/unc-11 | 17 | 104 | 0.069959 | 0.011206 | 6 | jnk-1/rab-3/rig-6/unc-3/unc-37/unc-4 | 0.352941 | 236 | 28 |
| GO:00508 | regulation of synapse structure or activity | 9/243 | 53/9281 | 8.08436E-06 | 6.75723E-05 | 4.10807E-05 | egl-1/egl-21/ins-4/unc-4/unc-37/unc-40/unc-69/unc-76/unc-104 | 9 | 53 | 0.037037 | 0.005711 | 3 | ins-4/unc-37/unc-4 | 0.333333 | 227 | 28 |
| GO:00226 | regulation of cell morphogenesis | 12/243 | 100/9281 | 1.07995E-05 | 8.79325E-05 | 5.34587E-05 | dnc-1/egl-20/rab-3/unc-4/unc-31/unc-37/unc-40/unc-69/unc-76/unc-104/unc-116/rig-6 | 12 | 100 | 0.049383 | 0.010775 | 4 | rab-3/rig-6/unc-37/unc-4 | 0.333333 | 229 | 28 |
| GO:00107 | regulation of cell morphogenesis involved in differentiation | 12/243 | 86/9281 | 2.17401E-06 | 2.08462E-05 | 1.26735E-05 | dnc-1/egl-20/rab-3/unc-4/unc-31/unc-37/unc-40/unc-69/unc-76/unc-104/unc-116/rig-6 | 12 | 86 | 0.049383 | 0.009266 | 4 | rab-3/rig-6/unc-37/unc-4 | 0.333333 | 230 | 28 |
| GO:01200 | regulation of plasma membrane bounded cell projection organization | 18/243 | 121/9281 | 2.06117E-09 | 3.32783E-08 | 2.02316E-08 | dnc-1/egl-8/egl-20/egl-30/goa-1/jnk-1/rab-3/unc-3/unc-4/unc-31/unc-37/unc-40/unc-69/unc-76/unc-104/unc-11 | 18 | 121 | 0.074074 | 0.013037 | 6 | jnk-1/rab-3/rig-6/unc-3/unc-37/unc-4 | 0.333333 | 237 | 28 |
| GO:00400 | positive regulation of locomotion | 22/243 | 119/9281 | 3.29752E-13 | 9.4379E-12 | 5.73779E-12 | ced-1/egl-8/egl-10/egl-19/egl-20/egl-21/egl-30/nca-2/pkc-1/rab-3/snf-6/snf-10/unc-31/unc-34/unc-40/unc-77/ur | 22 | 119 | 0.090535 | 0.012822 | 7 | nca-2/rab-3/rig-6/unc-77/unc-79/unc-80/unc-89 | 0.318182 | 249 | 28 |
| GO:00615 | axon development | 22/243 | 178/9281 | 1.20449E-09 | 2.04981E-08 | 1.24619E-08 | egl-8/egl-20/egl-30/egl-46/goa-1/jnk-1/rab-3/syg-1/tph-1/unc-4/unc-30/unc-31/unc-34/unc-37/unc-40/unc-44/u | 22 | 178 | 0.090535 | 0.019179 | 6 | jnk-1/rab-3/rig-6/unc-30/unc-37/unc-4 | 0.272727 | 239 | 28 |
| GO:00074 | axonogenesis | 17/243 | 159/9281 | 8.16521E-07 | 8.11793E-06 | 4.93531E-06 | egl-20/egl-46/rab-3/syg-1/unc-4/unc-30/unc-31/unc-34/unc-37/unc-40/unc-44/unc-69/unc-76/unc-104/unc-116, | 17 | 159 | 0.069959 | 0.017132 | 5 | rab-3/rig-6/unc-30/unc-37/unc-4 | 0.294118 | 240 | 28 |
| GO:00486 | cell morphogenesis involved in neuron differentiation | 18/243 | 170/9281 | 4.44359E-07 | 4.55081E-06 | 2.76667E-06 | dnc-1/egl-20/egl-46/rab-3/syg-1/unc-4/unc-30/unc-31/unc-34/unc-37/unc-40/unc-44/unc-69/unc-76/unc-104/ui | 18 | 170 | 0.074074 | 0.018317 | 5 | rab-3/rig-6/unc-30/unc-37/unc-4 | 0.277778 | 241 | 28 |
| GO:00009 | cell morphogenesis involved in differentiation | 18/243 | 182/9281 | 1.22125E-06 | 1.20153E-05 | 7.30471E-06 | dnc-1/egl-20/egl-46/rab-3/syg-1/unc-4/unc-30/unc-31/unc-34/unc-37/unc-40/unc-44/unc-69/unc-76/unc-104/ui | 18 | 182 | 0.074074 | 0.01961 | 5 | rab-3/rig-6/unc-30/unc-37/unc-4 | 0.277778 | 242 | 28 |
| GO:00329 | cell part morphogenesis | 19/243 | 182/9281 | 2.63019E-07 | 2.88862E-06 | 1.75614E-06 | dnc-1/egl-20/egl-46/rab-3/syg-1/unc-4/unc-30/unc-31/unc-34/unc-37/unc-40/unc-44/unc-69/unc-76/unc-104/ui | 19 | 182 | 0.078189 | 0.01961 | 5 | rab-3/rig-6/unc-30/unc-37/unc-4 | 0.263158 | 243 | 28 |
| GO:00488 | cell projection morphogenesis | 19/243 | 178/9281 | 1.85072E-07 | 2.12064E-06 | 1.28925E-06 | dnc-1/egl-20/egl-46/rab-3/syg-1/unc-4/unc-30/unc-31/unc-34/unc-37/unc-40/unc-44/unc-69/unc-76/unc-104/ui | 19 | 178 | 0.078189 | 0.019179 | 5 | rab-3/rig-6/unc-30/unc-37/unc-4 | 0.263158 | 244 | 28 |
| GO:01200 | plasma membrane bounded cell projection morphogenesis | 19/243 | 176/9281 | 1.54629E-07 | 1.80305E-06 | 1.09617E-06 | dnc-1/egl-20/egl-46/rab-3/syg-1/unc-4/unc-30/unc-31/unc-34/unc-37/unc-40/unc-44/unc-69/unc-76/unc-104/ui | 19 | 176 | 0.078189 | 0.018963 | 5 | rab-3/rig-6/unc-30/unc-37/unc-4 | 0.263158 | 245 | 28 |
| GO:00488 | neuron projection morphogenesis | 19/243 | 176/9281 | 1.54629E-07 | 1.80305E-06 | 1.09617E-06 | dnc-1/egl-20/egl-46/rab-3/syg-1/unc-4/unc-30/unc-31/unc-34/unc-37/unc-40/unc-44/unc-69/unc-76/unc-104/ui | 19 | 176 | 0.078189 | 0.018963 | 5 | rab-3/rig-6/unc-30/unc-37/unc-4 | 0.263158 | 246 | 28 |
| GO:00512 | regulation of cellular component movement | 12/243 | 131/9281 | 0.000159343 | 0.000958594 | 0.000582779 | ced-1/egl-17/egl-20/egl-27/egl-43/lon-2/rab-3/snf-10/unc-31/unc-40/unc-104/rig-6 | 12 | 131 | 0.049383 | 0.014115 | 2 | rab-3/rig-6 | 0.166667 | 247 | 28 |
| GO:00512 | positive regulation of cellular component movement | 7/243 | 49/9281 | 0.000259956 | 0.001474646 | 0.000896514 | ced-1/egl-20/rab-3/snf-10/unc-31/unc-104/rig-6 | 7 | 49 | 0.028807 | 0.00528 | 2 | rab-3/rig-6 | 0.285714 | 248 | 28 |
| GO:00313 | regulation of cell projection organization | 19/243 | 126/9281 | 5.69592E-10 | 1.05486E-08 | 6.41306E-09 | dnc-1/egl-8/egl-15/egl-20/egl-30/goa-1/jnk-1/rab-3/unc-3/unc-4/unc-31/unc-37/unc-40/unc-69/unc-76/unc-104 | 19 | 126 | 0.078189 | 0.013576 | 6 | jnk-1/rab-3/rig-6/unc-3/unc-37/unc-4 | 0.315789 | 238 | 28 |
| GO:00072 | neuropeptide signaling pathway | 33/243 | 129/9281 | 4.5611E-24 | 3.74605E-22 | 2.27742E-22 | flp-1/flp-3/flp-6/flp-7/flp-9/flp-11/flp-13/flp-16/flp-17/flp-18/flp-19/flp-21/nlp-2/nlp-3/nlp-5/nlp-8/nlp-12/nlp-1 | 33 | 129 | 0.135802 | 0.013899 | 2 | flp-1/nlp-3 | 0.060606 | 250 | 29 |
