## Supplemental Table 1(7) for "Rebalancing the motor circuit restores movement in a *Caenorhabditis elegans* model for TDP-43-toxicity"

| id | description | gene ratio | bg ratio | qvalue | gene id | hits count | hits list | fold change | cluster id |
|---|---|---|---|---|---|---|---|---|---|
| GO:0032228 | regulation of synaptic transmission, GABAergic | 4/243 | 12/9281 | 0.00067 | nca-2/rab-3/unc-75/unc-77 | 4 | nca-2/rab-3/unc-75/unc-77 | 5.92683 | 19 |
| GO:0032230 | positive regulation of synaptic transmission, GABAergic | 3/243 | 4/9281 | 0.00028 | nca-2/unc-75/unc-77 | 3 | nca-2/unc-75/unc-77 | 5.92683 | 19 |
| GO:0032224 | positive regulation of synaptic transmission, cholinergic | 8/243 | 18/9281 | 6.5E-08 | egl-8/egl-21/egl-30/nca-2/unc-10/unc-29/unc-75/unc-77 | 4 | nca-2/unc-10/unc-75/unc-77 | 2.96341 | 3 |
| GO:0072347 | response to anesthetic | 8/243 | 9/9281 | 2.7E-11 | egl-21/flp-16/nlp-1/nlp-14/unc-7/unc-31/unc-79/unc-80 | 4 | nlp-1/unc-7/unc-79/unc-80 | 2.96341 | 8 |
| GO:0032222 | regulation of synaptic transmission, cholinergic | 11/243 | 25/9281 | 1.5E-10 | egl-8/egl-21/egl-30/gar-2/goa-1/nca-2/unc-10/unc-29/unc-75/unc-77/unc-122 | 5 | nca-2/unc-10/unc-122/unc-75/unc-77 | 2.69401 | 3 |
| GO:0007271 | synaptic transmission, cholinergic | 17/243 | 43/9281 | 4.1E-15 | egl-8/egl-21/egl-30/gar-2/goa-1/nca-2/rab-3/unc-10/unc-17/unc-18/unc-29/unc-32/unc-63/unc-75/unc-77/unc-104/unc-122 | 7 | nca-2/rab-3/unc-10/unc-122/unc-32/unc-75/unc-77 | 2.44046 | 3 |
| GO:0050806 | positive regulation of synaptic transmission | 10/243 | 32/9281 | 4.7E-08 | egl-8/egl-21/egl-30/nca-2/unc-10/unc-18/unc-26/unc-29/unc-75/unc-77 | 4 | nca-2/unc-10/unc-75/unc-77 | 2.37073 | 3 |
| GO:0050770 | regulation of axonogenesis | 10/243 | 72/9281 | 7.8E-05 | egl-20/rab-3/unc-4/unc-31/unc-37/unc-40/unc-69/unc-76/unc-104/rig-6 | 4 | rab-3/rig-6/unc-37/unc-4 | 2.37073 | 28 |
| GO:0051960 | regulation of nervous system development | 21/243 | 146/9281 | 2.2E-09 | dnc-1/egl-8/egl-20/egl-21/egl-30/egl-43/goa-1/ins-4/jnk-1/rab-3/unc-3/unc-4/unc-31/unc-37/unc-40/unc-69/unc-75/unc-76/unc-104/unc-116/rig-6 | 8 | ins-4/jnk-1/rab-3/rig-6/unc-3/unc-37/unc-4/unc-75 | 2.25784 | 28 |
| GO:0050767 | regulation of neurogenesis | 19/243 | 126/9281 | 6.4E-09 | dnc-1/egl-8/egl-20/egl-30/egl-43/goa-1/jnk-1/rab-3/unc-3/unc-4/unc-31/unc-37/unc-40/unc-69/unc-75/unc-76/unc-104/unc-116/rig-6 | 7 | jnk-1/rab-3/rig-6/unc-3/unc-37/unc-4/unc-75 | 2.18357 | 28 |
| GO:0045664 | regulation of neuron differentiation | 19/243 | 116/9281 | 1.7E-09 | dnc-1/egl-8/egl-20/egl-30/egl-43/goa-1/jnk-1/rab-3/unc-3/unc-4/unc-31/unc-37/unc-40/unc-69/unc-75/unc-76/unc-104/unc-116/rig-6 | 7 | jnk-1/rab-3/rig-6/unc-3/unc-37/unc-4/unc-75 | 2.18357 | 28 |
| GO:0060284 | regulation of cell development | 22/243 | 167/9281 | 4.2E-09 | dnc-1/egl-8/egl-20/egl-30/egl-43/gld-1/goa-1/gsa-1/jnk-1/rab-3/unc-3/unc-4/unc-31/unc-37/unc-40/unc-69/unc-75/unc-76/unc-89/unc-104/unc-116/rig-6 | 8 | jnk-1/rab-3/rig-6/unc-3/unc-37/unc-4/unc-75/unc-89 | 2.15521 | 28 |
| GO:0010975 | regulation of neuron projection development | 17/243 | 104/9281 | 1.4E-08 | dnc-1/egl-8/egl-20/egl-30/goa-1/jnk-1/rab-3/unc-3/unc-4/unc-31/unc-37/unc-40/unc-69/unc-76/unc-104/unc-116/rig-6 | 6 | jnk-1/rab-3/rig-6/unc-3/unc-37/unc-4 | 2.09182 | 28 |
