## Supplemental Table 1(1) for "Rebalancing the motor circuit restores movement in a *Caenorhabditis elegans* model for TDP-43-toxicity"

| ID | temperature | strain | motion_mode_ | motion_mode_ | curvature_std_ | d_curvature_n | ang_vel_tail_ti | curvature_std_ | rel_to_head_b | width_tail_bas | minor_axis_w_ | d_rel_to_head | d_rel_to_body | rel_to_head_b | rel_to_body_r |
|---|---|---|---|---|---|---|---|---|---|---|---|---|---|---|---|
| 1 | 20 | hTDP-43 | 0.01444316 | 0.86332697 | 0.0045343 | 0.00662882 | 0.4835631 | 0.00177411 | -0.16467515 | 56.2257423 | 87.2518921 | 2.30032873 | 175.487457 | 0.27226293 | -42.8442726 |
| 2 | 20 | hTDP-43 | 0.017493 | 0.7445433 | 0.00521845 | 0.00733406 | 0.40803891 | 0.00261897 | -0.35115117 | 52.9379692 | 81.8096619 | 2.72409916 | 267.770783 | 0.42094535 | -28.7570114 |
| 3 | 20 | hTDP-43 | 0.00932794 | 0.87290048 | 0.00456011 | 0.00750887 | 0.40785611 | 0.00203064 | -0.1539321 | 56.8776054 | 88.2296753 | 2.00562668 | 211.354385 | 0.21756318 | -27.2551003 |
| 4 | 20 | hTDP-43 | 0.0108797 | 0.93087364 | 0.00445307 | 0.00553024 | 0.40169776 | 0.00178639 | -0.31771326 | 57.1667061 | 113.903877 | 2.11211252 | 174.225281 | 0.35546923 | -28.3319702 |
| 5 | 20 | hTDP-43 | 0.00740543 | 0.86005342 | 0.00421096 | 0.00611 | 0.47401047 | 0.0018017 | -0.28132713 | 56.6306801 | 108.783447 | 2.2491169 | 164.88295 | 0.34977397 | -43.6454811 |
| 6 | 20 | Control | 0.02710027 | 0.37125007 | 0.00439144 | 0.00839339 | 0.54031014 | 0.00241026 | -0.36610913 | 58.0646248 | 99.0361023 | 1.77348602 | 267.671387 | 0.3815138 | -34.4552345 |
| 7 | 20 | Control | 0.02177875 | 0.4511689 | 0.00389124 | 0.00779523 | 0.50133127 | 0.00063986 | -0.1941441 | 60.1346817 | 96.3233566 | 1.94200003 | 302.964111 | 0.28672591 | -29.0604897 |
| 8 | 20 | Control | 0.0169452 | 0.14593523 | 0.00455733 | 0.01047294 | 0.62107009 | 0.00239658 | -0.51365513 | 54.3745384 | 85.6060028 | 2.11329484 | 250.666367 | 0.4924742 | -29.7435379 |
| 9 | 20 | Control | 0.01597373 | 0.20746493 | 0.00415585 | 0.01014847 | 0.6720866 | 0.00230704 | -0.63407046 | 57.4022713 | 94.71241 | 2.07082367 | 310.018311 | 0.48941448 | -39.4850655 |
| 10 | 20 | Control | 0.01952172 | 0.16233216 | 0.00445104 | 0.01228079 | 0.69607127 | 0.00249467 | -0.64417076 | 55.276413 | 85.1674347 | 2.36798716 | 272.145813 | 0.54082227 | -42.75354 |
| 11 | 25 | hTDP-43 | 0.01576167 | 0.79421924 | 0.00379569 | 0.00452445 | 0.34279198 | 0.00106823 | -0.25417802 | 59.2138062 | 103.500191 | 1.71987832 | 152.701355 | 0.26655191 | -27.0940685 |
| 12 | 25 | hTDP-43 | 0.01265317 | 0.74212111 | 0.00408836 | 0.0051245 | 0.32139781 | 0.00142453 | -0.23984133 | 52.7531891 | 86.880806 | 1.65766501 | 146.64444 | 0.27887619 | -32.3685684 |
| 13 | 25 | hTDP-43 | 0.00666104 | 0.95091922 | 0.00506441 | 0.00682466 | 0.3740755 | 0.00267272 | -0.36420006 | 54.7979126 | 146.238281 | 2.10301781 | 206.91214 | 0.35209835 | -55.1193733 |
| 14 | 25 | hTDP-43 | 0.015317 | 0.90896852 | 0.00398619 | 0.00521128 | 0.36932927 | 0.0013451 | -0.1466859 | 57.7878571 | 106.374054 | 1.7046392 | 218.685669 | 0.25864956 | -41.2697144 |
| 15 | 25 | hTDP-43 | 0.01998934 | 0.75930517 | 0.00419249 | 0.00529149 | 0.3794117 | 0.00134371 | -0.238875 | 56.3930626 | 101.866676 | 1.89555526 | 171.048142 | 0.31580511 | -27.7499466 |
| 16 | 25 | Control | 0.02904954 | 0.41019722 | 0.00429867 | 0.00937527 | 0.60333407 | 0.00163502 | -0.35801294 | 67.0225983 | 119.666603 | 1.74231482 | 244.697357 | 0.35318202 | -71.731926 |
| 17 | 25 | Control | 0.01749922 | 0.65150595 | 0.00380916 | 0.00742122 | 0.53745896 | 0.00079705 | -0.08531979 | 60.7037926 | 122.418961 | 1.70739389 | 412.152588 | 0.22882512 | -58.1177788 |
| 18 | 25 | Control | 0.033035 | 0.47360716 | 0.00410493 | 0.00792043 | 0.49980354 | 0.00225542 | -0.23690078 | 62.0736427 | 124.058289 | 1.66062176 | 258.139008 | 0.329009 | -50.6728325 |
| 19 | 25 | Control | 0.01465755 | 0.23315405 | 0.0041899 | 0.01067463 | 0.6787281 | 0.00211745 | -0.75386947 | 63.3489533 | 119.457993 | 2.19671559 | 378.210907 | 0.50353921 | -68.784668 |
| 20 | 25 | Control | 0.0222084 | 0.39128674 | 0.00380775 | 0.00903571 | 0.67563558 | 0.00150144 | -0.17071502 | 66.9425888 | 120.615494 | 1.93258882 | 271.942078 | 0.30076894 | -74.2688828 |
| 21 | 20 | hTDP-43 | 0.02015294 | 0.80163841 | 0.00415651 | 0.00548237 | 0.38877523 | 0.00163712 | -0.34310654 | 58.720768 | 114.065231 | 2.33693314 | 192.846146 | 0.41081387 | -31.5830765 |
| 22 | 20 | hTDP-43 | 0.02906719 | 0.63790429 | 0.00417618 | 0.00632761 | 0.41227913 | 0.00197336 | -0.30531997 | 56.7497101 | 77.7536163 | 2.42114663 | 179.651032 | 0.41502941 | -26.7404785 |
| 23 | 20 | hTDP-43 | 0.02220347 | 0.79635534 | 0.00526514 | 0.00586201 | 0.40836638 | 0.0015042 | -0.35549614 | 58.3563767 | 106.915695 | 2.52340365 | 194.644989 | 0.40720242 | -28.1385174 |
| 24 | 20 | hTDP-43 | 0.02220347 | 0.39101569 | 0.00427725 | 0.00690866 | 0.43106338 | 0.00211833 | -0.45531178 | 56.4714203 | 95.9585648 | 2.50094175 | 230.724655 | 0.47635442 | -29.5101662 |
| 25 | 20 | hTDP-43 | 0.02913818 | 0.62973543 | 0.00407676 | 0.00667146 | 0.39668536 | 0.0019575 | -0.35254109 | 53.0702057 | 87.4567261 | 2.50319433 | 215.063141 | 0.45799768 | -24.9776497 |
| 26 | 20 | Control | 0.0316259 | 0.34064814 | 0.00396761 | 0.00685774 | 0.48491764 | 0.00209816 | -0.39412582 | 63.620327 | 92.1186371 | 1.92816138 | 241.282715 | 0.40773645 | -28.494278 |
| 27 | 20 | Control | 0.03626567 | 0.35555139 | 0.004009 | 0.00661289 | 0.47931445 | 0.00213969 | -0.48001313 | 64.4948196 | 101.276245 | 2.5151875 | 302.440826 | 0.52604032 | -27.4059906 |
| 28 | 20 | Control | 0.02975265 | 0.503047 | 0.00371433 | 0.00611935 | 0.51076949 | 0.00141209 | -0.16041274 | 61.4882202 | 110.416428 | 1.7834059 | 252.079895 | 0.27124742 | -32.5777016 |
| 29 | 20 | Control | 0.04469249 | 0.32818947 | 0.00396258 | 0.00639539 | 0.46635717 | 0.00224139 | -0.25691018 | 61.5382538 | 88.725708 | 1.94609165 | 238.888809 | 0.40331045 | -26.941452 |
| 30 | 20 | Control | 0.03278652 | 0.364612 | 0.00403026 | 0.00622797 | 0.46938121 | 0.00221481 | -0.21301723 | 62.3028183 | 100.921021 | 2.03904343 | 236.88501 | 0.39814571 | -27.0041199 |
| 31 | 25 | hTDP-43 | 0.01582911 | 0.67735202 | 0.00346419 | 0.00708441 | 0.44302499 | 0.00117806 | -0.50974417 | 59.9331055 | 118.315994 | 2.3146925 | 286.561096 | 0.39802417 | -44.4615479 |
| 32 | 25 | hTDP-43 | 0.00888336 | 0.9337636 | 0.00304397 | 0.00463916 | 0.29985637 | 0.00106518 | -0.10740827 | 57.1622582 | 80.7701569 | 2.25723481 | 177.117981 | 0.20826055 | -28.5432396 |
| 33 | 25 | hTDP-43 | 0.02248601 | 0.65360565 | 0.00447469 | 0.00638464 | 0.4114984 | 0.00136804 | -0.41032803 | 61.8474731 | 113.885529 | 2.28787136 | 292.106018 | 0.43184164 | -29.959795 |
| 34 | 25 | hTDP-43 | 0.01398508 | 0.61566329 | 0.00388487 | 0.00695165 | 0.44671702 | 0.00113297 | -0.39823398 | 60.4043465 | 124.975662 | 2.13647175 | 294.408447 | 0.38089341 | -33.5894699 |
| 35 | 25 | hTDP-43 | 0.03039177 | 0.71099113 | 0.00401069 | 0.00539834 | 0.3816492 | 0.00125662 | -0.33504269 | 56.4683876 | 114.677239 | 1.95747161 | 187.679306 | 0.35894191 | -34.8271904 |
| 36 | 25 | Control | 0.03349006 | 0.38108559 | 0.00399773 | 0.00736776 | 0.53370547 | 0.00178825 | -0.27128837 | 59.6122093 | 129.677734 | 1.48599076 | 218.429627 | 0.29256976 | -40.5452881 |
| 37 | 25 | Control | 0.04199254 | 0.56089975 | 0.00361561 | 0.00571044 | 0.50368249 | 0.00178468 | -0.19141141 | 58.0077133 | 132.566406 | 1.46252573 | 198.234467 | 0.26465756 | -33.3455048 |
| 38 | 25 | Control | 0.03287574 | 0.62097828 | 0.00387178 | 0.00677988 | 0.45889449 | 0.00145278 | -0.31037974 | 59.9791222 | 119.788933 | 1.36311507 | 231.099197 | 0.2740671 | -30.7699604 |
| 39 | 25 | Control | 0.03043227 | 0.51637693 | 0.0035946 | 0.00672744 | 0.55472791 | 0.00161784 | -0.20419613 | 61.5583992 | 114.834053 | 1.4012053 | 225.62796 | 0.26182136 | -39.3292198 |
| 40 | 25 | Control | 0.02976587 | 0.65459518 | 0.00369592 | 0.00567107 | 0.4703176 | 0.0017583 | -0.22634009 | 59.4548111 | 131.464203 | 1.35838139 | 191.64679 | 0.22623399 | -33.3131332 |

| ID | d_ang_vel_tail | rel_to_body_r | width_head_b | curvature_std_ | d_area_w_bac | d_curvature_m | rel_to_head_b | width_midbod | curvature_hea | d_curvature_s | curvature_me | curvature_std_ | d_width_head |
|---|---|---|---|---|---|---|---|---|---|---|---|---|---|
| 1 | 2.41593432 | 16.0771999 | 47.7109528 | 0.00288395 | -7521.29688 | 0.00285508 | -10.1458559 | 52.3325424 | 0.01079724 | 0.00192487 | 0.01034063 | 0.00216985 | 11.9718552 |
| 2 | 2.13768841 | 12.6854115 | 49.3824959 | 0.00246551 | -6651.05225 | 0.0029506 | -11.5017042 | 64.3122025 | 0.01059604 | 0.00256325 | 0.01011675 | 0.00304238 | 12.8214264 |
| 3 | 1.9602375 | 15.7235394 | 50.1784515 | 0.0036611 | -8004.61621 | 0.00299713 | -9.69245625 | 41.0981636 | 0.01069099 | 0.00229882 | 0.01014178 | 0.00229333 | 13.6085033 |
| 4 | 1.93636572 | 11.7000618 | 49.0441284 | 0.00224719 | -8072.00293 | 0.00152852 | -9.85307407 | 67.7210999 | 0.01101443 | 0.00199928 | 0.01038692 | 0.00219491 | 12.8643398 |
| 5 | 2.18455505 | 11.2251196 | 49.6295357 | 0.00213923 | -6953.31006 | 0.00121414 | -8.01690388 | 66.4573669 | 0.01001526 | 0.00140158 | 0.00971224 | 0.00245039 | 12.0448227 |
| 6 | 2.52872968 | 11.4265766 | 51.9610672 | 0.00210411 | -7966.47022 | 0.00272236 | -14.2920094 | 70.0623932 | 0.01044333 | 0.0024283 | 0.00987292 | 0.00284124 | 13.439292 |
| 7 | 2.27933168 | 12.2052336 | 52.8087769 | 0.00204795 | -7825.43262 | 0.00354964 | -13.6854076 | 39.4081268 | 0.00933158 | 0.00237593 | 0.00881769 | 0.00252571 | 10.9401894 |
| 8 | 2.98442054 | 10.4491091 | 49.7379036 | 0.00213658 | -7006.12256 | 0.00225472 | -12.7935963 | 64.8155212 | 0.01049306 | 0.00269266 | 0.00986526 | 0.00294691 | 13.8612032 |
| 9 | 3.07900333 | 14.2920961 | 51.2179871 | 0.00207558 | -8554.56152 | 0.00411834 | -14.2946491 | 67.9196625 | 0.00990748 | 0.00246817 | 0.00925938 | 0.00262621 | 14.6681194 |
| 10 | 3.67289639 | 11.8146162 | 50.4794426 | 0.00241415 | -7095.7334 | 0.00296063 | -13.0056686 | 67.0643234 | 0.0104674 | 0.00279523 | 0.00979536 | 0.00282771 | 14.902854 |
| 11 | 1.62127245 | 13.3114414 | 51.7227593 | 0.00219932 | -9152.91406 | 0.00091909 | -8.72495365 | 71.388031 | 0.00910846 | 0.00141924 | 0.00876512 | 0.00159969 | 11.3187552 |
| 12 | 1.58614302 | 13.0600328 | 50.9758873 | 0.002242 | -7210.60059 | 0.00134286 | -7.3689332 | 69.4837341 | 0.00907224 | 0.00132075 | 0.00867911 | 0.00180764 | 12.6010475 |
| 13 | 2.07763863 | 19.382658 | 52.1389275 | 0.00279672 | -8656.08008 | 0.00090557 | -10.6616278 | 70.1983337 | 0.01122623 | 0.00202788 | 0.0107508 | 0.00326921 | 13.017189 |
| 14 | 1.94877303 | 13.2192392 | 53.5249977 | 0.00236166 | -6925.80078 | 0.00100715 | -9.58933163 | 73.4824219 | 0.00960639 | 0.00166769 | 0.00920077 | 0.00180379 | 12.1790104 |
| 15 | 1.97265947 | 13.5323353 | 51.0482559 | 0.00236228 | -7653.34863 | 0.00113053 | -8.46138001 | 71.6088867 | 0.00973428 | 0.00168695 | 0.00932605 | 0.00165842 | 12.8058615 |
| 16 | 2.6874752 | 14.9158812 | 61.7689743 | 0.0015649 | -7990.87598 | 0.00167734 | -13.1433296 | 80.9814606 | 0.00899123 | 0.00190473 | 0.00851038 | 0.00264537 | 14.2625656 |
| 17 | 2.25124979 | 26.0176106 | 52.3182526 | 0.00193713 | -8172.33594 | 0.00835172 | -16.7065735 | 70.7212372 | 0.00873106 | 0.00224304 | 0.00831665 | 0.00267247 | 11.9907856 |
| 18 | 2.24581051 | 15.7842207 | 57.1822929 | 0.00189652 | -7654.25635 | 0.00248436 | -14.1308126 | 72.4537506 | 0.00900281 | 0.00211198 | 0.00853017 | 0.00265388 | 13.4775877 |
| 19 | 3.06840539 | 19.8277302 | 57.855545 | 0.00194819 | -7158.10791 | 0.00499133 | -14.7734737 | 76.1175079 | 0.00926872 | 0.00224318 | 0.00876072 | 0.00251256 | 15.4376392 |
| 20 | 2.72410989 | 14.7902536 | 59.7097778 | 0.00163355 | -6748.72461 | 0.00172825 | -13.9406376 | 78.2168427 | 0.00823695 | 0.00195994 | 0.00774425 | 0.00247361 | 12.7116919 |
| 21 | 1.94773018 | 12.0104561 | 53.4212303 | 0.00206129 | -7714.67481 | 0.0010602 | -10.1808958 | 73.4699097 | 0.01038184 | 0.00196621 | 0.00986545 | 0.00158038 | 12.7677326 |
| 22 | 1.84865701 | 11.3022518 | 51.7318802 | 0.00207831 | -6763.58643 | 0.00114439 | -10.1980114 | 70.430542 | 0.00930628 | 0.00188014 | 0.0088293 | 0.00188684 | 12.3631115 |
| 23 | 1.92217362 | 13.2392349 | 52.5191383 | 0.0034844 | -7292.17383 | 0.00116433 | -9.55302048 | 70.2848663 | 0.01102994 | 0.00194073 | 0.01045294 | 0.00199205 | 13.5371447 |
| 24 | 2.02731967 | 11.4943495 | 50.1629295 | 0.00202439 | -5468.76953 | 0.00154206 | -10.5290737 | 65.7524719 | 0.00899329 | 0.00219691 | 0.00855436 | 0.00231871 | 13.3949642 |
| 25 | 1.84829128 | 10.7465601 | 48.7404823 | 0.00173832 | -7053.81348 | 0.00126992 | -11.1405306 | 66.2020874 | 0.00904935 | 0.0023937 | 0.00858059 | 0.00204111 | 12.1551809 |
| 26 | 2.09885001 | 10.2068119 | 59.0257759 | 0.00198203 | -7666.09668 | 0.00151586 | -13.4079094 | 78.5143204 | 0.00932163 | 0.00235874 | 0.00881913 | 0.00245174 | 13.9466877 |
| 27 | 2.10100174 | 11.4341812 | 59.4798698 | 0.00203612 | -6590.271 | 0.00161668 | -16.1867619 | 79.9545975 | 0.00952958 | 0.00257121 | 0.00887082 | 0.00246256 | 14.3135309 |
| 28 | 1.98847473 | 10.9245415 | 57.3751373 | 0.00192609 | -9230.67188 | 0.00138805 | -14.3795109 | 78.0117493 | 0.00852517 | 0.00219113 | 0.0081032 | 0.00240957 | 10.8730021 |
| 29 | 2.02996397 | 11.111064 | 56.1188698 | 0.00198582 | -7957.59619 | 0.00182222 | -14.0480185 | 75.8099289 | 0.00946828 | 0.0024362 | 0.00886158 | 0.00255058 | 13.5868244 |
| 30 | 2.04105687 | 9.56194592 | 58.605545 | 0.00181128 | -8244.42773 | 0.00145156 | -14.5596428 | 79.2522583 | 0.00900401 | 0.00231755 | 0.00841194 | 0.00242227 | 13.094306 |
| 31 | 2.19599056 | 22.55966 | 55.2547989 | 0.00128202 | -7934.93262 | 0.00178978 | -11.8853846 | 73.87957 | 0.00973632 | 0.00177707 | 0.00950403 | 0.00150266 | 13.1053181 |
| 32 | 1.76820493 | 11.4244375 | 54.9697418 | 0.00205035 | -9542.34277 | 0.00090524 | -10.0381413 | 74.0897598 | 0.00900658 | 0.00183228 | 0.00845299 | 0.00150121 | 9.55016613 |
| 33 | 1.82778311 | 15.0059776 | 53.2861023 | 0.00180192 | -7829.76172 | 0.00192017 | -12.4231968 | 69.3155594 | 0.01079969 | 0.00240507 | 0.01027976 | 0.00233953 | 13.7630577 |
| 34 | 2.13923812 | 20.9929791 | 56.5685915 | 0.00150657 | -7547.98731 | 0.00265596 | -11.648118 | 71.7383652 | 0.00929475 | 0.00183725 | 0.00889266 | 0.00159847 | 12.9905815 |
| 35 | 1.933851 | 14.6820355 | 49.7998848 | 0.00215169 | -7743.59766 | 0.00107536 | -10.7476969 | 67.5416794 | 0.010104 | 0.00192708 | 0.00951318 | 0.00200549 | 13.4609823 |
| 36 | 2.35545445 | 17.5399742 | 55.1605721 | 0.00221496 | -9044.32422 | 0.00257063 | -11.3081741 | 74.5391159 | 0.00964054 | 0.00189077 | 0.00917237 | 0.00213716 | 12.6642466 |
| 37 | 2.03106237 | 14.7581768 | 53.7498741 | 0.00234967 | -8903.16211 | 0.00167641 | -11.298831 | 72.6284561 | 0.00861729 | 0.00193069 | 0.008187 | 0.00212065 | 11.812273 |
| 38 | 2.01757765 | 13.0220537 | 56.2346497 | 0.00212619 | -7036.99023 | 0.00225236 | -11.0704317 | 72.5795898 | 0.00969003 | 0.00187649 | 0.00919357 | 0.00210459 | 12.644661 |
| 39 | 2.39780593 | 17.1854458 | 56.8332329 | 0.00201144 | -7752.01514 | 0.00246587 | -10.8473997 | 78.2690888 | 0.00900106 | 0.00184393 | 0.00859671 | 0.00204997 | 12.5841646 |
| 40 | 1.98224521 | 13.3679676 | 55.8217392 | 0.00217489 | -8171.13965 | 0.00152434 | -11.583869 | 72.9779663 | 0.00946316 | 0.00182525 | 0.00898373 | 0.00204233 | 10.9477367 |

| ID | d_minor_axis_ | length_w_back | width_midbod | motion_mode_ | curvature_std_ | rel_to_neck_r | motion_mode_ | curvature_std_ | ang_vel_abs_ | length_IQR | d_ang_vel_hea | rel_to_hips_ra | d_speed_head |
|---|---|---|---|---|---|---|---|---|---|---|---|---|---|
| 1 | 33.8154068 | 792.705811 | 61.2333679 | 0.04636045 | 0.00131843 | -0.34656459 | 0.01129529 | 0.00096413 | 0.0444511 | 130.865112 | 0.38846636 | 0.00853099 | -1.42970896 |
| 2 | 37.5447502 | 812.663086 | 63.4838905 | 0.13112963 | 0.00158383 | -0.34455323 | 0.02054734 | 0.00098164 | 0.04725885 | 90.767273 | 0.75508666 | 0.17023301 | -4.36637783 |
| 3 | 36.0889435 | 807.171509 | 65.4950562 | 0.0951617 | 0.00142735 | -0.22040272 | 0.00577444 | 0.00097727 | 0.0353356 | 240.279419 | 0.374724 | 0.02246337 | 0.3281464 |
| 4 | 27.4043751 | 827.964356 | 68.5479965 | 0.0260041 | 0.00153759 | -0.61344129 | 0.00555087 | 0.00076769 | 0.03405818 | 63.8092041 | 0.61206275 | 0.00192466 | -2.66551828 |
| 5 | 33.754055 | 762.727112 | 67.6746979 | 0.10151826 | 0.00111044 | -0.58106995 | 0.00499867 | 0.00067557 | 0.05229347 | 143.071594 | 0.60186529 | 0.15486792 | -4.58838177 |
| 6 | 42.4319954 | 912.019898 | 70.2380981 | 0.38206788 | 0.00173412 | 0.51486677 | 0.03531921 | 0.00100732 | 0.05648562 | 60.9671021 | 0.5709089 | 0.35053995 | -3.5244503 |
| 7 | 40.8769302 | 978.182678 | 71.2171936 | 0.34704177 | 0.00155237 | 0.09211768 | 0.03289035 | 0.00084486 | 0.04610255 | 50.9763184 | 0.32017764 | 0.25979698 | -4.73138571 |
| 8 | 43.5398903 | 883.254517 | 64.4454041 | 0.60220531 | 0.00172807 | 0.42085707 | 0.04194631 | 0.00106565 | 0.06549604 | 73.9592285 | 0.80318332 | 0.36716133 | -6.72951794 |
| 9 | 54.6924706 | 930.976257 | 67.3447495 | 0.56042365 | 0.00162739 | -0.32972372 | 0.02506989 | 0.00099631 | 0.09780856 | 118.450623 | 0.70335597 | 0.32013148 | -7.81301737 |
| 10 | 47.9446411 | 886.773377 | 67.3582001 | 0.58410716 | 0.00173085 | 0.01963498 | 0.03926527 | 0.00109874 | 0.07475964 | 40.9709473 | 0.86311495 | 0.33200008 | -5.15084505 |
| 11 | 31.4715099 | 908.631775 | 73.2940521 | 0.12096968 | 0.00098847 | -0.55664498 | 0.00976779 | 0.00058706 | 0.03606755 | 73.9040527 | 0.45477858 | -0.04358589 | -1.93269539 |
| 12 | 27.7408371 | 899.367004 | 67.1391983 | 0.16467308 | 0.00092452 | -0.31720778 | 0.00954537 | 0.00071265 | 0.03831303 | 47.019043 | 0.49849924 | 0.00727744 | -0.42444521 |
| 13 | 21.5189934 | 864.954346 | 70 | 0.0223478 | 0.0012717 | -0.84425157 | 0.00471824 | 0.00071234 | 0.04499707 | 65.6224976 | 0.63999999 | -0.08472462 | -3.05176425 |
| 14 | 34.1601181 | 923.865295 | 73.3498993 | 0.04705354 | 0.00114517 | -0.26307163 | 0.00532765 | 0.00069565 | 0.04052549 | 47.6392212 | 0.47961003 | 0.06631409 | -0.96236235 |
| 15 | 31.7785931 | 849.247498 | 71.3446045 | 0.18341108 | 0.00105666 | -0.44703478 | 0.00999467 | 0.00063662 | 0.04095165 | 122.825867 | 0.54747099 | -0.02732479 | -0.70841539 |
| 16 | 68.3731461 | 1081.8468 | 80.8817444 | 0.41928355 | 0.00137513 | 0.74321145 | 0.02664417 | 0.00104644 | 0.05936801 | 48.911377 | 0.55490685 | 0.44990417 | -6.51932287 |
| 17 | 49.442482 | 1008.24988 | 68.5229874 | 0.2219213 | 0.0014973 | 0.18881711 | 0.01694369 | 0.00094737 | 0.03517304 | 250.524231 | 0.26166448 | 0.22176206 | -1.10559106 |
| 18 | 49.8761139 | 1021.84149 | 72.5356369 | 0.30355219 | 0.00138991 | 0.73049367 | 0.03109177 | 0.00103016 | 0.05179289 | 118.109863 | 0.48067385 | 0.53633094 | -4.32382774 |
| 19 | 60.6138878 | 1015.26178 | 75.4242706 | 0.56923183 | 0.00146744 | -0.38563526 | 0.02465133 | 0.00109396 | 0.11105686 | 77.4915772 | 0.70882005 | 0.57910585 | -7.5187521 |
| 20 | 67.2868347 | 1090.61377 | 78.0538406 | 0.40961882 | 0.00130837 | 0.31017587 | 0.03930887 | 0.00090975 | 0.05594064 | 72.3320313 | 0.36452305 | 0.31448215 | -4.92488003 |
| 21 | 32.5583763 | 875.044006 | 72.3852997 | 0.05470581 | 0.0011758 | -0.52632088 | 0.01442137 | 0.00079568 | 0.04243702 | 55.5606079 | 0.8182013 | 0.0696507 | -1.95907927 |
| 22 | 36.3250732 | 855.754395 | 70.7698975 | 0.18265217 | 0.00118071 | -0.50021142 | 0.02285435 | 0.00078676 | 0.04513678 | 70.8659668 | 0.75481343 | 0.09692347 | -4.64287615 |
| 23 | 28.3448219 | 826.578674 | 69.9969101 | 0.05463442 | 0.00126906 | -0.67044395 | 0.01842888 | 0.00090688 | 0.04294769 | 81.4138794 | 0.74827033 | 0.06204169 | -2.9075613 |
| 24 | 37.6151009 | 915.492676 | 65.5952759 | 0.31441577 | 0.00132954 | -0.11018801 | 0.0294196 | 0.00091992 | 0.05232074 | 40.2816162 | 0.84614199 | 0.24659184 | -5.61179352 |
| 25 | 31.7924976 | 849.720947 | 65.3846588 | 0.16999007 | 0.00137458 | -0.38586411 | 0.02775064 | 0.00085198 | 0.04114152 | 125.814453 | 0.878479 | 0.09843975 | -1.64363027 |
| 26 | 40.408535 | 964.984558 | 77.5498123 | 0.38722548 | 0.00147936 | 0.61287475 | 0.0434625 | 0.00092155 | 0.04528872 | 48.9431152 | 0.70228863 | 0.38266677 | -6.95396137 |
| 27 | 40.7730598 | 973.57196 | 79.99617 | 0.29595483 | 0.00154706 | 0.80388379 | 0.044777 | 0.0009598 | 0.04338998 | 43.2143555 | 0.93003815 | 0.43478149 | -10.1747026 |
| 28 | 39.4169998 | 1018.71326 | 77.7145233 | 0.25958302 | 0.00142218 | 0.32827529 | 0.03952218 | 0.00071684 | 0.03565303 | 91.3168945 | 0.29802376 | 0.37383622 | -0.91214734 |
| 29 | 36.2210159 | 949.997986 | 73.6032257 | 0.30604177 | 0.00161714 | 0.66080552 | 0.05825588 | 0.00100487 | 0.04592314 | 57.6121826 | 0.70858008 | 0.40192586 | -4.30182457 |
| 30 | 36.4545364 | 993.22876 | 79.2957916 | 0.34216273 | 0.00144235 | 0.83648884 | 0.04260395 | 0.00101471 | 0.03853514 | 43.9217529 | 0.62902772 | 0.45427722 | -4.02972078 |
| 31 | 61.1347885 | 1006.08563 | 75.899765 | 0.23475053 | 0.00099643 | -0.9982307 | 0.0138585 | 0.00056502 | 0.04776922 | 88.7394409 | 0.64468479 | -0.06282842 | -6.6259532 |
| 32 | 43.0916214 | 895.023926 | 71.9340897 | 0.04265028 | 0.00104874 | -0.19627968 | 0.00333126 | 0.00057332 | 0.0244643 | 44.4830322 | 0.09602317 | -0.01323612 | -0.27113676 |
| 33 | 37.5305748 | 886.732544 | 72.1010437 | 0.18159993 | 0.00141026 | -0.50071418 | 0.02276361 | 0.00084228 | 0.04269707 | 169.224182 | 0.68627441 | 0.08991709 | -1.17620802 |
| 34 | 43.2854958 | 880.654114 | 75.5494843 | 0.2734683 | 0.00110798 | -0.5534482 | 0.01709288 | 0.00068721 | 0.05627845 | 187.745056 | 0.6418246 | 0.09327582 | 0.21497926 |
| 35 | 30.658699 | 855.643799 | 67.3497467 | 0.14429575 | 0.0012742 | -0.72791386 | 0.02639869 | 0.00071459 | 0.04714416 | 69.6658325 | 0.60561657 | 0.03678212 | -1.23016644 |
| 36 | 37.8429337 | 936.273071 | 73.9396057 | 0.38083693 | 0.00130199 | 0.08975513 | 0.03947835 | 0.00090259 | 0.0635065 | 72.5788574 | 0.45643261 | 0.248693 | -1.29401791 |
| 37 | 33.258522 | 934.081543 | 72.7143784 | 0.18853081 | 0.00126304 | 0.12493935 | 0.03643797 | 0.00079215 | 0.04623573 | 81.9766235 | 0.40113717 | 0.31238276 | -1.67094779 |
| 38 | 37.871624 | 935.357117 | 72.3800049 | 0.20059532 | 0.00127191 | 0.0311929 | 0.02776667 | 0.00084637 | 0.04358691 | 76.8182983 | 0.40207657 | 0.22580132 | -2.71720028 |
| 39 | 44.337368 | 942.750061 | 77.9093628 | 0.31668619 | 0.00127118 | 0.08029535 | 0.02887734 | 0.00079014 | 0.05265093 | 50.7431641 | 0.39667293 | 0.26748329 | -3.4052279 |
| 40 | 35.4502411 | 952.286926 | 73.934082 | 0.17711674 | 0.00130347 | -0.0008071 | 0.02754454 | 0.00078936 | 0.03713945 | 41.8954468 | 0.33309281 | 0.2321122 | -1.58726287 |

| ID | rel_to_head_b | quirkiness_50t | rel_to_head_b | d_width_tail_t | minor_axis_w_ | d_rel_to_neck | d_rel_to_tail_t | curvature_hea | rel_to_head_b | d_path_curvat | rel_to_body_r | d_rel_to_head | d_length_50th | width_midbod |
|---|---|---|---|---|---|---|---|---|---|---|---|---|---|---|
| 1 | 8.80141068 | 0.93239808 | 8.9795208 | 9.67431068 | 232.090744 | 108.546471 | -0.35579139 | 0.00497941 | 8.78635788 | 1.33831513 | -2.74884176 | -1.3283 | -0.3048552 | 80.2289581 |
| 2 | 12.9128618 | 0.95632589 | 10.8328776 | 8.57578278 | 242.018219 | 200.676148 | -0.38284522 | 0.00580168 | 10.5593319 | 0.80717421 | 3.19879437 | -3.3401475 | 0.18697128 | 76.0765991 |
| 3 | 10.1593399 | 0.97716808 | 9.1975193 | 6.18629456 | 236.763886 | 139.788849 | -0.37030497 | 0.00521138 | 9.90799713 | 4.09700775 | 0.95703429 | -2.73148871 | -0.10953644 | 94.3142853 |
| 4 | 9.52521992 | 0.96704161 | 8.57346058 | 8.18660259 | 222.234467 | 119.710701 | -0.34627375 | 0.00743167 | 8.45871639 | 0.75531548 | -3.92412019 | -0.36636156 | -0.27711302 | 74.3606796 |
| 5 | 8.15858078 | 0.95318002 | 8.32953358 | 9.12025547 | 260.909546 | 107.512375 | -0.52132529 | 0.00363835 | 8.40371037 | 0.02555806 | 0.03983722 | -1.35809767 | -0.1066331 | 74.2154923 |
| 6 | 16.4720364 | 0.9857738 | 9.73876095 | 9.89122391 | 217.470795 | 223.83902 | -0.48074907 | 0.00616011 | 9.13979626 | 0.40405765 | 4.88636112 | -3.23101044 | 0.63714594 | 77.9435272 |
| 7 | 15.37432 | 0.98981702 | 10.5655632 | 9.45141316 | 208.496155 | 230.451431 | -0.58451527 | 0.00560405 | 10.010252 | 4.81128073 | 4.1485796 | -3.9514277 | 0.26477116 | 81.0635605 |
| 8 | 14.8402109 | 0.98688227 | 8.45788384 | 9.61140442 | 208.151428 | 207.32399 | -0.45311761 | 0.00595411 | 8.62098885 | 0.43286166 | 5.68054533 | -3.94460821 | 0.66597611 | 76.7294617 |
| 9 | 16.4722099 | 0.98527587 | 9.94126034 | 10.253212 | 240.51474 | 249.878815 | -0.75734901 | 0.00558352 | 9.07555199 | 0.14360042 | 5.17026186 | -3.47701287 | 0.3034749 | 80 |
| 10 | 15.0505619 | 0.98626781 | 9.76883698 | 10.6105986 | 212.874237 | 220.858368 | -0.50640607 | 0.00580468 | 9.1131382 | 0.21238029 | 5.32108212 | -4.12659836 | 0.28918076 | 72.1042481 |
| 11 | 9.22645855 | 0.96532965 | 8.28152084 | 8.81397724 | 267.361542 | 97.6939545 | -0.4602114 | 0.00443365 | 7.82075214 | 1.37668312 | -1.43375087 | -1.62634516 | -0.53057361 | 83.4116974 |
| 12 | 7.57666159 | 0.97615945 | 8.94677353 | 9.84324265 | 244.776856 | 89.7784119 | -0.26161668 | 0.00343936 | 8.72981548 | 0.61470377 | 0.20295298 | -2.17671728 | -0.45155406 | 81.0609665 |
| 13 | 11.6504746 | 0.92901039 | 8.9318552 | 8.7662487 | 286.335358 | 130.075165 | -0.19993375 | 0.00598176 | 9.64505959 | 0.4612655 | -2.74562216 | -2.46465278 | -1.01156426 | 79.4666443 |
| 14 | 9.72575378 | 0.95225281 | 10.7338047 | 10.5763607 | 251.771301 | 140.599945 | -0.20312342 | 0.00549536 | 9.49719048 | 0.56096602 | -3.88080168 | -1.4815234 | -0.222616 | 83.1887817 |
| 15 | 9.01502609 | 0.95883268 | 9.84234047 | 9.56443787 | 272.79361 | 106.004135 | -0.33984983 | 0.00461088 | 8.97540093 | 0.88511473 | -1.04664254 | -1.17888427 | -0.3151319 | 81.6802216 |
| 16 | 14.464201 | 0.98126686 | 9.62820244 | 10.5329695 | 280.683197 | 195.337845 | -0.53075111 | 0.0047983 | 9.07912827 | 0.23261683 | 5.35325003 | -3.54775238 | 0.87367952 | 90.8437958 |
| 17 | 18.8956146 | 0.98925316 | 11.4402933 | 9.13097191 | 255.41185 | 311.357575 | -0.3289952 | 0.0052095 | 10.6383467 | 3.00927353 | 3.66202998 | -3.74456215 | 0.34222981 | 94.6902466 |
| 18 | 15.8665962 | 0.9733212 | 10.4508266 | 10.6691055 | 277.201386 | 211.910263 | -0.60363394 | 0.00512401 | 9.80363655 | 0.41838792 | 5.50207233 | -2.78427076 | 1.03772867 | 91.9170685 |
| 19 | 16.907465 | 0.97889924 | 10.8099155 | 11.9949942 | 262.778778 | 280.215668 | -0.66242677 | 0.00488236 | 10.5812731 | 0.11466252 | 5.67713118 | -3.86589527 | 0.07977765 | 89.0015869 |
| 20 | 15.7264128 | 0.97307134 | 10.1666136 | 10.7960463 | 290.047852 | 216.268189 | -0.46280798 | 0.00445851 | 10.0995054 | 1.15724635 | 6.74119186 | -3.87583375 | 0.4498826 | 90.4252014 |
| 21 | 10.2169867 | 0.95143575 | 11.3332996 | 8.78324509 | 243.622192 | 131.73378 | -0.2206277 | 0.00482372 | 10.6729517 | 0.78913522 | -1.19434571 | -2.68996739 | -0.03739201 | 81.8978729 |
| 22 | 10.8147783 | 0.96364301 | 11.1306868 | 8.48456574 | 265.539215 | 131.143524 | -0.48734307 | 0.00438825 | 10.7807598 | 0.6430878 | 1.71154666 | -2.66395807 | 0.08288635 | 78.8442307 |
| 23 | 10.4216547 | 0.95195454 | 9.90679836 | 9.51042175 | 233.189041 | 133.616837 | -0.53732264 | 0.00537487 | 9.38746929 | 1.16273058 | 0.76032054 | -2.57159138 | -0.05498873 | 84.3615723 |
| 24 | 11.7362099 | 0.97759366 | 10.5544672 | 9.54104614 | 250.905853 | 170.423996 | -0.63524669 | 0.00450303 | 10.1273727 | 0.30992043 | 2.9043231 | -3.4238205 | 0.00689342 | 81.1053085 |
| 25 | 12.1214113 | 0.97416669 | 9.31995201 | 8.08189774 | 249.19046 | 162.780884 | -0.38931957 | 0.00480202 | 8.87830925 | 0.2166222 | 3.18741345 | -2.42545366 | 0.14248158 | 72.5858002 |
| 26 | 15.4865933 | 0.98782277 | 9.58936691 | 9.90712357 | 235.343247 | 196.320404 | -0.48089299 | 0.00525566 | 9.07158375 | 0.58752203 | 5.8643837 | -3.36993313 | 0.68354029 | 88.2728882 |
| 27 | 18.1219482 | 0.98507893 | 13.0332193 | 9.18917656 | 250.251267 | 246.857407 | -0.52380651 | 0.00542353 | 11.6843147 | 0.83010799 | 7.55361891 | -4.41305304 | 0.6462343 | 85.407753 |
| 28 | 16.5554752 | 0.98711121 | 10.5675249 | 8.23682594 | 271.341278 | 208.769791 | -0.35637367 | 0.00524639 | 9.94928455 | 0.99589241 | 6.26051283 | -3.2752595 | 0.38092592 | 86.6560974 |
| 29 | 15.7333412 | 0.98465246 | 10.3227091 | 9.81513214 | 241.740387 | 201.45636 | -0.70973313 | 0.0055756 | 9.93828011 | 0.83283174 | 5.79236412 | -3.51336074 | 0.73419732 | 86.2391815 |
| 30 | 16.3737259 | 0.98448831 | 10.4938183 | 9.02918625 | 248.181015 | 202.089935 | -0.53051448 | 0.00507348 | 9.91857338 | 0.79009044 | 5.45535755 | -3.56139731 | 0.78208792 | 85.6037064 |
| 31 | 12.7395659 | 0.96962917 | 12.6646442 | 8.75187492 | 325.304993 | 183.647034 | -0.43980181 | 0.00398588 | 11.1144667 | 0.4903267 | 2.5034585 | -3.23556376 | -0.55830663 | 100.746796 |
| 32 | 10.0890131 | 0.98880494 | 10.4225435 | 10.9525681 | 257.636414 | 116.439735 | -0.32535505 | 0.00431028 | 10.5087719 | 1.4338268 | 0.54827845 | -3.06033516 | -0.04292697 | 89.420784 |
| 33 | 14.1926518 | 0.97007799 | 12.3800707 | 9.22650051 | 276.352783 | 213.576508 | -0.34820831 | 0.0053531 | 10.7621069 | 0.64806861 | 4.88179493 | -4.35799599 | -0.22909883 | 81.9332504 |
| 34 | 12.9318419 | 0.96879149 | 10.6364679 | 9.35266018 | 336.103943 | 193.211166 | -0.44332486 | 0.00393979 | 10.3991928 | 0.32726583 | 6.08740711 | -4.00463247 | -0.10235172 | 82.9287415 |
| 35 | 12.7115154 | 0.95394987 | 8.78843117 | 9.14500904 | 264.908081 | 133.152573 | -0.2938259 | 0.00585241 | 8.7223444 | 0.84224206 | 0.3436577 | -2.19515395 | -0.4120343 | 85.1045837 |
| 36 | 12.6114941 | 0.96915925 | 6.96072245 | 9.92368507 | 287.723877 | 163.926437 | -0.33346552 | 0.00525133 | 7.00403643 | 0.32069835 | 5.73889923 | -1.8953141 | 0.18838188 | 82.0007858 |
| 37 | 12.6649714 | 0.96383309 | 6.99229813 | 9.2158556 | 298.584137 | 159.171082 | -0.32033646 | 0.00538961 | 6.46485853 | 0.4672831 | 3.23275995 | -1.7697556 | 0.26190871 | 81.6326065 |
| 38 | 12.7682829 | 0.97066498 | 5.8947134 | 9.27429199 | 266.003815 | 169.740784 | -0.32416931 | 0.00532948 | 6.01634741 | 0.3144207 | 4.38965797 | -1.56619144 | 0.10659665 | 87.6808014 |
| 39 | 12.1599255 | 0.97807258 | 5.82829666 | 9.80529213 | 286.371277 | 164.181061 | -0.36303374 | 0.00500886 | 5.96011543 | 0.26399905 | 6.36923742 | -1.46021628 | 0.26242253 | 84.9558411 |
| 40 | 13.808569 | 0.97029102 | 6.78751373 | 7.97894335 | 291.012482 | 156.695267 | -0.27509841 | 0.00552792 | 6.56186438 | 0.35624069 | 4.60366535 | -1.79045034 | 0.2674228 | 79.9790649 |

| ID | curvature_hips | path_transit_t | d_length_w_fo | motion_mode_ | speed_head_ti | curvature_std_ | rel_to_head_b | d_major_axis_ | d_curvature_s | curvature_me | path_coverage | d_rel_to_tail_t | d_rel_to_head |
|---|---|---|---|---|---|---|---|---|---|---|---|---|---|
| 1 | 0.00774891 | 27.984 | 58.3238602 | 0.09031257 | 84.2347488 | 0.00077208 | 0.4234139 | -13.0498886 | 0.00424666 | 0.01138216 | 4437500 | 2.67575336 | 4.58200216 |
| 2 | 0.0044227 | 35.608 | 40.7156906 | 0.12432706 | 70.0553589 | 0.00110526 | 0.5331794 | -7.78188705 | 0.00319385 | 0.00864805 | 7125000 | 2.11681366 | 5.39682007 |
| 3 | 0.00491414 | 31.66 | 46.5071945 | 0.03193783 | 60.9210129 | 0.00109066 | 0.36162198 | -11.06987 | 0.00420332 | 0.0084652 | 8687500 | 2.42684412 | 3.83864832 |
| 4 | 0.00413218 | 46.252 | 43.4180527 | 0.04312227 | 65.209877 | 0.00074164 | 0.4676176 | -7.19146729 | 0.00290512 | 0.00918561 | 4062500 | 1.64817357 | 4.07278347 |
| 5 | 0.00540612 | 41.864 | 46.5768814 | 0.03842833 | 42.3353729 | 0.00088598 | 0.459948 | -9.55810547 | 0.00216477 | 0.01241974 | 6750000 | 1.64114463 | 4.4321661 |
| 6 | 0.00438851 | 18.64 | 33.0753708 | 0.24668205 | 78.0131683 | 0.0007816 | 0.51267362 | -9.87472534 | 0.00246996 | 0.00733309 | 25375000 | 1.90464389 | 3.67325187 |
| 7 | 0.00344485 | 13.872 | 35.2561722 | 0.20178933 | 121.558586 | 0.00054687 | 0.55381507 | -8.79802704 | 0.00256202 | 0.0065385 | 33750000 | 2.03744555 | 4.02196789 |
| 8 | 0.00399661 | 13.536 | 32.7084236 | 0.25185946 | 61.0151062 | 0.00070715 | 0.58015889 | -8.72478485 | 0.00242955 | 0.00701143 | 21062500 | 1.79076409 | 4.28087091 |
| 9 | 0.00381934 | 10.96 | 36.1292801 | 0.23211142 | 125.06736 | 0.00076214 | 0.60870284 | -13.1298065 | 0.00280554 | 0.006946 | 39250000 | 2.10645533 | 4.18069029 |
| 10 | 0.00469082 | 11.732 | 33.1502838 | 0.25356069 | 72.8782349 | 0.00084332 | 0.63064373 | -12.5962257 | 0.00283701 | 0.00811912 | 30750000 | 1.97066498 | 4.80628538 |
| 11 | 0.0022412 | 33.344 | 47.6210365 | 0.08481108 | 87.1061249 | 0.00063335 | 0.36685982 | -5.86910248 | 0.00206943 | 0.00718999 | 8250000 | 1.46909154 | 3.42155099 |
| 12 | 0.00595826 | 41.856 | 44.6291008 | 0.09320582 | 52.4919891 | 0.00061851 | 0.37383905 | -5.20782471 | 0.00207667 | 0.00931463 | 9937500 | 1.44264591 | 3.42154288 |
| 13 | 0.00899305 | 96.532 | 56.768692 | 0.02673298 | 107.930252 | 0.00055885 | 0.44885874 | -14.6297455 | 0.00272527 | 0.01161 | 3000000 | 1.73500669 | 3.95609951 |
| 14 | 0.00508174 | 62.892 | 49.7901154 | 0.04397793 | 97.8024368 | 0.00063441 | 0.36029744 | -0.73165894 | 0.00229833 | 0.00920139 | 3750000 | 1.81322742 | 3.37190652 |
| 15 | 0.00439499 | 41.516 | 45.228157 | 0.05728375 | 67.8462524 | 0.00054572 | 0.41162413 | -3.26318741 | 0.00241125 | 0.00759405 | 8500000 | 1.71913827 | 3.85208821 |
| 16 | 0.00346378 | 13.956 | 35.3453217 | 0.17051923 | 71.548378 | 0.00054546 | 0.46315754 | -21.2165833 | 0.00190145 | 0.0073268 | 24687500 | 1.61728001 | 3.52149892 |
| 17 | 0.00346974 | 14.136 | 41.3914452 | 0.12657275 | 219.584274 | 0.00065658 | 0.4102571 | -16.9467926 | 0.00312512 | 0.00782409 | 32312500 | 2.89245009 | 3.48977351 |
| 18 | 0.00409318 | 19.712 | 37.4561005 | 0.22284065 | 69.3929215 | 0.00062581 | 0.44692308 | -13.3497238 | 0.0022448 | 0.00787308 | 18562500 | 1.84303153 | 3.42115569 |
| 19 | 0.00433816 | 9.434 | 37.7623406 | 0.19761412 | 153.437195 | 0.00068411 | 0.60988009 | -18.4732437 | 0.0026775 | 0.00820188 | 52562500 | 2.27228189 | 4.29333973 |
| 20 | 0.00314457 | 11.36 | 37.2543678 | 0.19909444 | 77.0679321 | 0.00035407 | 0.48696974 | -17.1621323 | 0.00185895 | 0.00689064 | 25562500 | 1.55599034 | 3.91265583 |
| 21 | 0.00471464 | 34.43 | 40.8695831 | 0.14365579 | 65.2994995 | 0.00067307 | 0.49724823 | -6.08558655 | 0.00264203 | 0.00909962 | 5250000 | 1.83760345 | 4.64051962 |
| 22 | 0.00459362 | 31.59 | 42.2748642 | 0.17944354 | 50.0823975 | 0.00073823 | 0.51169074 | -5.56602478 | 0.0023197 | 0.00869463 | 10625000 | 1.72222865 | 4.73168802 |
| 23 | 0.00444716 | 37.266 | 45.5805969 | 0.14901024 | 62.181263 | 0.00069801 | 0.50022143 | -6.57863617 | 0.00309102 | 0.00964263 | 5937500 | 2.10766912 | 4.90377283 |
| 24 | 0.00388942 | 17.2 | 40.2144089 | 0.29456854 | 55.5050621 | 0.0006756 | 0.54335946 | -5.53312302 | 0.00213774 | 0.00739378 | 16687500 | 1.6487807 | 4.84389591 |
| 25 | 0.00455221 | 27.428 | 39.1495972 | 0.2002745 | 50.1442452 | 0.00068706 | 0.52977467 | -6.07738495 | 0.00216735 | 0.00717675 | 8250000 | 1.65786886 | 4.90005064 |
| 26 | 0.00346766 | 21.152 | 34.1441956 | 0.27212638 | 55.5662689 | 0.00066093 | 0.51287592 | -8.94432068 | 0.00216942 | 0.00622415 | 20187500 | 1.65309298 | 3.91521549 |
| 27 | 0.0036238 | 21.828 | 34.5929489 | 0.34849378 | 58.9258347 | 0.00073775 | 0.5789997 | -5.95970154 | 0.00213396 | 0.00647175 | 15812500 | 1.69240844 | 4.98572397 |
| 28 | 0.00368509 | 18.58 | 36.8998604 | 0.23736998 | 54.6093674 | 0.00062085 | 0.46463341 | -8.20093155 | 0.00196038 | 0.00621043 | 14937500 | 1.65110922 | 3.78362536 |
| 29 | 0.00361983 | 20.72 | 34.1500359 | 0.36576876 | 62.9403153 | 0.00069796 | 0.4953506 | -6.2286377 | 0.00240828 | 0.00613967 | 18750000 | 1.8197155 | 4.00376225 |
| 30 | 0.00350746 | 17.36 | 35.0452232 | 0.29322527 | 55.9655457 | 0.00067157 | 0.50814194 | -6.42700195 | 0.00198581 | 0.00633474 | 17562500 | 1.59156406 | 4.13060045 |
| 31 | 0.0027624 | 21.496 | 50.2041512 | 0.08789746 | 113.062958 | 0.00053572 | 0.46447039 | -8.04548264 | 0.00195641 | 0.00665547 | 23437500 | 1.71816099 | 4.82876539 |
| 32 | 0.00292623 | 48.176 | 42.2989502 | 0.02358612 | 60.3498077 | 0.00055167 | 0.3902477 | -0.14610291 | 0.00276429 | 0.00706607 | 3625000 | 1.96656275 | 4.41291571 |
| 33 | 0.00278414 | 23.4 | 41.0708122 | 0.16479443 | 72.7496948 | 0.00058144 | 0.50346631 | -8.77437592 | 0.00228198 | 0.00687628 | 12312500 | 1.78208149 | 4.45323563 |
| 34 | 0.00346587 | 14.36 | 39.5650101 | 0.11086841 | 106.711853 | 0.00056373 | 0.46729684 | -6.78319931 | 0.00200341 | 0.0061994 | 30812500 | 1.72114611 | 4.22850657 |
| 35 | 0.00505084 | 37.352 | 40.9278259 | 0.14471312 | 62.1745377 | 0.00056289 | 0.45396841 | -7.41872788 | 0.00243857 | 0.00805239 | 5937500 | 1.73573339 | 3.9298029 |
| 36 | 0.00499128 | 16.656 | 35.2191925 | 0.23807748 | 72.4369202 | 0.00066505 | 0.41465461 | -10.3878021 | 0.00255958 | 0.00790359 | 31625000 | 1.89021421 | 3.09858012 |
| 37 | 0.00479401 | 27.712 | 36.2580414 | 0.25056943 | 54.6403198 | 0.00067734 | 0.37995499 | -11.1058235 | 0.0023128 | 0.00763336 | 11312500 | 1.76094115 | 3.00037742 |
| 38 | 0.00410872 | 25.616 | 31.143837 | 0.17842641 | 64.9920349 | 0.00062848 | 0.38228211 | -9.89952087 | 0.00237671 | 0.00719451 | 21812500 | 1.81296945 | 2.85345173 |
| 39 | 0.00439648 | 19.144 | 32.352253 | 0.16693688 | 81.8497315 | 0.00062292 | 0.3853389 | -17.1510696 | 0.00242646 | 0.00754447 | 27812500 | 1.90703559 | 2.91580558 |
| 40 | 0.0043063 | 29.532 | 35.5665398 | 0.16828808 | 46.8439522 | 0.00063582 | 0.34238017 | -12.4300957 | 0.00221207 | 0.00724313 | 14125000 | 1.66768742 | 2.8247993 |

| ID | rel_to_head_b | width_tail_bas | curvature_me | curvature_std_ | curvature_std_ | d_major_axis_ | curvature_hips | rel_to_body_r | curvature_tail_ | curvature_me | d_width_midb |
|---|---|---|---|---|---|---|---|---|---|---|---|
| 1 | 9.8183012 | 9.70088196 | 0.0052624 | 0.000848 | 0.002365 | -0.17461777 | 0.01056343 | -0.12111846 | 0.0114659 | 0.00466493 | -6.29787445 |
| 2 | 11.9464016 | 7.36750031 | 0.00458678 | 0.00123074 | 0.0027431 | -0.59490204 | 0.01019936 | 0.18560365 | 0.00941729 | 0.00545907 | -6.03914976 |
| 3 | 9.97755146 | 7.28633881 | 0.00250605 | 0.00096696 | 0.00226645 | -0.33454895 | 0.00847262 | -0.05479066 | 0.01062718 | 0.00596868 | -7.32878208 |
| 4 | 9.48021126 | 7.85738754 | 0.00359255 | 0.00096986 | 0.0021136 | -1.43642426 | 0.00861703 | -0.38753632 | 0.00919039 | 0.0058735 | -4.49200392 |
| 5 | 8.02357483 | 7.0736618 | 0.00360376 | 0.00087408 | 0.00164182 | -0.32615662 | 0.0103825 | -0.0718554 | 0.01398954 | 0.00470776 | -4.7683239 |
| 6 | 15.1308966 | 7.318676 | 0.0036026 | 0.00119196 | 0.0025365 | 0.9929657 | 0.00699072 | 1.73155594 | 0.01017008 | 0.0060676 | -5.71796894 |
| 7 | 14.2267494 | 4.36203766 | 0.00250838 | 0.00104032 | 0.00208357 | 0.5935669 | 0.00590736 | 0.76129764 | 0.00921464 | 0.0055016 | -5.82588673 |
| 8 | 13.7828922 | 6.03817749 | 0.00378955 | 0.00120886 | 0.0024894 | 2.67620087 | 0.00705396 | 2.49934125 | 0.00996576 | 0.00567491 | -5.8344841 |
| 9 | 15.1648197 | 8.51203156 | 0.0037218 | 0.00111751 | 0.00230944 | 1.97601318 | 0.00709593 | 1.95227671 | 0.0102069 | 0.00561975 | -5.65247536 |
| 10 | 13.9136105 | 5.45418167 | 0.00371964 | 0.00118981 | 0.00254618 | 1.97305679 | 0.00755319 | 1.8776294 | 0.01057586 | 0.0054675 | -5.00171661 |
| 11 | 8.80233383 | 5.58264542 | 0.00260968 | 0.00057321 | 0.0012388 | -0.89836121 | 0.00719702 | -0.38517404 | 0.00792289 | 0.00354589 | -5.40249348 |
| 12 | 7.42239046 | 6.93300629 | 0.00290804 | 0.0006831 | 0.00184077 | -0.83217621 | 0.00805206 | -0.31457284 | 0.00937058 | 0.00338979 | -5.34730673 |
| 13 | 10.1857309 | 8.39635468 | 0.00493 | 0.00104854 | 0.00253518 | -0.77495575 | 0.0104908 | -1.06448388 | 0.01426043 | 0.00505754 | -5.57572365 |
| 14 | 9.82576942 | 6.56369019 | 0.00306888 | 0.00070953 | 0.00169911 | 0.0919342 | 0.00920436 | 0.22469058 | 0.01022524 | 0.00497463 | -5.49179316 |
| 15 | 8.56600571 | 6.91809082 | 0.00260432 | 0.00065334 | 0.00153064 | -0.93698502 | 0.00873077 | -0.48896748 | 0.00969697 | 0.00341683 | -5.62940836 |
| 16 | 13.4368143 | 6.93723297 | 0.00349112 | 0.00108771 | 0.00191331 | 2.31819153 | 0.00602994 | 2.67869234 | 0.00915658 | 0.0049625 | -6.44988775 |
| 17 | 17.8058205 | 8.10037231 | 0.00295364 | 0.00116113 | 0.00149888 | 0.18901825 | 0.00573207 | 0.65479863 | 0.01024483 | 0.00589377 | -6.14771605 |
| 18 | 14.68188 | 12.0724106 | 0.00337293 | 0.00119895 | 0.00222967 | 2.01034546 | 0.00732599 | 2.24801588 | 0.01077531 | 0.00508977 | -5.85852623 |
| 19 | 15.7858877 | 8.59407806 | 0.00398855 | 0.0011238 | 0.00207351 | 2.05345154 | 0.00732599 | 2.15682234 | 0.01078261 | 0.00530465 | -6.00436926 |
| 20 | 14.2926884 | 7.9568634 | 0.00386686 | 0.00098398 | 0.0017973 | 1.0766983 | 0.00554431 | 1.04589939 | 0.00883518 | 0.00461527 | -6.10964298 |
| 21 | 9.90508556 | 6.03387833 | 0.003782 | 0.00068382 | 0.00189473 | -1.21192932 | 0.00944504 | -0.0503917 | 0.00866523 | 0.00386171 | -5.47729969 |
| 22 | 9.75586987 | 5.45858383 | 0.00349109 | 0.00073144 | 0.00219777 | -0.60148239 | 0.00935404 | 0.24750967 | 0.00895921 | 0.00404746 | -4.71909523 |
| 23 | 9.68504238 | 8.75399017 | 0.00377495 | 0.00073449 | 0.00192612 | -0.91533661 | 0.00919161 | -0.23292336 | 0.00901334 | 0.00461078 | -5.93144894 |
| 24 | 10.8993645 | 6.41118622 | 0.00360393 | 0.00092262 | 0.00226179 | 0.12602806 | 0.00751392 | 1.18332803 | 0.00886132 | 0.00423664 | -5.30884266 |
| 25 | 10.9500303 | 5.03127289 | 0.00364641 | 0.00082821 | 0.00244074 | -0.62160492 | 0.00830796 | 0.58566666 | 0.00776215 | 0.00403889 | -5.10613442 |
| 26 | 14.1380777 | 6.61857224 | 0.00336457 | 0.00096487 | 0.00215334 | 1.23653412 | 0.00603112 | 2.03499699 | 0.00937831 | 0.00481645 | -5.45827389 |
| 27 | 17.1248035 | 6.63259888 | 0.0032946 | 0.00104212 | 0.00230648 | 2.17399597 | 0.00642565 | 2.57837725 | 0.00888644 | 0.00492336 | -5.36134005 |
| 28 | 15.2761955 | 5.00784302 | 0.00279241 | 0.00098451 | 0.00154691 | 0.60119629 | 0.00576024 | 0.99537265 | 0.00931812 | 0.00481584 | -4.96934652 |
| 29 | 14.7148466 | 6.95841925 | 0.00338369 | 0.00113341 | 0.00217849 | 1.77116394 | 0.00638767 | 2.0798471 | 0.00863017 | 0.0051071 | -5.63048601 |
| 30 | 14.9095001 | 4.94083023 | 0.00329769 | 0.00106463 | 0.00227005 | 1.62763596 | 0.00581769 | 1.90932345 | 0.00853938 | 0.00456493 | -4.89230156 |
| 31 | 11.713028 | 5.26120377 | 0.00290764 | 0.00052462 | 0.00134844 | -0.9061365 | 0.00570655 | -0.12390426 | 0.00710505 | 0.00306981 | -5.31867504 |
| 32 | 10.2460785 | 5.33945084 | 0.00216714 | 0.00061098 | 0.00218167 | -0.10566711 | 0.00475553 | -0.03458335 | 0.00604763 | 0.00333399 | -6.08361721 |
| 33 | 12.696765 | 6.85233688 | 0.00347563 | 0.0008538 | 0.00154736 | -0.98500252 | 0.00791309 | 0.54617274 | 0.00759582 | 0.00537886 | -5.77796936 |
| 34 | 11.213439 | 5.59794235 | 0.00343638 | 0.0006237 | 0.00148692 | -0.62484741 | 0.00679211 | 0.64973176 | 0.0071824 | 0.0038533 | -5.33267498 |
| 35 | 10.8742933 | 7.02158356 | 0.00321024 | 0.00076644 | 0.00161979 | -0.43506622 | 0.0080096 | -0.43732402 | 0.00983263 | 0.00504605 | -5.38178444 |
| 36 | 11.5835228 | 5.4907341 | 0.00411122 | 0.00084009 | 0.00206945 | 0.47454834 | 0.00729266 | 1.48242974 | 0.01143188 | 0.00537489 | -5.96912146 |
| 37 | 11.3241272 | 5.93870544 | 0.00372124 | 0.00085712 | 0.00180943 | 0.04301071 | 0.00741503 | 0.95665932 | 0.01132084 | 0.00502211 | -5.42562723 |
| 38 | 11.6318932 | 5.6687088 | 0.00396834 | 0.00080412 | 0.0015444 | 0.15478134 | 0.00700441 | 1.45141792 | 0.01061626 | 0.00532902 | -5.54022789 |
| 39 | 11.1769867 | 5.75855963 | 0.00388326 | 0.00081123 | 0.00171213 | 0.21743774 | 0.00626865 | 1.38648391 | 0.0113681 | 0.00531587 | -5.59234142 |
| 40 | 12.3365669 | 4.96632385 | 0.00404308 | 0.00077026 | 0.00167936 | -0.09250641 | 0.00718939 | 0.86517048 | 0.01134563 | 0.00554858 | -5.53013563 |

| ID | width_midbod | rel_to_head_b | curvature_mid | curvature_tail_ | curvature_me | width_tail_bas | turn_intra_du | d_rel_to_head | width_midbod | d_curvature_s |
|---|---|---|---|---|---|---|---|---|---|---|
| 1 | 72.1328812 | -7.47756052 | 0.00841416 | 0.01063492 | 0.00360908 | 50.1640015 | 14.6 | 4.20090008 | 69.978447 | 0.00034824 |
| 2 | 69.4190598 | -9.7975378 | 0.0082168 | 0.01044272 | 0.00433275 | 49.5237961 | 212.16 | 6.08477831 | 64.3986664 | 0.00045161 |
| 3 | 73.2483139 | -7.82723236 | 0.00791583 | 0.01055195 | 0.00479029 | 51.1310425 | 15.92 | 5.26481772 | 70.090271 | 0.00042544 |
| 4 | 70.6701279 | -9.35743809 | 0.00693958 | 0.00852983 | 0.00620747 | 50.4631386 | 8.36 | 4.14779902 | 68.835083 | 0.00034664 |
| 5 | 70.4978714 | -8.40676117 | 0.00701964 | 0.0142646 | 0.00444512 | 53.4674034 | 47.04 | 3.55946732 | 66.628624 | 0.00023322 |
| 6 | 74.1531372 | -12.0030069 | 0.00681574 | 0.00816861 | 0.0041397 | 50.7383118 | 15.82 | 5.32899094 | 70.2513962 | 0.00043422 |
| 7 | 76.9749298 | -10.5236998 | 0.00608152 | 0.00709351 | 0.00394225 | 56.0557709 | 31.56 | 5.4486208 | 72.578537 | 0.00041482 |
| 8 | 69.2887192 | -12.0492144 | 0.00730168 | 0.00778803 | 0.00410529 | 48.875267 | 27.96 | 5.57551622 | 65.181427 | 0.0004839 |
| 9 | 71.6816101 | -13.1358824 | 0.00677223 | 0.00801991 | 0.00377887 | 50.5440903 | 16.16 | 5.82294273 | 68.5274582 | 0.00043956 |
| 10 | 70.2074051 | -12.3143377 | 0.00750879 | 0.00823588 | 0.00420869 | 49.9051132 | 10.72 | 6.21555376 | 66.8353195 | 0.00049946 |
| 11 | 78.6001434 | -7.71801519 | 0.00629738 | 0.00692788 | 0.00362973 | 54.6824341 | 900.92 | 3.27876353 | 75.2716217 | 0.00024568 |
| 12 | 72.5317688 | -7.83176756 | 0.00786166 | 0.00871338 | 0.00319859 | 48.3883591 | 270.7 | 3.44676447 | 69.5828857 | 0.00022879 |
| 13 | 76.0170288 | -9.36464787 | 0.01021636 | 0.01157679 | 0.00613185 | 50.6656265 | 900.76 | 4.33272934 | 70.0772324 | 0.00033468 |
| 14 | 78.3879013 | -7.87802315 | 0.0069317 | 0.01087857 | 0.00441827 | 52.5269699 | 1.74 | 3.75621176 | 75.3543778 | 0.00028597 |
| 15 | 76.9753342 | -8.29296398 | 0.00658637 | 0.01303237 | 0.00351294 | 51.3339615 | 67.84 | 4.03777885 | 71.8142319 | 0.00029406 |
| 16 | 87.1359863 | -12.1244135 | 0.00633868 | 0.0072403 | 0.00358919 | 60.006279 | 7.88 | 4.46454144 | 81.1013489 | 0.00033676 |
| 17 | 84.8511963 | -11.0777607 | 0.00584486 | 0.00720911 | 0.00393552 | 57.2422295 | 20.32 | 5.87892819 | 71.4479828 | 0.00040687 |
| 18 | 82.6133347 | -11.6528378 | 0.00736164 | 0.00874619 | 0.00389262 | 56.1623612 | 29.82 | 4.72151232 | 72.9771042 | 0.00036389 |
| 19 | 81.3350678 | -13.7124395 | 0.00679697 | 0.00840615 | 0.00379778 | 58.1170616 | 10.56 | 5.64580631 | 77.1755829 | 0.0003992 |
| 20 | 84.464325 | -11.406251 | 0.00628149 | 0.00709485 | 0.00348665 | 58.9268112 | 8.96 | 4.74071407 | 77.9882736 | 0.00034654 |
| 21 | 77.7588501 | -9.57222748 | 0.00799723 | 0.01048974 | 0.00378365 | 52.3016815 | 13.02 | 4.48665524 | 73.7712174 | 0.00036406 |
| 22 | 73.8153229 | -9.85623169 | 0.00842495 | 0.00802098 | 0.00340142 | 50.5170479 | 279.16 | 4.55455446 | 70.6242065 | 0.00033337 |
| 23 | 79.7984314 | -9.40894795 | 0.0075748 | 0.00908955 | 0.00370826 | 51.214386 | 900.76 | 4.84145927 | 71.8323822 | 0.00031762 |
| 24 | 69.800827 | -10.4912415 | 0.0073067 | 0.00817924 | 0.00364828 | 49.9355545 | 2.7 | 5.1574831 | 65.7145462 | 0.00039132 |
| 25 | 70.0800476 | -10.1226358 | 0.00748994 | 0.00684717 | 0.00366958 | 48.8604088 | 48.24 | 5.19771767 | 66.1724854 | 0.00042235 |
| 26 | 83.5695648 | -12.041091 | 0.00619935 | 0.00771929 | 0.00351418 | 58.8403015 | 7.04 | 5.16589117 | 79.0186386 | 0.00040614 |
| 27 | 82.5973358 | -14.5035029 | 0.0065322 | 0.00749919 | 0.00367308 | 59.3054848 | 14.16 | 5.92435408 | 79.9671555 | 0.0004489 |
| 28 | 82.744133 | -10.8317947 | 0.00531885 | 0.00835657 | 0.00358742 | 56.5978203 | 21.32 | 5.05360889 | 78.125473 | 0.00038979 |
| 29 | 81.421051 | -12.0396767 | 0.00643696 | 0.00742308 | 0.00377712 | 57.4359398 | 4.56 | 5.39251947 | 77.239563 | 0.00043329 |
| 30 | 81.8760376 | -12.3266535 | 0.00639836 | 0.00738501 | 0.00343255 | 57.4485245 | 12.72 | 5.08581924 | 79.4065628 | 0.00041593 |
| 31 | 81.6488953 | -10.9340153 | 0.00591792 | 0.00554099 | 0.0036967 | 55.5114098 | 10.44 | 4.83350134 | 78.8653565 | 0.00031316 |
| 32 | 77.2282486 | -7.43213558 | 0.00493815 | 0.00521363 | 0.00313012 | 49.2449188 | 17.52 | 4.05348778 | 73.0579681 | 0.00032365 |
| 33 | 79.8344955 | -10.1708088 | 0.00715266 | 0.01034465 | 0.0045303 | 55.6313515 | 63.1 | 5.43418169 | 72.3968277 | 0.00043951 |
| 34 | 79.9733353 | -9.96309185 | 0.00602773 | 0.00773532 | 0.00333273 | 54.0987701 | 88.6 | 4.83509636 | 74.3298416 | 0.00031946 |
| 35 | 72.7459412 | -9.26892376 | 0.00758697 | 0.0087169 | 0.00431683 | 50.808754 | 113.76 | 4.38967848 | 66.9731751 | 0.00033683 |
| 36 | 78.8481751 | -9.34714317 | 0.00705711 | 0.00878679 | 0.00396868 | 54.3949432 | 117.24 | 4.3318944 | 74.8801727 | 0.00033452 |
| 37 | 77.7553787 | -8.55631447 | 0.00699517 | 0.00985132 | 0.00413643 | 52.4604378 | 23.58 | 4.24370623 | 73.0655823 | 0.00033603 |
| 38 | 79.5352783 | -8.94185257 | 0.00654413 | 0.00869719 | 0.00417696 | 54.6057854 | 403.2 | 4.22446537 | 72.1573029 | 0.0003343 |
| 39 | 81.7446289 | -8.37017822 | 0.00644348 | 0.00854626 | 0.00388917 | 56.8696327 | 27.96 | 4.32100534 | 79.1417084 | 0.00031953 |
| 40 | 77.586647 | -8.64732933 | 0.00653898 | 0.01010189 | 0.00377138 | 54.3375549 | 33.76 | 3.78651285 | 73.8780823 | 0.00032434 |

| ID | curvature_me | speed_10th | d_ang_vel_he | rel_to_body_r | d_width_tail_t | width_tail_bas | d_speed_head | d_ang_vel_hea | d_width_head | speed_head_ti |
|---|---|---|---|---|---|---|---|---|---|---|
| 1 | 0.01047851 | -18.5475884 | 8.33495045 | 0.02824805 | 0.00500679 | 43.7248993 | -10.6537247 | 3.20114279 | 0.50213933 | -50.2145309 |
| 2 | 0.01052022 | -20.9930954 | 12.0410862 | 0.06983282 | 0.03558993 | 48.8111 | -12.1826382 | 4.91580486 | 0.36100149 | -55.129055 |
| 3 | 0.01058523 | -14.8357973 | 10.1018057 | -0.02940687 | 0.00494719 | 36.0904007 | -8.46518898 | 3.90181732 | 1.12357736 | -41.9884911 |
| 4 | 0.00859241 | -18.9012833 | 8.74620438 | 0.00691009 | 0.01496077 | 50.4843712 | -0.69660407 | 3.5123179 | 0.58064461 | -51.4260101 |
| 5 | 0.01464459 | -15.3630076 | 7.1554203 | 0.17219448 | 0.07870197 | 52.5215035 | -6.33359909 | 2.64933038 | 0.51792264 | -47.5000763 |
| 6 | 0.00763007 | -34.6356239 | 11.2907524 | 0.39535064 | 0.07317662 | 51.6003723 | -10.2709103 | 4.52627134 | 0.76394081 | -60.4416313 |
| 7 | 0.00643935 | -18.878273 | 11.7596407 | 0.21106675 | 0.04895926 | 30.5146961 | -12.1384592 | 4.85619831 | 0.51311255 | -45.4418068 |
| 8 | 0.00740031 | -55.8267479 | 11.9020567 | 0.46415854 | 0.12004972 | 49.5736275 | -11.9726448 | 5.04367161 | 0.73276758 | -88.5926285 |
| 9 | 0.00726907 | -51.1534691 | 11.9205141 | 0.30600634 | 0.06535053 | 51.0836983 | -10.9530516 | 4.92864943 | 0.49183369 | -82.117836 |
| 10 | 0.0077579 | -54.1696281 | 13.0873337 | 0.41433311 | 0.12009144 | 50.18293 | -9.57444763 | 5.44956017 | 0.55919886 | -84.8707047 |
| 11 | 0.00734362 | -18.7013474 | 7.3438592 | -0.06982589 | 0.01018047 | 49.3427391 | -12.2193928 | 2.82875633 | 0.4160285 | -51.0778542 |
| 12 | 0.0089066 | -19.6662769 | 7.21436214 | 0.00506412 | 0.02222061 | 49.7318649 | -8.89977169 | 2.89251328 | 0.49350262 | -52.0523491 |
| 13 | 0.01168616 | -19.4509525 | 9.93787003 | 0.06885769 | 0.04027486 | 48.7505684 | -4.2000494 | 4.42139721 | 0.37001967 | -61.9817848 |
| 14 | 0.01050219 | -17.9564419 | 8.18424988 | 0.08778884 | 0.00731945 | 52.2072601 | 1.35409701 | 3.52810407 | 0.65097809 | -49.4521599 |
| 15 | 0.01156026 | -19.9750977 | 8.95812702 | -0.02748337 | 0.00619292 | 50.4825058 | -5.21054554 | 3.79068589 | 0.61504245 | -53.3854218 |
| 16 | 0.00729618 | -33.615036 | 9.63879299 | 0.64921963 | 0.07938147 | 60.5962143 | -11.4081326 | 3.94838738 | 0.76880455 | -65.1722107 |
| 17 | 0.00701295 | -14.1854763 | 12.3908243 | 0.18774667 | -0.00436306 | 57.9306297 | -13.1986074 | 4.74779892 | 0.30808449 | -34.9902878 |
| 18 | 0.00832732 | -36.45578 | 10.2242756 | 0.68292999 | -0.0025034 | 56.9240685 | -11.6564274 | 4.19603014 | 0.55263042 | -76.4900818 |
| 19 | 0.0078885 | -25.0493393 | 11.9028482 | 0.38271216 | 0.04469156 | 59.0658188 | -14.0948715 | 4.78071404 | 0.74496269 | -62.5579987 |
| 20 | 0.00694192 | -37.0021629 | 10.3920546 | 0.3244428 | 0.02560616 | 59.8669701 | -18.084301 | 4.34246731 | 0.7576704 | -74.4161453 |
| 21 | 0.00921759 | -22.6706677 | 10.0891962 | 0.18489401 | 0.01705885 | 52.3628388 | -19.6199436 | 4.26865435 | 0.22367835 | -61.7439003 |
| 22 | 0.00790728 | -23.791008 | 9.56186867 | 0.1555391 | 0.07655025 | 50.4799957 | -11.3025703 | 3.8517766 | 0.35709143 | -61.2015839 |
| 23 | 0.00854872 | -19.780016 | 9.87173462 | 0.05144122 | 0.03646612 | 51.209652 | -10.0862837 | 3.99637604 | 0.4748106 | -57.1599731 |
| 24 | 0.00751438 | -38.3709221 | 10.7356482 | 0.24131015 | 0.08898974 | 50.5687408 | -13.4689474 | 4.65365934 | 0.21338463 | -76.0257568 |
| 25 | 0.00645537 | -30.6060429 | 11.0509977 | 0.17165265 | 0.06034374 | 48.698967 | -10.0966969 | 4.83966637 | 0.40872097 | -66.5546494 |
| 26 | 0.00719603 | -40.3394165 | 11.6723852 | 0.34742418 | 0.09553433 | 59.3356476 | -14.7389584 | 4.90396929 | 0.55992603 | -70.8819428 |
| 27 | 0.00702007 | -38.9499054 | 13.2120018 | 0.46227497 | 0.10354519 | 59.4687805 | -18.077755 | 5.91656542 | 0.38269758 | -79.5775909 |
| 28 | 0.00775712 | -29.5123081 | 11.3388929 | 0.29731908 | 0.02176762 | 56.6474266 | -8.92970657 | 4.57059574 | 0.54498315 | -56.6500168 |
| 29 | 0.0071245 | -42.3976822 | 11.6524754 | 0.46155492 | 0.06769896 | 56.8693199 | -13.7477684 | 4.94017983 | 0.35100579 | -71.6952286 |
| 30 | 0.00674599 | -36.6877441 | 11.4117308 | 0.44540685 | 0.07339716 | 57.8977051 | -15.3334703 | 4.85248852 | 0.53168535 | -67.4015427 |
| 31 | 0.00602782 | -20.4141827 | 10.182312 | 0.0263387 | 0.02570152 | 53.5052948 | -12.5340109 | 4.08831024 | 0.32427311 | -67.1927795 |
| 32 | 0.00506172 | -13.887598 | 9.61187553 | 0.01195539 | 0.02980232 | 51.0796318 | -15.8984766 | 4.09002447 | 0.1627326 | -43.3672791 |
| 33 | 0.00905443 | -18.2887363 | 11.2815332 | 0.07950179 | 0.05142689 | 53.5173454 | -11.1894226 | 4.826478 | 0.46553612 | -54.3285713 |
| 34 | 0.00730302 | -20.6326866 | 10.2296324 | 0.04970667 | 0.04174709 | 51.8707466 | -8.00439835 | 4.00646591 | 0.52154064 | -56.0461807 |
| 35 | 0.00825208 | -23.3029213 | 9.39056015 | 0.08676148 | 0.06700754 | 50.0003891 | -7.38489437 | 3.80190754 | 0.36239624 | -61.4822311 |
| 36 | 0.00818445 | -40.4951096 | 9.63406658 | 0.3117418 | 0.03395081 | 55.1280975 | -5.93558693 | 3.96927214 | 0.54271221 | -64.0684738 |
| 37 | 0.00890919 | -35.8598785 | 9.51150894 | 0.3609342 | 0.07866621 | 52.7470398 | -5.70719338 | 3.86124897 | 0.37696362 | -59.6508446 |
| 38 | 0.00772072 | -30.7720642 | 9.39520741 | 0.27040088 | 0.02264381 | 55.7140465 | -4.61995554 | 3.68619156 | 0.51891804 | -58.1247978 |
| 39 | 0.00796846 | -32.2115593 | 9.59149456 | 0.31968129 | 0.032866 | 57.3691902 | -7.46831608 | 3.69912267 | 0.67124367 | -52.8166847 |
| 40 | 0.00880314 | -27.6498375 | 8.4673214 | 0.24559036 | 0.01422167 | 55.0573654 | -2.76745009 | 3.3764019 | 0.47755241 | -49.079258 |

| ID | curvature_std_ | curvature_hips | d_width_midb | d_speed_head | length_w_forw | width_tail_bas | curvature_tail_ | d_width_head |
|---|---|---|---|---|---|---|---|---|
| 1 | 0.00144964 | 0.00569009 | 0 | 5.56507492 | 63.9310913 | 58.0712128 | 0.00088373 | 14.3136864 |
| 2 | 0.00156179 | 0.00512433 | -0.01242161 | -4.15572166 | 24.6695557 | 54.4522934 | 0.0012789 | 13.6348248 |
| 3 | 0.00091757 | 0.00495579 | 0.00314713 | 4.27274609 | 32.6384888 | 56.256958 | 0.00108915 | 16.9117451 |
| 4 | 0.0014744 | 0.00423474 | 0 | 0.7144897 | 58.0741577 | 57.4646492 | 0.00097536 | 14.4190006 |
| 5 | 0.00123598 | 0.00497524 | 0 | -3.22668791 | 46.291626 | 59.2742539 | 0.00124113 | 13.002265 |
| 6 | 0.00149965 | 0.00411975 | 0 | 8.02427959 | 34.975708 | 57.2210579 | 0.00081827 | 15.9713154 |
| 7 | 0.00149229 | 0.00366 | -0.00216365 | 10.7272301 | 28.3594971 | 59.893837 | 6.71E-05 | 16.2540073 |
| 8 | 0.00158924 | 0.00404638 | -0.02013445 | 11.5360584 | 73.6039429 | 52.6373749 | 0.00067708 | 14.7845268 |
| 9 | 0.0014323 | 0.00408698 | 0 | 2.51075459 | 69.2919922 | 57.0907669 | 0.00085222 | 16.2391815 |
| 10 | 0.00158304 | 0.0039436 | -0.00581741 | 8.61126232 | 35.755188 | 54.2793045 | 0.00089806 | 16.0691013 |
| 11 | 0.00115926 | 0.00259666 | 0 | 8.85044289 | 130.638367 | 59.7590179 | 0.00064865 | 13.3533239 |
| 12 | 0.0011059 | 0.00542315 | 0 | 10.7608585 | 28.1690674 | 54.9783897 | 0.00066015 | 12.9448681 |
| 13 | 0.00126559 | 0.00471061 | 0 | -0.19774908 | 72.6928101 | 60.3871803 | 0.00097941 | 14.0734434 |
| 14 | 0.00125252 | 0.00480328 | -0.0161171 | 3.47697997 | 44.3190308 | 58.1253853 | 0.00104681 | 13.9901009 |
| 15 | 0.0012097 | 0.00438345 | 0 | 5.57131958 | 64.0932007 | 57.4173355 | 0.00086625 | 14.062891 |
| 16 | 0.00130863 | 0.00349494 | 0 | 6.65072441 | 47.5941162 | 66.4428635 | 0.00066806 | 15.7809858 |
| 17 | 0.00124927 | 0.00415489 | 0 | 3.01165581 | 175.305664 | 64.9026489 | 0.00044897 | 17.7615356 |
| 18 | 0.00129638 | 0.00412617 | 0.00207424 | 10.2876844 | 62.3444824 | 62.9248238 | 0.00092279 | 15.047431 |
| 19 | 0.00130882 | 0.00479193 | 0.01937151 | 15.1573381 | 89.3963623 | 64.9952164 | 0.0009085 | 16.9440155 |
| 20 | 0.00134988 | 0.00324694 | -0.01928806 | 12.7159376 | 97.8947754 | 64.9963303 | 0.00068723 | 15.2151403 |
| 21 | 0.00133846 | 0.00464308 | -0.00789166 | 2.90414023 | 51.6884155 | 57.5023727 | 0.00108394 | 13.1469965 |
| 22 | 0.00126254 | 0.00514225 | -0.00354052 | 1.23659146 | 97.9023438 | 55.9341278 | 0.00071847 | 12.586957 |
| 23 | 0.00142191 | 0.00500567 | 0 | 0.23239313 | 55.8968506 | 58.289238 | 0.00074569 | 14.5477085 |
| 24 | 0.00133061 | 0.00432834 | -0.03156662 | 5.37773418 | 35.7930908 | 55.0744934 | 0.00073429 | 13.767767 |
| 25 | 0.0013093 | 0.00422373 | 0 | 5.12134552 | 139.19043 | 52.4106674 | 0.00062491 | 12.5814552 |
| 26 | 0.00134332 | 0.00316845 | -0.00705719 | 9.02217865 | 58.4341431 | 64.3990479 | 0.00072092 | 15.1521683 |
| 27 | 0.00143893 | 0.0036614 | -0.00905991 | 0.49749929 | 37.3220215 | 64.346817 | 0.00079524 | 16.0169373 |
| 28 | 0.0012173 | 0.0034252 | 0 | 10.604723 | 81.0148926 | 61.2587509 | 0.00068356 | 13.9461012 |
| 29 | 0.00137885 | 0.00339563 | -0.02017021 | 7.59275436 | 47.201294 | 62.9562569 | 0.00076462 | 15.1464348 |
| 30 | 0.00134625 | 0.00318856 | -0.01399517 | 11.4823141 | 48.184082 | 61.6558456 | 0.0008836 | 14.4653912 |
| 31 | 0.00123337 | 0.00306419 | 0 | -5.73724842 | 81.7424316 | 59.8791275 | 0.0004502 | 14.5641565 |
| 32 | 0.00097813 | 0.00382148 | 0 | -15.7894678 | 32.7747192 | 53.5337372 | 0.00038081 | 12.9984856 |
| 33 | 0.00135233 | 0.00402745 | 0 | 6.78534317 | 109.459961 | 61.879715 | 0.00056766 | 16.5111466 |
| 34 | 0.00118952 | 0.00308796 | 0 | 7.2589159 | 25.1643066 | 59.1564446 | 0.00063284 | 13.6837483 |
| 35 | 0.00132717 | 0.00407129 | 0 | 8.46783257 | 72.0571899 | 56.390564 | 0.00088873 | 15.3626328 |
| 36 | 0.00119186 | 0.00351639 | -0.02143383 | 8.93204403 | 62.6177979 | 59.6837692 | 0.00099527 | 14.9885864 |
| 37 | 0.00114659 | 0.00400528 | -0.01423359 | 5.59883022 | 78.8492432 | 57.6843681 | 0.00123911 | 14.091774 |
| 38 | 0.00119053 | 0.00372401 | -0.00993013 | 1.19618058 | 31.335083 | 59.8468399 | 0.0007903 | 14.7245646 |
| 39 | 0.00109204 | 0.00335541 | -0.02142191 | 2.06378794 | 60.3265991 | 61.6354523 | 0.00090133 | 15.31145 |
| 40 | 0.00113739 | 0.00367217 | -0.00911951 | 3.75691271 | 26.4476929 | 58.8138504 | 0.00084905 | 13.6202307 |

| d_curvature_st | d_width_head | width_head_b | d_length_w_b | d_curvature_st | motion_mode | d_curvature_st | d_length_w_b | curvature_std_ | curvature_std_ | rel_to_neck_ra | curvature_std_ | d_path_curvat | curvature_hea | rel_to_body_r | rel_to_neck_r | d_rel_to_body | width_head_b | curvature_me | curvature_mid | d_rel_to_head | length_w_back | d_rel_to_neck | d_ang_vel_mi | d_curvature_r | curvature_std_ | width_tail_ba | rel_to_body_r | ang_vel_head_ | curvature_std_ | d_rel_to_head | path_curvatur | speed_w_forw | d_rel_to_body | d_rel_to_body | d_ang_vel_hip | d_rel_to_body | width_midbod | d_curvature_n | d_quirkiness_ | speed_hips_w | d_curvature_h | curvature_me | d_curvature_m | ang_vel_tail_b | d_width_head | path_transit_t | path_curvatur | speed_w_forw | width_midbod | d_curvature_n | d_curvature_n | rel_to_hips_ra | ang_vel_head_ | d_curvature_n | curvature_std_ | ang_vel_head_ | curvature_std_ | d_curvature_st | speed_tail_ba | ang_vel_tail_b | d_rel_to_body | rel_to_tail_ba | d_ang_vel_mi | curvature_std_ | width_midbod | d_rel_to_body | curvature_std_ | d_rel_to_body | d_curvature_h | ang_vel_head_ | d_rel_to_body | d_rel_to_body | d_curvature_n | width_tail_ba | curvature_std_ | d_curvature_n | curvature_tail_ | curvature_std_ | ang_vel_head_ | speed_head_ti | d_curvature_h | ang_vel_head_ | d_curvature_h | d_rel_to_body | d_rel_to_tail_t | minor_axis_10 | speed_head_ti | d_rel_to_body | d_curvature_m | rel_to_tail_ba | rel_to_neck_ra | d_curvature_n | curvature_hips | d_major_axis_ | d_ang_vel_hea | curvature_std_ | rel_to_tail_ba | curvature_mid | speed_head_ti | curvature_std_ | width_midbod | d_ang_vel_mi | d_curvature_h | d_length_w_b | d_curvature_n | rel_to_body_r | rel_to_neck_r |
|---|---|---|---|---|---|---|---|---|---|---|---|---|---|---|---|---|---|---|---|---|---|---|---|---|---|---|---|---|---|---|---|---|---|---|---|---|---|---|---|---|---|---|---|---|---|---|---|---|---|---|---|---|---|---|---|---|---|---|---|---|---|---|---|---|---|---|---|---|---|---|---|---|---|---|---|---|---|---|---|---|---|---|---|---|---|---|---|---|---|---|---|---|---|---|---|---|---|---|---|---|---|---|---|---|---|---|---|
| 0.00011129 | -0.11261702 | 45.5632706 | 47.8792648 | 0.00399846 | 3.7 | 0.00504849 | -53.3395233 | 0.00434192 | 0.00580037 | -23.1103725 | 0.00191485 | 0.28743056 | 0.00479377 | 0.03952482 | -2.05931759 | -103.468681 | 53.4700775 | 0.01110816 | 0.00459023 | -1.10911012 | 51.817749 | -2.30974937 | 0.75328624 | 0.00213832 | 0.0019094 | 69.348259 | 0.31191748 | 0.0260989 | 0.00419914 | -43.501152 | 0.00054207 | 82.3128586 | 56.9042702 | 4.83074951 | 1.77685988 | 0.02660546 | 69.3134842 | 0.00029164 | 0.00020721 | 21.3567562 | 0.00652694 | 0.00488719 | 0.00305095 | 0.67837977 | 0.01817942 | 25.608 | 0.00023431 | 48.4566727 | 70.9925156 | 0.00039487 | 0.0049347 | 13.9784622 | 0.75493085 | 0.00102213 | 0.00237311 | 0.17480356 | 0.0018715 | 0.00023585 | 60.127121 | 0.2372492 | 107.892136 | -0.02307579 | 0.18388048 | 0.0023367 | 82.9551163 | -0.38099295 | 0.00082731 | 111.883797 | 0.00682201 | 0.04258753 | 0.02583838 | 0.02225816 | 0.00152362 | 72.1851883 | 0.00550124 | 0.00033382 | 0.01156747 | 0.00350589 | 0.2945022 | 45.0431595 | 0.00109672 | 1.0571568 | 0.00034529 | -59.899128 | -35.7189255 | 121.229157 | -39.7523308 | 0.47930744 | 0.0007936 | 4.49587774 | 22.3280678 | 0.00184337 | 0.0020801 | 67.8127747 | 3.3310523 | 0.00221052 | 5.98940849 | 0.00484217 | -87.8849793 | 0.00319014 | 75.615181 | 0.30597353 | 0.00461015 | 45.0428276 | 0.00569669 | -5.43262672 | 24.2975121 |
| 0.00010743 | -0.08677244 | 47.2093849 | 46.7148056 | 0.00328007 | 3.8 | 0.00610012 | -46.4970017 | 0.00404661 | 0.00563829 | -36.377121 | 0.0014954 | 0.99746627 | 0.00563607 | 0.08178912 | 0.24389246 | -86.3935089 | 53.113163 | 0.00903382 | 0.00392778 | -2.74504757 | 93.7081299 | -13.833458 | 0.65414602 | 0.00348241 | 0.00273945 | 64.4559097 | 0.4173013 | 0.03583617 | 0.00402039 | -52.2668152 | 0.00042978 | 85.5243683 | 53.1291122 | 0.68989414 | 1.21993792 | 0.02331788 | 66.7913208 | 0.00031655 | 0.00011558 | 30.027092 | 0.00728657 | 0.00491468 | 0.00466737 | 0.61211586 | 0.03883243 | 23.398 | 0.00018444 | 45.4683762 | 69.7328491 | 0.00036862 | 0.00623428 | 11.353714 | 1.22346461 | 0.00176928 | 0.00253012 | 0.22169836 | 0.00154619 | 0.00026028 | 55.8090744 | 0.20636773 | 88.2030411 | 0.04273238 | 0.1727958 | 0.00233456 | 76.5040283 | -0.6057452 | 0.00094049 | 89.2000656 | 0.0086778 | 0.06780779 | 0.01806842 | 0.01935961 | 0.00157473 | 69.8469238 | 0.00701761 | 0.00036921 | 0.0082537 | 0.00447454 | 0.4204573 | 70.8243408 | 0.00197972 | 1.30434632 | 0.00035023 | -51.734678 | -25.6866436 | 102.795822 | -36.8266449 | 0.38462153 | 0.001143 | 4.49244976 | 38.5770531 | 0.00295049 | 0.00083639 | 79.4308167 | 3.9604578 | 0.00249278 | 5.27763128 | 0.00418335 | -109.929588 | 0.00336378 | 71.3580933 | 0.27638343 | 0.00483914 | 42.5665512 | 0.00719803 | -3.51960492 | 37.2042465 |
| 0.00010997 | 0.02678633 | 51.7535324 | 45.4159927 | 0.00379949 | 3.78 | 0.00437089 | -53.7580681 | 0.00474217 | 0.0062542 | -27.9959221 | 0.00161586 | 0.48951697 | 0.00619738 | 0.0360227 | -1.7821455 | -82.3662567 | 56.0982513 | 0.00962521 | 0.00497505 | -1.22325325 | 73.2894898 | -7.64927578 | 0.90689868 | 0.00210308 | 0.00238877 | 78.0809631 | 0.01254073 | 0.02349025 | 0.00397297 | -47.1054115 | 0.00074579 | 75.5221405 | 51.6713753 | 2.24929166 | 1.02734768 | 0.02884079 | 70.4468765 | 0.00046829 | 3.63E-05 | 24.5546265 | 0.0060612 | 0.00456989 | 0.00404131 | 0.55712158 | 0.04742146 | 28.982 | 0.00037222 | 32.1714325 | 70.7807007 | 0.00047819 | 0.00574092 | 12.3726759 | 0.93865287 | 0.00125906 | 0.00177798 | 0.14808363 | 0.00184729 | 0.00022532 | 40.2302284 | 0.26126984 | 86.8024902 | -0.03777469 | 0.2123045 | 0.00245284 | 79.1829758 | -0.20617355 | 0.00090889 | 84.8883667 | 0.00621543 | 0.07889149 | 0.02326381 | 0.01697479 | 0.00133419 | 73.2761612 | 0.00546784 | 0.00037807 | 0.01089166 | 0.00325594 | 0.45246714 | 72.5015183 | 0.00151064 | 1.00553405 | 0.00032888 | -51.838501 | -26.7400532 | 48.7406006 | 3.62858677 | 0.45020855 | 0.00085293 | 4.45063591 | 30.0526753 | 0.00213404 | 0.00119596 | 86.0695801 | 3.98010707 | 0.00216859 | 5.50846624 | 0.0045326 | -102.15242 | 0.00266864 | 76.8328781 | 0.38593203 | 0.00430583 | 39.0218506 | 0.00499406 | -6.99591732 | 27.713398 |
| 9.10E-05 | -0.2415061 | 50.3107376 | 43.7810555 | 0.00247226 | 5.32 | 0.0053921 | -47.2179642 | 0.00401296 | 0.00562107 | -27.2245731 | 0.00245699 | 2.14726424 | 0.00597277 | -0.00315677 | -0.66096562 | -91.8559418 | 55.8581581 | 0.00917715 | 0.00293349 | -2.15378499 | 50.8965454 | -3.08917284 | 0.46441117 | 0.00200071 | 0.00202335 | 63.6423149 | 0.21935508 | 0.02988209 | 0.00459025 | -40.8666687 | 0.00043122 | 44.937542 | 49.3445511 | 1.44108069 | 0.85651213 | 0.01339567 | 70.6107254 | 0.00021227 | 0.00017042 | 30.7868748 | 0.00538455 | 0.00363894 | 0.0036359 | 0.51825094 | 0.03343821 | 33.248 | 0.00027022 | 33.9229736 | 70.5221024 | 0.00034636 | 0.00340281 | 9.90403366 | 0.83523035 | 0.00091423 | 0.00196778 | 0.18664408 | 0.00215092 | 0.00019177 | 35.9489517 | 0.14056115 | 97.1619339 | -0.0104791 | 0.15018308 | 0.00170154 | 73.4188385 | -0.5356546 | 0.00068367 | 99.7433548 | 0.00650634 | 0.03767441 | 0.01530167 | 0.01959536 | 0.00097631 | 63.9688263 | 0.00536284 | 0.00028261 | 0.00771158 | 0.00340303 | 0.27039716 | 33.2384338 | 0.00129214 | 1.05851412 | 0.00032373 | -47.5206223 | -25.9281979 | 112.94072 | -43.1445236 | 0.37399709 | 0.00075491 | 4.67696095 | 26.6114883 | 0.00173498 | 0.00069671 | 60.9483719 | 3.12444067 | 0.00196362 | 4.68651772 | 0.00391687 | -77.3775253 | 0.00325772 | 73.2845001 | 0.21802174 | 0.00360657 | 38.4511108 | 0.00532598 | -2.7251358 | 24.9487152 |
| 9.63E-05 | 0.10843277 | 49.5508041 | 45.4081726 | 0.00186632 | 4.6 | 0.00490734 | -47.4881401 | 0.00334119 | 0.00543513 | -21.1819992 | 0.00183244 | 0.65528548 | 0.00477539 | 0.18720658 | -0.19010158 | -100.69561 | 54.7227249 | 0.01400444 | 0.00327198 | -2.14413285 | 121.994446 | -4.15611458 | 0.55220085 | 0.00178547 | 0.00207493 | 65.3033829 | 0.62025034 | 0.03501316 | 0.00406766 | -41.2386589 | 0.00025922 | 52.9290314 | 51.4121552 | -1.18538523 | 1.26091981 | 0.01760576 | 70.1871719 | 0.00022406 | 0.00044395 | 33.9441147 | 0.00615058 | 0.00735411 | 0.00402762 | 0.66487187 | 0.03961325 | 44.816 | 0.00027259 | 25.5896645 | 70.3562088 | 0.00033482 | 0.00394288 | 10.873806 | 0.55611241 | 0.00058731 | 0.00190726 | 0.21267465 | 0.00224812 | 0.00017072 | 31.7990208 | 0.14910452 | 97.925705 | 0.04178701 | 0.17687072 | 0.00153373 | 73.2675934 | -0.6894123 | 0.00057276 | 99.3005524 | 0.00680887 | 0.04472848 | 0.01678938 | 0.02002561 | 0.00118554 | 68.4868317 | 0.00547628 | 0.00030702 | 0.00725641 | 0.00345889 | 0.24655725 | 44.1755028 | 0.0007446 | 1.08211577 | 0.00037224 | -52.2655335 | -27.4093533 | 121.387093 | -20.2433453 | 0.26634842 | 0.00094809 | 4.77450085 | 22.0634232 | 0.00210673 | 0.00105344 | 62.3960495 | 2.56220555 | 0.00210994 | 5.45352268 | 0.00385662 | -92.2907867 | 0.00270371 | 72.0937042 | 0.25715536 | 0.00421457 | 40.0435104 | 0.00572158 | -3.48358512 | 20.5863876 |
| 9.09E-05 | -0.24102926 | 52.4139061 | 36.3254433 | 0.00297732 | 4.56 | 0.00539261 | -40.4760094 | 0.0037681 | 0.00516237 | -47.1195717 | 0.00152274 | 1.67134285 | 0.00634223 | 0.43330729 | 0.88045263 | -87.0497437 | 57.3948898 | 0.00918342 | 0.00419978 | -3.11029053 | 57.9645386 | -13.560606 | 0.95986146 | 0.00163221 | 0.00191858 | 64.4857864 | 0.95175368 | 0.03917657 | 0.00355862 | -46.9276695 | 0.00016199 | 146.041367 | 46.5883064 | 1.19927418 | 1.65168107 | 0.02069003 | 73.7251511 | 0.00026502 | 0.00034288 | 33.1825218 | 0.00752174 | 0.00446141 | 0.00629733 | 0.88198102 | 0.06304979 | 18.384 | 0.0001603 | 56.7406921 | 73.8408203 | 0.00034923 | 0.00660096 | 11.0031605 | 1.28635573 | 0.00151598 | 0.00242644 | 0.26216334 | 0.00137551 | 0.00019098 | 66.9074783 | 0.1884194 | 78.7900696 | 0.09889164 | 0.17583214 | 0.00214423 | 78.2614746 | -0.24247916 | 0.00094108 | 78.4806061 | 0.00561437 | 0.0747209 | 0.01854429 | 0.01730008 | 0.00255862 | 64.1615295 | 0.00509835 | 0.00033019 | 0.00715875 | 0.00372161 | 0.45214176 | 75.9359207 | 0.00172206 | 1.1130867 | 0.00032834 | -46.3621864 | -22.3182182 | 93.0022354 | 20.6841869 | 0.55374795 | 0.00174339 | 4.43748665 | 47.3853912 | 0.00478799 | 0.00070676 | 84.8591766 | 3.98594141 | 0.00145584 | 5.35759306 | 0.00349437 | -157.478241 | 0.00311076 | 77.8600998 | 0.34048665 | 0.00606641 | 31.545414 | 0.00478934 | -4.54565716 | 46.5797043 |
| 8.94E-05 | -0.19958615 | 29.8994846 | 34.6870346 | 0.00360436 | 3.88 | 0.00495311 | -37.1388245 | 0.00322265 | 0.0044795 | -46.5860138 | 0.0012413 | 2.69259024 | 0.00576505 | 0.27079532 | 0.45981622 | -82.6987228 | 59.4181786 | 0.00813701 | 0.00391032 | -1.75043392 | 22.7069702 | -12.8216677 | 0.95071781 | 0.00176804 | 0.00128891 | 64.5852966 | 0.50549978 | 0.02297339 | 0.00305389 | -49.167553 | 0.00019787 | 189.062286 | 53.4171066 | 0.18474288 | 1.47501898 | 0.02439056 | 75.2059097 | 0.00028443 | 0.00011141 | 35.3064079 | 0.00640847 | 0.00394182 | 0.00533897 | 0.77794969 | 0.01032054 | 13.908 | 0.00021016 | 105.585533 | 74.1897202 | 0.00033691 | 0.00640255 | 11.9431162 | 1.32992315 | 0.00155183 | 0.00176496 | 0.18307231 | 0.00140042 | 0.00019936 | 119.857094 | 0.21902528 | 82.8968201 | 0.06348537 | 0.16732477 | 0.00217315 | 82.5455933 | -0.972399 | 0.00088259 | 82.2629089 | 0.00628358 | 0.07649706 | 0.01813279 | 0.0118689 | 0.00273368 | 65.3323975 | 0.00460966 | 0.00027759 | 0.00672505 | 0.00302291 | 0.50240916 | 86.2119064 | 0.00177153 | 1.13253903 | 0.00025444 | -53.0470543 | -22.6414318 | 97.5634613 | 20.2511177 | 0.51625222 | 0.00145181 | 3.99543881 | 45.891964 | 0.00477705 | 0.00029726 | 94.4411316 | 4.47371197 | 0.00259484 | 5.00908375 | 0.00410769 | -163.389023 | 0.00274288 | 80.4734879 | 0.3538177 | 0.00597868 | 31.7828426 | 0.00538314 | -3.10835743 | 47.1051254 |
| 0.00012068 | -0.24338961 | 49.836628 | 35.8839874 | 0.00273213 | 3.28 | 0.00627556 | -38.3052788 | 0.00380917 | 0.00539674 | -41.5908737 | 0.00156112 | 2.62895846 | 0.00596628 | 0.62335426 | 1.2428534 | -89.6328583 | 55.5116844 | 0.00928805 | 0.00396678 | -4.20148945 | 67.2236328 | -15.2570124 | 1.19471085 | 0.00190286 | 0.00176812 | 60.3834305 | 0.9130683 | 0.05700256 | 0.00363178 | -45.7292557 | 0.00014798 | 97.8239594 | 45.672821 | -0.38091388 | 2.00074077 | 0.02008211 | 68.7556534 | 0.00024136 | 0.00041693 | 30.9260693 | 0.00984825 | 0.00479862 | 0.00851271 | 1.03138053 | 0.04853606 | 12.058 | 0.00027486 | 46.6710968 | 68.9593887 | 0.0004067 | 0.00892747 | 9.50610733 | 1.32375968 | 0.00174918 | 0.00259987 | 0.37290525 | 0.0015815 | 0.00025229 | 57.128624 | 0.15941583 | 76.3532333 | 0.09394098 | 0.26055977 | 0.00210082 | 76.9184113 | -0.56002176 | 0.00087758 | 75.2104492 | 0.0070875 | 0.07579685 | 0.02475224 | 0.02093236 | 0.00321255 | 59.811882 | 0.00546448 | 0.00040052 | 0.00704471 | 0.0038402 | 0.44690293 | 71.6775208 | 0.00202332 | 1.2109009 | 0.00036213 | -44.7299271 | -20.7241478 | 86.1428528 | 15.3012657 | 0.70454878 | 0.00229072 | 4.42249012 | 40.6798554 | 0.00570789 | 0.00066233 | 82.4142456 | 3.91325879 | 0.00157097 | 5.20235682 | 0.0037757 | -183.469345 | 0.00328872 | 76.6968536 | 0.49545422 | 0.00739527 | 32.1224365 | 0.00625234 | -4.77362394 | 40.9976883 |
| 0.00011367 | -0.0649333 | 51.3101616 | 38.0536804 | 0.00353113 | 4.76 | 0.00568654 | -41.503212 | 0.00350332 | 0.0050065 | -49.6923332 | 0.00158241 | 2.24889374 | 0.00591774 | 0.51430237 | -0.24609523 | -109.458603 | 57.2236328 | 0.00893441 | 0.00388054 | -3.6478467 | 132.921082 | -13.548996 | 1.37240791 | 0.00161475 | 0.00192338 | 66.155777 | 0.72432017 | 0.05553084 | 0.00365405 | -50.7148743 | 0.0001503 | 174.477799 | 55.1845436 | -1.06847608 | 2.18806958 | 0.02807403 | 70.8768616 | 0.00032464 | 0.00061683 | 34.1563072 | 0.00992376 | 0.00465277 | 0.00856985 | 1.19511652 | 0.08831024 | 9.398 | 0.00028026 | 101.452118 | 71.2928925 | 0.00041839 | 0.00850983 | 13.2658939 | 1.40257692 | 0.00173572 | 0.00236798 | 0.41063649 | 0.00152476 | 0.00023415 | 111.907379 | 0.26281279 | 90.2834473 | 0.0819632 | 0.23731351 | 0.00196171 | 80.4083481 | -0.76457816 | 0.00077939 | 87.5011368 | 0.00662418 | 0.0950322 | 0.02206992 | 0.02397476 | 0.00436575 | 66.0752869 | 0.00513534 | 0.00038883 | 0.00685971 | 0.00364716 | 0.6087122 | 101.774353 | 0.00200929 | 1.17949629 | 0.00036649 | -57.0553818 | -24.3116112 | 92.5097427 | 28.7889404 | 0.74285388 | 0.00332845 | 5.18890905 | 51.4360962 | 0.0074852 | 0.000675 | 98.7220764 | 4.89806032 | 0.00157819 | 6.50975466 | 0.00347344 | -217.031387 | 0.00294979 | 79.0684204 | 0.49544129 | 0.00902085 | 34.2970352 | 0.00580547 | -7.32234097 | 51.7403259 |
| 0.00011741 | -0.11265278 | 50.578186 | 36.6730957 | 0.00318605 | 4.2 | 0.00652985 | -38.6132851 | 0.00398461 | 0.00538812 | -42.7746582 | 0.00173361 | 2.35651088 | 0.00575552 | 0.45686042 | 0.83060855 | -118.955399 | 56.0772926 | 0.00955422 | 0.00427106 | -4.31321335 | 44.7406006 | -16.2494831 | 1.76517606 | 0.00196259 | 0.00209975 | 60.6040955 | 0.77357113 | 0.05995767 | 0.00395941 | -49.7685356 | 0.00014965 | 118.605919 | 46.868515 | 0.63209504 | 2.76492119 | 0.02275077 | 70.0839081 | 0.00029749 | 0.00044784 | 33.3670883 | 0.01252433 | 0.00457063 | 0.01091494 | 1.42531204 | 0.06294251 | 11.002 | 0.00026969 | 54.8320465 | 70.1215515 | 0.00044658 | 0.01052362 | 10.6005688 | 1.42451298 | 0.00190762 | 0.00256568 | 0.39640337 | 0.00150785 | 0.00026464 | 64.2422791 | 0.20609582 | 79.6465302 | 0.04710245 | 0.26559243 | 0.00222914 | 72.1105576 | -0.85592932 | 0.00098653 | 79.4173584 | 0.00804229 | 0.07702817 | 0.02612088 | 0.02357225 | 0.00382379 | 59.6701737 | 0.00549947 | 0.00041326 | 0.00752128 | 0.00403715 | 0.4835211 | 78.8369904 | 0.00219809 | 1.32121384 | 0.00040112 | -47.0392838 | -22.4957867 | 84.9916763 | 20.6742764 | 0.97224951 | 0.00282345 | 4.93817902 | 43.2996864 | 0.0070146 | 0.00086593 | 84.8497162 | 4.25728035 | 0.00156847 | 6.1316042 | 0.00377229 | -224.104157 | 0.00327779 | 71.8998947 | 0.58326131 | 0.008809 | 34.18853 | 0.0070398 | -6.1700964 | 42.5975685 |
| 8.52E-05 | -0.09896159 | 51.4433289 | 43.367897 | 0.00173042 | 5.56 | 0.00401608 | -44.3351975 | 0.00219503 | 0.00342587 | -21.1901016 | 0.00134479 | 0.9712882 | 0.0038281 | -0.05349882 | -1.09067631 | -88.6492539 | 58.0453835 | 0.00901407 | 0.00246654 | -1.52899039 | 76.1708374 | -2.86508298 | 0.50598663 | 0.00151403 | 0.00142979 | 63.967926 | -0.03791989 | 0.0252396 | 0.00304608 | -39.4477463 | 0.00038882 | 52.5792923 | 52.0987167 | 0.87293357 | 0.7028538 | 0.01343966 | 80.7392273 | 0.00019435 | 7.76E-05 | 33.7735558 | 0.0040529 | 0.00437683 | 0.00240962 | 0.44621506 | 0.08413792 | 24.056 | 0.00031393 | 37.2423363 | 76.432991 | 0.00028037 | 0.0031003 | 10.8968754 | 0.70779973 | 0.00069763 | 0.00171675 | 0.16100636 | 0.00120883 | 0.0001652 | 34.7083359 | 0.11860623 | 92.5966644 | -0.02724777 | 0.16782506 | 0.00186821 | 84.4472656 | -0.43304586 | 0.0007017 | 94.717247 | 0.00468001 | 0.0374531 | 0.01487773 | 0.01730031 | 0.00094569 | 66.5796967 | 0.00412324 | 0.00025034 | 0.00686682 | 0.00229298 | 0.2259343 | 39.5219917 | 0.00083444 | 0.94828308 | 0.00025225 | -52.4671478 | -27.1264687 | 102.125961 | -55.6140213 | 0.21481766 | 0.00058382 | 4.40563011 | 22.080616 | 0.00133725 | 0.00041911 | 64.6443863 | 2.4459877 | 0.0017963 | 4.82278585 | 0.00323734 | -77.8581619 | 0.00263655 | 84.4441376 | 0.23025757 | 0.00308282 | 40.7959061 | 0.00395007 | -2.45179915 | 20.8698616 |
| 8.88E-05 | 0.00836253 | 52.3994751 | 45.3931885 | 0.00198214 | 7.64 | 0.00388816 | -46.4145203 | 0.00387842 | 0.0055277 | -18.9293442 | 0.00149507 | 0.18330362 | 0.00342517 | -0.02130036 | 0.1306392 | -88.1236725 | 55.660183 | 0.01008568 | 0.00316974 | -1.85569978 | 50.8949585 | -4.50775242 | 0.44995466 | 0.00152874 | 0.00166608 | 62.877758 | 0.06503673 | 0.0291071 | 0.00363962 | -41.0117836 | 0.00036324 | 51.5587082 | 51.2756348 | 4.08075428 | 0.98733836 | 0.01597309 | 70.4423065 | 0.00021094 | 6.93E-05 | 32.2326088 | 0.00456555 | 0.00586261 | 0.00255875 | 0.45896724 | 0.09183884 | 42.032 | 0.00036455 | 28.3539352 | 75.1269074 | 0.00031385 | 0.00399463 | 10.3082657 | 0.63850904 | 0.00074487 | 0.00193204 | 0.17706773 | 0.00219118 | 0.00020455 | 30.9921112 | 0.14577502 | 93.1307221 | -0.03559441 | 0.13962865 | 0.00238582 | 78.792038 | -0.20746809 | 0.00078979 | 90.3997116 | 0.00534071 | 0.03847801 | 0.01792585 | 0.01804605 | 0.00113109 | 62.5093651 | 0.00501686 | 0.00029705 | 0.00850663 | 0.00321137 | 0.2220901 | 42.1189728 | 0.00089785 | 0.87917543 | 0.00028364 | -52.1661186 | -25.2054863 | 107.645653 | -39.9053459 | 0.30857882 | 0.0007426 | 4.62238646 | 19.2329845 | 0.00173792 | 0.0009043 | 57.9892159 | 2.50485492 | 0.0018679 | 5.1261673 | 0.0042229 | -79.3924866 | 0.00246809 | 75.7949677 | 0.2039884 | 0.00333096 | 44.2475433 | 0.00444781 | -2.07181692 | 18.5461578 |
| 0.00010101 | -0.05575419 | 53.1238213 | 44.3525009 | 0.00228564 | 4.72 | 0.0049321 | -49.6070061 | 0.00441841 | 0.00580682 | -28.6750431 | 0.00140321 | 0.2826784 | 0.00510279 | 0.17900419 | -1.35472763 | -122.919388 | 58.9156494 | 0.01337252 | 0.0042356 | -2.68756318 | 49.4961548 | -2.68332052 | 0.60606492 | 0.00175398 | 0.00148639 | 63.734272 | 0.3574639 | 0.02290524 | 0.00315789 | -43.580986 | 0.000859 | 42.4975815 | 55.1530953 | 4.47959614 | 1.5075388 | 0.02595265 | 72.6555328 | 0.0001895 | 0.00017203 | 25.7026424 | 0.00520122 | 0.00366545 | 0.0021218 | 0.7004987 | 0.02766252 | 68.89 | 0.00043241 | 35.1434937 | 73.6313248 | 0.0004035 | 0.0043636 | 12.1317425 | 1.02022409 | 0.00092616 | 0.00236588 | 0.18324211 | 0.00224022 | 0.00019021 | 29.1543808 | 0.15365505 | 100.265205 | -0.08611683 | 0.19167668 | 0.00210905 | 79.1934662 | -0.62536544 | 0.00068895 | 100.108894 | 0.00665408 | 0.03849753 | 0.02306012 | 0.0296453 | 0.00174019 | 67.6362152 | 0.00711155 | 0.00035351 | 0.01030999 | 0.00479179 | 0.2525152 | 20.8295708 | 0.00133051 | 1.06947386 | 0.00034222 | -54.4924164 | -30.7126904 | 169.56546 | -68.3559265 | 0.43909389 | 0.00066388 | 4.85009861 | 29.6650677 | 0.00151032 | 0.00151118 | 66.1774139 | 3.32329655 | 0.00253952 | 5.983181 | 0.00569543 | -101.411667 | 0.0030034 | 77.262497 | 0.28834528 | 0.00378237 | 37.9162369 | 0.00554454 | -10.0818214 | 27.7305603 |
| 8.31E-05 | -0.24755597 | 52.0225143 | 48.442009 | 0.00201268 | 3.46 | 0.003988 | -49.5711784 | 0.00334746 | 0.00512385 | -31.1587601 | 0.00090004 | 0.65343279 | 0.0049535 | 0.10753363 | -1.41807675 | -124.681648 | 59.6812515 | 0.00977052 | 0.0038733 | -1.26592183 | 30.9784546 | -1.34204352 | 0.56000918 | 0.00152023 | 0.00143617 | 77.0276184 | 0.21346922 | 0.02524181 | 0.00229792 | -49.2102661 | 0.00016895 | 49.9847107 | 55.395752 | 1.71088255 | 1.35751879 | 0.01926744 | 77.137619 | 0.00016714 | 0.00013271 | 34.8007355 | 0.00399214 | 0.00329446 | 0.00183885 | 0.53262675 | 0.02846718 | 46 | 0.00027435 | 42.4974136 | 77.9024887 | 0.0003189 | 0.00370201 | 10.897316 | 0.77617484 | 0.00075278 | 0.00181272 | 0.16026421 | 0.00183839 | 0.00016573 | 38.8037338 | 0.13482156 | 101.456146 | 0.00456111 | 0.15211603 | 0.00196551 | 80.5001373 | -0.01613298 | 0.00074061 | 101.838928 | 0.00480542 | 0.03450501 | 0.01725183 | 0.02172743 | 0.00099543 | 66.3234482 | 0.00553254 | 0.00028099 | 0.00944824 | 0.00303941 | 0.22008453 | 29.7707024 | 0.00088321 | 0.85939914 | 0.00024292 | -55.9871445 | -30.0772114 | 110.549507 | -63.4590721 | 0.31443912 | 0.00053571 | 5.01921988 | 28.6026459 | 0.00121395 | 0.00106746 | 65.1248932 | 2.59534073 | 0.00261631 | 6.6292057 | 0.00359805 | -73.8569031 | 0.00282609 | 80.2333832 | 0.21280059 | 0.00304477 | 53.0156364 | 0.00390139 | -1.0960151 | 29.2263222 |
| 9.60E-05 | 0.06366968 | 51.8950501 | 44.8825913 | 0.00207132 | 3.44 | 0.00448807 | -45.7277603 | 0.00289013 | 0.00422747 | -22.2304516 | 0.00117844 | 0.3709771 | 0.00370867 | -0.01861595 | -0.94330764 | -82.9039536 | 56.9542694 | 0.00835137 | 0.00301274 | -1.61087024 | 91.7001343 | -4.46600437 | 0.5599975 | 0.00164522 | 0.0014952 | 74.6798019 | 0.06683178 | 0.02807915 | 0.00278182 | -44.0625458 | 0.00029895 | 56.5512123 | 52.5722466 | -0.12805817 | 1.12237191 | 0.0154974 | 77.5816345 | 0.00020415 | 2.25E-05 | 24.2854462 | 0.00415651 | 0.00354776 | 0.00235326 | 0.51599044 | 0.06613135 | 27.88 | 0.00026027 | 33.966095 | 78.0377045 | 0.0003323 | 0.00356012 | 10.4352074 | 0.87627107 | 0.00101723 | 0.00164051 | 0.18247287 | 0.00162795 | 0.00017808 | 38.3576431 | 0.14678314 | 87.701889 | -0.00662786 | 0.16847846 | 0.0021127 | 81.4470291 | -0.17960443 | 0.00081971 | 87.2405243 | 0.00546525 | 0.0402786 | 0.01765852 | 0.02040437 | 0.00113071 | 67.8017731 | 0.0047775 | 0.00026213 | 0.0074513 | 0.0030457 | 0.25230959 | 44.652153 | 0.00120053 | 0.96364266 | 0.00027475 | -51.1874924 | -26.5232925 | 141.314224 | -50.8855133 | 0.31002715 | 0.00063446 | 4.86634159 | 23.1037312 | 0.00147098 | 0.00097675 | 68.006134 | 2.80234313 | 0.00208917 | 5.5395627 | 0.00351096 | -82.1087036 | 0.00268514 | 80.3927918 | 0.25542247 | 0.00344143 | 47.1728516 | 0.00458806 | -3.01424074 | 20.9191055 |
| 0.00010942 | -0.14674664 | 62.0507278 | 39.1198578 | 0.00199314 | 3.3 | 0.00439141 | -43.2216339 | 0.00328878 | 0.00457639 | -43.6044426 | 0.00133837 | 1.09796298 | 0.00513041 | 0.70174372 | 1.10645413 | -131.636871 | 67.6111908 | 0.00865209 | 0.00313526 | -3.73545504 | 45.0917969 | -13.4697008 | 1.27716649 | 0.00134829 | 0.00143514 | 73.2549439 | 1.38964868 | 0.03955714 | 0.00316456 | -46.660759 | 0.00017897 | 106.447212 | 47.2303505 | 1.38227868 | 1.75512755 | 0.01873418 | 86.4251556 | 0.00020778 | 0.00043842 | 38.5981827 | 0.006686 | 0.00432215 | 0.00586966 | 1.13184559 | 0.04589558 | 13.064 | 0.00023955 | 49.7104759 | 86.5521851 | 0.0003954 | 0.00745234 | 10.6918755 | 1.05953705 | 0.00103291 | 0.00245254 | 0.28223118 | 0.00140673 | 0.0002206 | 61.0157623 | 0.17641611 | 78.7938538 | 0.1001097 | 0.24022828 | 0.00204367 | 91.2663498 | -0.68067956 | 0.00087826 | 77.2751083 | 0.00523377 | 0.07013787 | 0.02253328 | 0.01811668 | 0.00232535 | 72.8671188 | 0.00454487 | 0.00032112 | 0.00644763 | 0.00317624 | 0.42967758 | 82.4758377 | 0.00118657 | 1.07505059 | 0.00027773 | -48.3373451 | -21.3987007 | 116.80307 | 24.5650959 | 0.6583885 | 0.00133711 | 4.56602001 | 42.4323387 | 0.0033276 | 0.00043179 | 85.4734421 | 3.06168914 | 0.00149849 | 5.81071854 | 0.00330019 | -213.100632 | 0.00268609 | 90.6681061 | 0.48517507 | 0.00434482 | 35.0233879 | 0.00469863 | -7.81784201 | 43.1623421 |
| 8.67E-05 | -0.15230775 | 58.1158104 | 37.5134544 | 0.00481659 | 5.48 | 0.00387775 | -40.9352913 | 0.00299724 | 0.00445324 | -64.0386047 | 0.00146847 | 1.04765081 | 0.00609934 | 0.29980445 | 0.01007829 | -102.801506 | 60.7719803 | 0.00978482 | 0.00423946 | -1.38444567 | 47.8186035 | -11.8844204 | 0.92887777 | 0.0013985 | 0.00112022 | 76.6878967 | 0.55477542 | 0.02433445 | 0.00322771 | -52.8564339 | 0.00031755 | 240.054077 | 102.873505 | 1.23885274 | 1.35540021 | 0.03501576 | 72.462738 | 0.00034293 | 8.97E-05 | 43.0519829 | 0.00573694 | 0.00576255 | 0.00491086 | 0.91490132 | 0.02653599 | 14.476 | 0.00031579 | 166.672485 | 86.5532074 | 0.00033299 | 0.00652776 | 22.3468819 | 1.43218851 | 0.00147477 | 0.0019487 | 0.1668673 | 0.00161588 | 0.00018879 | 193.189987 | 0.41214508 | 122.703407 | 0.07672243 | 0.1781707 | 0.00214324 | 97.5170288 | -0.35237593 | 0.00089397 | 135.570679 | 0.00488057 | 0.10556706 | 0.0181324 | 0.01261174 | 0.00285293 | 73.9933319 | 0.0043863 | 0.00024658 | 0.00732268 | 0.00204054 | 0.77379274 | 147.966995 | 0.00170972 | 1.06747317 | 0.00022997 | -112.149376 | -29.5609131 | 68.943634 | 32.0136147 | 0.51242071 | 0.0008746 | 4.27236795 | 64.3550644 | 0.00460786 | 0.0002108 | 127.300934 | 7.77368259 | 0.00219178 | 5.94699526 | 0.00303636 | -174.647232 | 0.00273305 | 87.2735291 | 0.34309751 | 0.00560859 | 32.7907982 | 0.00436357 | -6.23171091 | 68.055191 |
| 9.69E-05 | -0.27920008 | 57.7373238 | 39.8330383 | 0.00252795 | 4.44 | 0.00438493 | -43.3060837 | 0.00353345 | 0.00518936 | -46.305687 | 0.001608 | 1.15669298 | 0.00520858 | 0.669536 | 1.45227432 | -102.193573 | 63.1184845 | 0.00978328 | 0.00430251 | -3.14592147 | 112.452759 | -13.150094 | 0.97591937 | 0.00140936 | 0.00175272 | 78.2850494 | 1.47873855 | 0.03411602 | 0.00324704 | -45.8333435 | 0.00020772 | 127.298584 | 52.8802338 | 1.79312742 | 1.50416493 | 0.02086104 | 80.8343124 | 0.00023545 | 0.00034503 | 39.9923935 | 0.00617637 | 0.00474075 | 0.00497717 | 0.8936249 | -0.0054121 | 17.498 | 0.0002252 | 51.006424 | 82.1103897 | 0.00027742 | 0.00631886 | 13.2276077 | 1.15852797 | 0.00117676 | 0.00236607 | 0.23578002 | 0.00151208 | 0.00019045 | 65.9633942 | 0.20197584 | 87.032486 | 0.12004642 | 0.23506521 | 0.00210947 | 92.3488236 | -0.78882211 | 0.00091854 | 90.1541214 | 0.00514139 | 0.06351239 | 0.02229043 | 0.01872996 | 0.00238697 | 77.2352905 | 0.00522546 | 0.00032155 | 0.00734348 | 0.00363282 | 0.40739822 | 75.8450775 | 0.00138629 | 1.01905632 | 0.00029757 | -51.9927216 | -24.38941 | 139.30957 | 17.596159 | 0.52890164 | 0.00133541 | 4.82340813 | 44.9346695 | 0.00361448 | 0.00092562 | 91.4519959 | 3.4271574 | 0.00150202 | 6.07197237 | 0.00376072 | -153.939056 | 0.00272138 | 90.4564972 | 0.40879288 | 0.00477451 | 36.5385017 | 0.00450284 | -7.68261051 | 46.1989822 |
| 0.00010762 | -0.02745986 | 58.5728188 | 38.1766434 | 0.00358462 | 3.16 | 0.00515166 | -40.5671082 | 0.00357345 | 0.00537175 | -55.5993462 | 0.0016636 | 1.35475946 | 0.00550152 | 0.64821738 | -0.05041597 | -131.303711 | 62.795578 | 0.00961471 | 0.00389041 | -5.07434845 | 72.2546387 | -16.1273384 | 1.71919096 | 0.00161551 | 0.00176312 | 71.2106018 | 0.93095136 | 0.0632084 | 0.00344752 | -52.4430389 | 0.00016166 | 187.19014 | 68.6250992 | 0.21779889 | 2.34473681 | 0.03478121 | 79.6440048 | 0.00037998 | 0.00083874 | 40.5891914 | 0.00882655 | 0.00497138 | 0.00798786 | 1.27913785 | 0.1129508 | 8.592 | 0.00026621 | 120.337524 | 80.178215 | 0.0004566 | 0.00928337 | 18.7351914 | 1.37763357 | 0.00159085 | 0.00246646 | 0.45619035 | 0.00181042 | 0.0002305 | 131.071259 | 0.35184178 | 101.090363 | 0.10718224 | 0.25901955 | 0.00222888 | 90.0559235 | -1.33274353 | 0.00095434 | 103.405136 | 0.00675301 | 0.10987537 | 0.02440338 | 0.02476961 | 0.00526647 | 72.132782 | 0.00522945 | 0.00032061 | 0.00713035 | 0.00358472 | 0.71394318 | 129.845444 | 0.00182647 | 1.31302083 | 0.00030713 | -70.7799225 | -26.6341782 | 122.984116 | 38.4453354 | 0.9544512 | 0.00384211 | 5.70157623 | 57.7955284 | 0.0080582 | 0.00071881 | 118.420982 | 5.55319738 | 0.00169885 | 6.06103087 | 0.00332247 | -235.612701 | 0.00267788 | 84.3097992 | 0.63690138 | 0.00922021 | 33.4133415 | 0.00602249 | -8.77995682 | 57.5660553 |
| 8.80E-05 | -0.17244816 | 59.872139 | 39.6853867 | 0.00188699 | 3.58 | 0.00409369 | -43.9068375 | 0.00295088 | 0.00423104 | -45.6398163 | 0.00122422 | 1.92945814 | 0.00472618 | 0.36260229 | 2.00308466 | -131.413727 | 65.5623169 | 0.0085962 | 0.00342265 | -2.1414144 | 91.3764648 | -13.4444771 | 1.3456775 | 0.00134603 | 0.001049 | 72.6077042 | 0.85466105 | 0.02884018 | 0.00270693 | -51.1456108 | 0.00021304 | 108.604912 | 51.3202858 | -0.32012811 | 1.80142736 | 0.02052464 | 84.3188782 | 0.00019035 | 0.00052276 | 43.631443 | 0.00670196 | 0.00469493 | 0.00605034 | 1.17075622 | 0.07758141 | 11.14 | 0.00031907 | 55.1384201 | 85.3542328 | 0.00036182 | 0.007921 | 11.4156227 | 1.10466576 | 0.00111592 | 0.00212474 | 0.22416475 | 0.00134866 | 0.00019238 | 66.078186 | 0.17390949 | 83.733017 | 0.04820534 | 0.26432163 | 0.00197426 | 89.5271378 | -0.72456676 | 0.00085762 | 84.2786102 | 0.00562461 | 0.06959403 | 0.025099 | 0.01622653 | 0.00221826 | 72.9194794 | 0.00437483 | 0.00030027 | 0.00651415 | 0.00284944 | 0.42641339 | 88.7440033 | 0.00125914 | 1.17079949 | 0.0002569 | -50.6149406 | -22.2795067 | 132.411713 | 19.1597004 | 0.70830178 | 0.00125445 | 4.53050423 | 44.8889198 | 0.00327705 | 0.00030579 | 95.4497833 | 3.33400726 | 0.00161845 | 6.29168749 | 0.00311805 | -226.406586 | 0.00274084 | 89.061882 | 0.5653137 | 0.0041425 | 35.7059593 | 0.004956 | -7.80593205 | 45.1929245 |
| 9.63E-05 | -0.09388924 | 53.9822426 | 43.8673515 | 0.0020968 | 4.76 | 0.00535684 | -45.2501679 | 0.00341607 | 0.00549963 | -25.5873184 | 0.00118763 | 0.67406201 | 0.00407652 | 0.12556472 | -0.00614538 | -85.832016 | 58.3666725 | 0.00897459 | 0.00346013 | -2.3324225 | 45.5332642 | -5.02447128 | 0.56699169 | 0.00204943 | 0.00166243 | 70.1614227 | 0.42384258 | 0.03261807 | 0.00306773 | -52.8600121 | 0.00032571 | 52.137001 | 47.9073792 | 0.61893183 | 1.00624609 | 0.01855583 | 76.3977509 | 0.00020831 | 0.00012875 | 33.4388771 | 0.00492139 | 0.00421556 | 0.00255489 | 0.53826886 | 0.00394583 | 34.83 | 0.0002931 | 38.0526009 | 77.9323807 | 0.00033179 | 0.0040434 | 10.2026901 | 0.95842785 | 0.00112713 | 0.00199221 | 0.20760256 | 0.00196611 | 0.00019314 | 42.0039902 | 0.14802748 | 89.1835175 | 0.02016728 | 0.15867449 | 0.00198609 | 79.9572449 | -0.32414651 | 0.00079745 | 85.3419647 | 0.00703972 | 0.03915367 | 0.01791493 | 0.02174071 | 0.0012785 | 63.8727379 | 0.00590731 | 0.00028053 | 0.00884047 | 0.00333246 | 0.2406179 | 41.6674881 | 0.00132686 | 1.22668886 | 0.00027714 | -45.3849564 | -27.4284458 | 118.02964 | -48.0244637 | 0.32153758 | 0.00072936 | 4.86125898 | 26.9942913 | 0.00166236 | 0.00086842 | 62.8566742 | 2.68549538 | 0.00237983 | 5.18192244 | 0.00396298 | -93.2541657 | 0.00295935 | 79.8330689 | 0.2586112 | 0.00403268 | 40.5669518 | 0.00583261 | -3.12952685 | 24.0688591 |
| 9.40E-05 | -0.24072528 | 51.4228249 | 44.8121185 | 0.00198282 | 5.6 | 0.00520801 | -46.401741 | 0.00362336 | 0.00597876 | -27.3829804 | 0.00139153 | 0.80434674 | 0.00419546 | 0.15700434 | 0.52406424 | -87.7881165 | 57.0416718 | 0.00814001 | 0.00331727 | -2.527879 | 144.47522 | -6.62106943 | 0.70237261 | 0.00202987 | 0.00178829 | 62.482151 | 0.39255798 | 0.03513214 | 0.00325141 | -53.3531303 | 0.00027232 | 52.9096069 | 47.6515427 | -0.16208804 | 1.12275577 | 0.01563985 | 75.6601181 | 0.00019709 | 0.00019627 | 37.210865 | 0.0052454 | 0.00417869 | 0.00316789 | 0.54485065 | 0.0328064 | 30.73 | 0.00028793 | 29.1693802 | 74.2950897 | 0.00037165 | 0.00458214 | 10.0521259 | 0.91368043 | 0.00104785 | 0.00204884 | 0.22243299 | 0.00197069 | 0.00019669 | 35.5283089 | 0.13658413 | 85.0522461 | 0.02608692 | 0.19293642 | 0.0021027 | 78.3033829 | -0.52350146 | 0.00084902 | 83.3162613 | 0.00716941 | 0.04708625 | 0.01882626 | 0.02059082 | 0.00135494 | 61.6790962 | 0.00595552 | 0.00030426 | 0.00648267 | 0.0036221 | 0.27069128 | 48.6018372 | 0.00121723 | 1.19619536 | 0.00028509 | -47.3703728 | -25.3179741 | 109.420631 | -27.939724 | 0.36148676 | 0.0009161 | 4.80088472 | 27.5250511 | 0.00210215 | 0.00073971 | 63.6279831 | 2.72788262 | 0.00207546 | 5.32958365 | 0.00465412 | -104.051277 | 0.00279745 | 79.4921494 | 0.31007725 | 0.00405095 | 41.5902863 | 0.00599195 | -3.40511894 | 25.0581055 |
| 0.00010084 | -0.28116107 | 51.8144684 | 42.813385 | 0.0023518 | 5.36 | 0.00557214 | -43.9885597 | 0.00369675 | 0.00573834 | -26.3622875 | 0.00136769 | 1.06907785 | 0.00500925 | 0.09267683 | 0.24979317 | -85.3032608 | 58.7206879 | 0.00847212 | 0.0035116 | -2.3499701 | 56.6975098 | -4.95928192 | 0.53449804 | 0.00210226 | 0.00168583 | 69.6797638 | 0.20190087 | 0.03536085 | 0.00327453 | -47.3742485 | 0.00033445 | 47.1023369 | 50.6002159 | -0.79270959 | 1.04663134 | 0.01872011 | 78.464035 | 0.00022997 | 0.00021529 | 32.4618187 | 0.00534759 | 0.00372841 | 0.00307112 | 0.53724986 | 0.06002188 | 35.976 | 0.00023852 | 28.5042152 | 79.5553055 | 0.00036028 | 0.00384822 | 10.3135185 | 0.90134049 | 0.00124915 | 0.00236292 | 0.21374942 | 0.00207461 | 0.00021351 | 31.5567303 | 0.16914773 | 94.488327 | 0.00047666 | 0.16939189 | 0.00232844 | 83.4985504 | -0.48923683 | 0.00089983 | 92.8806152 | 0.00739768 | 0.04160358 | 0.01807535 | 0.02164291 | 0.00118939 | 68.5108719 | 0.0057689 | 0.00031025 | 0.00879073 | 0.00311874 | 0.25839397 | 41.7190895 | 0.00132802 | 1.11506879 | 0.00033201 | -51.2652473 | -27.334198 | 117.200264 | -43.8369331 | 0.27907762 | 0.00075167 | 4.74944735 | 27.0386524 | 0.00171605 | 0.00089313 | 65.0524139 | 2.94422722 | 0.00245917 | 5.28827143 | 0.00376457 | -80.8234253 | 0.00324919 | 83.0622559 | 0.24552706 | 0.00404406 | 40.0305061 | 0.00609317 | -3.40348816 | 26.5942307 |
| 9.59E-05 | -0.20143986 | 49.9529266 | 43.319706 | 0.00208986 | 6.08 | 0.00559648 | -43.7367783 | 0.00358359 | 0.00506703 | -33.1601448 | 0.00155494 | 1.0639503 | 0.00436227 | 0.21504596 | 0.9233855 | -89.9760895 | 56.2739792 | 0.00834925 | 0.00362313 | -3.49100709 | 38.0814819 | -10.146246 | 0.78277051 | 0.00217005 | 0.00174866 | 63.6327782 | 0.4540343 | 0.04666783 | 0.00349796 | -50.9136887 | 0.00021913 | 69.8286667 | 50.4847527 | -0.3746776 | 1.17088997 | 0.01914583 | 72.0239716 | 0.00020086 | 0.00030967 | 35.224865 | 0.00600731 | 0.00425749 | 0.00444954 | 0.6365304 | -0.02236366 | 17.432 | 0.00027154 | 37.2610397 | 71.6743012 | 0.00036757 | 0.0054628 | 9.96753883 | 1.11033094 | 0.00141463 | 0.00229468 | 0.28170797 | 0.00154822 | 0.0002176 | 44.632431 | 0.14725062 | 88.7322846 | 0.03180853 | 0.19507356 | 0.00223735 | 80.1880493 | -0.92072874 | 0.00094844 | 88.3264465 | 0.00754781 | 0.05712817 | 0.01931066 | 0.02202819 | 0.00183429 | 61.8555985 | 0.00527571 | 0.00030332 | 0.00673824 | 0.00353747 | 0.32935625 | 62.6617737 | 0.00164673 | 1.20995569 | 0.00028094 | -48.6366348 | -23.6981812 | 104.488838 | -22.7396889 | 0.42565161 | 0.00134325 | 4.99064732 | 33.3776817 | 0.00315257 | 0.0006745 | 77.1656036 | 3.19660115 | 0.00165602 | 5.33003998 | 0.00361839 | -122.775101 | 0.002853 | 80.3227158 | 0.3344669 | 0.00473894 | 40.1094322 | 0.00629481 | -1.84757757 | 31.9050255 |
| 9.75E-05 | -0.17081499 | 49.6197891 | 41.321228 | 0.0019923 | 5.96 | 0.00557352 | -43.4643822 | 0.00408698 | 0.00569973 | -31.545826 | 0.00137672 | 0.51556438 | 0.0042382 | 0.22990167 | 0.7313019 | -82.2191849 | 52.6765251 | 0.00706658 | 0.00385588 | -3.07395816 | 123.065002 | -10.2892399 | 0.66126472 | 0.00219724 | 0.00152496 | 57.6785622 | 0.40024573 | 0.03929969 | 0.00326261 | -46.6973991 | 0.00023116 | 61.1747055 | 46.3955689 | 0.22872013 | 0.96673799 | 0.01907359 | 69.6953507 | 0.00019195 | 0.00015336 | 31.2115421 | 0.00528316 | 0.00348492 | 0.00319458 | 0.50888795 | 0.06325841 | 27.376 | 0.00019902 | 34.5273171 | 70.2458572 | 0.00036003 | 0.00482457 | 9.46937943 | 1.20593369 | 0.00160073 | 0.00206786 | 0.23012281 | 0.00179092 | 0.00019424 | 43.434967 | 0.12550907 | 84.9658508 | 0.05608525 | 0.17487755 | 0.00219268 | 72.7468872 | -0.45868012 | 0.00091473 | 82.2649002 | 0.007382 | 0.05170276 | 0.02041029 | 0.01976218 | 0.00154908 | 58.0880432 | 0.0054707 | 0.00034231 | 0.00624752 | 0.00371937 | 0.30058166 | 58.682106 | 0.00182365 | 1.19439554 | 0.00027995 | -46.7146835 | -22.8720627 | 99.3341751 | -24.2795658 | 0.34096631 | 0.00091698 | 4.35077667 | 32.6506424 | 0.00217318 | 0.0006829 | 76.4825363 | 2.92012644 | 0.0019717 | 4.77649927 | 0.00400918 | -107.388519 | 0.00281742 | 72.2641449 | 0.27807128 | 0.0040308 | 37.4481735 | 0.0061144 | -3.77865768 | 30.3967819 |
| 8.64E-05 | -0.27575493 | 59.6485024 | 37.4309082 | 0.00195123 | 4.04 | 0.00524036 | -39.9994431 | 0.00337363 | 0.00454589 | -40.5209351 | 0.00144826 | 2.21944857 | 0.00501339 | 0.46651667 | 1.97236419 | -86.5663681 | 64.598999 | 0.00847738 | 0.00350827 | -3.7039237 | 43.5893555 | -14.6204233 | 0.74236006 | 0.00165917 | 0.00165818 | 70.296257 | 0.89440483 | 0.04112951 | 0.00328792 | -47.958046 | 0.00016537 | 74.2329941 | 43.9407883 | 1.70730066 | 1.21450532 | 0.01683581 | 82.1660004 | 0.00021279 | 0.00023512 | 37.0468674 | 0.00573629 | 0.004432 | 0.00478902 | 0.68706763 | 0.01971722 | 18.156 | 0.00019909 | 40.1566925 | 83.1399689 | 0.00033851 | 0.00579061 | 9.15099239 | 1.29046416 | 0.00132797 | 0.00218912 | 0.26572529 | 0.00130254 | 0.00019427 | 49.4165573 | 0.14091466 | 78.8249741 | 0.09850698 | 0.17574111 | 0.00192243 | 88.6285095 | -0.56640142 | 0.0008311 | 75.8155518 | 0.0056576 | 0.06152261 | 0.01880227 | 0.01839919 | 0.00176467 | 70.3777847 | 0.00464774 | 0.00029734 | 0.0065888 | 0.00337595 | 0.37230846 | 69.3402252 | 0.00155506 | 1.16566563 | 0.00028006 | -42.7946816 | -22.0220738 | 92.403656 | 7.98448706 | 0.42540681 | 0.00133034 | 4.60643292 | 38.665184 | 0.00311931 | 0.00064958 | 81.9988709 | 3.1154964 | 0.0013572 | 5.12902641 | 0.00319267 | -146.16864 | 0.00283632 | 86.8063965 | 0.30685863 | 0.00432369 | 33.1550331 | 0.00482933 | -3.40182805 | 38.5590591 |
| 8.29E-05 | -0.13378263 | 59.4347267 | 37.237854 | 0.00208302 | 4.68 | 0.00588734 | -40.0596542 | 0.00353235 | 0.0048621 | -48.6587906 | 0.00141887 | 1.90418744 | 0.00518122 | 0.47160965 | 2.68292379 | -87.8646545 | 64.7949219 | 0.00854535 | 0.00369111 | -5.5799362 | 45.5435181 | -19.1473751 | 0.78410733 | 0.00216587 | 0.00170592 | 70.2923584 | 1.07866156 | 0.04490296 | 0.00335346 | -62.5788803 | 0.00015476 | 76.4687653 | 46.3296318 | 0.99029142 | 1.30086017 | 0.01938973 | 82.4931335 | 0.0002148 | 0.00025548 | 38.090538 | 0.00562095 | 0.00419226 | 0.00452605 | 0.6590122 | 0.11005402 | 20.144 | 0.00023338 | 42.7140808 | 82.4678726 | 0.00031454 | 0.00572769 | 9.40507507 | 1.44169855 | 0.00160472 | 0.00224668 | 0.28372121 | 0.00139928 | 0.00020065 | 51.9785843 | 0.15074503 | 78.121933 | 0.09962563 | 0.17745644 | 0.00207667 | 85.4338837 | -0.48536772 | 0.00089736 | 78.3412552 | 0.0075595 | 0.06199126 | 0.01849885 | 0.01864299 | 0.00178597 | 70.3987045 | 0.00480192 | 0.00027932 | 0.00639906 | 0.0034276 | 0.38397923 | 77.0694962 | 0.00190204 | 1.30732715 | 0.00027354 | -44.6768189 | -23.4982185 | 99.6737747 | 10.7143965 | 0.43492574 | 0.00131102 | 4.66239166 | 45.7798538 | 0.00317888 | 0.00067524 | 92.4087067 | 3.45913053 | 0.00140407 | 5.38631058 | 0.0033106 | -147.458237 | 0.00296738 | 85.5046768 | 0.30995154 | 0.0042886 | 33.9527588 | 0.00642126 | -3.03780174 | 45.5940247 |
| 7.63E-05 | -0.07116795 | 57.6488915 | 38.2419701 | 0.00184615 | 4.38 | 0.00442448 | -40.4476242 | 0.00288193 | 0.00407223 | -42.7354393 | 0.00145294 | 1.4805032 | 0.00502759 | 0.26770148 | 2.10362506 | -92.0057449 | 62.2372284 | 0.00841163 | 0.00306634 | -1.85756707 | 103.022705 | -14.0188055 | 0.68365675 | 0.00153726 | 0.00175039 | 67.6918564 | 0.77176249 | 0.02157184 | 0.00314644 | -50.944767 | 0.0001653 | 74.6171265 | 45.4630966 | 0.47242162 | 1.26289642 | 0.0172706 | 80.8671722 | 0.00019025 | 0.00010231 | 41.6601753 | 0.00527895 | 0.00442585 | 0.00440605 | 0.68492103 | 0.06012917 | 16.324 | 0.00014464 | 40.3576584 | 82.7073669 | 0.00029899 | 0.00534185 | 9.6933012 | 1.18663907 | 0.00124686 | 0.00195767 | 0.175124 | 0.00146578 | 0.00016968 | 50.4301643 | 0.1390475 | 80.9833374 | 0.08915024 | 0.1585469 | 0.00200865 | 86.6120911 | -0.1484526 | 0.00090034 | 77.0320892 | 0.00511361 | 0.05711087 | 0.01703675 | 0.01386153 | 0.00135312 | 67.1247864 | 0.00411918 | 0.00023259 | 0.00651294 | 0.00277084 | 0.34778369 | 70.3660355 | 0.00143026 | 1.06226599 | 0.0002381 | -44.195919 | -22.2593689 | 93.1363678 | 14.1121769 | 0.39087135 | 0.00093003 | 4.43284369 | 41.5186005 | 0.00234098 | 0.00064585 | 83.5108795 | 2.90009713 | 0.00144923 | 5.5144887 | 0.00295836 | -152.155548 | 0.00264554 | 85.337677 | 0.27376741 | 0.00340181 | 33.8286514 | 0.00438677 | -3.34925604 | 41.338398 |
| 8.21E-05 | -0.31431317 | 56.9894524 | 37.3728294 | 0.00236013 | 4.5 | 0.00520879 | -40.1271782 | 0.00341198 | 0.00459917 | -41.3626709 | 0.00138133 | 1.53321958 | 0.00538195 | 0.49318478 | 1.82550323 | -83.8527832 | 61.8086281 | 0.00798895 | 0.00362229 | -4.01490784 | 58.2520752 | -13.2290125 | 0.73287052 | 0.0018407 | 0.00165958 | 69.7427368 | 1.02635896 | 0.03820638 | 0.00324625 | -49.2747994 | 0.00015709 | 97.2029648 | 44.365799 | 0.02510114 | 1.24865651 | 0.01756323 | 79.8618927 | 0.00022955 | 0.00025172 | 38.4484024 | 0.0054015 | 0.00380883 | 0.004181 | 0.62198973 | 0.00743866 | 17.288 | 0.00025003 | 46.760128 | 81.0245132 | 0.00030557 | 0.00532712 | 9.80342865 | 1.26785398 | 0.00130806 | 0.00226104 | 0.24248439 | 0.0013701 | 0.00017123 | 57.6280518 | 0.16569504 | 77.2322388 | 0.10540978 | 0.17592783 | 0.00204799 | 86.481308 | -0.57062078 | 0.00089289 | 77.4113693 | 0.00564011 | 0.06289725 | 0.01718547 | 0.01754823 | 0.00175909 | 70.3144608 | 0.00464959 | 0.00028071 | 0.00658627 | 0.0034466 | 0.38140583 | 71.2223816 | 0.00154026 | 1.12195182 | 0.00025407 | -42.5555725 | -22.4256096 | 96.6235046 | 6.79120827 | 0.39675838 | 0.00130232 | 4.6035552 | 39.7571259 | 0.00329055 | 0.00072863 | 80.8141708 | 3.35552526 | 0.00128562 | 5.17821741 | 0.00330574 | -134.027771 | 0.00289098 | 85.0277557 | 0.30063564 | 0.00454429 | 34.2145844 | 0.00475115 | -2.45611 | 39.9351807 |
| 8.36E-05 | -0.27496815 | 58.3217659 | 36.8487816 | 0.00184228 | 4.56 | 0.005168 | -40.5485344 | 0.00332177 | 0.00458907 | -41.7339745 | 0.00126056 | 2.4989655 | 0.00480381 | 0.47802728 | 2.31308746 | -85.3783493 | 63.5757828 | 0.00770339 | 0.00344581 | -3.76247025 | 49.1898804 | -14.0210762 | 0.75649756 | 0.00183816 | 0.00167404 | 68.3667755 | 0.90958726 | 0.03408969 | 0.00303537 | -50.6479378 | 0.00014772 | 72.4564667 | 42.2409058 | 0.5995791 | 1.20780778 | 0.01682622 | 81.4453735 | 0.00018884 | 0.00018522 | 38.2505684 | 0.00533671 | 0.00381521 | 0.00443355 | 0.65768701 | 0.01988411 | 16.352 | 0.00023254 | 42.7996597 | 81.7563782 | 0.00030519 | 0.0049725 | 9.0155859 | 1.2380445 | 0.00125263 | 0.00231354 | 0.22572032 | 0.00130439 | 0.00017032 | 51.9798508 | 0.14088081 | 75.1309357 | 0.09158704 | 0.17456666 | 0.00201932 | 85.7843323 | -0.45862582 | 0.00087363 | 73.4453735 | 0.00574832 | 0.05180024 | 0.01694905 | 0.0167471 | 0.00155697 | 67.7856751 | 0.00464016 | 0.00028555 | 0.0062522 | 0.00341804 | 0.31939143 | 65.8248444 | 0.00146299 | 1.13310564 | 0.0002592 | -41.4775353 | -21.2404842 | 99.7177124 | -15.4095631 | 0.42030865 | 0.00117937 | 4.43738079 | 39.270462 | 0.00279579 | 0.00068564 | 78.1202316 | 2.86792517 | 0.00131769 | 5.32256746 | 0.00341382 | -137.401016 | 0.0028304 | 84.7963486 | 0.31317043 | 0.00392205 | 34.860199 | 0.00479223 | -2.90251398 | 39.3689385 |
| 8.91E-05 | 0.01538992 | 56.1537743 | 43.0828705 | 0.00200113 | 6.12 | 0.00451017 | -45.7744141 | 0.00240427 | 0.00348559 | -36.8221436 | 0.00108533 | 0.33800128 | 0.0032613 | -0.02226585 | -0.46426141 | -116.288284 | 59.5224686 | 0.00708577 | 0.00256625 | -3.71000814 | 52.8022461 | -11.4926729 | 1.31705082 | 0.00169234 | 0.00129172 | 68.9227753 | 0.18514609 | 0.03635876 | 0.00240209 | -58.435051 | 0.00031752 | 160.744782 | 78.2151108 | -0.69828332 | 1.43912005 | 0.02050595 | 79.810318 | 0.00022175 | 0.00028974 | 39.8480721 | 0.00542793 | 0.00402742 | 0.00429471 | 0.96711296 | 0.0407517 | 16.63 | 0.00027923 | 67.860733 | 80.4703674 | 0.00035165 | 0.00629204 | 13.1056337 | 1.08450294 | 0.00125296 | 0.00157395 | 0.22438936 | 0.00126817 | 0.00016894 | 89.4264374 | 0.21010046 | 110.415543 | -0.05540256 | 0.21970105 | 0.00143078 | 101.419067 | -1.07003379 | 0.0005741 | 118.948372 | 0.00628417 | 0.052848 | 0.01704574 | 0.01817645 | 0.00144783 | 65.8985748 | 0.00373331 | 0.00028358 | 0.00518357 | 0.00216589 | 0.346766 | 79.4875565 | 0.00145695 | 1.2277813 | 0.00022374 | -85.5884628 | -27.1642971 | 104.871803 | -18.9920864 | 0.61335111 | 0.00074393 | 4.85346556 | 37.8872147 | 0.0019793 | 0.00055119 | 96.1977005 | 3.91293454 | 0.00137207 | 5.99731016 | 0.00316251 | -242.60672 | 0.00273546 | 82.7370911 | 0.4350926 | 0.00355518 | 39.3402023 | 0.00534507 | -3.67728138 | 33.5024948 |
| 9.63E-05 | -0.11329651 | 52.6408806 | 44.1690941 | 0.0021495 | 5.04 | 0.00382216 | -45.6517944 | 0.00327734 | 0.00431652 | -25.1943741 | 0.00175434 | 0.53395611 | 0.00353607 | -0.00892938 | -0.17986633 | -89.7019043 | 59.1440849 | 0.00612287 | 0.00229082 | -1.01112735 | 35.5496216 | -7.51355457 | 0.37661451 | 0.00188774 | 0.00114026 | 71.6034851 | 0.03434255 | 0.01562686 | 0.00335659 | -46.6637878 | 0.00103052 | 55.6724243 | 46.83424 | -1.10267794 | 0.78612977 | 0.01386186 | 75.5477295 | 0.00020415 | 3.50E-05 | 28.158741 | 0.00403644 | 0.00309574 | 0.00251868 | 0.42579734 | 0.00457764 | 42.55 | 0.00028534 | 31.7818718 | 78.5419312 | 0.00029326 | 0.00288503 | 9.86565018 | 1.08871973 | 0.00123013 | 0.00126689 | 0.12370374 | 0.0011229 | 0.00016304 | 35.6299057 | 0.15767777 | 84.6534119 | -0.00511326 | 0.1331247 | 0.00177128 | 80.2048035 | -0.17123245 | 0.00067811 | 88.3110352 | 0.00731358 | 0.04111105 | 0.01294356 | 0.01186283 | 0.00060314 | 58.6237793 | 0.00332257 | 0.00026786 | 0.00612285 | 0.00187921 | 0.24417441 | 48.980835 | 0.00141714 | 1.23128486 | 0.00024044 | -44.9068451 | -26.603447 | 56.3895035 | -43.8953514 | 0.34630916 | 0.00051152 | 3.87947154 | 25.6576271 | 0.00112844 | 0.00030872 | 70.7993469 | 2.58706617 | 0.00111746 | 5.22276115 | 0.00214833 | -95.4085388 | 0.00240725 | 92.7303467 | 0.18112892 | 0.00292327 | 41.539753 | 0.00611653 | 0.59797108 | 23.9721451 |
| 9.87E-05 | -0.09719133 | 52.0673904 | 41.1956749 | 0.00240686 | 5.16 | 0.00566139 | -41.4946022 | 0.00312203 | 0.00473927 | -41.9347153 | 0.00099249 | 0.76833206 | 0.0054721 | 0.15182516 | -0.0253739 | -88.6408463 | 59.1631241 | 0.00708925 | 0.00325313 | -2.36600065 | 109.333679 | -14.0864983 | 0.73839569 | 0.00206883 | 0.00151145 | 67.0554504 | 0.16148672 | 0.03930448 | 0.002437 | -53.7465744 | 0.00026802 | 101.294968 | 57.3111839 | -1.30069542 | 0.99309719 | 0.02410332 | 78.3086929 | 0.00021022 | 0.00019141 | 33.9249497 | 0.0048643 | 0.00335191 | 0.0033824 | 0.55865562 | 0.035429 | 23.276 | 0.00036057 | 48.7202988 | 74.442543 | 0.00033079 | 0.00471857 | 11.1090727 | 1.24220276 | 0.00174867 | 0.00215107 | 0.2429478 | 0.0017033 | 0.00018421 | 64.2367096 | 0.16310284 | 94.9763336 | 0.01234466 | 0.19303846 | 0.0019689 | 83.537468 | -0.17516817 | 0.00081204 | 97.521492 | 0.00716463 | 0.07015341 | 0.01873085 | 0.01896885 | 0.00146591 | 68.7581635 | 0.00536339 | 0.00029376 | 0.00685746 | 0.00288214 | 0.40624893 | 76.8031464 | 0.00202372 | 1.18874991 | 0.0002622 | -55.5606804 | -24.1195183 | 95.9123535 | -9.98625279 | 0.35567325 | 0.00089505 | 4.29794121 | 42.7298241 | 0.00221823 | 0.00060587 | 99.521019 | 3.73147488 | 0.00197895 | 4.9266758 | 0.00374686 | -124.376617 | 0.0029915 | 82.8327103 | 0.30625659 | 0.00436447 | 37.174408 | 0.00590628 | -4.20887995 | 43.7957153 |
| 9.67E-05 | -0.02728701 | 53.8638573 | 41.3312225 | 0.00250201 | 4.5 | 0.00450029 | -43.9916649 | 0.00289108 | 0.00438956 | -38.0399666 | 0.0011476 | 0.63845241 | 0.00395015 | 0.14386439 | 0.64911169 | -102.608101 | 60.5593033 | 0.00715402 | 0.0032823 | -3.01368618 | 50.9301758 | -14.4431953 | 1.10115552 | 0.00181697 | 0.00128977 | 65.2314606 | 0.14414203 | 0.03663154 | 0.00287004 | -50.9211578 | 0.00018845 | 180.774719 | 63.6057625 | -1.24389577 | 1.32405436 | 0.02447202 | 79.7950668 | 0.00025002 | 0.00025535 | 34.8932495 | 0.00603712 | 0.00377408 | 0.00477492 | 0.84302938 | 0.05362034 | 14.52 | 0.00031057 | 88.9509583 | 78.5550461 | 0.00034035 | 0.00530982 | 13.835947 | 1.11738265 | 0.00128016 | 0.00176256 | 0.23756999 | 0.00155124 | 0.00016805 | 110.03891 | 0.25479427 | 99.7194977 | -0.0126518 | 0.22224644 | 0.00172545 | 83.6902771 | -0.57188094 | 0.00066681 | 102.848457 | 0.00603318 | 0.07136627 | 0.02172244 | 0.02007341 | 0.00226769 | 65.2719727 | 0.0042036 | 0.00029234 | 0.00600566 | 0.00252644 | 0.43917292 | 100.659103 | 0.00149895 | 1.10807383 | 0.00026225 | -64.1425171 | -22.8552551 | 101.505615 | 24.4026184 | 0.53396398 | 0.00108965 | 4.63349581 | 38.5606346 | 0.00321553 | 0.00070591 | 98.7803497 | 3.47749186 | 0.00174341 | 5.16043234 | 0.00294669 | -185.398285 | 0.00261792 | 83.1537933 | 0.43139625 | 0.00454396 | 39.2704659 | 0.00506806 | -4.68071413 | 37.4387207 |
| 8.92E-05 | -0.24849176 | 50.0911026 | 39.680912 | 0.00187658 | 3.48 | 0.00487755 | -43.5552063 | 0.00347112 | 0.00545741 | -27.9421883 | 0.00141094 | 0.6717211 | 0.00531354 | 0.14226876 | -0.75050557 | -87.0493774 | 57.4974594 | 0.00919445 | 0.00398139 | -2.59009385 | 49.6130371 | -6.08077097 | 0.57076144 | 0.0018405 | 0.00137997 | 64.1717758 | 0.36878541 | 0.03189827 | 0.00302539 | -42.6374359 | 0.00039605 | 51.691124 | 46.3805962 | 3.04079914 | 1.08929563 | 0.01562921 | 71.9470367 | 0.00018362 | 0.00027474 | 27.6653976 | 0.00470324 | 0.00395654 | 0.00286047 | 0.55622029 | 0.04376173 | 34.808 | 0.0003306 | 31.1837025 | 73.7024536 | 0.00030525 | 0.00395545 | 9.76108837 | 0.94586009 | 0.00103108 | 0.0018231 | 0.19768338 | 0.00220248 | 0.00017814 | 35.8518829 | 0.14899015 | 80.6166382 | -0.00890504 | 0.15907647 | 0.00180781 | 83.3426209 | -0.17304656 | 0.00072892 | 81.1073303 | 0.00587721 | 0.0455587 | 0.01910801 | 0.02054782 | 0.00124198 | 63.5836601 | 0.00497943 | 0.00029756 | 0.00907686 | 0.00293189 | 0.25798425 | 41.5096741 | 0.00123704 | 1.04668427 | 0.00026086 | -48.5943337 | -25.6536713 | 115.1763 | -44.291256 | 0.32203579 | 0.00070487 | 4.60069609 | 32.5687637 | 0.00166071 | 0.0010701 | 64.5190277 | 2.96643114 | 0.00220857 | 5.09117937 | 0.0039043 | -98.5972595 | 0.00314065 | 85.0346298 | 0.24643928 | 0.00353191 | 36.9698563 | 0.00481372 | -3.01054144 | 27.5530338 |
| 0.00010605 | -0.071913 | 54.7052994 | 34.9766808 | 0.00245537 | 5.08 | 0.00435864 | -37.2844925 | 0.00352299 | 0.00511533 | -36.7023163 | 0.00177382 | 0.83469886 | 0.00554417 | 0.32436076 | -0.01681704 | -95.3277817 | 59.7297745 | 0.01004712 | 0.00410917 | -1.9725498 | 65.984375 | -10.1419239 | 1.00597417 | 0.00132992 | 0.00178578 | 65.2397003 | 0.51887018 | 0.03269592 | 0.00387538 | -35.5489883 | 0.00021298 | 125.536575 | 54.0719795 | 0.55386037 | 1.46873796 | 0.02616176 | 77.8880539 | 0.00027784 | 0.00028383 | 30.9767151 | 0.00663853 | 0.0046834 | 0.00551045 | 0.87903845 | 0.01814365 | 15.736 | 0.00031223 | 58.2055969 | 78.5372009 | 0.00036978 | 0.00569252 | 11.6179142 | 1.07040465 | 0.00120474 | 0.00220968 | 0.23409861 | 0.00171502 | 0.00019373 | 76.8179627 | 0.2419429 | 88.9822083 | 0.04272404 | 0.22320607 | 0.00204869 | 82.0544663 | -0.53963482 | 0.00085215 | 88.2024384 | 0.00426839 | 0.06862158 | 0.0224616 | 0.02070033 | 0.0029926 | 65.569046 | 0.0049857 | 0.00035214 | 0.00750669 | 0.0033621 | 0.40675905 | 78.0345972 | 0.00133547 | 0.92202747 | 0.00034546 | -55.9130478 | -22.5634689 | 126.878067 | 22.296175 | 0.52040362 | 0.0017608 | 4.5023427 | 36.3179283 | 0.00417977 | 0.00111389 | 78.4300842 | 3.04096031 | 0.00173043 | 5.43477011 | 0.00350672 | -143.090942 | 0.00261508 | 81.1451035 | 0.40822965 | 0.00538232 | 31.7995281 | 0.0036975 | -5.00061846 | 36.9404221 |
| 0.00010209 | -0.12267232 | 53.1861382 | 32.2808533 | 0.00211902 | 5.24 | 0.00424195 | -34.6850815 | 0.00338164 | 0.00488442 | -33.7904167 | 0.00167922 | 0.80013543 | 0.00511283 | 0.35103816 | 0.2159756 | -76.9148941 | 58.5954056 | 0.01026029 | 0.00418082 | -1.97060466 | 82.1817627 | -6.52280045 | 0.72713041 | 0.00131515 | 0.00188512 | 64.1170425 | 0.76284897 | 0.02579962 | 0.00353667 | -33.5037499 | 0.00025353 | 73.6371155 | 44.643898 | 0.35459557 | 1.11353672 | 0.01954584 | 77.2682953 | 0.00021977 | 0.00017692 | 36.3823853 | 0.00555176 | 0.00469969 | 0.00433205 | 0.71086478 | 0.00676513 | 24.848 | 0.00026694 | 40.6344757 | 77.4617233 | 0.00030937 | 0.00429509 | 10.5891438 | 0.99311888 | 0.001111 | 0.00189877 | 0.18233395 | 0.00176812 | 0.00016073 | 56.9535561 | 0.19326466 | 81.4879227 | 0.07121776 | 0.20131615 | 0.0018399 | 82.5121918 | -0.38017833 | 0.00078735 | 82.1250382 | 0.00393628 | 0.05054919 | 0.01796871 | 0.01819374 | 0.00170079 | 64.0960846 | 0.00482632 | 0.0002976 | 0.00866376 | 0.00329091 | 0.31785789 | 58.9694786 | 0.00122798 | 0.85381538 | 0.00031956 | -46.4271507 | -23.6185665 | 134.100372 | -8.97347832 | 0.36119109 | 0.00113381 | 4.31335592 | 33.8372688 | 0.00281247 | 0.0009829 | 72.0882874 | 2.74558258 | 0.00168897 | 4.9844079 | 0.00338548 | -107.540207 | 0.002506 | 81.270752 | 0.32197875 | 0.0041043 | 28.7622776 | 0.00333029 | -4.94594288 | 32.759079 |
| 0.0001127 | -0.22414923 | 56.4388504 | 29.9087048 | 0.00249257 | 5.34 | 0.00410475 | -32.0778198 | 0.00311821 | 0.00458962 | -36.2272911 | 0.00168223 | 0.50319439 | 0.00541566 | 0.32092798 | -0.15195575 | -70.9080353 | 60.5323792 | 0.00958337 | 0.00390916 | -2.09583902 | 66.6917725 | -11.9138784 | 0.71377641 | 0.00123872 | 0.00159642 | 67.4311142 | 0.66758096 | 0.03000619 | 0.00366759 | -30.8574238 | 0.00023235 | 132.658463 | 46.4221001 | 0.18844324 | 1.03402782 | 0.02243554 | 78.0896378 | 0.00025794 | 0.00027375 | 25.3082638 | 0.00563935 | 0.00475563 | 0.00465684 | 0.72776747 | -0.00475049 | 23.35 | 0.00030246 | 53.5088501 | 79.8106232 | 0.00037924 | 0.00536947 | 9.62838554 | 1.07200575 | 0.00126799 | 0.00216341 | 0.20209071 | 0.00175491 | 0.00018176 | 63.7754974 | 0.19641616 | 75.5625153 | 0.04604247 | 0.18499069 | 0.00197066 | 86.1081772 | -0.18717872 | 0.00079276 | 75.9684677 | 0.00388298 | 0.06599799 | 0.01693902 | 0.01609155 | 0.00182604 | 66.5253296 | 0.00455951 | 0.00029955 | 0.00658193 | 0.00306204 | 0.40970135 | 76.1654205 | 0.00144048 | 0.89353335 | 0.00031819 | -45.8985367 | -21.4827061 | 121.583649 | 25.0849934 | 0.35754946 | 0.00095457 | 3.92755246 | 37.5801888 | 0.00269532 | 0.00064819 | 76.0742188 | 3.26082277 | 0.00179934 | 4.60652256 | 0.00370231 | -127.911705 | 0.00254797 | 81.0243683 | 0.29979578 | 0.00401802 | 25.5677395 | 0.00336734 | -4.68497849 | 37.7619934 |
| 0.00011424 | -0.07570982 | 57.3078232 | 29.9277782 | 0.00233286 | 4.2 | 0.00402284 | -32.9556541 | 0.00297284 | 0.00433795 | -34.9360275 | 0.00171814 | 0.82523906 | 0.00546662 | 0.31968827 | 0.05760833 | -85.4590988 | 61.4585342 | 0.01043711 | 0.00383338 | -1.60017157 | 45.5472412 | -10.1027241 | 0.94826049 | 0.00119575 | 0.00153494 | 68.1473541 | 0.48513258 | 0.02898481 | 0.00367351 | -31.0608482 | 0.0001702 | 135.529327 | 52.0641518 | -0.73085195 | 1.25182426 | 0.02446927 | 80.5834045 | 0.00028034 | 0.00019208 | 33.1896095 | 0.00628953 | 0.00528045 | 0.00547875 | 0.87293929 | 0.08274317 | 17.108 | 0.00028995 | 62.8887863 | 81.3228302 | 0.00035988 | 0.0053621 | 10.8084383 | 1.03618598 | 0.00116994 | 0.00193114 | 0.20912954 | 0.00149424 | 0.00018223 | 83.5960617 | 0.24535126 | 85.7814407 | 0.06037031 | 0.22082134 | 0.00166288 | 85.2498245 | -0.18146698 | 0.00068208 | 86.2235031 | 0.00404855 | 0.07144611 | 0.0203289 | 0.01758925 | 0.00254319 | 68.5758667 | 0.0041893 | 0.00031893 | 0.00721094 | 0.00303216 | 0.43272746 | 83.4874115 | 0.00129902 | 0.88210648 | 0.00034413 | -53.5939179 | -21.393671 | 110.434616 | 26.5951805 | 0.47251952 | 0.00141291 | 4.18870306 | 36.3482819 | 0.00370912 | 0.00090464 | 83.9477081 | 3.09665155 | 0.0014682 | 5.27249908 | 0.00312802 | -152.174088 | 0.00240725 | 84.5108185 | 0.37572479 | 0.00471522 | 24.7354507 | 0.00349831 | -7.85935211 | 36.2912445 |
| 9.94E-05 | -0.07237792 | 54.4430161 | 32.1588898 | 0.00192622 | 4.92 | 0.00382061 | -33.8836365 | 0.00326454 | 0.00455752 | -34.3689537 | 0.00168209 | 0.34747407 | 0.00571012 | 0.20841217 | -0.6352669 | -75.8989792 | 59.8450165 | 0.00989522 | 0.00430569 | -1.57658196 | 33.5059815 | -9.27119255 | 0.67115212 | 0.00127024 | 0.00158383 | 64.9473114 | 0.53588647 | 0.0230768 | 0.00349637 | -32.2542381 | 0.00025259 | 65.4775848 | 43.5230904 | -0.28949401 | 1.02488446 | 0.02025607 | 77.2676697 | 0.00018815 | 0.00015799 | 27.4385262 | 0.00516518 | 0.00458402 | 0.00399087 | 0.68444657 | 0.03947616 | 29.31 | 0.00017409 | 33.7073288 | 76.9084168 | 0.00030972 | 0.00459704 | 9.42066193 | 0.94724596 | 0.00100965 | 0.00197964 | 0.16844559 | 0.00173928 | 0.0001659 | 47.7511482 | 0.16840679 | 74.7319794 | 0.0697017 | 0.18456082 | 0.00190453 | 80.3552475 | -0.13264677 | 0.00077355 | 72.3323517 | 0.00367901 | 0.05685908 | 0.01703198 | 0.0157315 | 0.0014196 | 64.3888321 | 0.00460189 | 0.00027814 | 0.00757096 | 0.00334321 | 0.32826513 | 62.6315002 | 0.00119523 | 0.82632268 | 0.00029766 | -43.5611 | -21.3875446 | 118.91938 | 11.7306938 | 0.34778857 | 0.00082875 | 4.16525173 | 36.6846657 | 0.00217061 | 0.00104317 | 74.9105911 | 2.54406667 | 0.00158833 | 4.89437485 | 0.00303214 | -116.015938 | 0.00252965 | 79.7383423 | 0.2957252 | 0.00339582 | 27.9865074 | 0.00307367 | -4.52709484 | 35.4239655 |

| d_curvature_n | d_ang_vel_abs | ang_vel_head_ | d_major_axis_ | curvature_tail_ | d_length_10th | d_curvature_st | d_ang_vel_hea | d_length_90th | d_ang_vel_tail | d_curvature_st | curvature_mea | speed_tail_bas | ang_vel_head_ |
|---|---|---|---|---|---|---|---|---|---|---|---|---|---|
| 0.00543468 | 0.01933818 | 0.67648608 | -65.1937714 | 0.0016434 | -36.3364563 | 0.00142607 | 8.4464407 | 35.7500763 | 0.7513054 | 0.00252189 | 0.00366038 | 22.3199234 | 0.32509071 |
| 0.00908325 | 0.02061996 | 0.8654967 | -78.1130219 | 0.00092577 | -36.3838882 | 0.001296 | 10.2731142 | 35.836792 | 0.674707 | 0.00310116 | 0.00433562 | 31.3657894 | 0.412184 |
| 0.00723564 | 0.01288839 | 0.74673593 | -79.8521042 | 0.00103694 | -34.8472862 | 0.00116222 | 7.04516268 | 34.2675705 | 0.71506292 | 0.0020406 | 0.00524423 | 23.4321365 | 0.37220356 |
| 0.00540961 | 0.01694686 | 0.60714281 | -56.9590378 | 0.00059673 | -38.3774262 | 0.00117914 | 9.00603771 | 37.8155441 | 0.69044387 | 0.00272103 | 0.00499638 | 31.4756393 | 0.29458219 |
| 0.00337919 | 0.01837958 | 0.64134079 | -60.8369446 | 0.00093288 | -38.2190895 | 0.00131299 | 8.95371723 | 37.8349037 | 0.69436038 | 0.00241397 | 0.0050391 | 35.6156273 | 0.32539898 |
| 0.00745645 | 0.01358668 | 0.9424749 | -85.0255432 | 0.00076326 | -31.9803639 | 0.00125053 | 9.45306873 | 30.856842 | 0.72016555 | 0.00272478 | 0.00372544 | 34.7016182 | 0.42228073 |
| 0.00768786 | 0.00932296 | 0.92712683 | -97.3660278 | 0.00078526 | -29.4269104 | 0.00111997 | 9.37217712 | 28.9966145 | 0.66731948 | 0.00266864 | 0.00372903 | 36.3028984 | 0.43372873 |
| 0.00819213 | 0.01750706 | 1.15057671 | -82.586174 | 0.000648 | -32.162487 | 0.00158457 | 11.0729485 | 31.8803368 | 0.85250145 | 0.00319669 | 0.00360671 | 31.6037083 | 0.54212976 |
| 0.00851423 | 0.01936019 | 1.14683962 | -101.895065 | 0.00078215 | -35.4086418 | 0.00163989 | 10.7618866 | 34.5463677 | 0.84285903 | 0.00289909 | 0.00353855 | 34.9220581 | 0.54300195 |
| 0.00912668 | 0.01959936 | 1.25373197 | -84.7376633 | 0.00081517 | -32.9024963 | 0.00176479 | 12.1653633 | 33.2765465 | 0.86885631 | 0.00331808 | 0.00397729 | 34.1921463 | 0.57812953 |
| 0.00384014 | 0.0143271 | 0.54880595 | -60.7898712 | 0.00082008 | -37.0811653 | 0.00091069 | 7.25676632 | 36.9043503 | 0.58531624 | 0.00199012 | 0.0042851 | 34.1920967 | 0.2750358 |
| 0.00405338 | 0.01401131 | 0.59497124 | -54.8744392 | 0.00095982 | -38.9305267 | 0.00096931 | 7.05996275 | 39.264389 | 0.55976504 | 0.00194912 | 0.00477916 | 33.1966362 | 0.29125425 |
| 0.00609256 | 0.02328328 | 0.68675637 | -57.1764069 | 0.00060519 | -40.6395149 | 0.00114992 | 9.0711937 | 40.713913 | 0.74119669 | 0.00246939 | 0.00243277 | 27.3666935 | 0.34707007 |
| 0.00417393 | 0.0163713 | 0.63710016 | -70.3306198 | 0.00115382 | -39.8073769 | 0.00087064 | 6.93405247 | 40.2644119 | 0.63623315 | 0.00196468 | 0.00389 | 34.4212952 | 0.31541613 |
| 0.00524665 | 0.01583853 | 0.64737749 | -61.4570618 | 0.000864 | -38.9450531 | 0.00099755 | 8.00478649 | 38.8653221 | 0.6580233 | 0.00223738 | 0.00360564 | 23.9627018 | 0.30733526 |
| 0.00529812 | 0.01406999 | 1.09288132 | -88.4329224 | 0.00059918 | -34.9099617 | 0.00116727 | 8.62054348 | 33.4947586 | 0.78166306 | 0.00224362 | 0.00367518 | 38.7366295 | 0.53522342 |
| 0.00802593 | 0.00928487 | 1.119699 | -135.393982 | 0.00071679 | -29.514431 | 0.00095494 | 7.77871656 | 29.4324608 | 0.6378746 | 0.00187181 | 0.00437108 | 43.2764778 | 0.5360266 |
| 0.00596243 | 0.01321644 | 1.01030076 | -96.0092316 | 0.00090497 | -35.2701531 | 0.00102329 | 8.19835663 | 34.3101387 | 0.71146309 | 0.0022247 | 0.00431783 | 40.8662453 | 0.51160502 |
| 0.00798561 | 0.02168536 | 1.33620715 | -125.513954 | 0.00088095 | -36.2496605 | 0.00139956 | 10.8469057 | 36.0048561 | 0.81088668 | 0.00264413 | 0.00414053 | 41.3707962 | 0.5959115 |
| 0.00540116 | 0.01440222 | 1.24447751 | -90.5619965 | 0.00065157 | -33.4917984 | 0.00104605 | 8.88935566 | 33.2640877 | 0.8095777 | 0.00210166 | 0.00345139 | 43.8003616 | 0.59728956 |
| 0.00569612 | 0.0191456 | 0.6707868 | -56.1963272 | 0.00105356 | -37.2315865 | 0.00104169 | 10.198678 | 37.4116936 | 0.66042769 | 0.00273215 | 0.0042788 | 34.4401016 | 0.33396715 |
| 0.00547991 | 0.0173169 | 0.74503893 | -62.1792412 | 0.00070485 | -38.7232781 | 0.00107609 | 9.8869772 | 38.3132324 | 0.61368525 | 0.00261958 | 0.00424413 | 38.2606773 | 0.36835331 |
| 0.00650997 | 0.0202352 | 0.63046229 | -60.2714539 | 0.00128705 | -36.8351631 | 0.001158 | 9.86947918 | 37.3243561 | 0.65829545 | 0.00278973 | 0.00575866 | 33.4116516 | 0.31659597 |
| 0.0068827 | 0.01895196 | 0.82095599 | -75.76931 | 0.0006052 | -39.9449883 | 0.00114713 | 10.5976725 | 39.8464432 | 0.67388755 | 0.00285559 | 0.00416114 | 35.7182121 | 0.39980024 |
| 0.00745524 | 0.01656961 | 0.72176397 | -70.9955444 | 0.00050757 | -36.7466927 | 0.00108711 | 10.6530685 | 36.2699699 | 0.61996353 | 0.00285355 | 0.00388514 | 32.0656662 | 0.35001367 |
| 0.00687653 | 0.01447162 | 0.84813786 | -80.8616638 | 0.00060695 | -33.6192131 | 0.00111719 | 10.2300301 | 33.2824516 | 0.67759806 | 0.00267066 | 0.00310749 | 38.137825 | 0.40848127 |
| 0.00779872 | 0.01482246 | 0.91034395 | -94.456955 | 0.00070225 | -33.8125877 | 0.00111604 | 11.9854155 | 33.1647873 | 0.64660263 | 0.0030343 | 0.00344251 | 39.0995445 | 0.43387979 |
| 0.00604883 | 0.00958083 | 0.84637296 | -82.7041626 | 0.0006755 | -31.8329773 | 0.00106916 | 8.64340496 | 31.7780552 | 0.63924003 | 0.00224197 | 0.00304615 | 42.7820664 | 0.39813829 |
| 0.00701471 | 0.01385064 | 0.8118813 | -77.862587 | 0.00071151 | -33.2757568 | 0.00104536 | 9.92199802 | 32.9766617 | 0.66691893 | 0.00265171 | 0.00327063 | 39.9110451 | 0.38429147 |
| 0.00644905 | 0.01292228 | 0.77459586 | -79.8201752 | 0.00079095 | -33.7621422 | 0.00101862 | 9.95258427 | 32.8973618 | 0.65187943 | 0.00264416 | 0.00345507 | 39.5307961 | 0.37504441 |
| 0.00576729 | 0.01608559 | 1.24497795 | -102.483826 | 0.0005917 | -40.4425125 | 0.00095149 | 8.95220757 | 41.3436508 | 0.64108706 | 0.00229261 | 0.00394275 | 42.2753868 | 0.48793328 |
| 0.00560771 | 0.00863461 | 0.62525106 | -65.2267075 | 0.00086116 | -29.9199257 | 0.00098415 | 7.53595591 | 30.4914684 | 0.59526086 | 0.00188256 | 0.00318654 | 27.8354187 | 0.32217711 |
| 0.00772083 | 0.01766449 | 0.764265 | -95.4961777 | 0.00062906 | -33.8689728 | 0.00098754 | 9.57641602 | 34.2618637 | 0.6242851 | 0.00281201 | 0.00356884 | 34.9213104 | 0.37229744 |
| 0.0061673 | 0.01775595 | 0.90251643 | -101.620773 | 0.00060324 | -36.8343735 | 0.00106212 | 9.0284338 | 37.238224 | 0.69091082 | 0.0022902 | 0.00311286 | 36.0525818 | 0.44126713 |
| 0.00555738 | 0.01845465 | 0.66242188 | -56.7810822 | 0.00094334 | -36.3137474 | 0.00097718 | 8.7021904 | 37.0008316 | 0.65469676 | 0.00241969 | 0.00373639 | 28.0144959 | 0.32150096 |
| 0.00588063 | 0.01560072 | 0.84719545 | -78.4506226 | 0.00081842 | -31.7910271 | 0.00124735 | 7.96492577 | 31.3818359 | 0.7451033 | 0.00216062 | 0.00392912 | 31.491415 | 0.41294727 |
| 0.00533995 | 0.01374908 | 0.70392501 | -65.8348846 | 0.00116281 | -30.6559582 | 0.00109146 | 7.65303516 | 30.0661831 | 0.69834709 | 0.00210593 | 0.00378281 | 38.2507782 | 0.34933919 |
| 0.00595499 | 0.01278703 | 0.82546461 | -77.5253296 | 0.00067748 | -27.8875065 | 0.00114673 | 7.14064884 | 27.9540577 | 0.70074272 | 0.00203707 | 0.00359964 | 25.5175591 | 0.40664417 |
| 0.00573485 | 0.01248511 | 0.89581394 | -85.4755936 | 0.00075479 | -28.3695145 | 0.00122833 | 7.43336964 | 28.2174072 | 0.79746562 | 0.00198539 | 0.00363883 | 33.5708885 | 0.4286139 |
| 0.00492527 | 0.01240656 | 0.78720939 | -66.3566742 | 0.00084379 | -29.1730042 | 0.00106242 | 6.52896929 | 29.14361 | 0.70918655 | 0.00191365 | 0.00352739 | 28.2362499 | 0.37614757 |

| rel_to_neck_rad | d_minor_axis_ | d_rel_to_tail_l | ang_vel_head_ | motion_mode_ | width_tail_bas | d_curvature_n | ang_vel_tail_b | curvature_std_ | speed_hips_w_ | d_ang_vel_tail | d_rel_to_body | d_rel_to_body | d_curvature_n |
|---|---|---|---|---|---|---|---|---|---|---|---|---|---|
| -2.72016239 | -0.33750534 | 30.7887325 | 0.63039887 | 0.00444405 | 55.3605499 | 0.00324182 | 0.39470428 | 0.00143607 | 57.2017517 | 1.09492469 | -165.038651 | -121.411339 | 0.00024317 |
| -3.05469751 | -0.4216671 | 24.9592896 | 0.96625114 | 0.014994 | 54.153595 | 0.00445247 | 0.3381817 | 0.00150156 | 52.6919441 | 0.98077732 | -188.830414 | -87.6773682 | 0.00029767 |
| -4.50639153 | -0.31024218 | 27.9173412 | 0.52509964 | 0.0035535 | 55.2533913 | 0.00435343 | 0.36363483 | 0.00130693 | 37.6717606 | 0.94948006 | -170.210449 | -85.4704895 | 0.00032835 |
| -2.40770865 | -0.58218241 | 24.0814953 | 0.8252725 | 0.00288645 | 56.308876 | 0.00195869 | 0.22468476 | 0.00139247 | 34.9948845 | 0.91928321 | -153.472626 | -93.4711609 | 0.00019554 |
| -1.94723427 | -1.07159615 | 28.4707642 | 0.78277463 | 0.00277704 | 57.9947624 | 0.00205844 | 0.21294886 | 0.00121929 | 30.2712612 | 1.00736761 | -170.661163 | -94.9105225 | 0.00023549 |
| -2.0715816 | -1.75619125 | 27.2582073 | 0.98149413 | 0.02776667 | 57.9211731 | 0.00443419 | 0.36176357 | 0.00143115 | 64.6149216 | 1.11827362 | -152.614868 | -79.6775436 | 0.00026709 |
| -1.77039373 | -1.31239891 | 26.1357269 | 0.7242333 | 0.02911241 | 58.8861084 | 0.00530621 | 0.44450682 | 0.00098603 | 116.617577 | 1.05130363 | -169.296936 | -83.1433487 | 0.00026152 |
| -1.68879414 | -2.06502676 | 25.4019146 | 1.19535971 | 0.04277968 | 53.9669647 | 0.00411607 | 0.28980979 | 0.00154209 | 54.5797348 | 1.33775735 | -165.469406 | -73.6772995 | 0.00034066 |
| -2.86220503 | -3.38704586 | 31.9652901 | 1.1570183 | 0.02684474 | 57.704773 | 0.006196 | 0.47857931 | 0.00138801 | 110.14846 | 1.36485159 | -189.998276 | -90.6013107 | 0.00036514 |
| -2.37818146 | -2.21049786 | 29.9737377 | 1.28781939 | 0.03970895 | 55.0523949 | 0.00493782 | 0.3637135 | 0.00153922 | 62.4036293 | 1.52116394 | -191.121384 | -80.895813 | 0.0003947 |
| -1.66561174 | -0.47979951 | 24.6427803 | 0.6510247 | 0.01132176 | 57.2488632 | 0.00171769 | 0.18388082 | 0.00104677 | 34.8804894 | 0.78374887 | -137.915772 | -94.9487915 | 0.00018777 |
| -1.38110328 | -0.39834976 | 25.1431789 | 0.65796202 | 0.0106553 | 55.6079407 | 0.00192493 | 0.20559502 | 0.00105151 | 29.5209217 | 0.77661651 | -149.724381 | -91.6043854 | 0.00022734 |
| -4.86881781 | -0.38614273 | 29.4763451 | 0.87721729 | 0.00305298 | 55.6512909 | 0.00266252 | 0.23876858 | 0.00119968 | 31.4622116 | 0.95895672 | -166.801422 | -101.111763 | 0.00022737 |
| -0.76392531 | -0.32725334 | 30.9683666 | 0.61343378 | 0.00599361 | 59.9039002 | 0.00181622 | 0.21837246 | 0.001119 | 39.2501717 | 0.86571443 | -187.891296 | -111.494339 | 0.0001873 |
| -1.62458456 | -0.24850369 | 28.8574123 | 0.73438454 | 0.01399254 | 56.9691925 | 0.00196213 | 0.22254853 | 0.0011578 | 36.9613152 | 0.96080411 | -164.682938 | -86.530983 | 0.000205 |
| -2.73468447 | -2.06003189 | 28.5923653 | 0.89210463 | 0.03201001 | 66.7711563 | 0.00350413 | 0.28928238 | 0.00122114 | 58.5545998 | 1.22110283 | -185.342163 | -81.2308273 | 0.00028355 |
| -3.11561036 | -0.93369484 | 29.0510521 | 0.58889902 | 0.01638816 | 68.1054382 | 0.01080572 | 0.90414733 | 0.00131267 | 184.749329 | 0.97381449 | -188.988159 | -144.332199 | 0.00028702 |
| -3.16053534 | -1.1357069 | 28.5869751 | 0.83137393 | 0.02637248 | 63.4376869 | 0.0037084 | 0.36927131 | 0.00118145 | 62.5676575 | 1.03490865 | -163.383652 | -91.7392578 | 0.00029097 |
| -3.5658493 | -3.98752689 | 29.9409752 | 1.18843532 | 0.02487341 | 64.3317337 | 0.00711098 | 0.63215721 | 0.0013184 | 129.167572 | 1.30860043 | -228.664703 | -107.851746 | 0.00034888 |
| -2.8002336 | -1.72805786 | 30.3278332 | 0.74117667 | 0.04219597 | 66.0638733 | 0.00365723 | 0.30314523 | 0.00122499 | 63.8905792 | 1.23984039 | -217.083374 | -87.2422485 | 0.00029675 |
| -2.60376763 | -0.3235817 | 26.1075192 | 0.98380363 | 0.00647113 | 57.9451408 | 0.00174302 | 0.22725323 | 0.00129946 | 40.9589348 | 0.96084642 | -196.51387 | -92.2390976 | 0.00025579 |
| -2.3615315 | -0.77559948 | 27.0119572 | 0.9554432 | 0.01952605 | 55.9016686 | 0.00196092 | 0.21800315 | 0.00122157 | 33.4428406 | 0.90155512 | -197.396301 | -81.9735107 | 0.00025182 |
| -2.05105305 | -0.50349236 | 26.1230774 | 0.92974168 | 0.01043563 | 59.1915665 | 0.00207091 | 0.24720821 | 0.00133916 | 30.0908794 | 0.9513191 | -166.134995 | -91.7589874 | 0.00024496 |
| -1.68494201 | -1.16949081 | 26.2360458 | 1.10043764 | 0.02590405 | 56.253376 | 0.00260882 | 0.23407111 | 0.00130939 | 42.3446808 | 0.97993714 | -181.885986 | -87.4620209 | 0.00028321 |
| -2.531811 | -0.5158186 | 24.4890041 | 1.09071076 | 0.01859293 | 52.3243141 | 0.00229854 | 0.2099212 | 0.00126575 | 40.8909607 | 0.89598 | -175.664322 | -81.9527054 | 0.00026399 |
| -1.568362 | -1.3363719 | 26.9183292 | 1.04783356 | 0.03587968 | 64.4079742 | 0.00303893 | 0.2308999 | 0.00129064 | 47.2331467 | 0.99484336 | -159.918518 | -77.4859772 | 0.00024874 |
| -1.15322065 | -1.32555962 | 27.1194058 | 1.28139567 | 0.03830099 | 64.5008545 | 0.0031291 | 0.25758848 | 0.00138384 | 49.8023682 | 0.96848369 | -200.621674 | -79.1972427 | 0.00027042 |
| -1.74500608 | -0.68237782 | 27.4204617 | 0.69319797 | 0.03041876 | 61.6870346 | 0.00282814 | 0.23197395 | 0.00106606 | 47.7635384 | 0.94504917 | -166.210403 | -79.0454788 | 0.0002298 |
| -1.12118876 | -1.16701126 | 26.2059307 | 1.0229876 | 0.04069018 | 62.4728699 | 0.00347986 | 0.281443 | 0.00134101 | 54.7814102 | 0.98814476 | -157.812714 | -76.7478943 | 0.00021389 |
| -1.46759701 | -0.87735653 | 26.2732773 | 0.99883813 | 0.03612074 | 61.9859848 | 0.00262257 | 0.23126146 | 0.0012738 | 49.7202187 | 0.96889746 | -156.564194 | -74.2312164 | 0.00021698 |
| -3.06580162 | -1.37367249 | 28.6477642 | 0.94414699 | 0.01360748 | 59.3060188 | 0.00288523 | 0.36809877 | 0.00115371 | 85.7538757 | 0.96241367 | -239.622955 | -120.473488 | 0.00024897 |
| -0.79309458 | -0.3195405 | 25.9318066 | 0.52815998 | 0.00444168 | 59.4647141 | 0.00169528 | 0.2280777 | 0.00101659 | 34.1704369 | 0.85810173 | -209.812576 | -87.5959625 | 0.00024066 |
| -2.61364317 | -0.78498125 | 25.3786335 | 0.98467374 | 0.0163787 | 59.3921509 | 0.00339409 | 0.28968412 | 0.00132453 | 60.685318 | 0.91540188 | -180.773224 | -94.1817703 | 0.00025199 |
| -2.76851225 | -1.22051239 | 24.7409515 | 0.92393595 | 0.01465104 | 58.643055 | 0.00318765 | 0.45740131 | 0.00113882 | 106.51751 | 1.01349163 | -198.532745 | -100.252998 | 0.00023625 |
| -2.00054765 | -0.61278343 | 27.5816383 | 0.84971064 | 0.02329296 | 56.8415413 | 0.00195429 | 0.23172627 | 0.00121814 | 34.7565537 | 0.92549777 | -146.545624 | -85.6430206 | 0.00020257 |
| -2.12515831 | -1.01833344 | 27.6382351 | 0.77035224 | 0.0328247 | 59.8171158 | 0.00353772 | 0.42982853 | 0.00122213 | 73.7601318 | 1.11284685 | -135.739014 | -90.9204865 | 0.00025465 |
| -1.88949335 | -0.40588379 | 25.3119087 | 0.68912393 | 0.02510665 | 58.0089607 | 0.00250893 | 0.32896143 | 0.00107114 | 53.2956162 | 0.99735904 | -112.766182 | -81.3549347 | 0.00021873 |
| -2.37829661 | -0.95980167 | 24.1069336 | 0.7157349 | 0.0193256 | 60.6808853 | 0.00352649 | 0.33161545 | 0.00113438 | 62.4583206 | 0.96702248 | -123.618919 | -76.2291107 | 0.00029121 |
| -3.02741981 | -0.86050034 | 27.0639 | 0.68562412 | 0.0246568 | 61.8075905 | 0.00364931 | 0.4265044 | 0.00105604 | 80.8927841 | 1.11377895 | -129.871811 | -88.8003693 | 0.00031167 |
| -2.10306597 | -0.61509609 | 25.2088604 | 0.61545479 | 0.01510507 | 59.545887 | 0.00246615 | 0.26922002 | 0.00114179 | 44.9534912 | 0.97147036 | -111.45369 | -72.7116547 | 0.00025831 |

| d_rel_to_head | rel_to_neck_a | rel_to_head_b | d_length_w_b | curvature_hips | d_curvature_st | d_ang_vel_nec | d_speed_neck_ | d_rel_to_body | d_major_axis_ | d_curvature_st | d_path_curvat | d_rel_to_body | d_rel_to_hips_ | width_tail_base_w_forward_10th |
|---|---|---|---|---|---|---|---|---|---|---|---|---|---|---|
| 43.8909988 | 0.0438419 | -9.29111767 | -5.30112314 | 0.00518808 | 0.00129315 | 0.09469182 | -0.67125523 | -0.98103899 | -79.4706345 | 0.00037391 | 0.54429513 | 0.4870522 | -0.45514256 | 43.8992653 |
| 63.0061417 | 0.05574584 | -11.1265812 | -3.57276845 | 0.00569089 | 0.0013906 | 0.11046472 | -1.22990084 | -0.69989377 | -76.4592133 | 0.00043918 | 0.81691885 | 0.40239516 | -0.57217848 | 48.7274971 |
| 47.0572395 | 0.04454071 | -10.995863 | -8.08189392 | 0.00321682 | 0.00130105 | 0.11652988 | -0.01981496 | 0.83837813 | -76.7614365 | 0.00035522 | 0.31850502 | 0.43564349 | -0.41641781 | 50.9500237 |
| 48.2986832 | 0.03672851 | -9.28220749 | -3.68246222 | 0.00464038 | 0.00103134 | 0.05280021 | -1.5994612 | 0.53858423 | -61.9876289 | 0.00034439 | 1.16746926 | 0.26709527 | -0.45580742 | 50.4134026 |
| 41.2671738 | 0.04171243 | -8.6139946 | -4.95836163 | 0.00517866 | 0.00103256 | 0.075871 | -2.62779021 | 0.38738143 | -74.0176773 | 0.0003406 | 0.52056211 | 0.26638284 | -0.87611038 | 51.5269737 |
| 74.147583 | 0.04472694 | -11.4586535 | -2.6301558 | 0.00362596 | 0.00110923 | 0.09386142 | 1.00908113 | -0.49985319 | -82.5551987 | 0.00027147 | 2.55467939 | 0.54578942 | -0.5190981 | 52.4690552 |
| 70.4679413 | 0.05249623 | -11.6936102 | -1.97260249 | 0.00403922 | 0.00115784 | 0.10607826 | -1.15035868 | -0.86331844 | -82.6633453 | 0.00031539 | 2.67427278 | 0.52702785 | -0.77508265 | 55.8308373 |
| 70.151741 | 0.05822089 | -10.6535072 | -2.37126422 | 0.00384385 | 0.00141095 | 0.09386934 | -0.06332966 | -0.72183228 | -81.7106247 | 0.00032786 | 2.78741908 | 0.69258624 | -0.69896686 | 49.7848434 |
| 75.948616 | 0.06033097 | -11.5888958 | -3.04267716 | 0.00377099 | 0.00128955 | 0.1246782 | -0.77562797 | -1.23216569 | -98.657074 | 0.00028536 | 1.59588087 | 0.74997109 | -1.19470453 | 51.2768211 |
| 72.1561279 | 0.06691259 | -10.9177218 | -2.17416167 | 0.00370094 | 0.00145104 | 0.10729976 | 0.02716114 | -0.9599613 | -98.2453308 | 0.00033964 | 2.51208663 | 0.98944223 | -0.84601444 | 50.3254509 |
| 42.0378838 | 0.03290146 | -8.41590214 | -2.9509902 | 0.00343241 | 0.00090247 | 0.05946162 | -2.01750779 | 0.62837154 | -60.628624 | 0.00026337 | 1.11767706 | 0.25662795 | -0.585199 | 51.5745049 |
| 37.9384689 | 0.03830395 | -9.27753639 | -1.73533559 | 0.00366298 | 0.0010743 | 0.06483112 | -1.19639802 | 1.43194687 | -69.4996338 | 0.00028627 | 0.18732345 | 0.30402404 | -0.35147062 | 48.2978478 |
| 47.2181091 | 0.04600322 | -10.5133228 | -6.8279047 | 0.00544228 | 0.00111417 | 0.1248427 | -2.42633152 | -1.85347807 | -95.8396912 | 0.00030557 | 0.93266422 | 0.37921092 | -0.28924239 | 48.1542091 |
| 48.7900047 | 0.03414389 | -10.4931288 | -1.30082774 | 0.00532681 | 0.00100164 | 0.08527114 | -0.25007877 | 0.20110899 | -59.901619 | 0.00024274 | 1.25599039 | 0.34599328 | -0.31590527 | 51.2610855 |
| 43.0303268 | 0.03457631 | -9.35549831 | -1.87677598 | 0.00481849 | 0.00100485 | 0.06864144 | -0.54288673 | 0.07938117 | -62.9059982 | 0.00027501 | 0.20095256 | 0.33204916 | -0.28909022 | 50.6136589 |
| 65.0172958 | 0.04858185 | -11.7165184 | -3.10360265 | 0.00310083 | 0.00125418 | 0.08794549 | -1.54414392 | -1.18809712 | -127.131577 | 0.0002421 | 2.95330524 | 0.69413096 | -0.62147415 | 60.640213 |
| 81.1902161 | 0.05092929 | -13.25105 | -3.58669925 | 0.00291087 | 0.00109638 | 0.14597583 | -0.11673518 | -0.20730671 | -107.752571 | 0.00024416 | 1.34379387 | 0.47519779 | -0.36256161 | 53.1295662 |
| 69.9384918 | 0.05369118 | -11.90868 | -3.18566871 | 0.00361159 | 0.00108863 | 0.08929524 | -0.33543971 | -0.04307667 | -98.1107483 | 0.00024293 | 1.34072256 | 0.57704431 | -0.79185027 | 57.2523041 |
| 77.600853 | 0.06883032 | -12.1322861 | -4.03158331 | 0.00369938 | 0.00131976 | 0.14487313 | -2.08385849 | -1.19470239 | -124.242325 | 0.00027661 | 2.14682007 | 0.9450801 | -1.1699059 | 58.8485222 |
| 71.3189163 | 0.05920632 | -12.1040697 | -3.50838876 | 0.00313914 | 0.00111168 | 0.09150149 | -1.19151616 | -2.1939826 | -132.051498 | 0.00023468 | 2.73089743 | 0.72278339 | -0.63297659 | 60.0818253 |
| 49.0614166 | 0.04512648 | -11.1324062 | -3.23945427 | 0.00513573 | 0.00105663 | 0.07151423 | -1.52135217 | -0.27655453 | -75.1700974 | 0.00035741 | 0.64904726 | 0.31903204 | -0.20930478 | 54.1489296 |
| 49.7724609 | 0.04998378 | -11.6037912 | -3.0674839 | 0.00511849 | 0.00109993 | 0.06561908 | -2.23038864 | -0.32592517 | -76.8562317 | 0.00035713 | 1.29100192 | 0.35441351 | -0.73930997 | 51.9865532 |
| 49.4562645 | 0.04584275 | -9.99295425 | -2.73089695 | 0.00459304 | 0.00118521 | 0.08985266 | -1.71987665 | -0.38616926 | -68.7495422 | 0.0003588 | 1.12458074 | 0.32600385 | -0.77802503 | 51.285286 |
| 57.9477615 | 0.05251209 | -10.817359 | -1.57480991 | 0.00400946 | 0.0011768 | 0.09033778 | -1.50705147 | -0.8124674 | -76.9747162 | 0.00038499 | 1.25241494 | 0.41053006 | -1.05096054 | 51.1738052 |
| 57.7338486 | 0.04707624 | -9.98952389 | -3.60868621 | 0.00441826 | 0.00108757 | 0.09203408 | -1.04520357 | -0.20621274 | -71.8055725 | 0.00039134 | 0.65772796 | 0.35678506 | -0.55399913 | 49.7667084 |
| 72.5420151 | 0.04300747 | -11.0677509 | -2.66903877 | 0.0030574 | 0.00106806 | 0.08163862 | -0.19319922 | -0.53485555 | -80.4000473 | 0.0002876 | 3.01643205 | 0.44693804 | -0.57464236 | 58.7345085 |
| 89.9403381 | 0.05382709 | -13.8009892 | -1.90748584 | 0.00327554 | 0.00111669 | 0.08861801 | 0.80990821 | -0.07276341 | -93.0700684 | 0.0003769 | 2.5911994 | 0.48098713 | -0.81358445 | 59.5895462 |
| 76.2874146 | 0.04420646 | -11.9747791 | -2.38306093 | 0.00312004 | 0.00098932 | 0.08095846 | 0.67108119 | 0.11181579 | -84.7321472 | 0.00026839 | 1.68848896 | 0.42513961 | -0.56395781 | 56.6709099 |
| 75.1908569 | 0.04040535 | -11.5326023 | -2.06728935 | 0.00323679 | 0.00097881 | 0.08451445 | 1.0606606 | -0.10143518 | -78.7381134 | 0.00032213 | 1.8254292 | 0.42722622 | -0.80167568 | 55.4302063 |
| 75.8337479 | 0.04327429 | -11.709507 | -1.56049609 | 0.00298201 | 0.0009687 | 0.07696484 | 0.22450028 | -0.61204082 | -78.5661316 | 0.00030799 | 2.36980844 | 0.43607298 | -0.65909398 | 58.2562714 |
| 59.6807594 | 0.05272211 | -13.2818832 | -4.07823563 | 0.00286095 | 0.00098317 | 0.08449395 | -3.42603254 | -1.19829512 | -107.320595 | 0.00029612 | 0.15967982 | 0.60879147 | -0.6630491 | 55.6076202 |
| 51.4577942 | 0.05070612 | -9.68861961 | 0.78862715 | 0.00282235 | 0.00099872 | 0.05530656 | -0.91545993 | -0.82142282 | -81.8081665 | 0.0003685 | 0.80483806 | 0.257949 | -0.41454491 | 51.1308975 |
| 65.6095276 | 0.05516936 | -12.5745258 | -4.12106228 | 0.0043055 | 0.00107379 | 0.11691114 | -0.29722199 | -0.54242158 | -75.1380539 | 0.00037201 | 0.49691871 | 0.39257976 | -0.47132117 | 57.0834427 |
| 58.3625565 | 0.05178631 | -11.6106319 | -2.87356806 | 0.00321752 | 0.0009747 | 0.10256242 | -0.73326683 | -0.93104148 | -90.4499817 | 0.00031026 | 0.52622563 | 0.5466789 | -0.52729034 | 56.2790451 |
| 52.5419731 | 0.03597426 | -9.99479103 | -3.33639288 | 0.00400665 | 0.00097284 | 0.0723102 | 0.01691404 | 0.22208827 | -74.891571 | 0.0003098 | 0.99223566 | 0.35718226 | -0.3378723 | 50.0007629 |
| 56.7455559 | 0.03986714 | -8.76600266 | -3.34231615 | 0.00336284 | 0.00107283 | 0.10166004 | -0.39382815 | -0.16779456 | -75.27668 | 0.00023023 | 1.17230225 | 0.53607905 | -0.34355605 | 55.1646004 |
| 54.9970856 | 0.03348539 | -8.39079762 | -3.10343456 | 0.00368824 | 0.00092677 | 0.07906774 | 0.75458068 | -0.02649017 | -69.66259 | 0.00023532 | 0.69966602 | 0.36785293 | -0.35672814 | 52.7804146 |
| 55.5688858 | 0.04042415 | -7.5851078 | -3.88050413 | 0.00384834 | 0.00105328 | 0.09158738 | -0.05438014 | -0.30943957 | -71.1220703 | 0.00022232 | 0.47480452 | 0.35919535 | -0.32529861 | 55.4889069 |
| 54.4505081 | 0.04530213 | -8.01202679 | -3.83950973 | 0.00294987 | 0.00098425 | 0.09728859 | -0.60303128 | 0.54575557 | -82.9036713 | 0.00020911 | 1.59725618 | 0.43865594 | -0.32096228 | 57.3357925 |
| 57.3047867 | 0.0344226 | -8.23572731 | -3.16146755 | 0.00351079 | 0.00091814 | 0.08127739 | 0.48085213 | 0.00118505 | -69.3158112 | 0.00022804 | 0.48489949 | 0.34801984 | -0.23694438 | 55.5756722 |
