## Supplemental Table 1(3) for "Rebalancing the motor circuit restores movement in a *Caenorhabditis elegans* model for TDP-43-toxicity"

| worm_names | motion_mode_ | motion_mode_ | curvature_std_d | curvature_n | ang_vel_tail_ti | curvature_std_ | rel_to_head_b | width_tail_bas | minor_axis_w_d | rel_to_head_d | rel_to_body_ | rel_to_head_b | rel_to_body_rad |
|---|---|---|---|---|---|---|---|---|---|---|---|---|---|
| hTDP-43_20C | 0.018192535 | 0.533940247 | 0.004906863 | 0.010545515 | 0.622973615 | 0.002139234 | -0.280496631 | 58.6053551 | 109.5435392 | 2.319621222 | 256.6804177 | 0.382051686 | -60.00184638 |
| hTDP-43_25C | 0.017016281 | 0.622362248 | 0.004690056 | 0.009287602 | 0.51172099 | 0.001898178 | -0.246155679 | 55.87494353 | 106.7620299 | 2.256429486 | 243.392806 | 0.366218559 | -43.25115743 |
| clh-3(ok763)II | 0.041939227 | 0.42350346 | 0.003912099 | 0.007251564 | 0.403385058 | 0.002602589 | -0.104560882 | 68.02299118 | 135.7241593 | 1.838262856 | 266.925415 | 0.395005495 | -33.4989605 |
| egl-42(n995)II | 0.027896145 | 0.288842245 | 0.003489907 | 0.008263709 | 0.507211636 | 0.002106454 | -0.207087186 | 75.78104342 | 106.6817028 | 2.252483959 | 280.9539947 | 0.409482469 | -40.20489084 |
| egl-50(n1086)II | 0.037327526 | 0.382500013 | 0.002875891 | 0.006377593 | 0.430371981 | 0.001531977 | -0.002607185 | 79.79548462 | 133.1128152 | 2.186398435 | 246.4780231 | 0.341831423 | -37.12480164 |
| unc-9(e101)X | 0.025864072 | 0.174840853 | 0.004399289 | 0.010519662 | 0.436835333 | 0.002371067 | -0.643442026 | 66.75863476 | 88.98151855 | 3.406576991 | 407.3491364 | 0.714942148 | -35.53465891 |
| unc-77(gk9)IV | 0.031022028 | 0.275268313 | 0.003310469 | 0.009176045 | 0.532246863 | 0.001842022 | -0.139275525 | 70.52527539 | 116.7020206 | 2.332092539 | 334.5235367 | 0.411608002 | -34.78473091 |
| C24G7.1(ok1822)II | 0.000889225 | 0.004011582 | 0.003356183 | 0.01732719 | 1.256806636 | 0.002198288 | -0.294793862 | 75.6475708 | 120.2838944 | 2.827907085 | 435.7892883 | 0.578996932 | -55.49352951 |
| cat-2(e1112)II | 0.011544792 | 0.160868844 | 0.002937667 | 0.016710915 | 1.668207407 | 0.00175872 | 0.009402621 | 78.95588493 | 135.9485855 | 2.750572324 | 694.1011505 | 0.53934525 | -145.2312584 |
| egl-30(ep271)I | 0.002265452 | 0.016909802 | 0.003808389 | 0.029256685 | 2.577332388 | 0.003031419 | -0.217372306 | 73.88448207 | 150.4724231 | 4.135085479 | 586.4503267 | 0.849250184 | -170.7361016 |
| gon-2(q362)I | 0.022399454 | 0.138776143 | 0.003352801 | 0.011144384 | 0.700295684 | 0.00238154 | -0.207913255 | 76.96950947 | 115.9214301 | 2.868896181 | 357.944723 | 0.537632102 | -51.32275443 |
| gpa-9(pk438)V | 0.019606928 | 0.153711502 | 0.003328045 | 0.011950934 | 0.746661446 | 0.002193981 | -0.135106705 | 79.09006405 | 127.1621935 | 2.295905071 | 328.6909687 | 0.440514704 | -54.46960037 |
| gpb-2(sa603)I | 0.022119489 | 0.18507198 | 0.002902807 | 0.009062328 | 0.617337646 | 0.001847874 | -0.285944587 | 89.02498043 | 132.2655873 | 2.359958591 | 303.3630534 | 0.444928953 | -48.59048179 |
| gpc-1(pk298)X | 0.018649179 | 0.122600662 | 0.003075174 | 0.012375646 | 0.789922124 | 0.002105278 | -0.28780668 | 79.56737518 | 114.3119283 | 2.243708019 | 391.9215914 | 0.473942032 | -54.91254142 |
| ins-18(ok1672)I | 0.015760159 | 0.19178761 | 0.00278457 | 0.009162595 | 0.667771695 | 0.001574587 | -0.217953372 | 70.66766866 | 110.6125634 | 1.761522412 | 316.0448208 | 0.349208937 | -43.19850246 |
| ins-22(ok3616)III | 0.016059709 | 0.278318404 | 0.003021607 | 0.011204817 | 0.822342756 | 0.002207518 | -0.279893359 | 77.43160082 | 120.3995212 | 2.498839207 | 366.1591094 | 0.432411633 | -68.56086581 |
| ins-25(ok2773)I | 0.029541663 | 0.515140056 | 0.003115225 | 0.009971245 | 0.677768067 | 0.002359775 | -0.121588368 | 71.87534004 | 142.2674462 | 1.891690599 | 342.1175701 | 0.267773309 | -67.15066105 |
| ins-3(ok2488)II | 0.027468527 | 0.312979651 | 0.003816563 | 0.010463797 | 0.663393948 | 0.002185322 | -0.254116422 | 73.69208395 | 122.4943252 | 2.288458104 | 362.4105225 | 0.418085146 | -61.10010272 |
| ins-35(ok3297)V | 0.021697168 | 0.193727817 | 0.003310055 | 0.010910524 | 0.854962682 | 0.002315929 | -0.200505446 | 73.29563405 | 128.4761118 | 2.280357489 | 378.6926105 | 0.452990589 | -70.63577608 |
| ins-4(ok3534)II | 0.033328589 | 0.357024236 | 0.003460906 | 0.008801521 | 0.576070736 | 0.002238399 | -0.208061843 | 74.53350639 | 132.1254727 | 2.219992891 | 357.5305144 | 0.405875283 | -46.81528394 |
| flr-1(ut11)X | 0.035803207 | 0.36203861 | 0.004545554 | 0.009583861 | 0.455707724 | 0.002713476 | -0.213916663 | 51.0511726 | 111.6889971 | 1.684236012 | 197.7365466 | 0.277196481 | -36.15961751 |
| lite-1(ce314)X | 0.030320056 | 0.352620531 | 0.003471415 | 0.007701015 | 0.399177894 | 0.002151443 | -0.163058097 | 74.14054775 | 116.2601948 | 2.909087032 | 353.838089 | 0.487505235 | -40.11975026 |
| mir-124(n4255)IV | 0.021337283 | 0.158698748 | 0.003082535 | 0.011028433 | 0.762319665 | 0.00208088 | -0.23533953 | 78.75367336 | 116.931283 | 2.742427406 | 419.8999069 | 0.50790652 | -63.50734573 |
| mec-12(u76)III; bzIs17[pmec-4:: | 0.040945332 | 0.277003336 | 0.002937126 | 0.007937976 | 0.578244414 | 0.002037941 | -0.113419915 | 78.82426961 | 124.8602334 | 1.906447627 | 299.670886 | 0.357993671 | -45.01625298 |
| nlp-1(ok1469)X | 0.015578721 | 0.244540915 | 0.003798366 | 0.010281252 | 0.516608784 | 0.001967585 | -0.445112866 | 70.64437223 | 131.9488355 | 2.199054987 | 335.7821834 | 0.419504624 | -40.48259997 |
| nzIs29[punc-17::rho-1(G14V); p | 0.016919279 | 0.401329945 | 0.007004664 | 0.02123063 | 1.04333831 | 0.003399934 | -0.36751662 | 60.49092048 | 143.1707738 | 3.570119631 | 409.1638968 | 0.762630511 | -91.83395586 |
| tmc-1(ok1859)X | 0.002123564 | 0.015851423 | 0.003073528 | 0.0147704 | 1.524820461 | 0.001734822 | -0.3750011 | 74.94385425 | 121.9755051 | 3.090758871 | 544.1969299 | 0.813338137 | -137.4454626 |
| ser-4(ok512)III | 0.017168262 | 0.128937599 | 0.003439652 | 0.012363026 | 0.870752715 | 0.002434032 | -0.234938782 | 78.05789003 | 123.3903485 | 2.766154091 | 340.7757423 | 0.523488629 | -53.98374794 |
| unc-108(n501)I | 0.034776128 | 0.569429027 | 0.003939296 | 0.005704611 | 0.305112805 | 0.002208915 | -0.143344509 | 83.63742337 | 99.02620782 | 2.621389754 | 280.9639842 | 0.487844471 | -29.887204 |
| acr-10(ok3064)X | 0.008677417 | 0.046314146 | 0.003301549 | 0.013179062 | 0.928366363 | 0.00224104 | -0.250807049 | 79.07959938 | 120.0032894 | 2.931003511 | 397.9917709 | 0.552268271 | -62.16795177 |
| acr-11(ok1345)I | 0.008226683 | 0.052284763 | 0.003284822 | 0.01436437 | 0.946912319 | 0.002277439 | -0.210631159 | 76.46811848 | 117.3356879 | 3.041445009 | 380.7220095 | 0.56248557 | -56.71076399 |
| lev-8(x15)X | 0.01802821 | 0.114706933 | 0.002991192 | 0.011222748 | 0.792078146 | 0.002055224 | -0.076709023 | 81.04426438 | 111.3638374 | 2.382223959 | 330.5670471 | 0.429487831 | -56.41499424 |
| acr-14(ok1155)II | 0.018682763 | 0.115255522 | 0.003193566 | 0.01126252 | 0.663955232 | 0.002068438 | -0.241702768 | 80.27079352 | 119.0449127 | 2.738021916 | 332.9078027 | 0.494784736 | -45.7929844 |
| acr-15(ok1214)V | 0.016754335 | 0.116013847 | 0.003582282 | 0.013601439 | 0.922072679 | 0.002393081 | -0.235770544 | 76.16208627 | 125.2941271 | 2.719761348 | 374.7185329 | 0.510647937 | -62.81966008 |
| acr-18(ok1285)V | 0.004183146 | 0.024929694 | 0.003523714 | 0.015784009 | 1.111355438 | 0.002235461 | -0.306543964 | 72.93904659 | 112.9463494 | 3.376989864 | 387.907632 | 0.59281165 | -66.63978776 |
| acr-19(ad1674)I | 0.013417262 | 0.09020416 | 0.003273571 | 0.012041913 | 0.857927047 | 0.002189995 | -0.257933036 | 78.38009042 | 114.723345 | 2.611984907 | 344.8351676 | 0.498545775 | -55.94008012 |
| acr-2(ok1887)X | 0.013330203 | 0.108736948 | 0.00326832 | 0.012953403 | 0.879023584 | 0.002190818 | -0.280113349 | 80.59694481 | 124.6988389 | 2.806534324 | 394.6908646 | 0.531262509 | -62.20682941 |
| acr-21(ok1314)III | 0.006532952 | 0.085820667 | 0.003166637 | 0.015783526 | 1.211274847 | 0.00213948 | -0.212331245 | 80.52164596 | 120.2841911 | 3.014469959 | 437.7992576 | 0.562688217 | -71.55542232 |
| acr-23(ok2804)V | 0.018832905 | 0.202836048 | 0.003146076 | 0.010617196 | 0.740347172 | 0.002074478 | -0.227038483 | 78.18383462 | 115.1080649 | 2.350785052 | 315.2940442 | 0.465216066 | -61.69144419 |
| acr-3(ok2049)X | 0.008981828 | 0.059431568 | 0.003539379 | 0.01479806 | 0.986843871 | 0.002272204 | -0.258446278 | 75.82518938 | 118.2454619 | 2.934331642 | 391.5630278 | 0.552998674 | -61.24284236 |
| acr-6(ok3117)I | 0.007192799 | 0.044154743 | 0.003278395 | 0.015428204 | 1.051638273 | 0.002148814 | -0.230495775 | 79.54300935 | 122.5072493 | 3.07668301 | 415.9414782 | 0.574594117 | -62.77049569 |
| acr-7(tm863)II | 0.027715093 | 0.222491074 | 0.003333847 | 0.008658643 | 0.610708456 | 0.002037971 | -0.222433062 | 77.21559931 | 112.2891429 | 2.426095005 | 321.5535823 | 0.4757675 | -41.69600264 |
| acr-9(ok933)X | 0.011697945 | 0.068123576 | 0.003524976 | 0.014583915 | 1.00494385 | 0.002213381 | -0.31184377 | 75.2619939 | 130.0987153 | 2.986263721 | 373.0215922 | 0.545002677 | -69.92824259 |
| asic-1(ok415)I | 0.040010318 | 0.381913208 | 0.0032775 | 0.007948591 | 0.471716825 | 0.002469746 | 0.041795615 | 77.44496187 | 139.4047082 | 1.813961213 | 267.5357418 | 0.287543853 | -41.61584449 |
| bas-1(ad446)III | 0.011561078 | 0.148525526 | 0.003745804 | 0.014899347 | 1.063541314 | 0.002327933 | -0.436100001 | 78.49137582 | 131.1232736 | 3.065756644 | 450.8669093 | 0.632913619 | -84.32989749 |
| cat-4(e1141)V | 0.021275902 | 0.186243252 | 0.003540334 | 0.012035589 | 0.748787774 | 0.002321071 | -0.462855908 | 73.09953213 | 126.857193 | 3.056196213 | 442.5399297 | 0.551485725 | -62.27972857 |
| ced-1(e1735)I | 0.027809199 | 0.202128997 | 0.003135072 | 0.007554624 | 0.504882763 | 0.002170097 | -0.096187448 | 75.34712982 | 90.42413902 | 1.928135574 | 253.8671455 | 0.370785534 | -32.89291668 |
| ocr-3(ok1559)X | 0.015695223 | 0.093422743 | 0.003343264 | 0.012164419 | 0.934573177 | 0.002293084 | -0.319766972 | 77.45222231 | 134.3794616 | 2.812253768 | 477.7089374 | 0.621827132 | -79.93265468 |
| daf-3(e1376)X | 0.024527017 | 0.224390461 | 0.003107343 | 0.010011104 | 0.661890806 | 0.002073728 | -0.177536898 | 80.96404739 | 121.0673556 | 2.378430832 | 356.3530557 | 0.424696169 | -48.52642995 |
| daf-5(e1386)II | 0.022065359 | 0.230909408 | 0.003310132 | 0.01053634 | 0.772312439 | 0.002154367 | -0.136245916 | 80.15201537 | 139.0896352 | 2.101598794 | 356.4642111 | 0.375081983 | -55.11824068 |
| daf-7(m62)III | 0.020832982 | 0.120675335 | 0.002816268 | 0.008791995 | 0.629478235 | 0.001819556 | -0.059260859 | 82.27969395 | 125.2714618 | 2.560355132 | 385.4381062 | 0.461151875 | -48.67491566 |
| dat-1(ok157)III | 0.011480975 | 0.058823892 | 0.003259814 | 0.012791051 | 0.814480146 | 0.002088188 | -0.265475391 | 83.48225923 | 118.6851286 | 2.666301922 | 357.26828 | 0.512723091 | -54.54539039 |
| unc-116(e2310)III | 0.026877357 | 0.646836569 | 0.003426302 | 0.006911285 | 0.418828133 | 0.001913906 | 0.040257644 | 76.9153856 | 95.4261298 | 1.786588082 | 197.0610107 | 0.220080932 | -36.82227275 |
| del-1(ok150)X | 0.025008694 | 0.172783128 | 0.003297929 | 0.011240571 | 0.725351633 | 0.002224733 | -0.172575764 | 73.54569421 | 120.4431989 | 2.360322624 | 312.9700203 | 0.462828839 | -47.92772341 |
| dnc-1(or404)IV | 0.012481935 | 0.161923404 | 0.003382613 | 0.011949439 | 0.758055293 | 0.002054366 | -0.096412826 | 81.53836823 | 119.1502745 | 2.679020252 | 341.6110567 | 0.528717073 | -52.62108571 |
| dop-1(vs101)X | 0.023726433 | 0.220747515 | 0.002848672 | 0.008008956 | 0.522076994 | 0.001559798 | -0.422934651 | 73.43077342 | 124.0117671 | 1.977995396 | 333.6308492 | 0.395603309 | -36.73054187 |
| dop-2(vs105)V | 0.026182382 | 0.299068115 | 0.003310692 | 0.01091044 | 0.72960673 | 0.001964636 | -0.09407636 | 82.62771301 | 121.499205 | 2.313941502 | 345.4512176 | 0.386039326 | -60.08975601 |
| dop-2(vs105)V; dop-3(vs106)X | 0.035624271 | 0.38926329 | 0.003158293 | 0.008380347 | 0.610511899 | 0.00138751 | 0.141291857 | 80.24346161 | 128.8059087 | 2.155893326 | 413.8879395 | 0.291687906 | -60.16529083 |
| dop-3(vs106)X | 0.038137761 | 0.620330844 | 0.003122397 | 0.005659846 | 0.462441832 | 0.001568369 | 0.029864438 | 83.67870331 | 169.7440033 | 1.956972957 | 391.7306214 | 0.229362458 | -49.80765343 |
| dpy-20(e1282)IV | 0.023018266 | 0.186546387 | 0.003572656 | 0.012543811 | 0.737737375 | 0.002040553 | -0.146024992 | 75.96300233 | 89.05501283 | 2.482367788 | 370.3104135 | 0.456266678 | -37.09030127 |
| odr-3(n2150)V | 0.017641162 | 0.211832204 | 0.003617594 | 0.011959507 | 0.823363459 | 0.002289828 | -0.362642322 | 76.58059156 | 111.9083388 | 2.595367511 | 368.8781942 | 0.521482324 | -55.86915737 |
| eat-16(sa609)I | 0.001177568 | 0.005111939 | 0.004367291 | 0.034680446 | 2.464317685 | 0.003357347 | -0.145790195 | 61.53519749 | 111.7251176 | 4.982576438 | 578.9784517 | 0.874905058 | -144.3425093 |
| egl-11(n587)V | 0.025558347 | 0.291664538 | 0.003895009 | 0.01013371 | 0.592766284 | 0.002426443 | -0.154258555 | 68.03834182 | 143.1724396 | 2.361325151 | 294.8230037 | 0.429522799 | -59.02487298 |
| egl-2(n693)V | 0.010805413 | 0.093024154 | 0.003130113 | 0.013201089 | 1.015454707 | 0.00206694 | -0.264946589 | 76.83606949 | 135.3691902 | 2.666418827 | 406.0968216 | 0.506224626 | -73.49943514 |
| egl-1(n487)V | 0.029573255 | 0.396468859 | 0.003257063 | 0.007899313 | 0.568918676 | 0.001620925 | -0.077955447 | 84.89748822 | 129.3267579 | 1.934527466 | 336.5578525 | 0.339378378 | -60.97964183 |
| egl-10(md176)V | 0.030594018 | 0.456004739 | 0.003237123 | 0.006088901 | 0.415088097 | 0.001534776 | -0.009390466 | 76.59346164 | 133.7637015 | 1.329984408 | 237.3047081 | 0.245520719 | -45.10524633 |
| egl-14(n549)X | 0.029638001 | 0.636273532 | 0.003392754 | 0.005646135 | 0.378048001 | 0.001400155 | 0.001778248 | 73.8852635 | 129.1640113 | 1.59845123 | 260.4975396 | 0.217808086 | -51.58463484 |
| egl-12(n602)V | 0.014541108 | 0.880292808 | 0.004093319 | 0.008917219 | 0.593265898 | 0.001898502 | -0.216240788 | 88.00142924 | 133.7291549 | 2.102239549 | 428.7347349 | 0.346646196 | -53.4698407 |
| egl-13(n483)X | 0.024888904 | 0.352686969 | 0.003592433 | 0.009535174 | 0.592816522 | 0.001842872 | -0.160861558 | 80.21835767 | 131.3452407 | 1.905362918 | 311.0961979 | 0.337276641 | -54.20076385 |
| egl-15(n484)X | 0.019985262 | 0.488125794 | 0.003431413 | 0.007650744 | 0.542677273 | 0.001506832 | -0.158760831 | 80.29222361 | 121.219138 | 1.806221298 | 334.3663853 | 0.278816 | -49.91974304 |
| egl-17(e1313)X | 0.027214232 | 0.298726528 | 0.003346009 | 0.009538103 | 0.507561733 | 0.002384163 | -0.101519364 | 73.45473157 | 112.7814944 | 1.958958809 | 231.7492576 | 0.329313066 | -33.23875104 |
| egl-18(ok290)IV | 0.031294063 | 0.576446867 | 0.003447815 | 0.006308129 | 0.440382356 | 0.001751709 | -0.079241726 | 78.87671248 | 119.9593384 | 1.533646837 | 264.164793 | 0.245396566 | -48.19786318 |
| egl-19(n2368)IV | 0.007234902 | 0.093832961 | 0.004739651 | 0.019462387 | 1.179215322 | 0.002494896 | -0.063433623 | 66.0478197 | 93.99268656 | 2.41699542 | 368.1068752 | 0.513544866 | -71.24528418 |
| egl-20(mu39)IV | 0.021364858 | 0.442921962 | 0.003581979 | 0.010649202 | 0.795298266 | 0.002351408 | 0.094662521 | 82.76700127 | 117.987306 | 2.089026718 | 309.9300819 | 0.333701058 | -57.08102913 |
| egl-21(n476)IV | 0.027502795 | 0.201820412 | 0.002950469 | 0.007716059 | 0.520375878 | 0.001652165 | -0.105482003 | 82.64650238 | 166.0763193 | 2.967483273 | 375.3079553 | 0.51798208 | -65.8248114 |
| egl-21(n611)IV | 0.010591077 | 0.063463627 | 0.002929698 | 0.009986956 | 0.699709314 | 0.00188387 | -0.185972015 | 81.98867798 | 142.8326387 | 3.212328245 | 426.490195 | 0.60130796 | -86.65112909 |
| egl-23(n601)IV | 0.010253348 | 0.544981475 | 0.003266486 | 0.008768129 | 0.575747142 | 0.00142178 | -0.131304099 | 87.10017621 | 104.5865846 | 1.78433682 | 265.252579 | 0.254376757 | -43.45055167 |
| egl-27(ok151)II | 0.021086316 | 0.249643096 | 0.004768084 | 0.014041079 | 0.789636364 | 0.002776276 | -0.274217165 | 59.07854323 | 106.4551579 | 2.527083489 | 272.2020118 | 0.446790184 | -60.13327235 |
| egl-24(n572)III | 0.029187578 | 0.238230724 | 0.00308652 | 0.008812231 | 0.555576143 | 0.001815757 | -0.234699257 | 80.03552662 | 120.7154531 | 2.3491813 | 334.8340843 | 0.448900105 | -54.3702377 |
| egl-28(n570)II | 0.018175563 | 0.209394681 | 0.003562219 | 0.009763558 | 0.539067808 | 0.001810451 | -0.19908096 | 76.52905443 | 111.2906107 | 1.969958177 | 243.1521601 | 0.37349949 | -44.12253959 |
| egl-30(n686)goa-1(n1134)I | 0.027969022 | 0.525809882 | 0.00305616 | 0.005456422 | 0.312956223 | 0.001401418 | -0.086271675 | 74.19494556 | 94.91692098 | 1.453051542 | 199.5491159 | 0.277910005 | -29.70193808 |
| egl-31(n472)I | 0.022399565 | 0.552211953 | 0.003822272 | 0.006485891 | 0.519550607 | 0.00123149 | -0.206756073 | 83.0687832 | 106.8448699 | 1.624673488 | 261.6788477 | 0.273505106 | -37.24551232 |
| egl-32(n155)I | 0.031204594 | 0.266858099 | 0.003241745 | 0.008455851 | 0.674840122 | 0.001782504 | 0.018046516 | 91.02689743 | 128.8828945 | 2.451207483 | 345.4203079 | 0.364043644 | -72.95116291 |
| egl-33(n151)I | 0.030425743 | 0.224187791 | 0.003543214 | 0.009355317 | 0.607688508 | 0.002034957 | -0.224672326 | 80.67725107 | 119.5637857 | 2.35496696 | 339.8111982 | 0.431805326 | -63.00761587 |
| egl-36(n728)X | 0.006140933 | 0.298648084 | 0.002742731 | 0.010109818 | 0.882631478 | 0.001602962 | -0.140725954 | 86.92654488 | 122.4885115 | 1.836002986 | 360.8653995 | 0.334082593 | -57.58832486 |
| egl-37(n742)II | 0.025553252 | 0.188210514 | 0.003103118 | 0.009772151 | 0.660950922 | 0.001997471 | -0.218040926 | 82.11749366 | 122.023551 | 2.605889317 | 365.1837217 | 0.464646568 | -53.67061258 |
| pkg-1(n478)IV | 0.017618395 | 0.126858528 | 0.002668673 | 0.008877839 | 0.861736991 | 0.001625598 | -0.068406765 | 87.27314949 | 111.9569035 | 2.411008621 | 362.174422 | 0.468159426 | -74.84220362 |
| egl-40(n606)IV | 0.010777019 | 0.128196764 | 0.003156319 | 0.011917754 | 0.969950695 | 0.001966175 | -0.147858479 | 76.27925992 | 96.66358003 | 2.538845593 | 324.7468247 | 0.489447715 | -78.70684273 |
| egl-47(n1081)V | 0.018260949 | 0.356368563 | 0.003207887 | 0.008517992 | 0.705701444 | 0.001689114 | -0.309254679 | 78.69650284 | 117.9774378 | 1.82779236 | 305.1460556 | 0.414501823 | -51.69309331 |
| egl-43(n1079)II | 0.026143433 | 0.258811063 | 0.004135211 | 0.01672152 | 1.123866242 | 0.00219929 | -0.142709843 | 74.46860194 | 160.9987224 | 2.998026909 | 430.6306002 | 0.418075457 | -90.18826389 |
| egl-44(n1080)II | 0.018012664 | 0.211394255 | 0.003173001 | 0.010678055 | 0.697698737 | 0.001723609 | -0.160415442 | 78.823729 | 131.9446476 | 2.427829958 | 375.6248234 | 0.439073278 | -65.85045198 |
| egl-46(n1127)V | 0.030476614 | 0.313933083 | 0.00348799 | 0.008189065 | 0.55689409 | 0.001617832 | -0.241195152 | 82.46408871 | 132.0504431 | 2.022221438 | 310.7349216 | 0.37135004 | -60.74069752 |
| egl-49(n1107)X | 0.037979045 | 0.333490947 | 0.002997051 | 0.006067517 | 0.411639911 | 0.001714007 | -0.173425908 | 83.32086978 | 112.1341815 | 2.023345657 | 288.834472 | 0.388658938 | -40.69458936 |
| egl-5(n486)III | 0.035701332 | 0.471550206 | 0.003553647 | 0.007501959 | 0.645501906 | 0.001816992 | -0.130050165 | 86.60310884 | 121.9666925 | 2.09426189 | 270.1466203 | 0.331282921 | -118.2118737 |
| egl-6(n592)X | 0.031538088 | 0.647917725 | 0.003450083 | 0.005285325 | 0.314289839 | 0.001933802 | 0.042608278 | 84.36193593 | 102.2867406 | 1.59063205 | 212.9063993 | 0.234934193 | -36.55655829 |
| egl-7(n575)III | 0.024828676 | 0.452808323 | 0.003570328 | 0.007276383 | 0.425537957 | 0.001893515 | -0.065667175 | 86.33615657 | 134.0725149 | 2.11061646 | 312.1455367 | 0.305411801 | -54.84017281 |
| egl-8(n488)V | 0.025270253 | 0.261792507 | 0.003153997 | 0.008187677 | 0.553181702 | 0.00188382 | -0.182377395 | 60.08032442 | 95.55279619 | 0.992155723 | 166.4783804 | 0.247910561 | -31.22432063 |
| egl-9(n586)V | 0.02240163 | 0.141005634 | 0.003489125 | 0.009521084 | 0.622632585 | 0.001912007 | -0.384250551 | 81.22665558 | 129.7639087 | 2.141878214 | 312.6318811 | 0.468263817 | -46.62143555 |
| sem-4(ga82)I | 0.019252503 | 0.29630362 | 0.005060046 | 0.012829648 | 0.949590419 | 0.00278047 | -0.003217848 | 61.57194675 | 90.81767655 | 4.109674335 | 434.0601141 | 0.66640253 | -68.1499039 |
| ftn-2(ok404)I | 0.015294214 | 0.112499106 | 0.003405146 | 0.013079474 | 0.890272334 | 0.002342211 | -0.260134172 | 69.7492733 | 100.9917564 | 2.656490564 | 303.4363022 | 0.463824868 | -57.953907 |
| flp-1(yn2)IV | 0.027963466 | 0.235522324 | 0.003701808 | 0.009068915 | 0.548420089 | 0.002206148 | -0.336816208 | 74.84252824 | 117.1446128 | 2.43605557 | 306.9525399 | 0.465827576 | -41.03823517 |
| flp-10(ok2624)IV | 0.016011759 | 0.090827317 | 0.003378224 | 0.012023774 | 0.867360973 | 0.002043122 | -0.248393091 | 70.36211661 | 118.5391461 | 2.685658963 | 372.8245133 | 0.524810413 | -55.34923454 |
| flp-11(tm2706)X | 0.016848223 | 0.123346698 | 0.003598543 | 0.012072417 | 0.750308735 | 0.002232806 | -0.210090936 | 73.22733525 | 122.6618155 | 2.778123685 | 360.7923468 | 0.535103602 | -58.90054957 |
| flp-12(ok2409)X | 0.028395206 | 0.234235135 | 0.003902823 | 0.009984397 | 0.602606887 | 0.002403248 | -0.145612785 | 76.5180169 | 126.7939687 | 2.589856023 | 342.7455819 | 0.454782727 | -53.64144924 |
| flp-13(tm2427)IV | 0.027679337 | 0.175977266 | 0.00299653 | 0.009612864 | 0.649435455 | 0.001932749 | -0.267713277 | 76.25188479 | 112.3821478 | 2.737956327 | 367.3982332 | 0.516865593 | -55.77285112 |
| flp-16(ok3085)II | 0.023165053 | 0.177952956 | 0.003570545 | 0.011014835 | 0.668089685 | 0.002349696 | -0.212056126 | 75.5578202 | 123.9796906 | 2.926929173 | 361.8928395 | 0.541329513 | -56.04621514 |
| flp-17(ok3587)IV | 0.017643093 | 0.117457834 | 0.003401226 | 0.013851756 | 0.868168956 | 0.002278899 | -0.324634116 | 72.46864357 | 116.6418716 | 2.978536749 | 438.7540802 | 0.5854546 | -62.47171478 |
| flp-18(db99)X | 0.018457887 | 0.139705689 | 0.003064849 | 0.010127407 | 0.737493645 | 0.001603592 | -0.387046151 | 75.88937863 | 120.139174 | 2.322425766 | 362.2440858 | 0.491269889 | -58.61362934 |
| flp-19(ok2460)X | 0.016049298 | 0.180991853 | 0.00355375 | 0.013745004 | 1.00613627 | 0.00231218 | -0.358289478 | 70.51068824 | 117.282648 | 2.557158804 | 337.879529 | 0.487270285 | -81.268812 |
| flp-20(ok2964)X | 0.01936935 | 0.255539764 | 0.003041229 | 0.010318064 | 0.810779849 | 0.001890748 | -0.190621 | 79.25150276 | 114.4828699 | 2.574903127 | 416.8402192 | 0.46150583 | -74.05736895 |
| flp-21(ok889)V | 0.020257978 | 0.12139028 | 0.003555477 | 0.011136453 | 0.755341607 | 0.002168776 | -0.222284306 | 76.87350311 | 118.1983711 | 2.732495046 | 346.0665459 | 0.524948069 | -59.32102299 |
| flp-25(gk1016)III | 0.024425865 | 0.189944347 | 0.003403003 | 0.010290346 | 0.648588368 | 0.002111235 | -0.191666987 | 78.93763003 | 118.3548989 | 2.804079585 | 361.0769992 | 0.511996769 | -53.84858927 |
| flp-28(gk1075)X | 0.009775781 | 0.056533811 | 0.003216738 | 0.015572417 | 1.215655863 | 0.002137127 | -0.331269217 | 73.72519744 | 116.8601131 | 2.900614588 | 427.8760432 | 0.609529486 | -79.7259643 |
| flp-3(ok3265)X | 0.01093925 | 0.177529723 | 0.003475301 | 0.014084283 | 0.874991863 | 0.002340205 | -0.388553965 | 74.80499252 | 123.7643582 | 2.814358567 | 386.5146332 | 0.54552581 | -65.7646753 |
| flp-33(gk1038)I | 0.019472359 | 0.175393436 | 0.003026542 | 0.010647875 | 0.821611579 | 0.001848628 | -0.26100032 | 75.82398698 | 124.4153374 | 2.612447655 | 395.6594318 | 0.503654057 | -74.38399449 |
| flp-6(ok3056)V | 0.018854819 | 0.203756526 | 0.004049534 | 0.012458661 | 0.786850272 | 0.002472742 | -0.239817175 | 74.79869588 | 119.3477013 | 2.853594462 | 371.4477772 | 0.511949086 | -55.97000921 |
| flp-7(ok2625)X | 0.029953309 | 0.232599677 | 0.003479296 | 0.008673354 | 0.502644733 | 0.00229538 | -0.123021284 | 76.88357506 | 124.3467018 | 2.60494287 | 337.4410248 | 0.466026247 | -42.10787497 |
| flp-9(ok2730)IV | 0.018237124 | 0.173558684 | 0.00371759 | 0.012294959 | 0.811618593 | 0.002246184 | -0.311889916 | 71.08661571 | 124.1646105 | 2.579507647 | 336.0967952 | 0.501160471 | -53.72117414 |
| pdl-1(gk157)II | 0.015285779 | 0.093461624 | 0.003346118 | 0.013327381 | 1.010226391 | 0.002275115 | -0.222007365 | 72.42388859 | 119.2334504 | 2.769486003 | 401.6870818 | 0.539946079 | -67.55865535 |
| gsa-1(ce81)I | 0.004097787 | 0.125571833 | 0.003625456 | 0.020596187 | 2.09481214 | 0.002340949 | -0.038709502 | 75.51848602 | 77.1652174 | 3.170696497 | 358.7754555 | 0.676146485 | -143.4119454 |
| gar-2(ok520)III | 0.027737941 | 0.2279241 | 0.003000349 | 0.01007684 | 0.651367735 | 0.002081313 | 0.023514118 | 80.13371952 | 120.9080253 | 2.580913539 | 354.4262566 | 0.377578709 | -41.5193816 |
| gld-1(op236)I | 0.022390422 | 0.144738872 | 0.003410386 | 0.012520941 | 0.822791329 | 0.002450862 | -0.343831564 | 65.07388783 | 125.413764 | 2.41147203 | 348.1964302 | 0.482726196 | -64.13747788 |
| gly-2(gk204)I | 0.031971981 | 0.287045363 | 0.003842582 | 0.009429866 | 0.528806968 | 0.002549529 | -0.251275608 | 71.39234001 | 120.654118 | 2.500064834 | 354.8792173 | 0.445172048 | -50.5395773 |
| goa-1(sa734)I | 0.012136936 | 0.113277143 | 0.003197462 | 0.022365151 | 2.341545484 | 0.002239728 | -0.003254076 | 74.69710895 | 120.1469637 | 4.113453678 | 448.7347701 | 0.577515138 | -170.6017032 |
| gpa-16(ok2349)I | 0.008036246 | 0.052123326 | 0.002932856 | 0.015107257 | 1.419566835 | 0.002008976 | -0.293319879 | 82.90261358 | 132.8466193 | 2.736613059 | 412.3877355 | 0.575130533 | -119.5412462 |
| gpa-4(pk381)IV | 0.009391602 | 0.062005689 | 0.003420828 | 0.014314949 | 0.95503371 | 0.002165798 | -0.305626923 | 74.61279865 | 119.0741221 | 2.862710065 | 382.4307624 | 0.573417863 | -61.65805605 |
| gpa-1(pk15)V | 0.019838679 | 0.143896405 | 0.003367674 | 0.011621028 | 0.771410768 | 0.002144278 | -0.249708102 | 77.68273584 | 121.7125457 | 2.505072672 | 347.8513047 | 0.479281928 | -51.20371727 |
| gpa-10(pk362)V | 0.019265 | 0.168754036 | 0.003419878 | 0.011363179 | 0.755926013 | 0.002069133 | -0.273559961 | 77.38772635 | 127.6519866 | 1.940258915 | 287.1317589 | 0.391588419 | -54.9753681 |
| gpa-11(pk349)II | 0.013711456 | 0.078957225 | 0.00326783 | 0.013100766 | 0.990764451 | 0.002013454 | -0.145365277 | 78.46028036 | 126.4514081 | 2.384534087 | 349.2304042 | 0.480488873 | -76.14776387 |
| gpa-12(pk322)X | 0.020595668 | 0.137677321 | 0.003451334 | 0.011643769 | 0.750384834 | 0.002288595 | -0.213348266 | 78.70138411 | 126.8155846 | 2.530560955 | 309.5250224 | 0.471587279 | -49.28135933 |
| gpa-13(pk1270)V | 0.009892142 | 0.055114691 | 0.003452676 | 0.013298376 | 0.970901976 | 0.002070273 | -0.318214911 | 84.79052365 | 119.6672828 | 2.558713106 | 358.0719437 | 0.522267908 | -69.22175586 |
| gpa-14(pk347)I | 0.013139437 | 0.134552816 | 0.003405973 | 0.01551412 | 1.066168136 | 0.002289085 | -0.194572552 | 79.71433767 | 148.5090161 | 2.411130865 | 417.8994252 | 0.458516949 | -84.97325585 |
| gpa-15(pk477)I | 0.011002761 | 0.093372916 | 0.003165235 | 0.011589887 | 0.823539619 | 0.002200657 | -0.246922531 | 87.66353501 | 139.248916 | 2.3805297 | 323.3382224 | 0.457184925 | -63.35500218 |
| gpa-17(ok2334)III | 0.037047462 | 0.163908863 | 0.003162977 | 0.011395225 | 0.748025977 | 0.002217418 | -0.226571644 | 82.55614268 | 122.6062991 | 2.449904561 | 335.0972306 | 0.481457798 | -60.17159935 |
| gpa-2(pk16)V | 0.017554974 | 0.135605948 | 0.003126193 | 0.01048141 | 0.699795546 | 0.001972684 | -0.387104214 | 83.85727967 | 122.0973555 | 2.48376238 | 312.6357945 | 0.49579864 | -47.15506379 |
| gpa-3(pk35)V | 0.015992582 | 0.102693024 | 0.003628297 | 0.013354132 | 0.782294957 | 0.002405677 | -0.22311166 | 76.45328958 | 124.5561094 | 2.436261636 | 287.5161211 | 0.450303425 | -51.42631976 |
| gpa-5(pk376)X | 0.014318157 | 0.090882576 | 0.003102749 | 0.012410105 | 0.916157899 | 0.002052088 | -0.258867945 | 79.64903358 | 123.658232 | 2.637934354 | 373.84986 | 0.514871975 | -67.64780981 |
| gpa-6(pk480)X | 0.025492984 | 0.171922027 | 0.003484394 | 0.01040373 | 0.641019212 | 0.002319584 | -0.192958326 | 79.40349274 | 124.8391719 | 2.185812652 | 312.8066465 | 0.425647804 | -50.90989332 |
| gpa-7(pk610)IV | 0.0119759 | 0.137827146 | 0.003197514 | 0.012321787 | 0.965413807 | 0.001991049 | -0.130842274 | 86.28178324 | 141.1484567 | 2.310031973 | 327.816905 | 0.446854226 | -76.15920222 |
| gpa-8(pk345)V | 0.014878568 | 0.107781007 | 0.003226068 | 0.012472051 | 0.907338917 | 0.002088488 | -0.21412269 | 78.06605005 | 128.6518683 | 2.407750443 | 346.6929283 | 0.478261959 | -68.49332118 |
| hcf-1(ok559)IV | 0.033425998 | 0.238214801 | 0.003672228 | 0.009257437 | 0.529845662 | 0.002582705 | -0.392011081 | 66.14910761 | 123.8226424 | 2.230922954 | 303.7726499 | 0.447720103 | -43.56835102 |
| nhr-95(gk836)V | 0.01924203 | 0.115560732 | 0.003464952 | 0.011373194 | 0.739701652 | 0.002302471 | -0.28550074 | 70.12909741 | 127.6794199 | 2.446477206 | 369.8532026 | 0.487848517 | -56.53345793 |
| ins-11(tm1053)II | 0.014414978 | 0.123604042 | 0.003122898 | 0.011613659 | 0.924896341 | 0.002006039 | -0.246343951 | 74.40565395 | 132.951059 | 2.538453674 | 385.0848319 | 0.485585606 | -67.53017044 |
| ins-15(ok3444)II | 0.010402233 | 0.242096782 | 0.003824691 | 0.014408734 | 1.02925671 | 0.002463289 | -0.197744065 | 71.71103489 | 122.0319761 | 2.586612828 | 414.0488945 | 0.441136892 | -67.56153612 |
| ins-16(ok2919)III | 0.011124811 | 0.09301033 | 0.003223499 | 0.013232808 | 0.985473635 | 0.002013193 | -0.352557005 | 71.78745933 | 130.8240045 | 2.75651061 | 378.4236749 | 0.538979203 | -78.41164796 |
| ins-27(ok2474)I | 0.024707413 | 0.293152179 | 0.003485499 | 0.011145272 | 0.685188967 | 0.002305333 | -0.033612558 | 76.29772345 | 134.1345803 | 2.217086683 | 340.0628707 | 0.358771843 | -64.34773175 |
| ins-28(ok2722)I | 0.030968889 | 0.379488082 | 0.003504003 | 0.009699156 | 0.672139692 | 0.002249741 | -0.073504696 | 72.77881961 | 133.4876675 | 2.241551495 | 392.2895842 | 0.405789483 | -47.37329298 |
| ins-30(ok2343)I | 0.018912255 | 0.162154699 | 0.003006803 | 0.011146804 | 0.891543936 | 0.002002634 | -0.230330109 | 76.02432979 | 128.5087655 | 2.320917975 | 360.7217705 | 0.481007325 | -63.65095139 |
| ins-31(ok3543)II | 0.014318484 | 0.216031178 | 0.003456513 | 0.013638764 | 0.958398757 | 0.002400349 | -0.27292713 | 72.60975547 | 128.1085102 | 2.390809095 | 406.5532134 | 0.49401385 | -81.16228684 |
| jnk-1(gk7)IV | 0.030107187 | 0.354609651 | 0.004025181 | 0.010519956 | 0.593075868 | 0.002394014 | -0.08715219 | 69.21911818 | 163.2605783 | 2.488084897 | 343.674214 | 0.408243824 | -47.54618753 |

| rel_to_neck_ra | curvature_std_d | path_curvat | curvature_hea | rel_to_body_ra | rel_to_neck_ra | d_rel_to_body | width_head_b | curvature_mea | curvature_mid | d_rel_to_head | length_w_back | d_rel_to_neck | d_ang_vel_mi | d_curvature_s | curvature_std | width_tail_bas | [illegible] |
|---|---|---|---|---|---|---|---|---|---|---|---|---|---|---|---|---|---|
| -30.88441073 | 0.002087021 | 0.515530105 | 0.005275485 | 0.239924985 | -0.238996792 | -133.6626933 | 59.31803039 | 0.011520777 | 0.004598996 | -2.3194277 | 78.46128476 | -9.024991842 | 1.215914389 | 0.002198748 | 0.002361107 | 70.80096475 | [illegible] |
| -30.64321605 | 0.001814342 | 0.57865513 | 0.004919661 | 0.173833872 | -0.311367517 | -107.4164488 | 56.76571829 | 0.011442206 | 0.004155562 | -2.243452798 | 67.59054566 | -8.745527965 | 1.038645663 | 0.0020858 | 0.002076218 | 67.60152701 | [illegible] |
| -48.42143441 | 0.001388355 | 1.71842897 | 0.004532109 | 0.795343518 | 3.0488379 | -83.84773255 | 65.40021324 | 0.007240308 | 0.004394575 | -2.745266318 | 42.2916565 | -11.6706996 | 1.026327044 | 0.001429897 | 0.001727998 | 76.68685532 | [illegible] |
| -44.20219121 | 0.001364492 | 2.231196604 | 0.004511373 | 0.594266728 | 1.327410375 | -112.0882028 | 72.0682109 | 0.007680652 | 0.003423847 | -2.803219085 | 29.17341027 | -12.42097436 | 1.26929065 | 0.001490304 | 0.001681552 | 84.62694608 | [illegible] |
| -32.80184387 | 0.001032104 | 2.63066458 | 0.003323847 | 0.278157381 | 2.088640901 | -109.0193353 | 72.58209412 | 0.005739309 | 0.003318154 | -2.375675855 | 28.47222656 | -9.644277192 | 1.305407042 | 0.001584241 | 0.001263917 | 86.8306311 | [illegible] |
| -50.85837289 | 0.002065643 | 0.67395742 | 0.00442634 | 0.078046004 | 0.921561325 | -116.4478188 | 62.00474052 | 0.007838668 | 0.004379752 | -6.826206684 | 25.78149109 | -26.18268519 | 1.261190805 | 0.002295044 | 0.001933245 | 74.39314804 | [illegible] |
| -45.03643274 | 0.001119628 | 2.973135062 | 0.004267327 | 0.813542715 | 1.544034301 | -119.5535005 | 68.48934237 | 0.006159817 | 0.003704094 | -2.787441457 | 26.18763987 | -14.27020135 | 1.85047853 | 0.00139798 | 0.001346017 | 75.80730375 | [illegible] |
| -49.34680176 | 0.001094635 | 4.811555123 | 0.005083661 | 1.678098178 | 0.931759083 | -203.7266707 | 73.28674774 | 0.006645879 | 0.003672367 | -3.295718861 | 23.83565674 | -12.27723198 | 4.512131405 | 0.001411029 | 0.001625041 | 87.2354065 | [illegible] |
| -64.96261215 | 0.001174442 | 1.342981279 | 0.004543651 | 1.460678697 | 0.891955897 | -475.8077164 | 74.00198365 | 0.007610428 | 0.00348335 | -1.98208642 | 28.58932495 | -5.815431446 | 5.299910903 | 0.001146579 | 0.001423455 | 88.27399635 | [illegible] |
| -62.54043828 | 0.001773505 | 3.358172018 | 0.005307565 | 1.801014688 | 1.160573474 | -594.8043452 | 74.20961055 | 0.011133911 | 0.00495592 | -3.760589537 | 26.9272913 | -12.93606986 | 9.462635289 | 0.001748794 | 0.002349239 | 80.55867485 | [illegible] |
| -54.02872987 | 0.001266278 | 3.782140789 | 0.004330573 | 1.062900321 | 2.401655295 | -159.1032739 | 75.67920234 | 0.006818658 | 0.003715317 | -3.765997464 | 29.4971258 | -16.00038606 | 2.100358367 | 0.00159703 | 0.001691178 | 84.91489098 | [illegible] |
| -49.59897491 | 0.001196282 | 3.773338081 | 0.004658451 | 1.285957611 | 1.45164973 | -162.628872 | 76.36165833 | 0.00646178 | 0.003653702 | -2.952801489 | 29.78875732 | -11.44181478 | 2.341581591 | 0.001248676 | 0.001597415 | 88.51232076 | [illegible] |
| -45.81119796 | 0.001062355 | 2.940866432 | 0.004033856 | 0.888164663 | 1.463258704 | -146.2667485 | 84.92978758 | 0.006400282 | 0.003230632 | -2.934678266 | 29.79378554 | -13.52382889 | 1.76891378 | 0.001398934 | 0.001470876 | 97.77868787 | [illegible] |
| -47.26535047 | 0.001121318 | 3.583101902 | 0.004825086 | 1.03738924 | 0.628094224 | -170.2211472 | 74.89433736 | 0.006463892 | 0.003747774 | -2.87582234 | 23.86991093 | -12.27010689 | 2.520814121 | 0.00132159 | 0.001501129 | 86.43977172 | [illegible] |
| -33.23250111 | 0.001043277 | 3.424794555 | 0.004434497 | 0.633600845 | 0.133578795 | -144.6487247 | 69.01099873 | 0.005499232 | 0.003207053 | -2.182240687 | 22.29590098 | -8.744001805 | 2.06165417 | 0.001058504 | 0.001111188 | 76.32968299 | [illegible] |
| -47.64813747 | 0.00126402 | 3.634484058 | 0.004449665 | 1.135113217 | 1.023080845 | -200.7179423 | 71.24011065 | 0.006650902 | 0.003526625 | -3.083615941 | 24.46284817 | -12.63396597 | 2.736330377 | 0.001231475 | 0.001430814 | 83.51531286 | [illegible] |
| -42.818342 | 0.00160853 | 1.901729278 | 0.004600072 | 0.756755558 | 0.702251081 | -160.5713292 | 72.33583182 | 0.008767313 | 0.004354596 | -1.683646195 | 24.52839717 | -13.08469759 | 2.036344517 | 0.001165241 | 0.001858557 | 77.8773815 | [illegible] |
| -48.06590066 | 0.001412328 | 1.813135749 | 0.004828932 | 0.984729531 | -0.933721297 | -152.4040064 | 69.62285436 | 0.007881407 | 0.00381944 | -2.944428641 | 28.11861244 | -13.41935624 | 1.806213776 | 0.001518147 | 0.001463322 | 80.23838542 | [illegible] |
| -51.74356152 | 0.001307308 | 2.367004342 | 0.004396329 | 1.254485089 | 0.963369649 | -205.8969521 | 70.24271422 | 0.007453928 | 0.003902671 | -3.051186938 | 27.08205473 | -14.13803097 | 2.576395367 | 0.001291771 | 0.001605752 | 78.41455195 | [illegible] |
| -51.65994867 | 0.001292294 | 2.615593751 | 0.00439474 | 0.80267746 | 2.235402572 | -137.3271357 | 70.63010168 | 0.006583756 | 0.003710499 | -3.293058528 | 24.8953654 | -15.47316329 | 1.824829914 | 0.001389344 | 0.001540695 | 81.61584059 | [illegible] |
| -30.3510978 | 0.001813435 | 2.137372642 | 0.006742499 | -0.047954786 | 0.727310467 | -103.009843 | 54.12843583 | 0.010487349 | 0.005304413 | -0.935248687 | 25.76733676 | -7.242581806 | 1.396063582 | 0.001772524 | 0.002634782 | 55.39907976 | [illegible] |
| -54.90691328 | 0.001293493 | 2.810753122 | 0.004657071 | 0.764469441 | 2.671423182 | -112.1126757 | 72.53535843 | 0.007273287 | 0.003883946 | -3.633568034 | 28.64050293 | -19.67195082 | 1.227394044 | 0.001837453 | 0.00179068 | 81.74177458 | [illegible] |
| -57.67074204 | 0.001122171 | 3.63995949 | 0.004488949 | 1.35554729 | 1.657207549 | -197.4594048 | 74.52612286 | 0.006318337 | 0.003431861 | -3.941040421 | 29.99308014 | -15.82661052 | 2.354328699 | 0.001416923 | 0.001397232 | 86.61124497 | [illegible] |
| -47.46790844 | 0.001143083 | 2.645220211 | 0.004181633 | 0.977190011 | 1.858739564 | -130.9530869 | 75.55092699 | 0.006363046 | 0.003436547 | -2.958520298 | 27.85645322 | -13.32796771 | 1.55514021 | 0.001226726 | 0.001465666 | 86.40648415 | [illegible] |
| -57.72064939 | 0.001268357 | 3.654534641 | 0.0057618 | 0.813371306 | -0.097169599 | -130.1910433 | 70.58726392 | 0.006047777 | 0.003923654 | -3.322698322 | 29.88976353 | -19.767684 | 1.72410304 | 0.001561999 | 0.001517094 | 79.02790244 | [illegible] |
| -55.15237808 | 0.003915744 | 0.066999515 | 0.007927128 | -1.741261948 | -2.113255735 | -201.6176402 | 56.92069535 | 0.018042422 | 0.00821401 | -4.294645713 | 24.73885963 | -15.05613799 | 2.003117581 | 0.00378904 | 0.00328533 | 68.89323425 | [illegible] |
| -62.24511996 | 0.000986573 | 2.8729201 | 0.005017967 | 2.28729998 | 0.872680844 | -390.8549894 | 74.64464274 | 0.006583974 | 0.003142087 | -3.887017623 | 19.95547306 | -14.81130492 | 4.648207117 | 0.001347985 | 0.001267946 | 79.23706506 | [illegible] |
| -55.99153228 | 0.001280205 | 4.4690412 | 0.004473833 | 1.142455683 | 2.706226746 | -163.8608878 | 73.59431603 | 0.007062217 | 0.003900058 | -3.781787112 | 26.39330473 | -14.94810509 | 2.3714329 | 0.001460483 | 0.001822702 | 85.16975911 | [illegible] |
| -43.08406514 | 0.001218909 | 2.011233265 | 0.003608514 | 0.543007932 | 3.16451758 | -102.1034793 | 75.81404199 | 0.005273947 | 0.003255473 | -4.248158013 | 30.35077037 | -17.19137088 | 0.640748812 | 0.001965888 | 0.001513707 | 92.93046911 | [illegible] |
| -54.68322029 | 0.001172255 | 4.536212063 | 0.004567905 | 1.255044642 | 1.533853403 | -201.9104721 | 76.81969948 | 0.006577088 | 0.003677202 | -3.543041265 | 29.12438049 | -14.6523247 | 2.838685888 | 0.001427529 | 0.001531368 | 88.9099781 | [illegible] |
| -52.59700209 | 0.00118692 | 5.530092051 | 0.004512838 | 1.187149705 | 1.541270987 | -195.4136763 | 73.87083976 | 0.006699811 | 0.003801966 | -3.539797133 | 29.9476023 | -15.79580982 | 2.986980265 | 0.001558583 | 0.001714228 | 86.58021152 | [illegible] |
| -44.6920305 | 0.001022127 | 4.043842018 | 0.004489217 | 1.046470816 | 1.171954951 | -196.8651134 | 84.10256032 | 0.006119317 | 0.003388026 | -2.30647452 | 31.62275478 | -9.622460297 | 2.493303906 | 0.001348643 | 0.001535949 | 91.38811738 | [illegible] |
| -47.51926659 | 0.00116231 | 3.895385179 | 0.00424303 | 1.042079083 | 2.273036414 | -167.0999027 | 77.54343046 | 0.006008495 | 0.003652362 | -3.510864562 | 30.53697468 | -14.60471065 | 2.223131935 | 0.0015166 | 0.00149275 | 90.3899072 | [illegible] |

[illegible]

| width_tail_bas | d_curvature_m | ang_vel_tail_b | curvature_std_ | speed_hips_w_ | d_ang_vel_tail | d_rel_to_body_ | d_rel_to_body_ | d_curvature_md | rel_to_head_ | rel_to_neck_arr | rel_to_head_b | d_length_w_ba | curvature_hips | d_curvature_std | ang_vel_neck | d_speed_neck | d_rel_to_body_ | d_major_axis_ | d_curvature_st | d_path_curvat | d_rel_to_body_ | d_rel_to_hips_ | width_tail_base_w_forward_ |
|---|---|---|---|---|---|---|---|---|---|---|---|---|---|---|---|---|---|---|---|---|---|---|---|
| 58.63168303 | 0.00431055 | 0.443268048 | 0.001444649 | 58.45157656 | 1.33345422 | -209.2744191 | -108.5096462 | 0.00039425 | 47.533951 | 0.055162353 | -10.73101956 | -4.683041476 | 0.004978914 | 0.001570773 | 0.125159265 | -2.023371728 | -1.347645931 | -96.91120121 | 0.000386234 | 0.624916117 | 0.671800149 | -0.772444798 | 51.24629113 |
| 56.725112 | 0.004051242 | 0.405750555 | 0.001319689 | 57.97682973 | 1.166619051 | -192.6201437 | -104.5735985 | 0.000330187 | 47.54119981 | 0.053186355 | -10.39455276 | -4.036331225 | 0.004571291 | 0.001446062 | 0.120305341 | -1.912843189 | -0.308491027 | -85.81176186 | 0.000366147 | 0.818345573 | 0.578331297 | -0.636568994 | 49.52771071 |
| 68.78332902 | 0.003904172 | 0.295210555 | 0.001032161 | 69.0507679 | 0.729478094 | -188.9321518 | -72.4311142 | 0.000252499 | 51.47471237 | 0.043030583 | -11.16983843 | -2.360770464 | 0.003692468 | 0.00096172 | 0.108927831 | -2.266234636 | -0.846285642 | -91.98913193 | 0.000244605 | 2.287140131 | 0.559035927 | -0.785368353 | 62.51092339 |
| 75.67967077 | 0.004601262 | 0.35951555 | 0.000970662 | 82.88567895 | 1.090806218 | -219.6205632 | -94.45743253 | 0.000303307 | 59.39384651 | 0.05732179 | -11.62187463 | -3.330092559 | 0.002901941 | 0.001037552 | 0.127640557 | -2.427390352 | -0.969689968 | -102.6045607 | 0.000256403 | 2.761023058 | 0.750670067 | -0.808274698 | 69.53536533 |
| 80.01416504 | 0.003179676 | 0.366895939 | 0.000895149 | 81.02811706 | 0.980927837 | -231.3117169 | -90.77507904 | 0.000254714 | 44.14232475 | 0.04966737 | -10.15557743 | -2.979043219 | 0.00284693 | 0.000821528 | 0.110754316 | -1.597440028 | -0.517053455 | -97.45394593 | 0.000280443 | 3.251242714 | 0.65556438 | -0.749643085 | 74.27829071 |
| 66.60906067 | 0.004745841 | 0.214671442 | 0.00134121 | 48.25867519 | 1.004142636 | -309.6526291 | -94.37233849 | 0.000582683 | 76.30805244 | 0.116292223 | -14.34793496 | -0.609420099 | 0.004615592 | 0.001880602 | 0.179243344 | -5.991763884 | -0.507397693 | -124.2083267 | 0.000409046 | 1.054682559 | 0.743615267 | -1.506232697 | 61.39722691 |
| 69.63976733 | 0.004763127 | 0.381134984 | 0.00091908 | 86.30071115 | 1.289093524 | -266.3979111 | -94.17313273 | 0.000368391 | 55.68750429 | 0.056999088 | -10.40705363 | -3.328790692 | 0.003075337 | 0.001107917 | 0.16302077 | -2.812749644 | -2.539013733 | -107.2279139 | 0.000236297 | 3.447121633 | 0.97504746 | -1.081998677 | 65.99289672 |
| 75.24030151 | 0.007872984 | 0.682571113 | 0.001030014 | 119.8812637 | 3.737607574 | -472.8259277 | -145.7662613 | 0.000826416 | 54.60908203 | 0.147554341 | -13.2902813 | -5.725697422 | 0.003085414 | 0.001757799 | 0.271996123 | -9.411938287 | -4.318649149 | -139.0622269 | 0.000270447 | 6.274723816 | 2.472510266 | -2.24539733 | 68.47046356 |
| 78.76278687 | 0.011571818 | 1.092740923 | 0.000894261 | 194.8565331 | 4.818897247 | -659.4876099 | -261.2080536 | 0.000477924 | 64.97429943 | 0.078021517 | -16.28260755 | -8.278149604 | 0.002782991 | 0.001357141 | 0.510055192 | -5.18344 | -6.748421669 | -255.3523903 | 0.000187668 | 1.895428151 | 2.867776394 | -1.699466437 | 74.33069611 |
| 74.04909317 | 0.015047377 | 1.573075377 | 0.001194388 | 135.2952038 | 8.643810314 | -719.4841932 | -337.6134365 | 0.002285057 | 65.04647893 | 0.299605366 | -16.15048616 | -8.226445945 | 0.004024856 | 0.002697348 | 0.929487195 | -4.918388432 | -15.57295131 | -259.3263086 | 0.000308421 | 5.024025088 | 5.365765354 | -7.855637374 | 70.03856433 |
| 77.07067143 | 0.005233291 | 0.452924812 | 0.001016944 | 104.1841795 | 1.650984466 | -297.6459378 | -112.4278058 | 0.000369506 | 72.1223207 | 0.070656873 | -14.48242387 | -4.765520583 | 0.003090132 | 0.001207275 | 0.161431718 | -5.354737536 | -1.766280028 | -129.2618852 | 0.000261517 | 4.670895934 | 1.209267324 | -1.516708669 | 71.68349804 |
| 78.95797098 | 0.005768896 | 0.451521697 | 0.000998154 | 84.03390823 | 1.792019834 | -323.1929203 | -107.1231751 | 0.000602564 | 59.22199052 | 0.101982784 | -14.01531127 | -6.032003279 | 0.002994668 | 0.001344368 | 0.191912015 | -4.032334426 | -4.071299761 | -140.6095209 | 0.000216168 | 3.510034156 | 1.282311805 | -1.565640294 | 73.89827502 |
| 89.07648939 | 0.004322341 | 0.398077501 | 0.00097878 | 80.5747122 | 1.308153259 | -286.9115222 | -106.9855046 | 0.00044002 | 61.8867665 | 0.082192572 | -13.19675113 | -4.988136792 | 0.002679329 | 0.001092159 | 0.140869942 | -3.982899587 | -1.908821609 | -123.4447145 | 0.00022898 | 3.180205018 | 0.910856386 | -1.225871591 | 83.2797984 |
| 79.33483387 | 0.006063197 | 0.470290932 | 0.000949442 | 100.5670161 | 1.972331394 | -317.3226787 | -108.7107307 | 0.000707813 | 52.70712294 | 0.101582446 | -11.45679281 | -5.633986749 | 0.002990601 | 0.001400457 | 0.213173273 | -6.872171259 | -3.335418631 | -133.8927941 | 0.00022899 | 5.272094825 | 1.397642717 | -1.240625901 | 75.2583947 |
| 70.57709408 | 0.004978387 | 0.388506146 | 0.000918109 | 87.60323445 | 1.685186597 | -261.5639184 | -95.58615035 | 0.000473812 | 38.39829636 | 0.077139269 | -9.046777586 | -3.539503658 | 0.002580705 | 0.001104622 | 0.153122114 | -3.250874256 | -3.851505243 | -101.8302099 | 0.000186472 | 3.525136342 | 1.087456383 | -1.409108823 | 66.52657556 |
| 77.35098134 | 0.006119354 | 0.475039939 | 0.000941048 | 105.467429 | 2.264945401 | -370.3184894 | -103.4410721 | 0.000543442 | 57.47788686 | 0.101647588 | -12.77974498 | -4.828413397 | 0.002782477 | 0.001248829 | 0.214634821 | -2.962224815 | -5.671983118 | -154.4815916 | 0.000213147 | 4.209669735 | 1.513245097 | -1.312453001 | 72.96772766 |
| 71.89470828 | 0.004558779 | 0.414716036 | 0.000731129 | 76.05179638 | 1.549570949 | -274.3432903 | -110.3081419 | 0.000327426 | 51.8864253 | 0.057473507 | -10.80478476 | -5.156432231 | 0.00372227 | 0.001043126 | 0.140980943 | -1.649659385 | -1.700994399 | -124.7585893 | 0.000201305 | 2.918173953 | 1.081609991 | -1.363223055 | 67.08277409 |
| 73.55223876 | 0.005216863 | 0.326831187 | 0.001048185 | 77.07278986 | 1.380294357 | -310.4933841 | -100.7461733 | 0.000452556 | 55.59555465 | 0.079166823 | -12.1623113 | -5.227139815 | 0.00298426 | 0.001240636 | 0.190405258 | -3.868126461 | -1.604373153 | -132.5311969 | 0.000265981 | 2.268242409 | 1.025534554 | -1.176185945 | 68.32479477 |
| 72.91189781 | 0.005408478 | 0.483496179 | 0.000933223 | 104.3565159 | 1.983266941 | -314.5458474 | -115.5358403 | 0.000459662 | 59.01106541 | 0.075995071 | -12.09901439 | -3.542039282 | 0.003200702 | 0.00109804 | 0.211148786 | -3.464274769 | -2.46081285 | -146.3626069 | 0.000222856 | 3.817412072 | 1.431038858 | -1.714010744 | 69.12223713 |
| 73.96284199 | 0.004554035 | 0.365479646 | 0.000952853 | 92.09161314 | 1.415160465 | -268.8787791 | -89.07044601 | 0.000305104 | 63.82461294 | 0.065947211 | -12.40112309 | -2.584737097 | 0.002911934 | 0.000970717 | 0.143825297 | -2.941595467 | -2.048281404 | -117.9399042 | 0.000246791 | 3.082482385 | 1.028674214 | -1.065356293 | 70.00186793 |
| 50.80694008 | 0.004318544 | 0.351418461 | 0.00143924 | 56.29580714 | 1.04060001 | -152.8244133 | -82.60055941 | 0.000355789 | 39.83553314 | 0.048038443 | -9.153210184 | -3.173234609 | 0.004347895 | 0.001083914 | 0.107468245 | -1.519486306 | 0.897778011 | -71.55637238 | 0.000334672 | 3.496873069 | 0.743239193 | -0.354844666 | 46.66023029 |
| [illegible] | [illegible] | [illegible] | [illegible] | [illegible] | [illegible] | [illegible] | [illegible] | [illegible] | [illegible] | [illegible] | [illegible] | [illegible] | [illegible] | [illegible] | [illegible] | [illegible] | [illegible] | [illegible] | [illegible] | [illegible] | [illegible] | [illegible] | [illegible] |
